# Human bone marrow adipocytes drive prostate cancer bone metastasis progression via lipid-mediated induction of Angiopoietin-like 4

**DOI:** 10.64898/2025.12.16.694397

**Authors:** Marine Hernandez, Sauyeun Shin, David Rengel, Stéphanie Dauvillier, Nancy Geoffre, Carine Valle, Alessandro Taccini, Mohamed Moutahir, Justine Bertrand-Michel, Nicolas Reina, Catherine Muller, Camille Attané

**Affiliations:** Institut de Pharmacologie et de Biologie Structurale (IPBS), Université de Toulouse, CNRS, Toulouse, France; Equipe Labellisée Ligue Contre Le Cancer, Toulouse, France; MetaboHUB-MetaToul, National Infrastructure of Metabolomics and Fluxomics, Toulouse, 31077, France; Université de Toulouse, Inserm, I2MC, Toulouse, France; Université de Toulouse, Inserm, CRCT, Toulouse, France; Département de Chirurgie Orthopédique et Traumatologique, Hôpital Pierre-Paul Riquet, CHU de Toulouse, Toulouse, France; Laboratoire AMIS, UMR 5288 CNRS, Université de Toulouse, Toulouse, France

**Keywords:** Prostate cancer metastases, primary human bone marrow adipocytes, metabolic crosstalk, lipids, migration, ANGPTL4

## Abstract

Bone is the main metastatic site in advanced prostate cancer (PCa) and contains bone marrow adipocytes (BMAds), which account for more than 70% of adult bone marrow. However, their role in the progression of PCa metastases remains poorly understood. Herein, we developed a physiologically relevant 3D culture model using primary human BMAds from red hematopoietic rich-areas (rBMAds) and we showed that rBMAds engage in metabolic crosstalk with PCa cells. Specifically, rBMAds release free fatty acids (FFAs) through a non-canonical lipolytic pathway and these FFAs are taken up by PCa cells inducing a transcriptional reprogramming that promotes motility. Among the responsive genes, Angiopoietin like 4 (ANGPTL4) is the most upregulated via a peroxisome proliferator-activated receptor γ-dependent mechanism, and its silencing abolishes the rBMAd-driven migratory and invasive phenotype. Clinically, elevated ANGPTL4 expression is associated with poor overall survival specifically in PCa bone metastases, is maintained during metastatic progression, and is indicative of aggressive disease. Together, our findings uncover a novel adipocyte-tumor interaction and identify ANGPTL4 as a key mediator and potential therapeutic target in bone-metastatic PCa.

## Introduction

Prostate cancer (PCa) is the second most commonly diagnosed cancer and the fifth leading cause of cancer-related death among men worldwide^1,2^. While many patients benefit from curative therapies, approximately one-third progress to advanced metastatic disease, which remains a major life-limiting condition^3,4^. Among distant metastases, the bone is the predominant site of colonization, with nearly 80% of advanced PCa patients developing bone lesions, predominantly osteoblastic metastases^4–6^. The bone microenvironment supports metastatic establishment, progression and resistance to therapy, involving a complex interplay among osteoblasts, osteoclasts, stromal and immune cells and the extracellular matrix^7,8^. Despite their proximity to PCa bone metastases, bone marrow adipocytes (BMAds) remain one of the most abundant yet understudied components of the metastatic bone niche^9^. As metabolically active and secretory cells, BMAds are known to regulate normal hematopoiesis, bone remodeling and onset of hematologic malignancies, primarily based on evidence from preclinical models^9,10^. However, their role in PCa bone metastasis remains poorly defined.

Growing evidence, including our own, strongly supports the idea that adipocyte–tumor cell interactions, particularly metabolic exchanges, play a pivotal role in cancer, including PCa progression.

At the primary tumor site, periprostatic adipose tissue (PPAT), a white adipose tissue (WAT) surrounding the prostate^11^, contributes to PCa progression through paracrine mechanisms^12^. While soluble factors such as chemokines and pro-inflammatory cytokines from PPAT have been shown to enhance PCa aggressiveness and local dissemination^12,13^, a more specific and increasingly recognized mechanism involves the direct transfer of free fatty acids (FFAs) from adipocytes to tumor cells^14^. Tumor cells induce FFA release from adipocytes by activating lipolysis, a major metabolic pathway in adipocytes that involves the hydrolysis of triglycerides (TGs) into FFAs and glycerol^15,16^. In turn, these FFAs are taken up by cancer cells and exploited for their own advantage^16^. This form of metabolic symbiosis is now recognized across multiple cancer types including melanoma, ovarian, breast, and colon cancers^16^. Depending on the tumor type, FFAs taken up by cancer cells can act directly as substrates for a range of pathways, including fatty acid β-oxidation (FAO), signaling and transcriptional remodeling or be converted into neutral lipids (mainly, TGs) and stored in lipid droplets (LDs) to avoid lipotoxicity^16^. In PCa, at the primary tumor site, we have shown that FFA uptake induces expression of the pro-oxidant enzyme NADPH oxidase 5 (NOX5), triggering signaling cascades that promote tumor invasion^14^. Therefore, the fate of lipids in tumor cells depends both on the tumor type and the nature of the proximal adipose depot. Understanding the mechanisms and consequences of lipid transfer from adipocytes to tumor cells is essential in the context of PCa metastasis in lipid-rich environments such as the bone marrow. The dynamic interplay between tumor cells and adipocytes represents not only a critical axis in cancer progression but also a potential therapeutic target that remains largely underexplored in the field of PCa metastasis.

Accumulating evidence indicates that BMAds exhibit specific metabolic and functional properties that differ markedly from other “classical” adipocytes^9,10^. Two distinct adipocyte populations exist in rodents: constitutive bone marrow adipocytes (cBMAds), located in areas devoid of hematopoietic cells and resistant to metabolic cues, and regulatory BMAds (rBMAds), interspersed within hematopoietic niches and responsive to energy stress and other signals, likely via lipolysis^9,10^ at least in rodents. Using femoral bone marrow aspirate of patients undergoing hip replacement surgery^17^, we have demonstrated that in humans, both types of adipocytes lack lipolytic activity^18,19^. Furthermore, we recently demonstrated that rBMAds, frequently found in proximity to metastatic lesions^20^, are anucleate but maintain metabolic and secretory functions that support hematopoiesis *in vitro*^19^. In both acute myeloid and lymphoblastic leukemias, which occur in areas of active hematopoiesis, a decrease in the number and size of rBMAds is observed, highlighting that dynamic remodeling of rBMAds also occurs in humans^19^. One very important point that needs to be underlined from our studies is that the metabolic specificity of human primary BMAds, particularly their altered lipolysis, is not recapitulated in *in vitro*–differentiated adipocytes, derived from preadipocyte cell lines or human bone marrow mesenchymal stem cells (BM-MSCs)^18^. Cocultures of *in vitro* differentiated adipocytes and PCa cell lines have demonstrated that lipid transfer occurs^21^, driven by adipocytes lipolysis^22,23^, leading to increased invasion, enhanced survival, and chemoresistance of PCa cells^22,24^. However, as stated below, since these models fail to recapitulate the metabolic characteristics of primary BMAds^18^, these results should be interpreted with caution. The use of primary human rBMAds, which is essential for investigating their involvement in cancer progression, has been limited to date by technical challenges associated with their isolation, anatomical localization, and the difficulty of maintaining them in culture for *in vitro* studies^25^.

To overcome these limitations, we developed a three-dimensional (3D) culture model of primary human rBMAds isolated from the femoral bone marrow of patients undergoing hip surgery. This model preserves rBMAds viability and metabolic function and reveals a previously unrecognized metabolic crosstalk in which rBMAds release FFAs via a non-canonical lipolytic mechanism. These FFAs are taken up by PCa cells, inducing a transcriptional program that promotes motility and invasion. We further identify angiopoietin-like 4 (ANGPTL4) as a peroxisome proliferator-activated receptor γ (PPARγ)-dependent effector of FFA signaling. Extending our previous findings that rBMAds promote PCa bone homing^26^, we show here that they also enhance PCa cell motility and invasion. Together, these data suggest that rBMAds facilitate both bone colonization and subsequent metastatic dissemination and identify ANGPTL4 as a key driver of the migratory phenotype in PCa, positioning it as a potential therapeutic target in advanced PCa.

## Materials and Methods

### Collection of adipose tissue samples and adipocyte isolation

BMAT and subcutaneous adipose tissue (SCAT) were harvested from patients undergoing hip surgery at the Orthopedic Surgery and Traumatology Department of the Hospital Pierre Paul Riquet (Toulouse, France) as previously reported^18^. All patients gave informed consent and the samples were obtained according to national ethics committee rules (authorization DC-2019-3784). For all experiments, samples were obtained from 185 men and 31 women (mean age: 66 years and mean body mass index (BMI): 24.26 kg/m^2^). All patient information’s are summarized in Table S1. Woman samples were used only for the development of rBMAd 3D culture model, lipolysis experiments and for the quantification of FFA and glycerol release by rBMAds cultured in BSA medium.

After obtention, the samples were immediately placed in 37°C pre-warmed KRBHA (Krebs Ringer Buffer HEPES Albumin buffer) corresponding to Krebs Ringer buffer (Sigma-Aldrich) supplemented with 100mM HEPES (Sigma-Aldrich) and 0.5% FFA-free bovine serum albumin (BSA) (Sigma Aldrich) at pH 7.4 and rapidly transferred to the laboratory (within 1h) where they were processed.

Periprostatic adipose tissue (PPAT) obtained from patients undergoing radical prostatectomy were transported to the laboratory within 1h in 10mL of DMEM (Thermo Fisher Scientific, Courtaboeuf, France)^11^. Periprostatic adipocytes, subcutaneous adipocytes (SCAds) from SCAT and rBMAds from “red” BMAT were isolated as we previously described^11,19^. Briefly, PPAT and SCAT were digested with collagenase (250 UI/ mL) diluted in calcium and magnesium free PBS supplemented with 2% FA-free BSA during 30 minutes at 37°C under constant shaking and samples were filtered with 200µm cell strainers. For BMAT, “red” BMAT was directly filtered with 100µm cell strainer and floating rBMAds were collected. SCAds and rBMAds were washed several times with KRBHA until a clear suspension was obtained. For rBMAds, a centrifugation at 300 g for 5 minutes at room temperature (RT) was performed to remove cell debris and obtain pure cell suspension.

### 3D culture and preparation of conditioned medium (CM) of rBMAds

Fibrin gels of SCAds and rBMAds were performed as previously described^27^. Briefly, 70µL of fibrinogen at 18mg/mL (Sigma Aldrich #F8630-5G) in NaCl 0.9% (4.2mg/mL final concentration) were mixed with 1.5µL or 7µL of aprotinin (Sigma Aldrich #A1153) at 5mg/mL (25µg/mL and 110µg/mL final concentration for 3 days and 5 days of coculture respectively) and 30µL of PBS-CaCl_2_ in 2mL eppendorf tube. One hundred microliters of SCAds or rBMAds (corresponding to approximatively 1.3×10^5^ SCAds and 2.3×10^5^ rBMAds) were added and homogenized quickly with 100µL of thrombin 25UI/mL (Sigma Aldrich #T6634-1KU) in PBS-CaCl_2_ (8.3UI/mL final concentration). After gel polymerization, gels were transferred in 12-well plates, containing 2.5mL of medium, to be cultivated alone or with PCa cell lines.

To validate the 3D culture model, a suspension of 50µL of rBMAds were included in fibrin gel and fixed with paraformaldehyde 3.7% during 20 minutes. Cells were then stained with BODIPY® 493/503 (for neutral lipids) at 1µg/mL. For each adipocyte sample, at least 3 z-stacks were acquired with LSM710 confocal microscope (Zeiss). Cell diameter measurement was performed manually using ImageJ/Fiji® software (Image J, Bethesda, MD, USA). For the quantification of adiponectin secretion overtime, rBMAds gel was incubated during 3h in RPMI without FBS at day 1, day 2 and day 3 of culture at 37°C in a humidified atmosphere with 5% CO_2_. Medium were cautiously collected and centrifuged at 500g for 5 minutes at RT to eliminate cell debris. The quantification of adiponectin was realized with Human DuoSet ELISA kits from R&D Systems (#DY1065-05) according to the manufacturer’s instruction.

To performed conditioned medium (CM) of rBMAds, fibrin gels were incubated in 1mL of RPMI (ThermoFisher Scientific #11879020) without FBS and supplemented with 5mM of glucose and 0.5% BSA (Sigma Aldrich). After 3 days of incubation, at 37°C in a humidified atmosphere with 5% CO_2_, CM were collected, centrifuged at 300g for 5 minutes at RT to remove cellular debris. CM were stored at -20°C until treatment of PCa cell lines. Delipidated CM (dCM) were prepared with Cleanascite™ lipid removal reagent (Gentaur, #0337-X2555-10) at ratio 1:1. Samples were gently shaken for 20 minutes, followed by 10 minutes of centrifugation at 5000g and the supernatant was used as dCM.

### Cell line culture and coculture

The human PCa cell lines, PC3 (ATCC CRL-1435), LNCaP (ATCC CRL-1740) and Du145 (ATCC HTB-81) were used in this study and derived from bone, lymph node and brain metastasis respectively. Leukemia cell lines U937 control and knock-out (KO) for CD36 were provided by Dr G. Bossis, IGMM, Montpellier, France. U937 cells KO for CD36 were generated by lentiviral Crispr/Cas9 system with a CD36-targeting single guide RNA (sgRNA) (5’AGAAAAGAACTGCAATACCTGG 3’). Control cells were generated using a non-targeting sgRNA (5’CACCGACGGAGGCTAAGCGTCGCAA 3’). PC3 and LNCaP were cultivated in RPMI medium (ThermoFisher Scientific # 11879020) supplemented with 5mM glucose (Sigma Aldrich), 5% FBS (Foetal Bovine Serum) and 1% Penicillin–Streptomycin (P/S). Du145, U937 and U937 KO CD36 cells were cultivated in RPMI GlutaMAX™ medium (ThermoFisher Scientific #61870044) supplemented with 10% FBS and 1% P/S. Cells were grown at 37°C, in a humidified atmosphere with 5% CO_2_ and used within 2 months after resuscitation of frozen aliquots. The cells were tested every month by polymerase chain reaction for mycoplasma contamination.

For coculture experiments, PCa cell lines were seeded at 10 000 cells/well for PC3 and 30 000 cells/well for Du145 and LNCaP in 12-well plates. For confocal microscopy, PCa cell lines were seeded on a coverslip. One day after seeding, medium was replaced by 2.5mL of fresh medium and rBMAds gel were added in a well and float above tumor cells. For the direct transfer of fluorescent palmitate (BODIPY™ FL-C16, Fisher #D3821), rBMAds suspension was loaded during 4 hours with 5µM of BODIPY™ FL-C16 prior to be embedded in fibrin gel and used for coculture with PCa cells.

### Lipolysis assay

Lipolysis activity was performed as we previously described^18,19^. Briefly, SCAds and rBMAds in suspension were incubated for 3 hours at 37°C in KRBHA with or without isoproterenol 1µM (Sigma Aldrich, #I6504) or forskolin 10µM (Sigma Aldrich, #F6886) to evaluate stimulated and basal lipolysis. For extended lipolysis assays, SCAds and rBMAds were incubated at 37°C for 3, 18, or 24 h in RPMI supplemented with 5 mM glucose and 0.5% BSA, with or without 1 µM isoproterenol. For experiments with lipase inhibitors, suspension of isolated SCAds and rBMAds were preincubated 1 hour at 37°C with or without 100µM orlistat (Sigma Aldrich #O4139) or 100µM paraoxon ethyl (Sigma Aldrich #D9286) before adding isoproterenol 1µM. Moreover, rBMAds cultured alone or in co-culture with PC3 cells were treated or not with either 100 µM orlistat or 100 µM paraoxon for 24 h. At the end of the incubation, media were collected to measure glycerol and FFA release using commercial kits (Sigma F6428 and Wako diagnostic NEFA-HR, respectively). Absorbance was read at a wavelength of 550nm and 540 nm for FFA and glycerol respectively according the manufacturer’s instructions.

### FFA release and neutral lipid quantification

PC3 alone (NC), rBMAds alone (rBMAds NC) or PC3 cultivated with rBMAds (COC) were incubated during 72 hours in 12-well plates in medium containing 5% FBS or 0.5% BSA depending on the experiment. Medium were then centrifuged for 5 minutes at 300g at RT to remove any cellular debris. FFA and glycerol released into medium were measured as described above. For FFA species quantification, lipids were extracted from the medium of rBMAds NC, PC3 NC and COC and analyzed as we previously described^19^.

For evaluation of TG content, tumors cells cultivated or not with rBMAds were lyzed in TE 1X buffer (10 mM Tris-HCl, 1 mM EDTA, pH 7.5) containing 0.1% Triton and TG content was quantified using a Triglyceride reagent (Sigma Aldrich) according the manufacturer’s instructions. The results were normalized by the concentration of protein evaluated by BCA Assay (Biorad). For neutral lipid quantification of PC3 NC and COC, lipids were extracted according to Bligh and Dyer^28^ in CH_2_Cl_2_:MeOH :water (2.5:2.5:2, v/v/v) in the presence of the internal standards. Organic phase was dried and dissolved in 20 µL of ethyl acetate. 1 µL of the lipid extract was analyzed by gas chromatography flam ionization detector on a GC TRACE 1300 Thermo Electron system using an RTX-5 column (Restek). Oven temperature was programmed from 210°C to 350°C at a rate of 5°C/min and the carrier gas was hydrogen (5ml/min). The injector and the detector were at 315°C and 345°C respectively. Finally, peak detection, integration and relative quantitative analysis were done using Chromeleon 7.3.1 (Thermo Scientific).

### Cell line treatments

For exogenous FFA treatment, PCa cell lines were seeded at 10 000 cells/well for PC3 and 30 000 cells/well for Du145 and LNCaP in 12 well-plates and at 2000 cells/well for PC3 in 96 well-plates. One day after seeding, cells were treated with 100µM palmitate (Cayman #29558), 100µM oleate (Sigma Aldrich #O3008), 100µM linoleate (Sigma Aldrich #L9530), or with the mix of 3 FFAs (100µM palmitate/100µM oleate/100µM linoleate) during 3 days.

For PPARγ inverse agonist treatment, PC3 and Du145 were seeded at 10 000 cells/well and 30 000 cells/well in 12 well-plates respectively. Three days after seeding, cells were preincubated during 1h with T0070907 10µM (Cayman #10026) followed by exogenous FFA treatment for 24h at the same concentration mentioned above or by coculture with rBMAds for 24h.

### Boyden chamber migration and invasion assay

After coculture, CM treatment or exogenous FFA treatment, 50 000 cells, were seeded in 8.0µm Transwells (Corning, #353097) placed in 24-well plates for migration assay. For invasion assay, Matrigel (Corning, #356231) was diluted 1:80 and 150 000 cells were seeded on the top of Matrigel contained in 8.0µm Transwells (Corning, #353182) placed in 12-well plates. The lower chamber was filled with RPMI (ThermoFisher Scientific) without serum (used as control) or containing 10% FBS. After 24h of migration or invasion, non-migrated cells were removed with a cotton swab except for a control assay to evaluate the total number of cells at the end of the experiment. Migratory cells were fixed in methanol and stained with Toluidine Blue 1% supplemented with 0.1M borax (Sigma). Transwell membranes were cut and incubated in a lysis buffer (Tris HCl 6.25mM pH 6.8, Glycerol 10%, SDS 2% and β-mercaptoethanol 5%). Absorbance was measured at 570nm and the percentage of migration and invasion is determined with migrating cells and the total number of cells.

### Scratch assay

After coculture or exogenous FFA treatment for 72 hours, cells are confluent and a wound was realized with a tip for 12 well-plate (coculture condition) or with IncuCyte® WoundMaker Tool for 96-well plate (treatment with exogeneous FFA). Medium was changed to remove cellular debris. Plates were imaged during 24h, every hour, with IncuCyte® live-cell imaging system. Wound closure was determined with IncuCyte® software for 96-well plate and with ImageJ/Fiji® plugin *Wound Healing Size Tool* for 12 well-plates. Wound healing was normalized to the initial wound.

### Proliferation assay

PC3, Du145, LNCaP NC and COC were counted with Malassez hemocytometer at 24, 48 and 72 hours of coculture. Proliferation was also evaluated by BrdU assay for PC3. Briefly, cells were incubated 24h with BrdU and the proliferation is evaluated according to the manufacturer’s protocol (Cell Signaling, #6813) each day during 72 hours.

### Short interfering RNA (siRNA) transfection

PC3 and Du145 were transiently transfected with ANGPTL4 siRNA pool (final concentration 25nmol/L) using ON-TARGETplus SMARTpool (Thermo Scientific Dharmacon) or ON-TARGETplus nontargeting pool used as a control (Thermo Scientific Dharmacon). Transfection was done according to the manufacturer’s instructions with Lipofectamine RNAiMAX (Invitrogen Life Technologies) followed by 24h of coculture with rBMAds. After coculture, a second transfection with siRNA was done to maintain ANGPTL4 silencing during migration and invasion assays. Gene knockdown efficiency, migration and invasion were then evaluated. The sequences of the four siRNA against ANGPTL4 are 5’ GGAGACAUCCUUCCGCAAA 3’; 5’GAUGGGAUCUGCACUGUAA 3’; 54GAAAUUAGGUCUGGGGAUA 3’ and 5’GCAGGAAGGACCUCUAGAA 3’. The sequence of the four siRNA against SLC27A are 5’ AAGACCAUCAGGCGCGAUA 3’ ; 5’ GGCUGAGACUGACGGGUUU 3’ ; 5’ GGUUCUGGGACGAUUGUAU 3’ ; 5’UCGCAUGGCACUAGGCAAU 3’.

### RNA extraction and quantitative real-time PCR

PCa cell lines NC and COC and U937 ctl and U937 KO CD36 were trypsinized and centrifuged for 5 minutes at 800g at 4°C and washed 3 times with PBS. RNA extraction was performed on pellets using the RNeasy® Mini kit columns (Qiagen) following the manufacturer’s instructions. Gene expression was quantified by quantitative real-time PCR (ANGPTL4: Hs01101123_g1; CD36: Hs00354519_m1, SLC27A1: Hs01587911_m1; SLC27A2: Hs00186324_m1; SLC27A3: Hs00225680_m1; SLC27A4: Hs00192700_m1; SLC27A5: Hs00202073_m1) with TaqMan primers with a CFX96 real time PCR system (BioRad). Results were normalized with 18S rRNA (Hs99999901_s1).

### RNA sequencing (RNA-seq)

RNA was extracted as described above. RNA was quantified by Qubit RNA kit (Thermofischer Scientific) and qualified with the HS RNA kit for the Fragment Analyzer (Agilent Technologies). The mRNAseq libraries were prepared with the Illumina Stranded mRNA Prep Ligation kit using an input of 500ng of high-quality RNA (RQN =10) following manufacturer’s instructions. The libraries were profiled with the HS NGS kit for the Fragment Analyzer and quantified by qPCR using the KAPA library quantification kit (Roche Diagnostics), pooled and sequenced on the Illumina NextSeq550 High Output flow cell 150 cycles. The dual-indexed sequencing parameters were 74, 10, 10, 74 cycles (read 1, index i7, index i5, read 2). Sequencer output files have been demultiplexed with the Illumina’s software bcl2fastq. Reads were mapped to the annotated Homo sapiens genome (GRCh38v97) using bowtie2 aligner (v2.4.4)^29^. RSEM software (v1.3.3)^30^ was used to quantify the gene expression. Statistical analysis and visualization were carried out in R working environment^31^ using the RStudio interface^32^. Data wrangling and visualization were carried out using the Tidyverse meta-package^33^. RNA-seq data analysis was performed using the edgeR package^34^. Genes with a sufficient number of reads were considered as differentially expressed when their Bonferroni-corrected p-value obtained from Quasi-Likelihood F-Test was inferior to 0.05. Gene Ontology analysis were completed using the clusterProfiler package^35^. GO terms were declared as significantly enriched when the p-value of the enrichment test fell below 0.01. R source codes are available upon request. Differentially expressed genes belonging in GO terms (i) hosting fewer than 500 genes, (ii) presenting « migration » or « motility » keywords on their name and (iii) being significantly enriched with a q-value smaller than 0.05, where considered as cell migration-related differentially expressed genes. Interactions among those differentially expressed genes, as well as those with ANGPTL4 were set using the STRINGdb library in R^36^, using a score threshold of 600. Nodes (i.e. differentially expressed genes) and edges (i.e. interactions) where then loaded into Cytoscape^37^ to build the shown network.

### Western Blot

Proteins were extracted from PC3 previously transfected with siCT and siSLC27A4 with lysis buffer (20mM TrisHCl, 150mM NaCl, 1mM EDTA, 1% Triton X100, 2.5mM sodium pyrophosphate, 1mM β glycerophosphate, 1mM Na3VO4, antiproteases 1X). Twenty micrograms of proteins were reduced with modified Laemmli buffer for 5min at 95°C, loaded on 4%–10% gradient SDS-PAGE gel (Biorad) and transferred to nitrocellulose membrane. Membrane was blocked with 5% skimmed milk in TBS (20mM Tris, 150mM NaCl) and incubated with rabbit anti-FATP4 (1:1000, abcam, #200353,) and mouse anti-actin (1:1000, Sigma, #A5441,). The membrane was washed with TBS complemented with 0.1% Tween-20 and incubated with HRP conjugated secondary antibodies (1:5000, Santa Cruz Biotechnology). The immunoreactive protein bands were revealed by ECL prime western blotting detection reagent (Ammersham^TM^) and detected with Odyssey imaging system (LI-COR Biosciences). Signal intensities were normalized to β-Actin.

### ELISA ANGPTL4

Culture medium and PCa cells were collected after 24 and 72 hours of culture of rBMAds NC, PCa NC and COC with or without siANGPTL4 transfection. The quantification of intracellular ANGPTL4 and ANGPTL4 secretion was performed with Human ANGPTL4 ELISA kit (HUES01556, Ozyme) according to the manufacturer’s protocol.

### Confocal microscopy on PCa cell lines

After coculture or exogenous FFA treatment, PCa cells were fixed with paraformaldehyde 3.7% during 20 minutes at RT. Cells were then stained with BODIPY® 493/503 (for neutral lipids) at 2.5 ng/mL, with rhodamine-phalloidin 1:1000 (for actin cytoskeleton) (Abcam) and with DAPI (for nucleus) at 1µM (Thermo Fisher Scientific) as previously described^27^. Fluorescence images were acquired with a confocal microscope (LSM710, Zeiss International, Germany) and analyzed with ImageJ/Fiji® software.

### Fluorescence quantification

For neutral lipid quantification after coculture, PC3 siCT and PC3 siSLC27A4 were stained with Bodipy 4µg/mL for 1h, trypsinized and centrifuged for 5 minutes at 800g and washed one time with PBS. For Bodipy FLC-16 uptake during coculture, PC3 were trypsinized and centrifuged for 5 minutes at 800g and washed one time with PBS. Fluorescence was quantified using a BD Accuri® C6 flow cytometer. The results were analyzed using FlowJo™ v10.8 Software (BD Life Sciences). For kinetic of palmitate and oleate uptake, fluorescence was measured with FLx800 Fluorescence™ Microplate Reader. Fluorescence values were normalized to the fluorescence measured at time 0.

### ANGPTL4 expression in human databases

ANGPTL4 expression levels were compared between primary prostate tumors and bone metastatic using available Metastatic Prostate Adenocarcinoma datasets (*SU2C/PCF Dream Team*^38^) and the *Prostate Cancer MDA PCa PDX* (*MD Anderson*^39^ cohorts available on cBioportal (http://www.cbioportal.org). ANGPTL4 expression levels were compared between metastatic sites (bone, lung, liver, lymph node) using three independent cohorts from the *Metastatic Prostate Adenocarcinoma* (*SU*2*C*/*PCF Dream Team*, *PNAS* 2019^38^) available on cBioportal (http://www.cbioportal.org), as well database under the accession number GSE118435, GSE126078 available on PCaDB (http://bioinfo.jialab-ucr.org/PCaDB/). Differences in ANGPTL4 expression between tumor sites were assessed with linear or with mixed linear models using, respectively, the stats package or the lme4 package in R^40^. R source codes are available upon request.

### Survival analysis

Survival analyses were performed using available PCa datasets. RNA expression and clinical data were obtained from The Cancer Genome Atlas database (TCGA) containing a total of 459 primary prostate tumors. Data from bone metastatic samples were retrieved from the *Metastatic Prostate Adenocarcinoma* (*SU*2*C*/*PCF Dream Team*, *PNAS* 2019^38^) cohort available on cBioportal (http://www.cbioportal.org) which includes 26 bone metastases PCa patients. For each dataset, we stratified patients according to ANGPTL4 expression with the upper and lower tertiles (33% lowest and 33% highest expression). Overall survival and disease-free survival were analyses and p-value were generated by using log-rank test.

### Statistical analyses

Data correspond to the mean ± standard error of the mean (S.E.M) of independent experiments. Statistical analysis was carried out using the Student’s t test, one-way ANOVA, two-way ANOVA or Wilcoxon test depending on the experiments (details in figure legends) after testing data normality by Shapiro-Wilk test. The Tukey or Sidak post hoc test correction was applied for one-way ANONVA and two-way ANOVA. *, **, *** and **** in the figures refer to statistical probabilities (P) of <0.05, <0.01, <0.001 and <0.0001, respectively. GraphPad Prism 10.6 software was used for all the statistical analyses.

## Results and discussion

### Transfer of FFA is observed between rBMAds and PCa cells in a 3D co-culture model

To investigate the potential metabolic crosstalk between primary human rBMAds and PCa cells, we developed a 3D co-culture model that maintains rBMAds viability and function over time. Due to their buoyancy, rBMAds lyse rapidly in 2D culture, as other adipocytes. To overcome this limitation, we adapted a 3D fibrin-matrix model previously developed for mammary adipocytes^27^. This model provides the dual advantage of preserving adipocyte viability for 3–5 days and enabling co-culture with tumor cells without compromising the integrity of the matrix^27^. Primary rBMAds isolated from BMAT were embedded in a 3D fibrin matrix. The morphology and size of the adipocytes were monitored for up to 5 days using BODIPY 493/503 staining (Supplementary Fig. 1A-B). Confocal microscopy revealed that rBMAds were homogeneously distributed within the matrix and maintained their spherical morphology (Supplementary Fig. 1A) and size (Supplementary Fig. 1B) over time. Adiponectin secretion also remained stable (Supplementary Fig. 1C). Together, these results demonstrate that the 3D fibrin matrix effectively maintains rBMAds morphology, size and function over time.

To assess metabolic interactions between rBMAds and PCa cells, PC3 cells were co-cultured (COC) or not (NC) with rBMAds for 72 hours. Unlike PC3 NC, PC3 COC accumulated numerous neutral lipid-containing LDs over time, visible as early as 24 hours post-co-culture and displayed morphological modifications (Fig. 1A). This phenotype was replicated in two additional PCa cell lines, Du145 and LNCaP (Supplementary Fig. 1D). Lipidomic profiling of PC3 cells after 72 hours of co-culture revealed a marked increase in TG content, while cholesteryl ester (CE) levels showed a modest increase, and free cholesterol (FC) levels remained unchanged (Fig. 1B). These findings were corroborated in the Du145 and LNCaP cell lines (Supplementary Fig. 1E).

**Figure 1:**
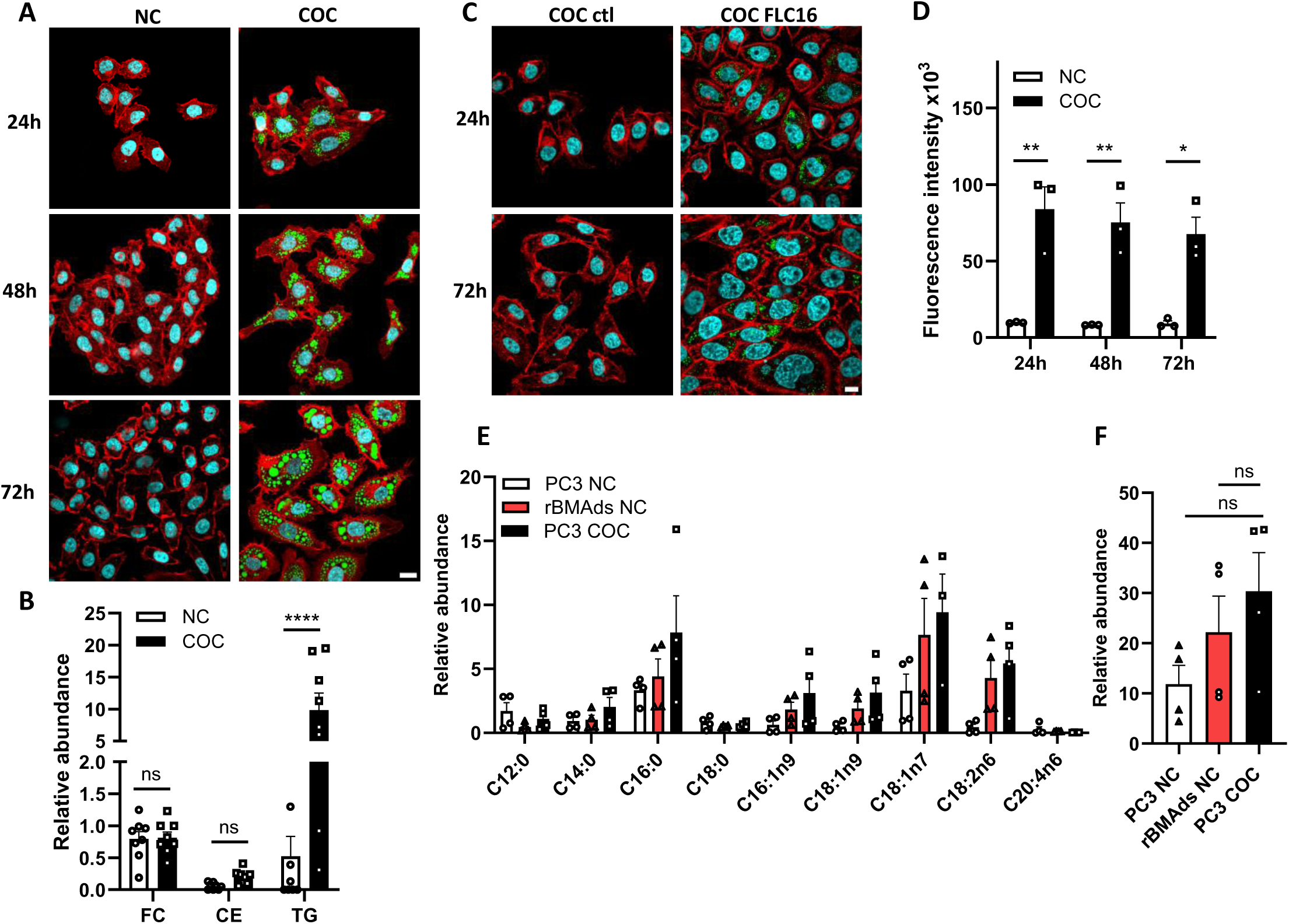
Transfer of FFAs is observed between rBMAds and PCa cells in a 3D co-culture model. **A.** Representative images of PC3 cultivated alone (NC) or with rBMAds (COC) during 24h, 48h or 72h. PC3 NC and COC were stained with rhodamine-phalloidin (red) for F-Actin, DAPI (cyan) for nuclei and Bodipy 493/503 (green) for neutral lipids. Scale bar, 20µm. **B.** Relative abundance of free cholesterol (FC), cholesterol ester (CE) and triglycerides (TG) in PC3 NC and COC after 72 hours of culture (n=8). **C.** Representative images at 24h and 72h of culture of PC3 COC previously loaded with Bodipy FLC16 (COC FLC16, in green) or without (COC ctl). Cells were stained with rhodamine-phalloidin (red) for F-Actin and DAPI (cyan) for nuclei. Scale bar, 10µm. **D.** Fluorescence intensity of Bodipy FLC16 at 24h, 48h and 72h of culture of PC3 NC or COC with rBMAds previously loaded with Bodipy FLC16 (n=3). **E.** Relative abundance of FFA species contained in culture medium of rBMAds alone (rBMAds NC), PC3 NC or PC3 COC after 72 hours of culture (n=4). **F.** Relative abundance of total FFA contained in culture medium of rBMAds alone (rBMAds NC), PC3 NC and PC3 COC after 72 hours of culture (n=4). Bar plot represent mean ± SEM, non- significant (ns), *P < 0.05, **P < 0.01, ****P < 0.0001; (B, D, E) were analyzed by two-way ANOVA with post hoc Sidak’s multiple comparisons test, (F) were analyzed by one-way ANOVA with post hoc Tukey’s multiple comparisons test.

To directly trace FFA transfer from rBMAds to tumor cells, we used a pulse-chase assay previously established by our group^41^. rBMAds were preloaded with BODIPY FL-C16, a fluorescently labeled saturated FA (palmitate), and then co-cultured with PC3 for up to 72 hours. Fluorescent LDs increased in PC3 between 24 and 72 hours (Fig. 1C), indicating FFA transfer, which was further confirmed by flow cytometry (Fig. 1D). These findings demonstrate that FFAs released by 3D-cultured rBMAds are efficiently taken up by PCa cells and subsequently re-esterified into TGs. To further characterize the FFA species released by rBMAds, we quantified the FFA composition in the culture medium collected from rBMAds cultured alone, PC3 NC and PC3 COC (Fig 1E). We demonstrated that, under both basal conditions and during co-culture with cancer cells, rBMAds predominantly release oleate (C18:1n9), palmitate (C16:0), and linoleate (C18:2n6) (Fig. 1E-F). This FFA profile mirrors their abundance in rBMAds^19^, where these three FFAs together account for approximately 85% of total TGs.

To better understand how FFA are taken up in PCa cells, we quantified, in PC3, Du145 and LNCaP cells, the expression of several FFA transmembrane transporters such as CD36 and FATP family members (SLC27A1-5 gene). CD36 expression was undetectable at both the mRNA and protein levels in all three PCa cell lines. These experimental results were validated using the U937 leukemic cell line as a positive control and CD36-knockout U937 cells as a negative control (Supplementary Fig. 1F–G). Among the FATP family members, FATP4 (encoded by *SLC*27*A*4) was consistently the most highly expressed transporter in all three PCa cell lines (Supplementary Fig. 1H). Based on its predominant expression, FATP4 was silenced in PC3 cells using siRNA (Supplementary Fig. 1I–J). FATP4 depletion modestly reduced the uptake of exogenous fluorescent oleate (Supplementary Fig. 1K) but had no detectable effect on the uptake of fluorescent palmitate (Supplementary Fig. 1L), suggesting that FATP4 contributes to the uptake of specific FFA species. Importantly, FATP4 knockdown did not affect lipid transfer in the co-culture model (Supplementary Fig. 1M), in which cancer cells are exposed to the complex mixture of FFAs released by rBMAds. Collectively, these findings indicate that FATP4 is dispensable for the transfer of rBMAd-derived lipids to PCa cells under our experimental conditions. These results suggest that additional mechanisms are likely to contribute to the uptake of adipocyte-derived FFAs, potentially including passive diffusion or extracellular vesicle-mediated lipid transfer.

In summary, these results demonstrate for the first time a direct metabolic crosstalk between primary human rBMAds and PCa cells. Notably, despite their absence of lipolytic activity upon β-adrenergic stimulation^19^, rBMAds can release FFAs via a mechanism yet to be elucidated.

### rBMAds exhibit non-canonical lipolysis with uncoupled FFA and glycerol release

We previously demonstrated that both human cBMAds and rBMAds lack canonical lipolytic activity^18,19^. Canonical lipolysis involves catecholamine-mediated stimulation of TG hydrolysis, through sequential activation of three lipases, and release of glycerol and FFAs^15^ (Fig. 2A). To test their lipolytic activity, isolated rBMAds and SCAds (used as controls) were treated with isoproterenol (ISO, a β-adrenergic agonist) or forskolin (FK, an adenylyl cyclase activator) for 3 hours. As previously reported^19^, rBMAds released minimal glycerol at baseline compared to SCAds, and this release was not enhanced by ISO or FK treatment (Fig. 2B). Moreover, while basal FFA release was comparable between rBMAds and SCAds, only SCAds exhibited a significant increase in FFA release upon lipolytic stimulation (Fig. 2B). To exclude the possibility that rBMAds display slower lipolytic kinetics, lipolytic stimulation was extended to 18 and 24 h. These experiments confirmed that rBMAds do not undergo complete lipolysis under either basal or stimulated conditions (Supplementary Fig. 2A). Interestingly, FFA release by rBMAds was not accompanied by a proportional increase in glycerol release (Fig. 2B and Supplementary Fig. 2A), suggesting that FFAs may be released, at least in part, through a non-canonical mechanism. To further investigate this hypothesis, we extended the incubation time to 72 hours in the presence of 0.5% BSA, which stabilizes both FFAs and glycerol by preventing their degradation (Fig. 2C). In this condition, FFAs were released at similar level between SCAds and rBMAds. In contrast, glycerol release was significantly lower in rBMAds compared to SCAds. SCAds exhibited a FFA:glycerol ratio of ∼3:1, consistent with complete TG hydrolysis, while rBMAds showed a strikingly elevated ratio of ∼25:1, indicating substantial FFA release uncoupled from glycerol liberation (Fig. 2D).

**Figure 2:**
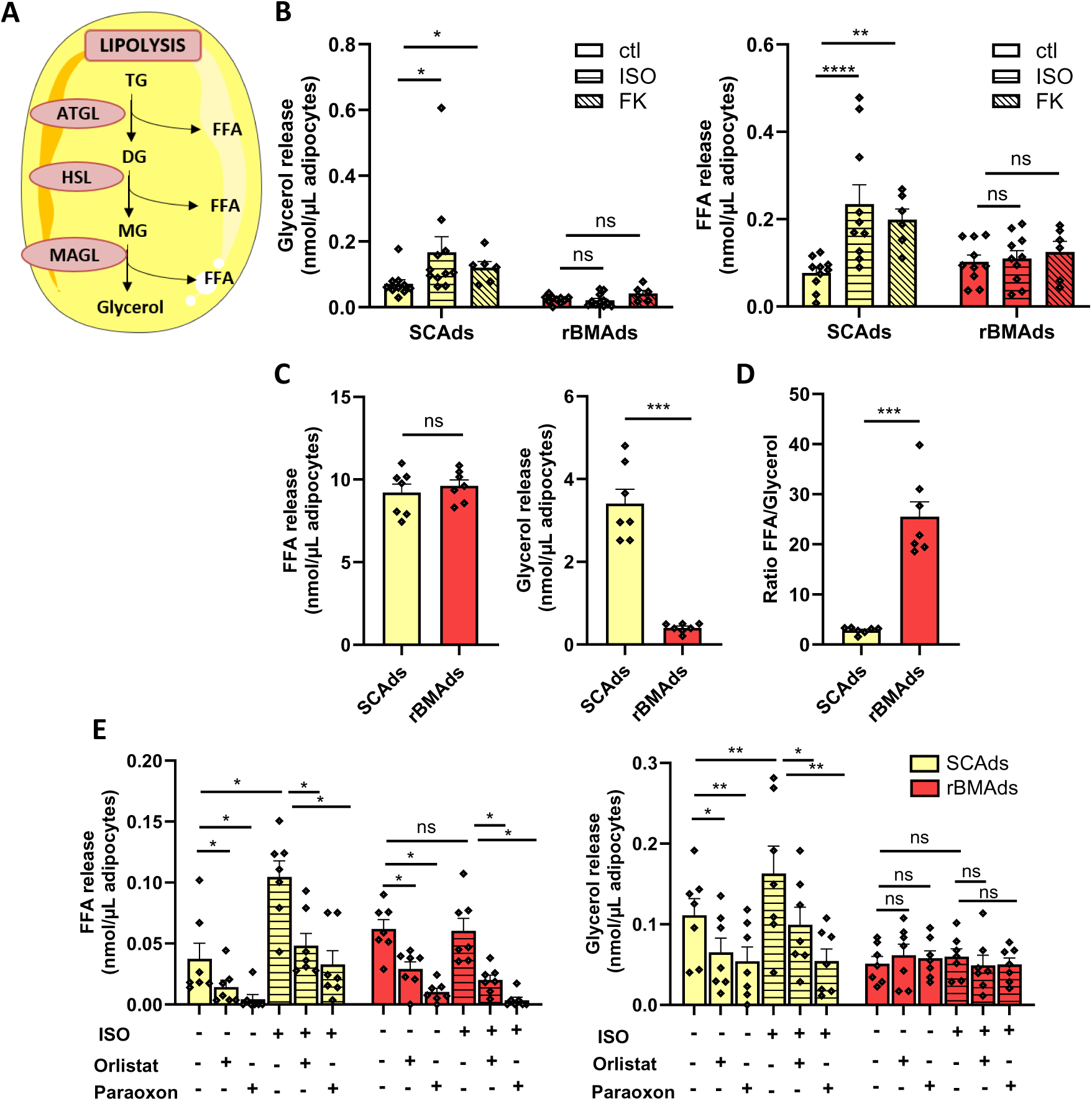
rBMAds exhibit non canonical lipolysis with uncoupled FFA and glycerol release. **A.** Scheme representing lipolysis pathway in “classical” adipocytes. TG: Triglycerides, DG: Diglycerides, MG: Monoglycerides, FFA: Free Fatty Acid, ATGL: Adipose Triglyceride Lipase, HSL: Hormone Sensitive Lipase, MAGL: Monoacylglycerol Lipase. **B.** Glycerol and FFA released by subcutaneous adipocytes (SCAds) and rBMAds at basal level (ctl, n=10) or after 3 hours of lipolysis stimulation with 1µM isoproterenol (ISO, n=10) or 10µM Forskolin (FK, n=6). **C.** FFA and glycerol released by SCAds and rBMAds during 72h of culture in medium containing 0.5% Bovine serum albumin (BSA) (n=7). **D.** Ratio of FFA/Glycerol released by SCAds and rBMAds after 72h of culture in medium containing 0.5% BSA (n=7). **E.** FFA and glycerol released by SCAds and rBMAds at basal level or after 3 hours of treatment with 100µM orlistat or 100µM paraoxon ethyl with or without isoproterenol (ISO) stimulation (n=7). Bar plot represent mean ± SEM, ns non- significant, *P < 0.05, **P < 0.01, ***P < 0.001, ****P < 0.0001; (B) was analyzed by two-way ANOVA with post hoc Sidak’s multiple comparisons test, (C-D), were analyzed by paired Student’s t-test, (E) was analyzed by Wilcoxon test.

Given that rBMAds contain over 90% of TG^18,19^, the observed FFA release is likely derived from TG hydrolysis. rBMAds express Adipose Triglyceride Lipase (ATGL), the first enzyme in the lipolytic cascade^42^. Mice with BMAd-specific deletion of ATGL, show enlarged rBMAds implicating ATGL in TG breakdown^43^. Since human ATGL inhibitors are unavailable and rBMAds lack nuclei, preventing genetic manipulation^19^, we tested the effect of pan-lipase inhibitors (orlistat and paraoxon ethyl)^44–46^ on FFA and glycerol release under basal and stimulated conditions (Fig. 2E) or during coculture with PCa cells (Supplementary Fig. 2B). As expected, both inhibitors reduced FFA and glycerol release in SCAds under all conditions (Fig. 2E and Supplementary Fig. 2B). In contrast, rBMAds showed significant reduction in FFA release with no changes in glycerol levels (Fig. 2E and Supplementary Fig. 2B), reinforcing the notion of incomplete lipolysis and absence of full TG hydrolysis in these cells. In conclusion, these findings reveal that rBMAds exhibit a non-canonical lipolytic pathway, potentially implicating ATGL, characterized by selective FFA release uncoupled from glycerol liberation, distinguishing them functionally from “classical” adipocytes.

### rBMAds reprogram PCa cells toward a pro-migratory and invasive phenotype

To investigate the impact of rBMAds on PCa progression, we performed RNA-seq on five matched PC3 NC and COC. Unsupervised multivariate analysis clearly partitioned the PC3 cells based on their culture condition (Fig. 3A) with the two first components explaining 39% and 20.1% of the total variance. Of the 15 686 genes, 749 were significantly upregulated and 443 were significantly downregulated in response to coculture (Fig. 3B). Gene Ontology (GO) analysis of differentially expressed genes revealed a strong enrichment in migration-associated processes among upregulated genes. Notably, pathways related to cell migration and motility, cell adhesion, including dynamic remodeling of cell-cell interactions and regulation of locomotion were among the most significantly induced (Fig. 3C, Table 1). In contrast, downregulated genes were predominantly associated with the regulation of cell population proliferation including both pro and anti-proliferative processes (Fig. 3C, Table 1). Given the strong induction of migration-related pathways, we next assessed PC3 cell migration using Boyden chamber and scratch assay following coculture. As shown in Fig. 3D, PC3 cells previously cocultured with rBMAds exhibited enhanced migration toward serum-containing medium compared to control cells. Similarly, in the scratch assay, wound closure was significantly accelerated in PC3 COC, indicating increased migratory capacity (Fig. 3E). This pro-migratory effect was also confirmed in Du145 and LNCaP cells (Supplementary Fig. 3A-B). In addition to increased migration, coculture with rBMAds promoted invasion of PC3 cells (Fig. 3F), as well as Du145 and LNCaP cells, through Matrigel-coated Boyden chambers (Supplementary Fig. 3C-D). Consistent with these findings, cocultured PCa cells displayed a more scattered morphology and developed multiple elongated protrusions, as revealed by actin staining (Fig. 3G, Supplementary Fig. 3E) compatible with the establishment of a pro-migratory phenotype. Finally, we did not observe any significant changes in cell number or BrdU incorporation after 72 hours of coculture in PC3 cells (Fig. 3H-I), nor in other PCa cell lines (Du145 and LNCaP) (Supplementary Fig. 3F-G).

**Figure 3:**
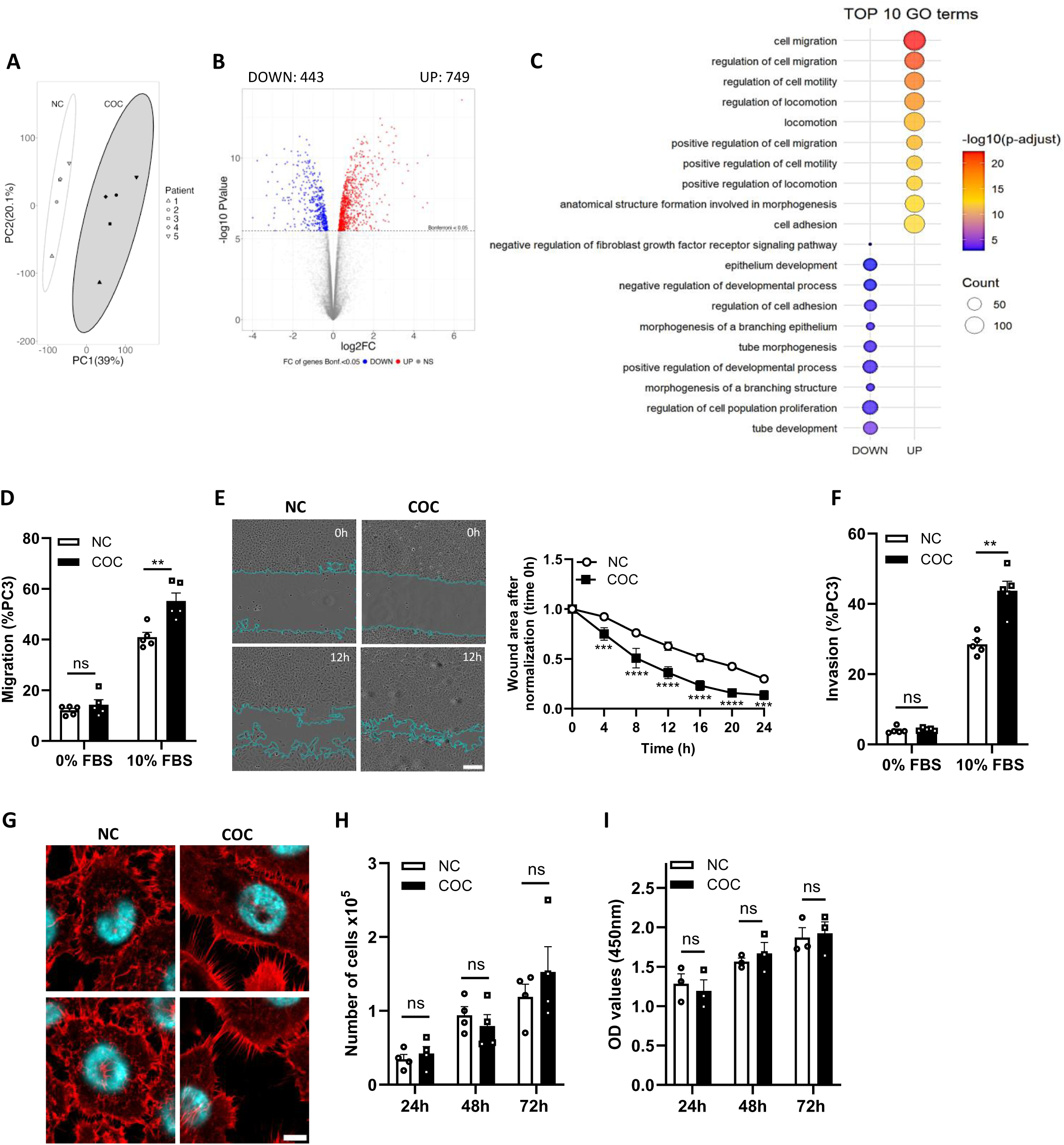
rBMAds reprogram PCa cells toward a pro-migratory and invasive phenotype. **A.** Principal Component Analysis of PC3 NC and COC during 72h (n=5). **B.** Volcano plot of genes detected by RNAseq with downregulated and upregulated genes by coculture represented in blue and in red respectively. **C.** Balloon plot representing the top 10 most downregulated and upregulated Gene Ontology (GO) terms in PC3 COC. **D.** Boyden chamber assay on PC3 NC and COC after 5 days of culture (n=5). **E.** Scratch assay on PC3 NC and COC after 5 days of culture. Images at the beginning (0h) and 12h after the wound are represented. Scale bar, 100µm. The wound was normalized to time 0 (when the scratch was performed) and followed during 24h (n=4). **F**. Invasion assay on PC3 NC and COC after 5 days of culture (n=5). **G.** Representative images of PC3 NC and COC during 72h and stained with rhodamine-phalloidin (red) for F-Actin and DAPI (cyan) for nuclei. Scale bar, 10µm. **H.** Number of PC3 NC and COC after 24h, 48h and 72h of culture (n=4). **I**. OD values of BrdU incorporated at 24h, 48h and 72h into PC3 NC and COC (n=3). Bar plot and curve represent mean ± SEM, ns non- significant, **P < 0.01, ***P < 0.001, ****P < 0.0001; (D-F, H, I) were analyzed by two-way ANOVA with post hoc Sidak’s multiple comparisons test.

**Table 1:**
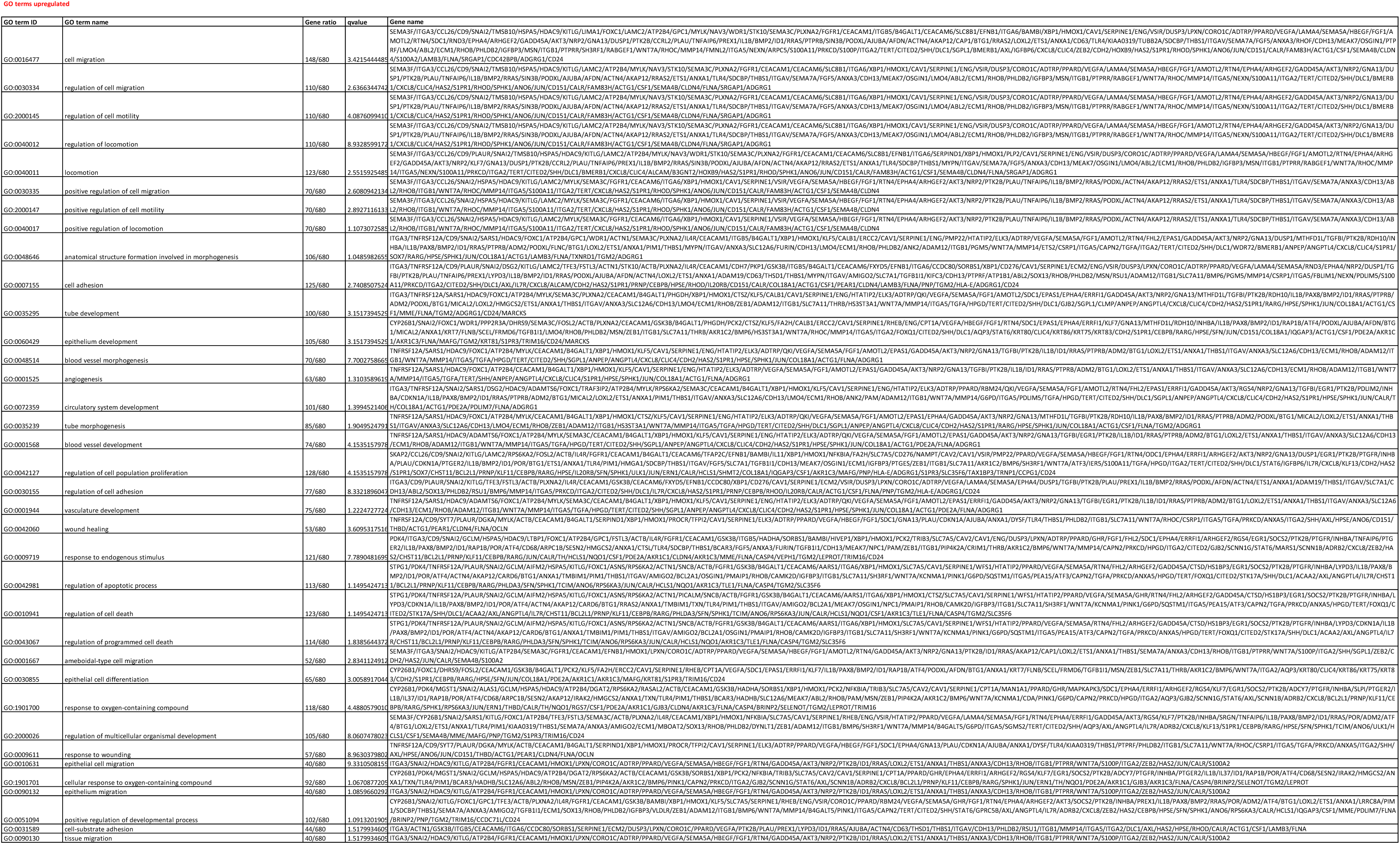
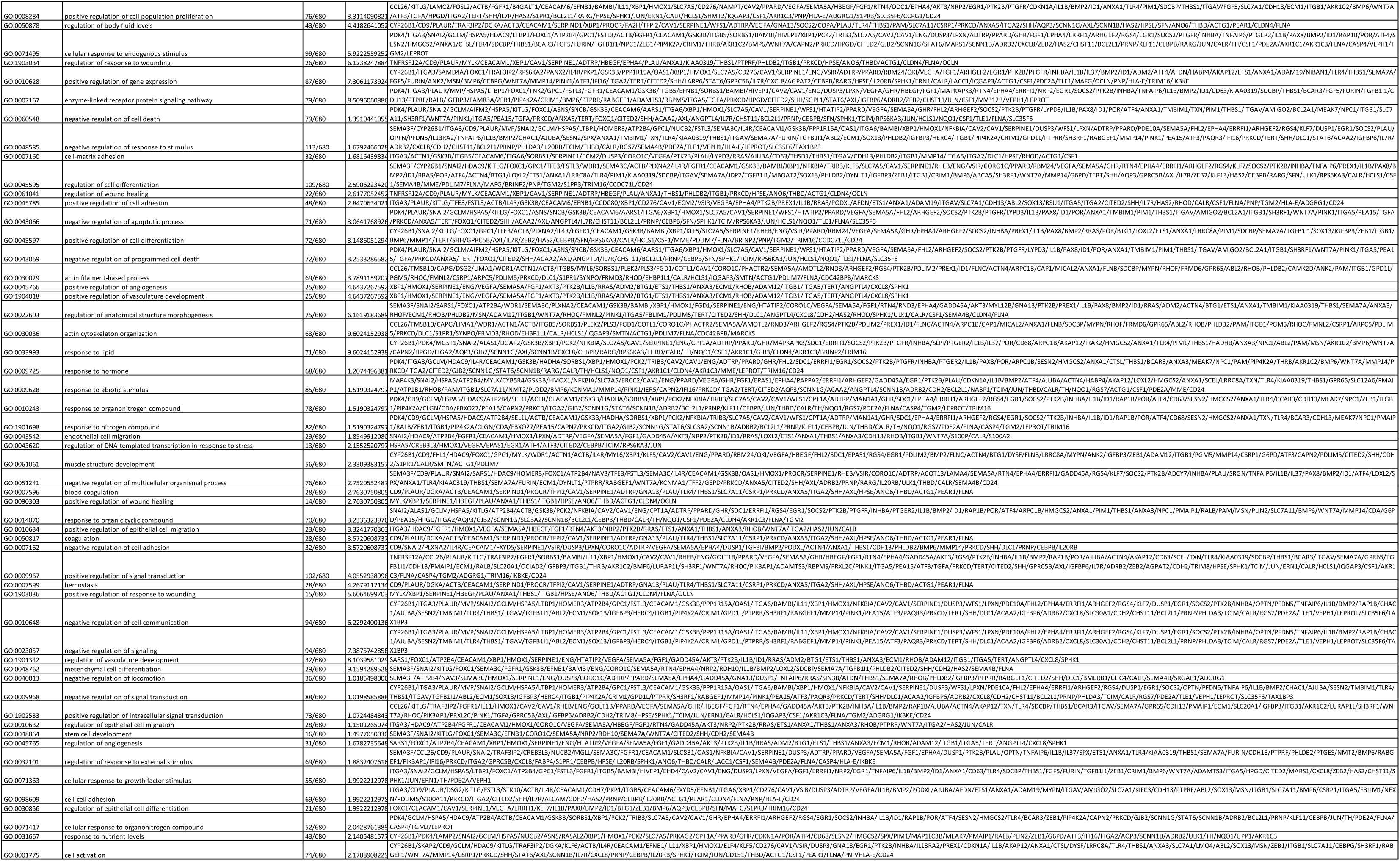
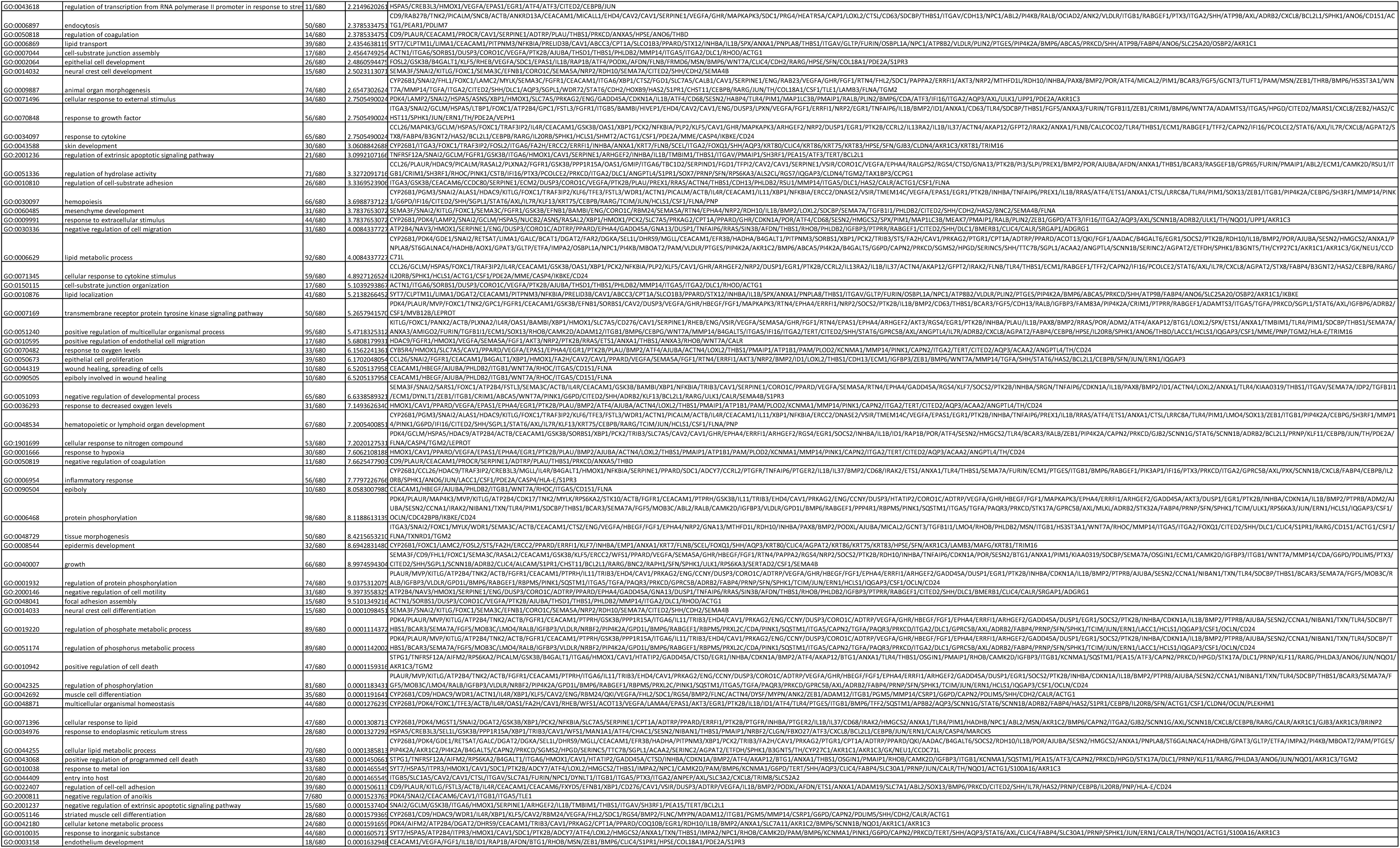
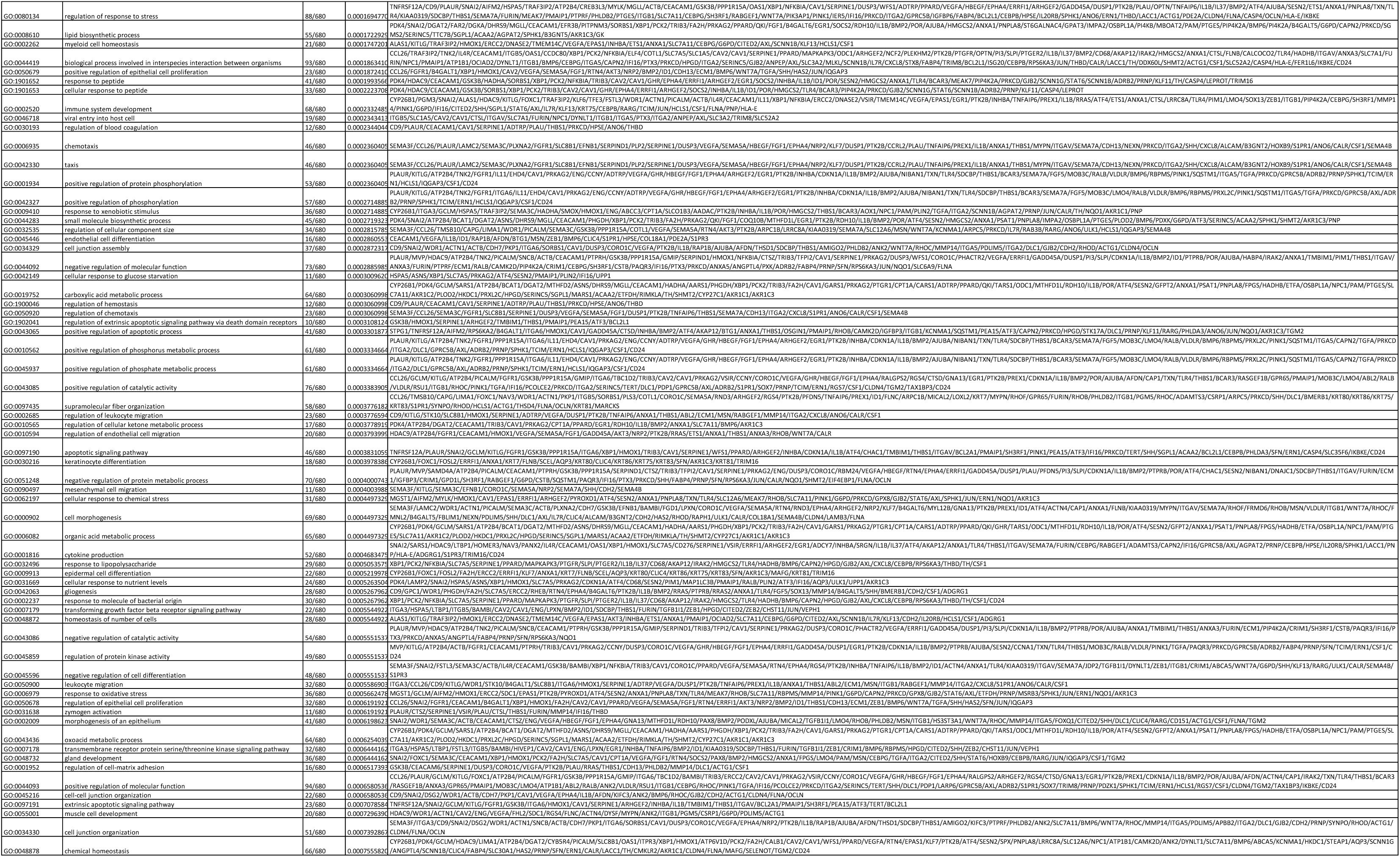
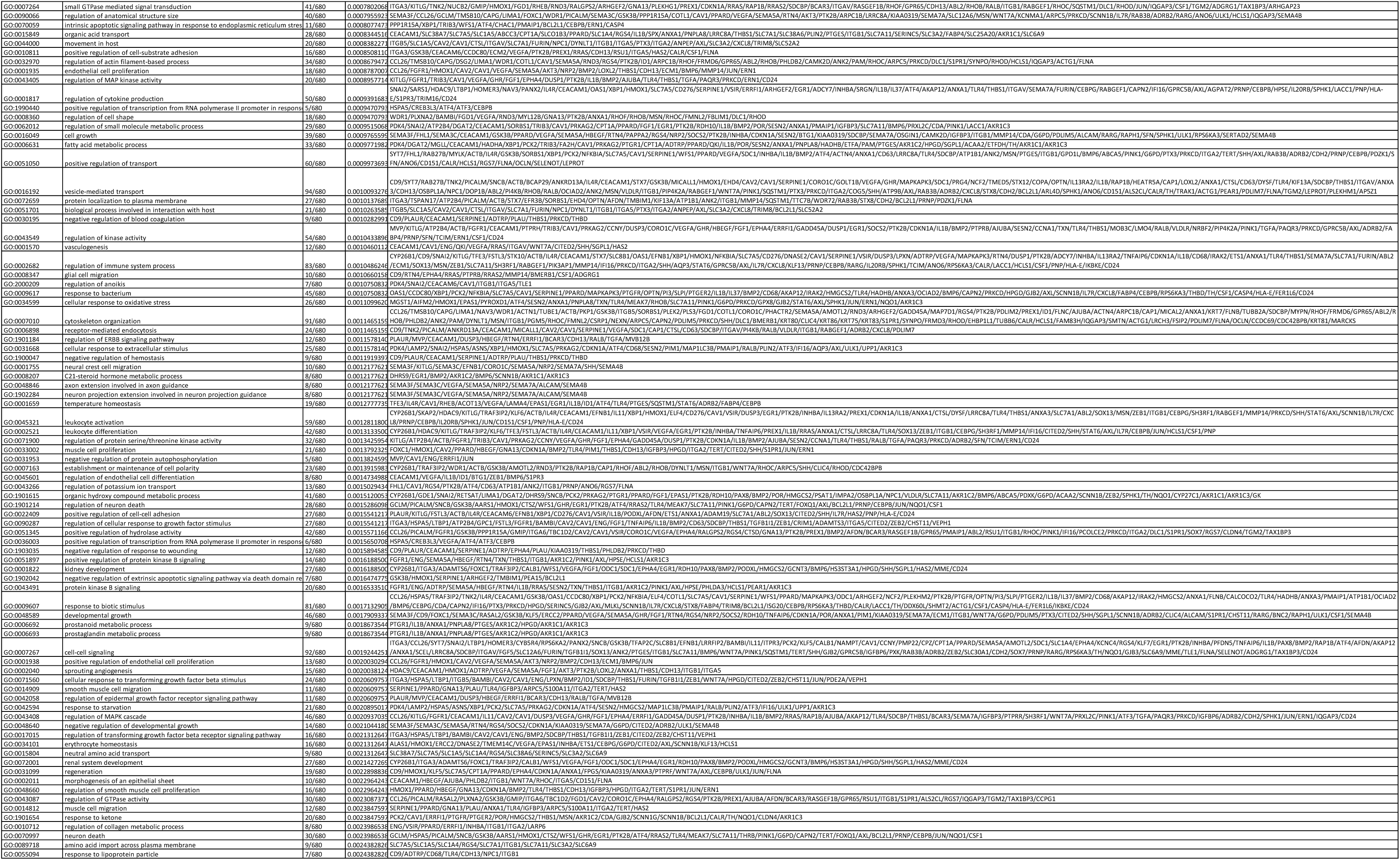
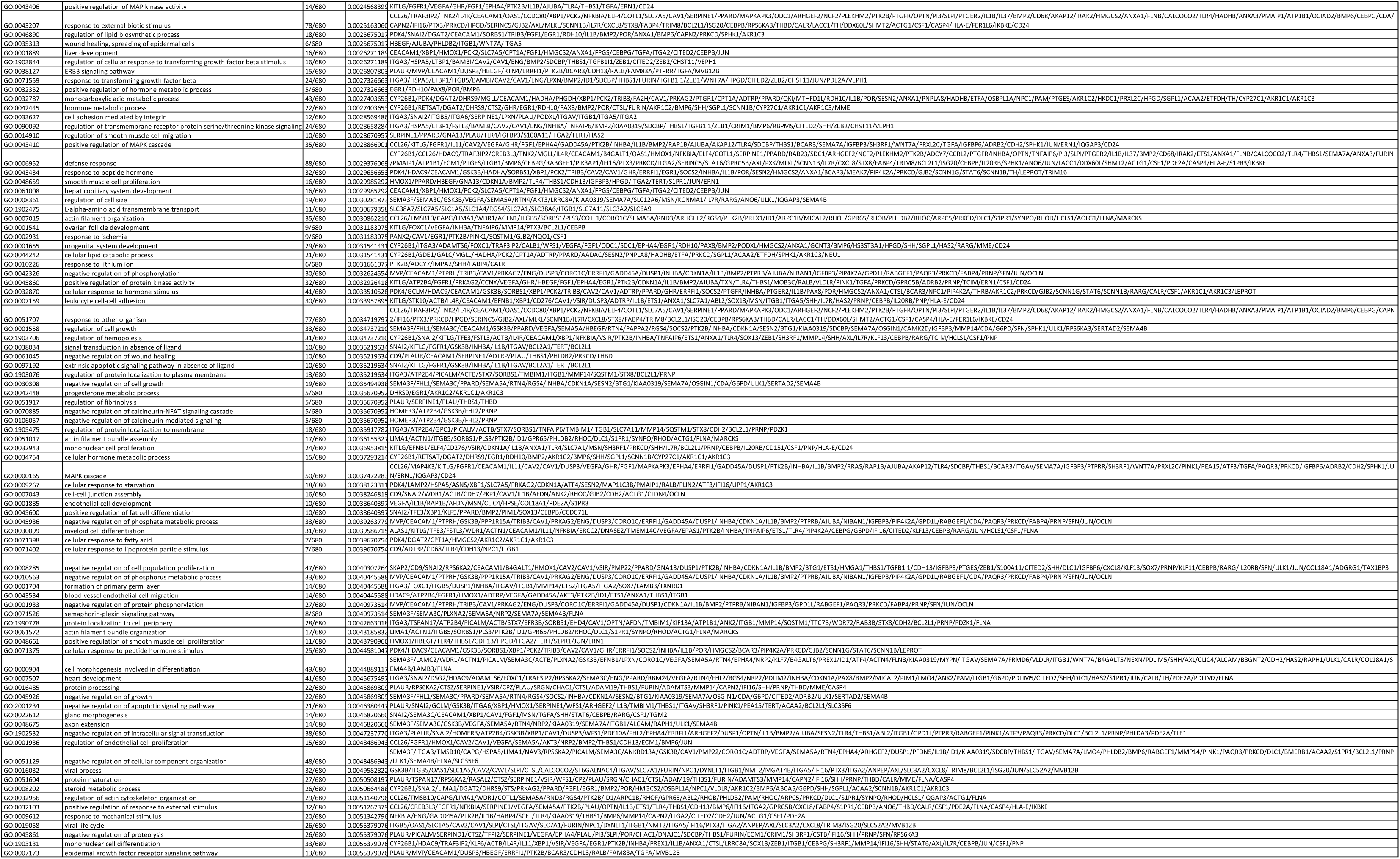
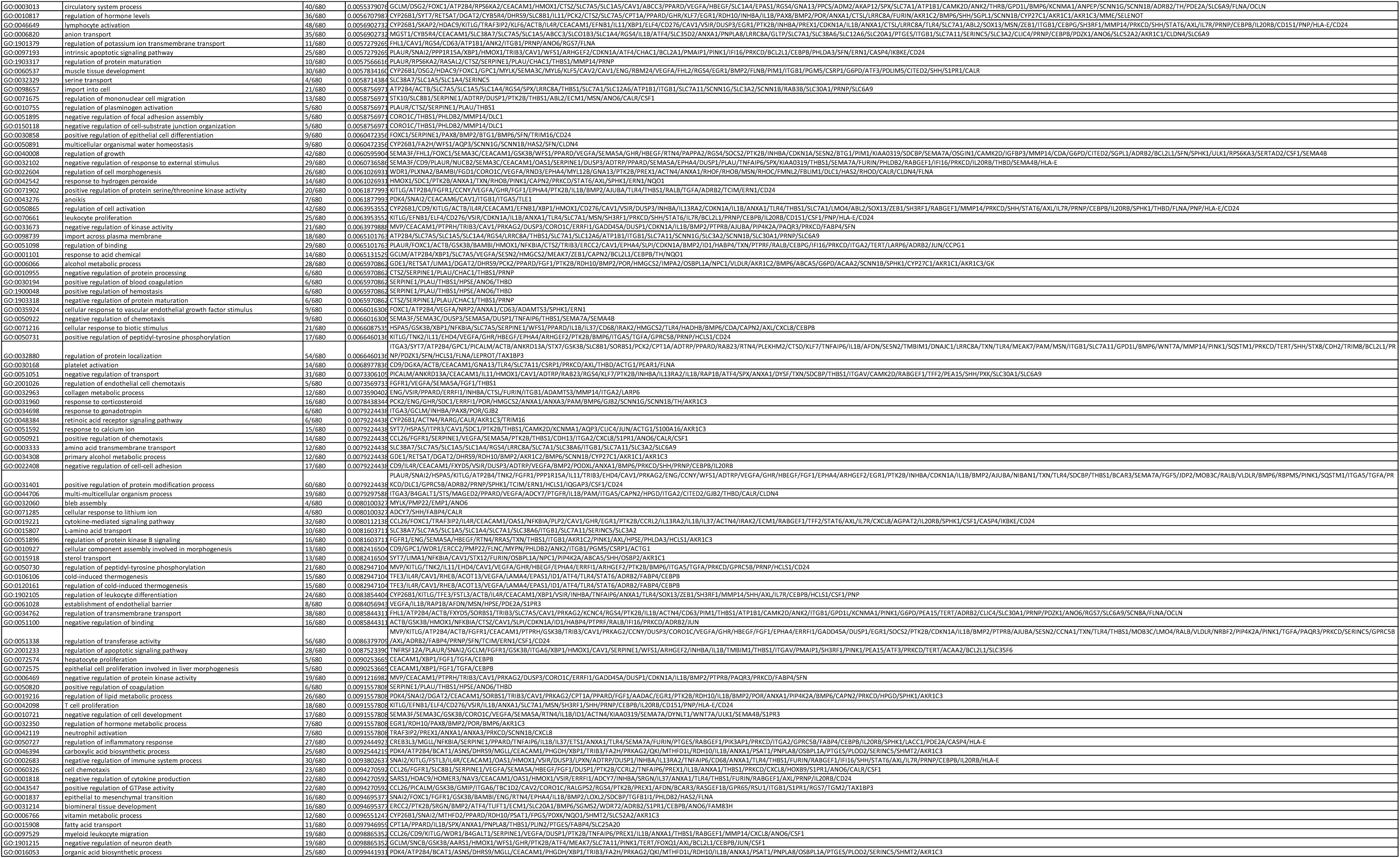
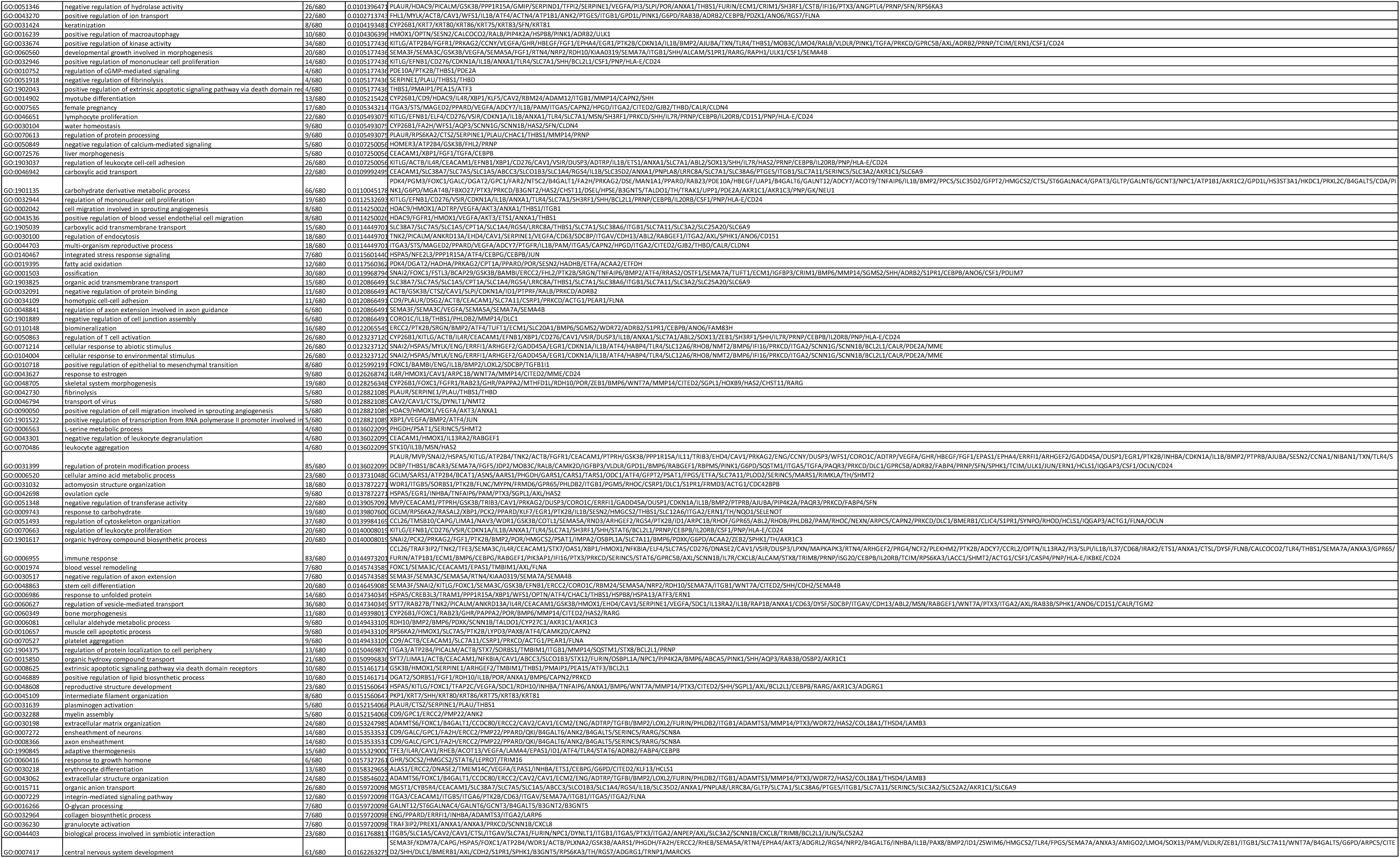
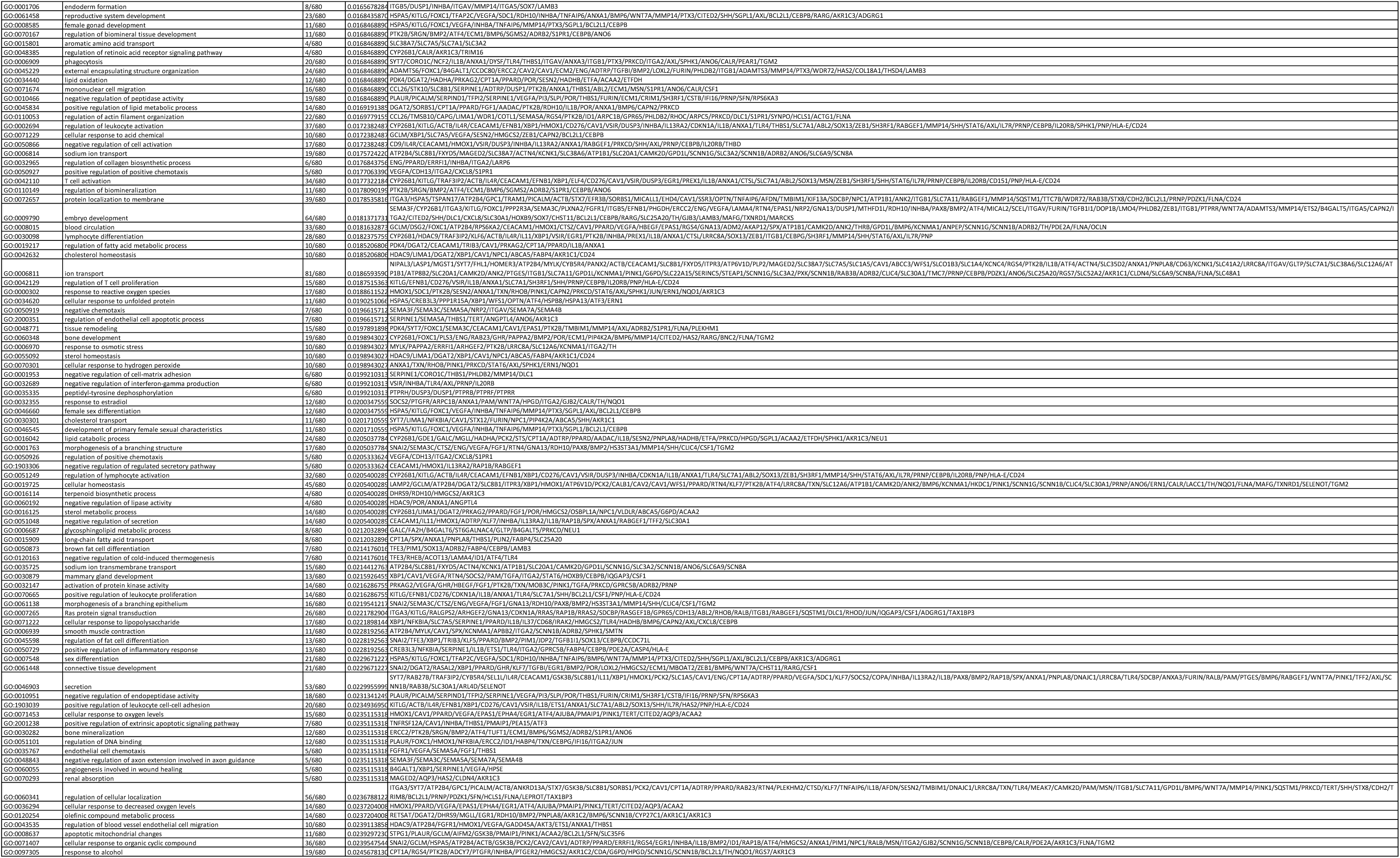
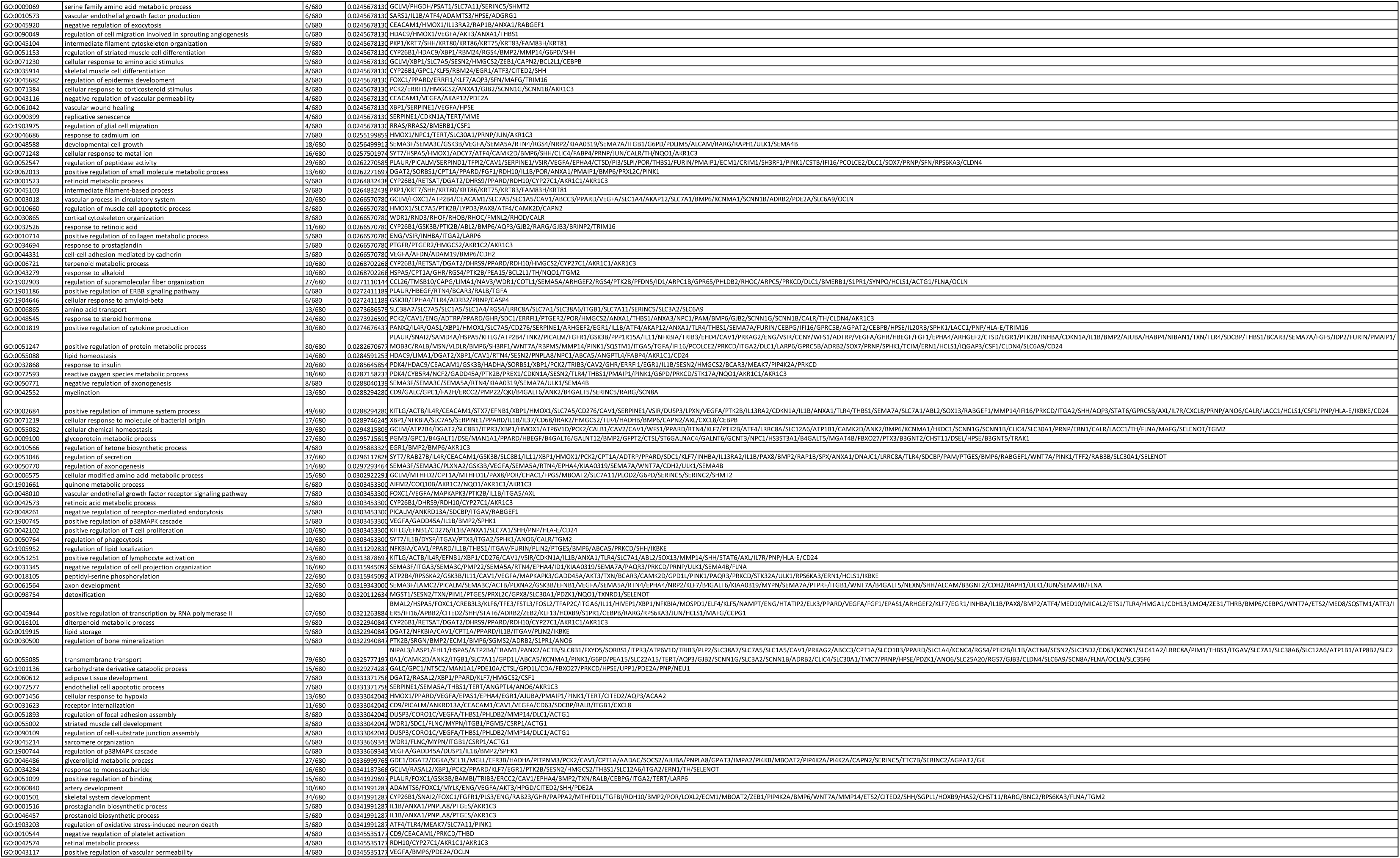
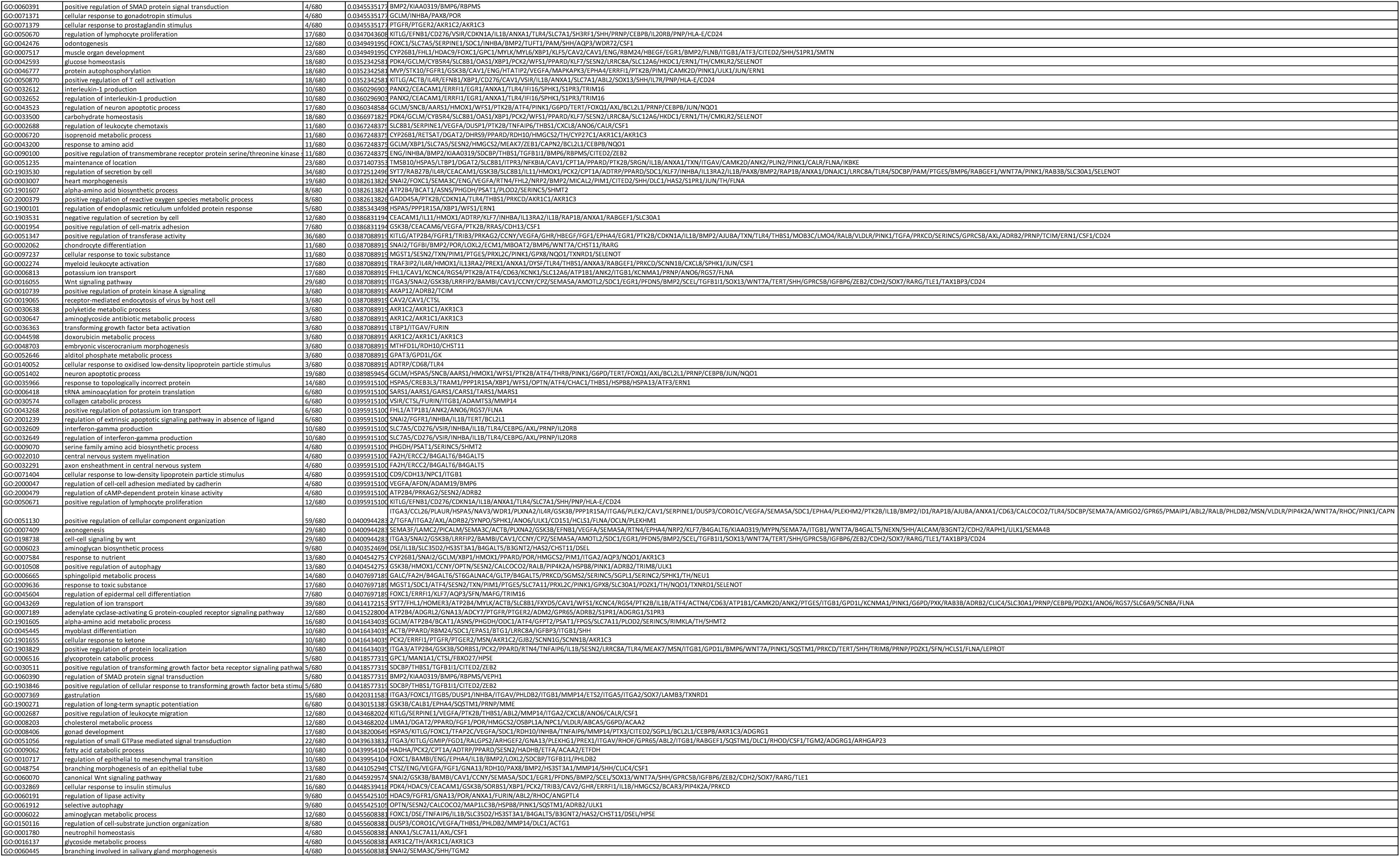
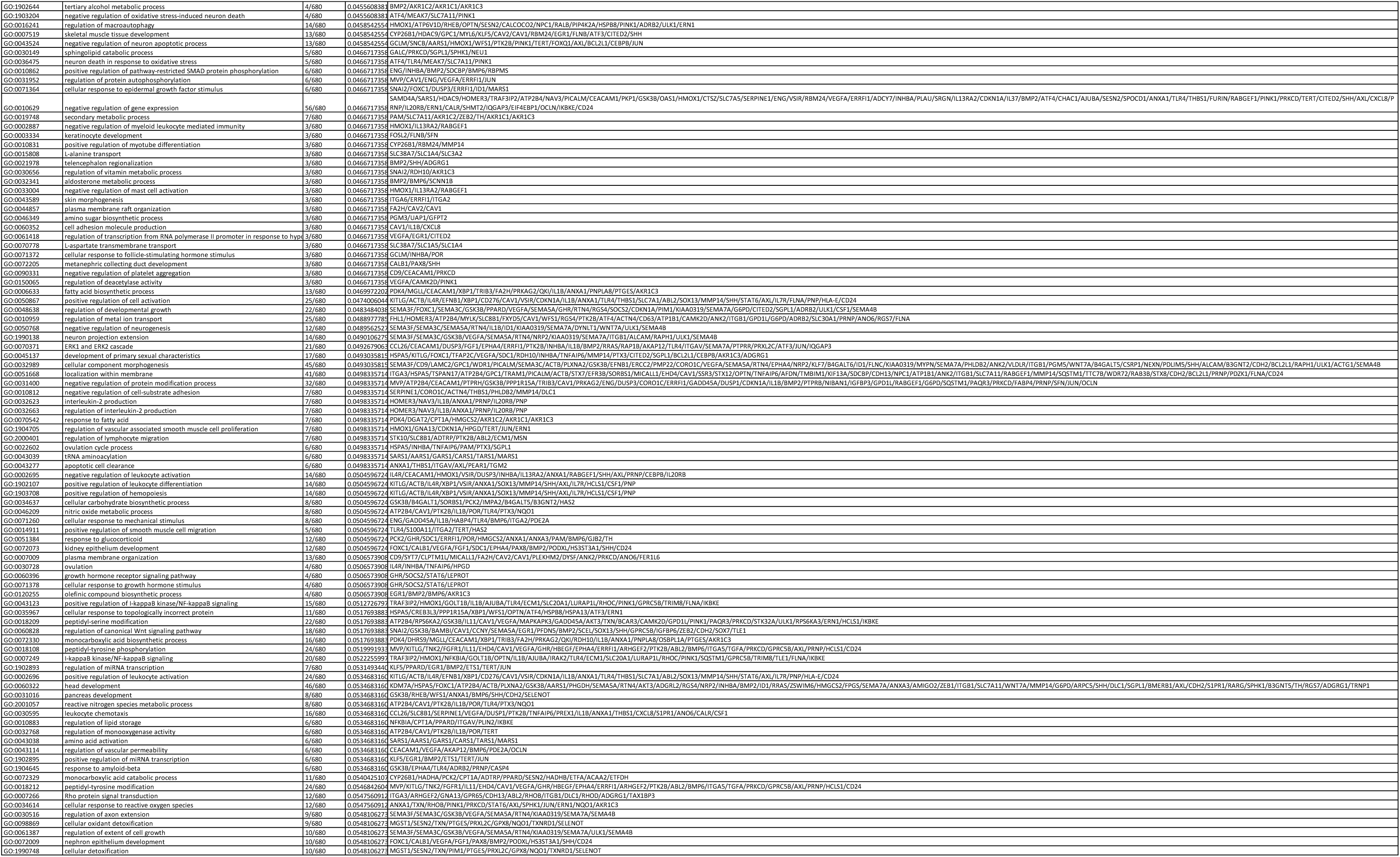
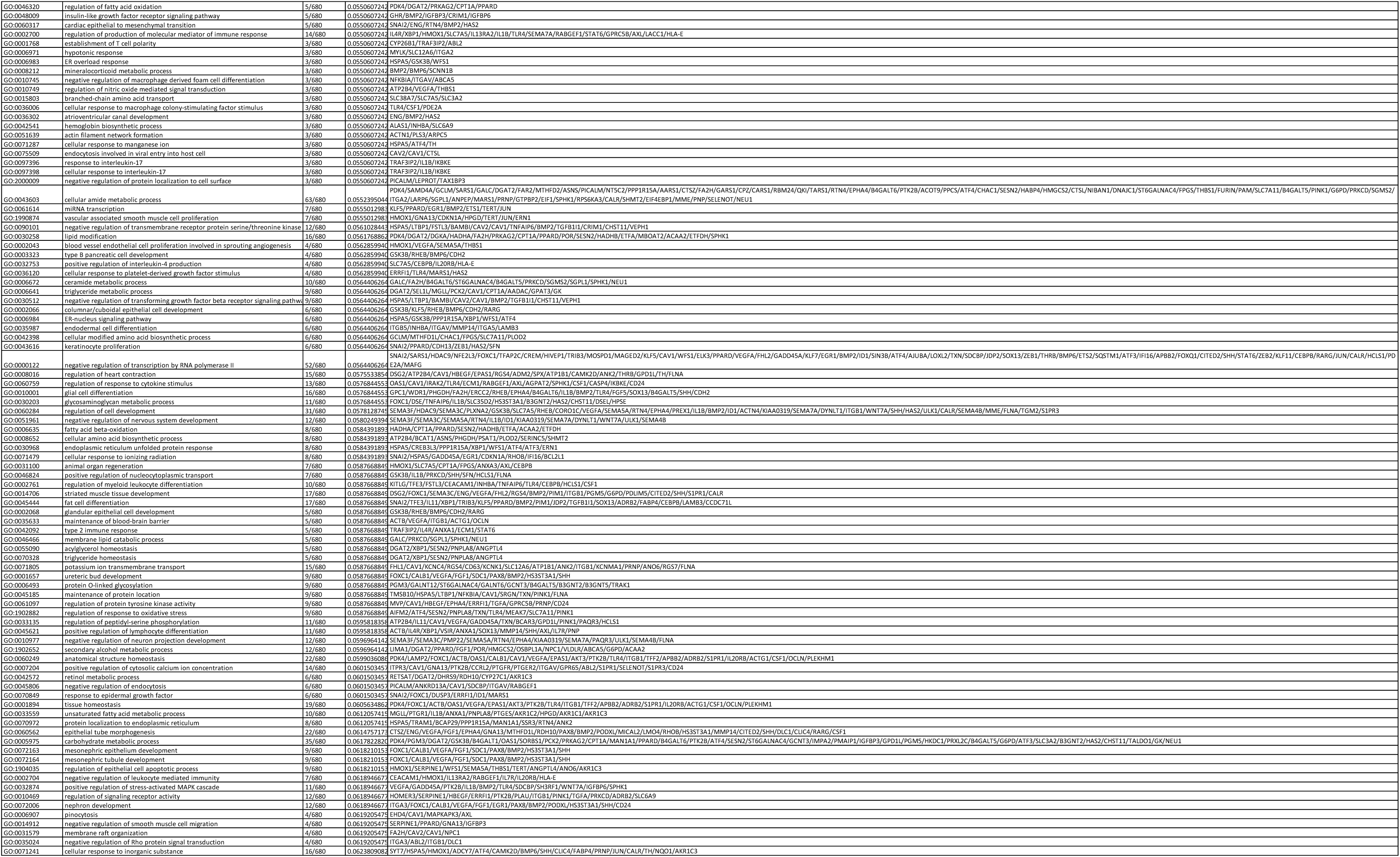
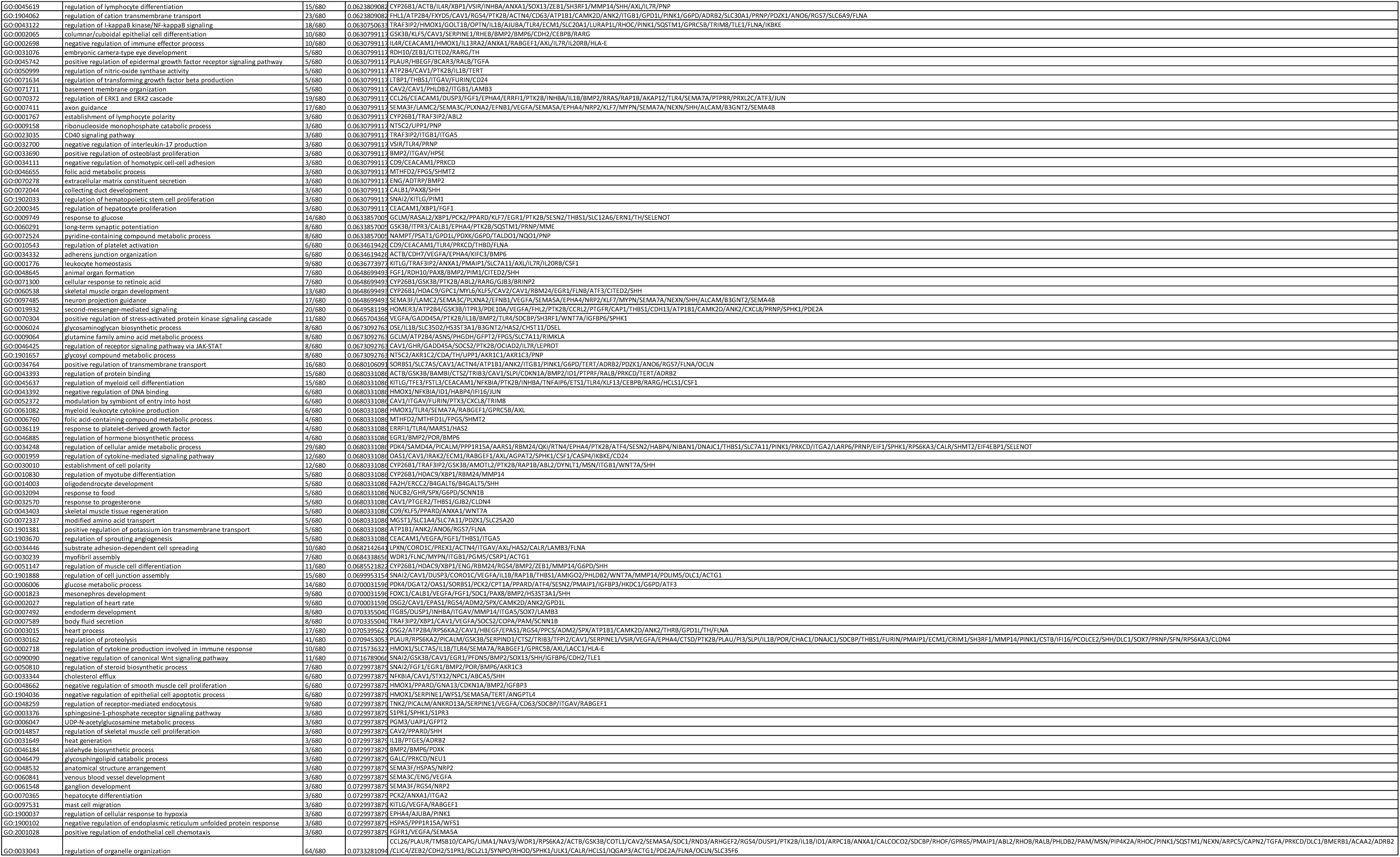
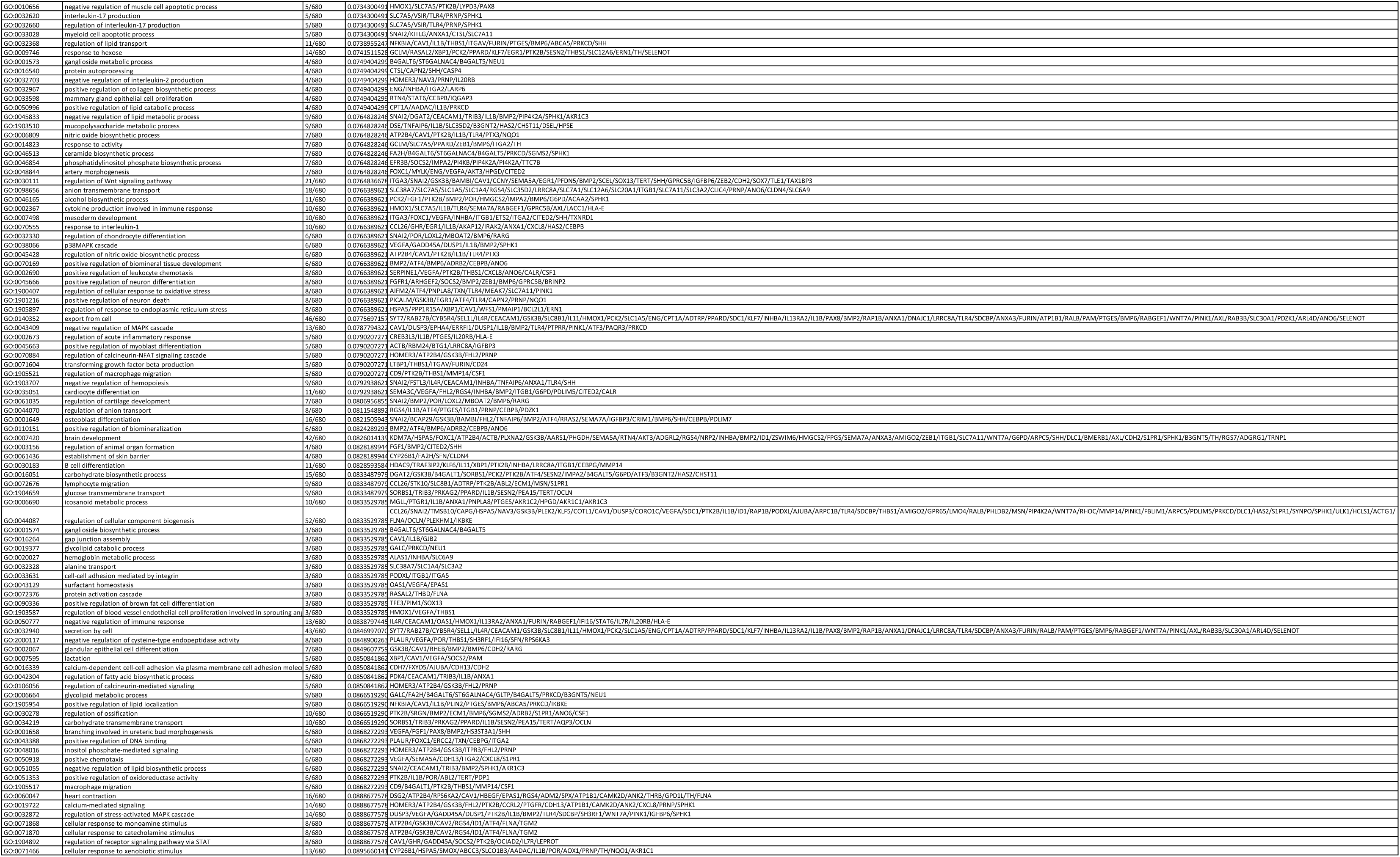
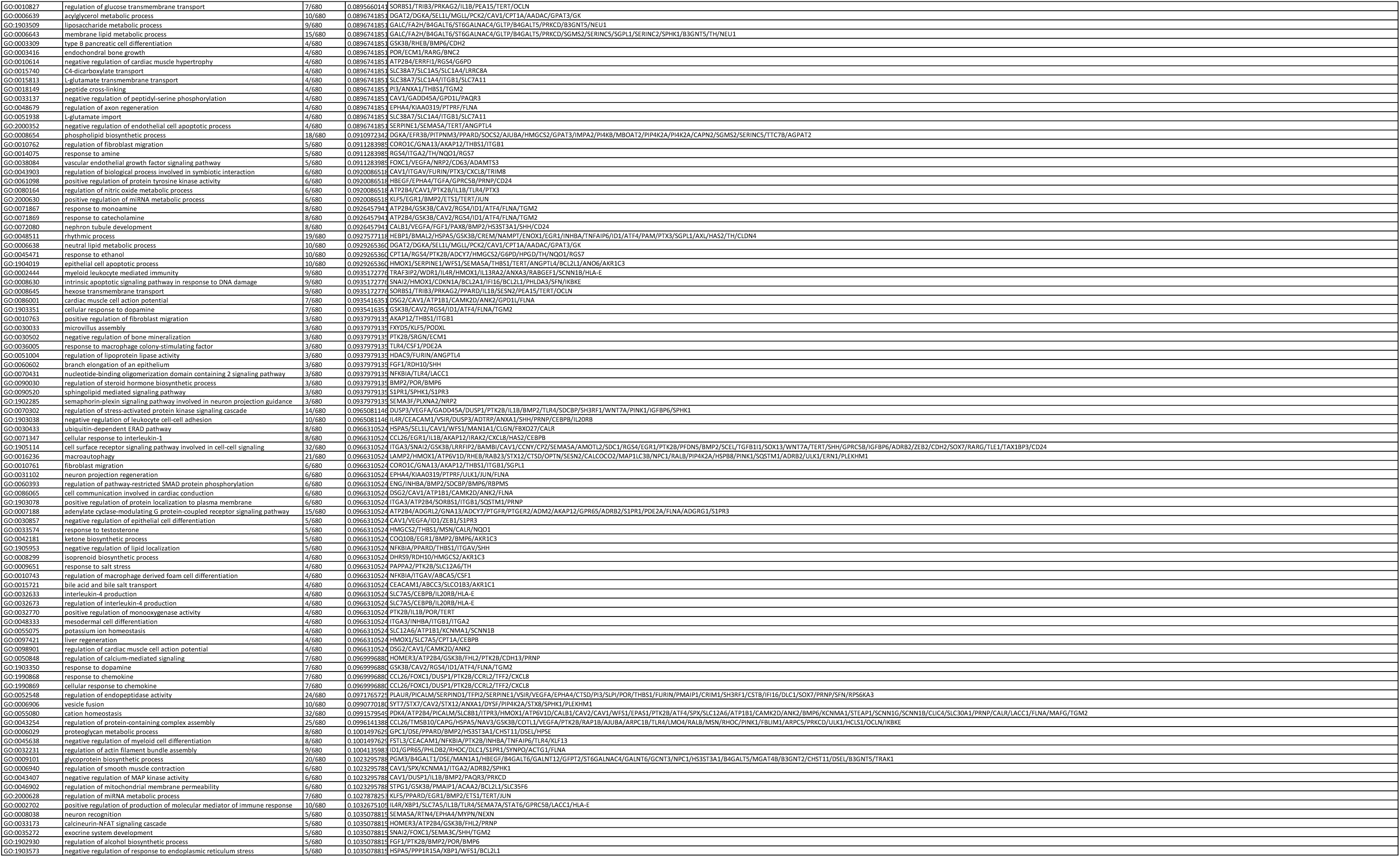
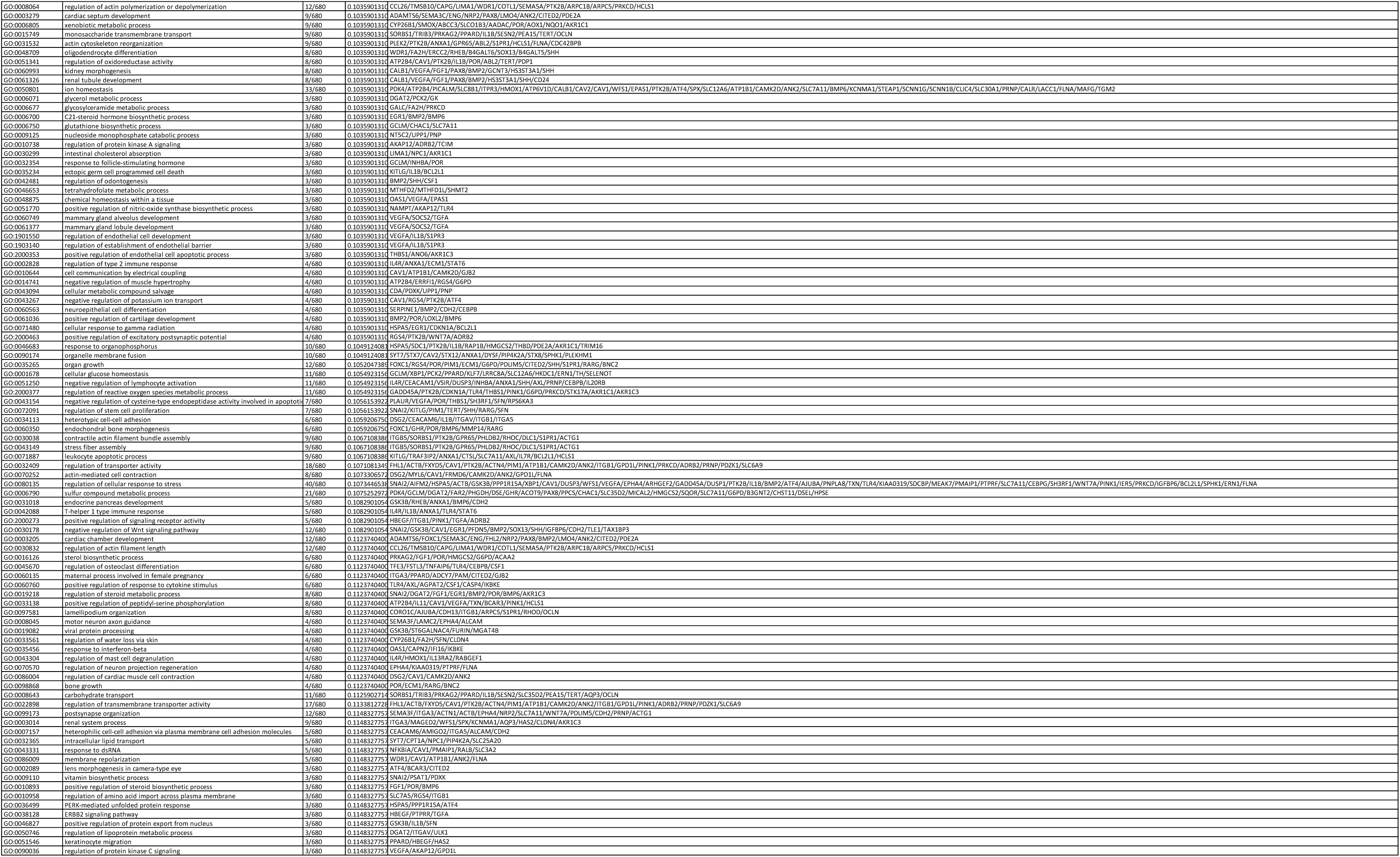
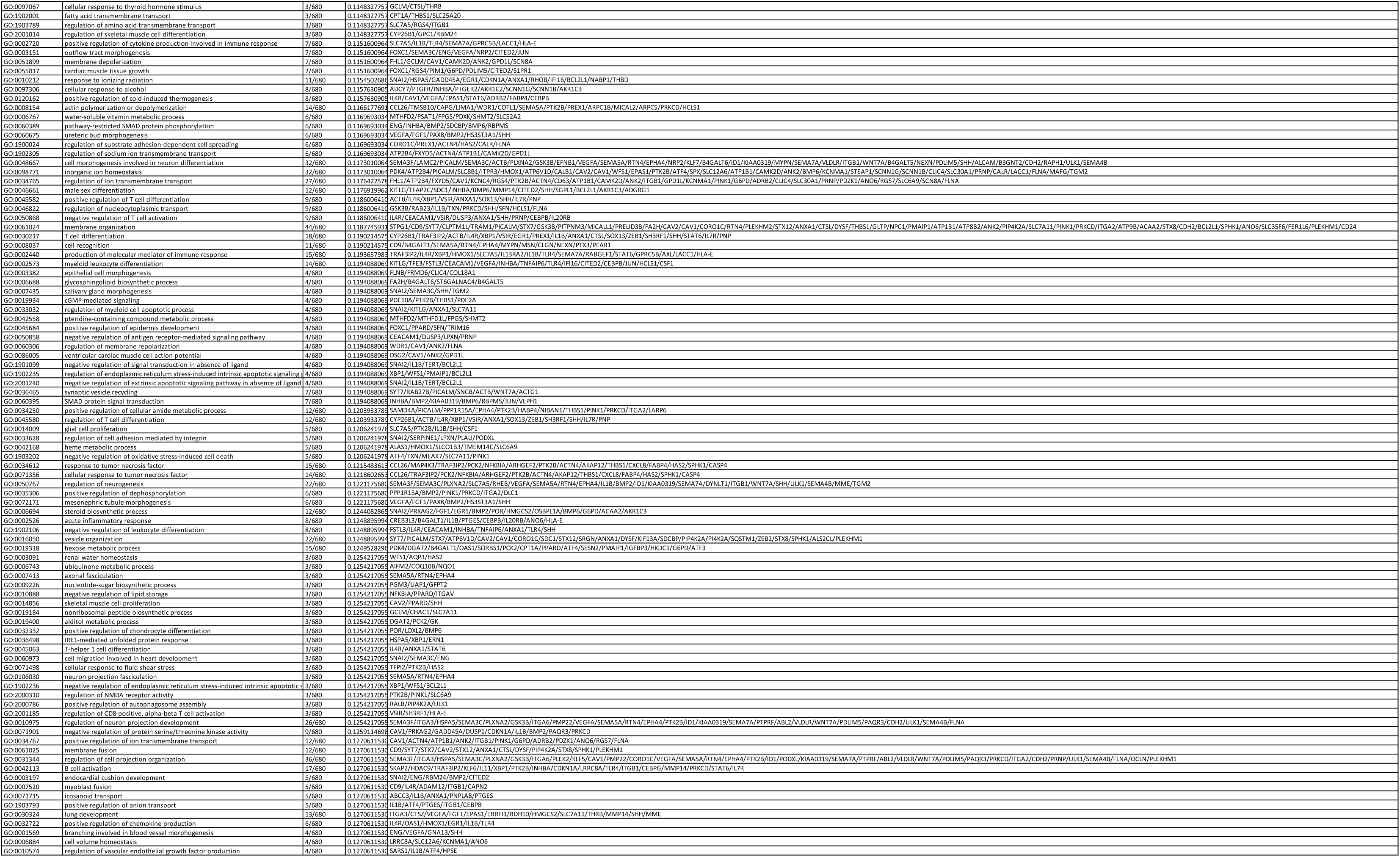
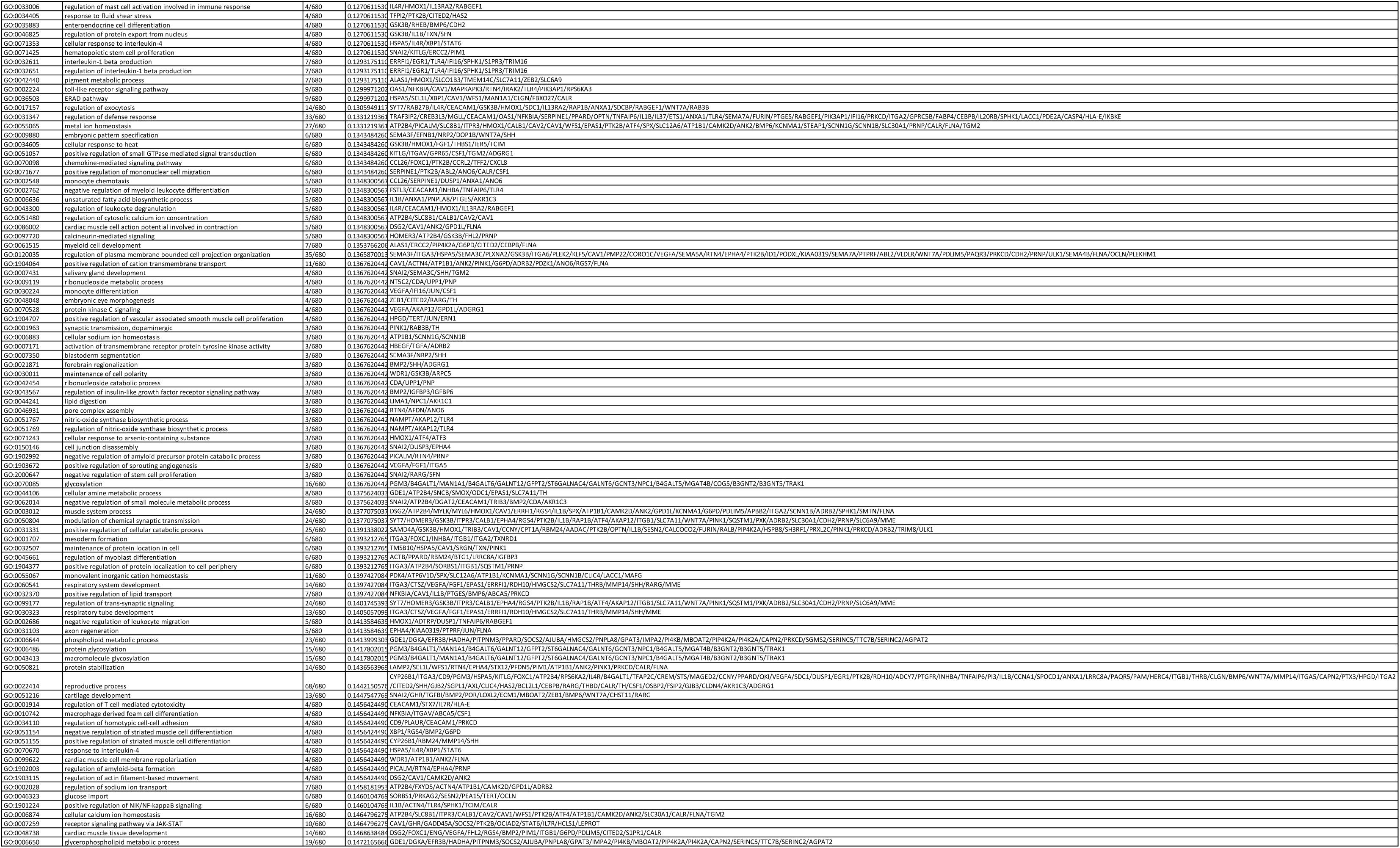
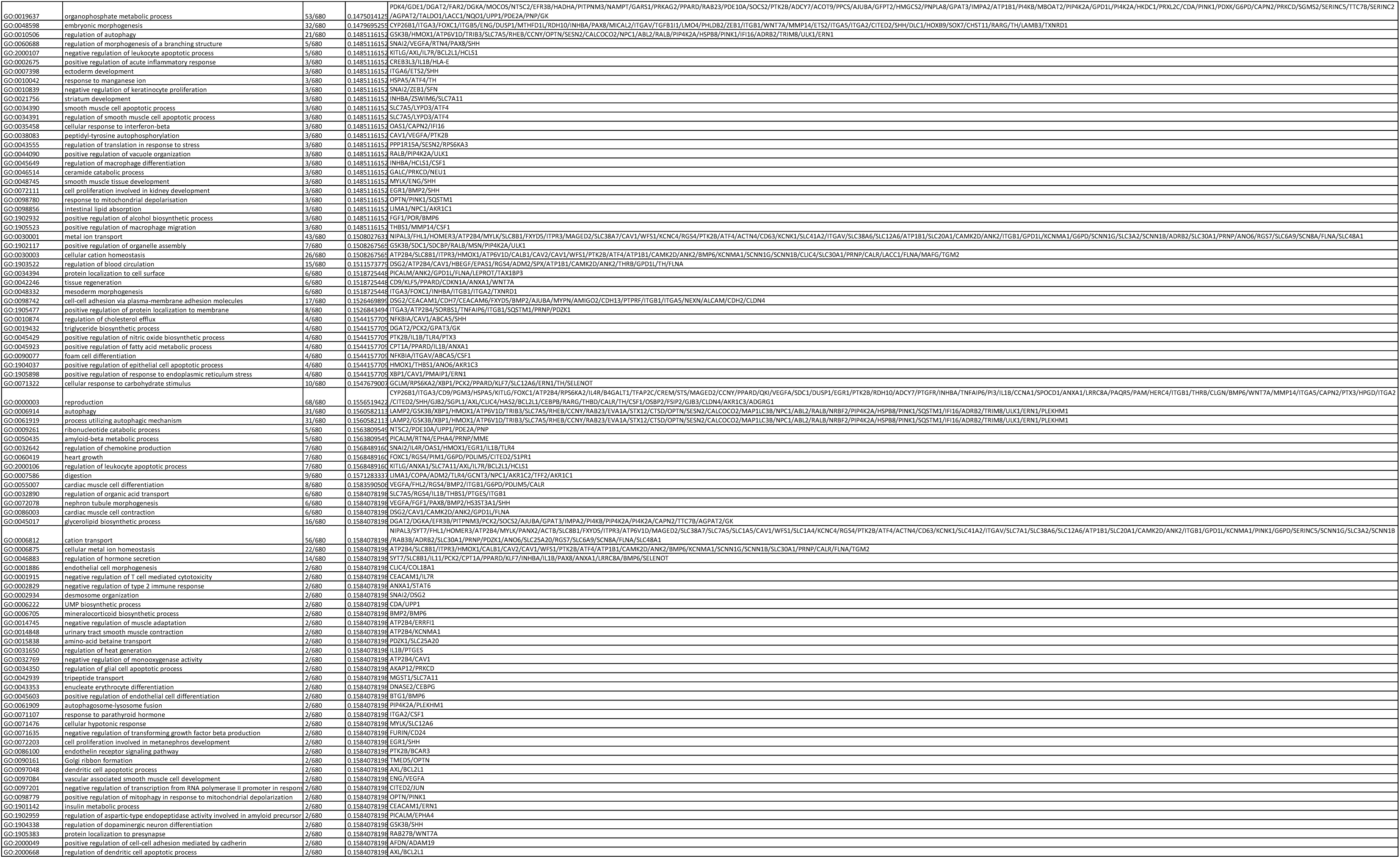
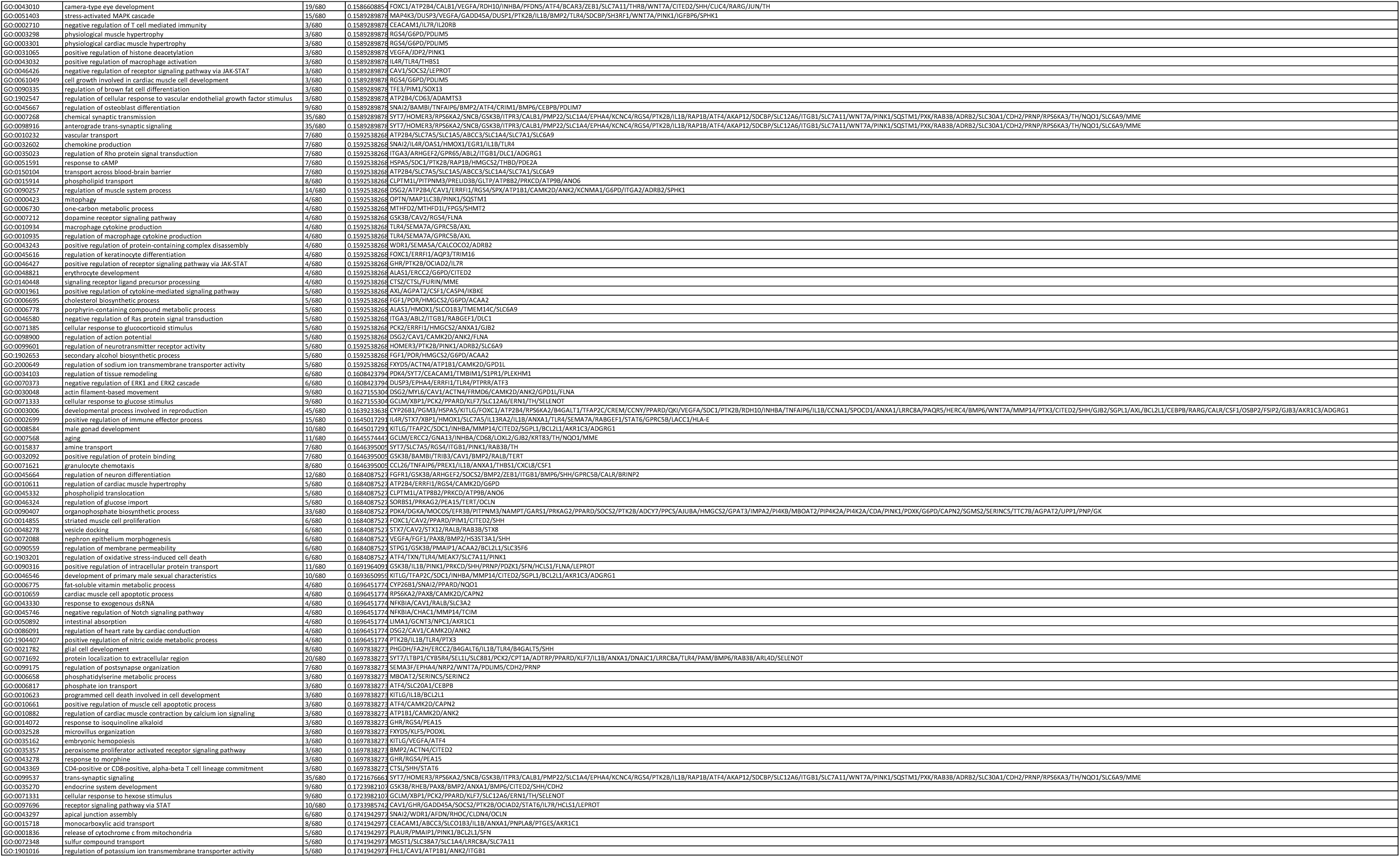
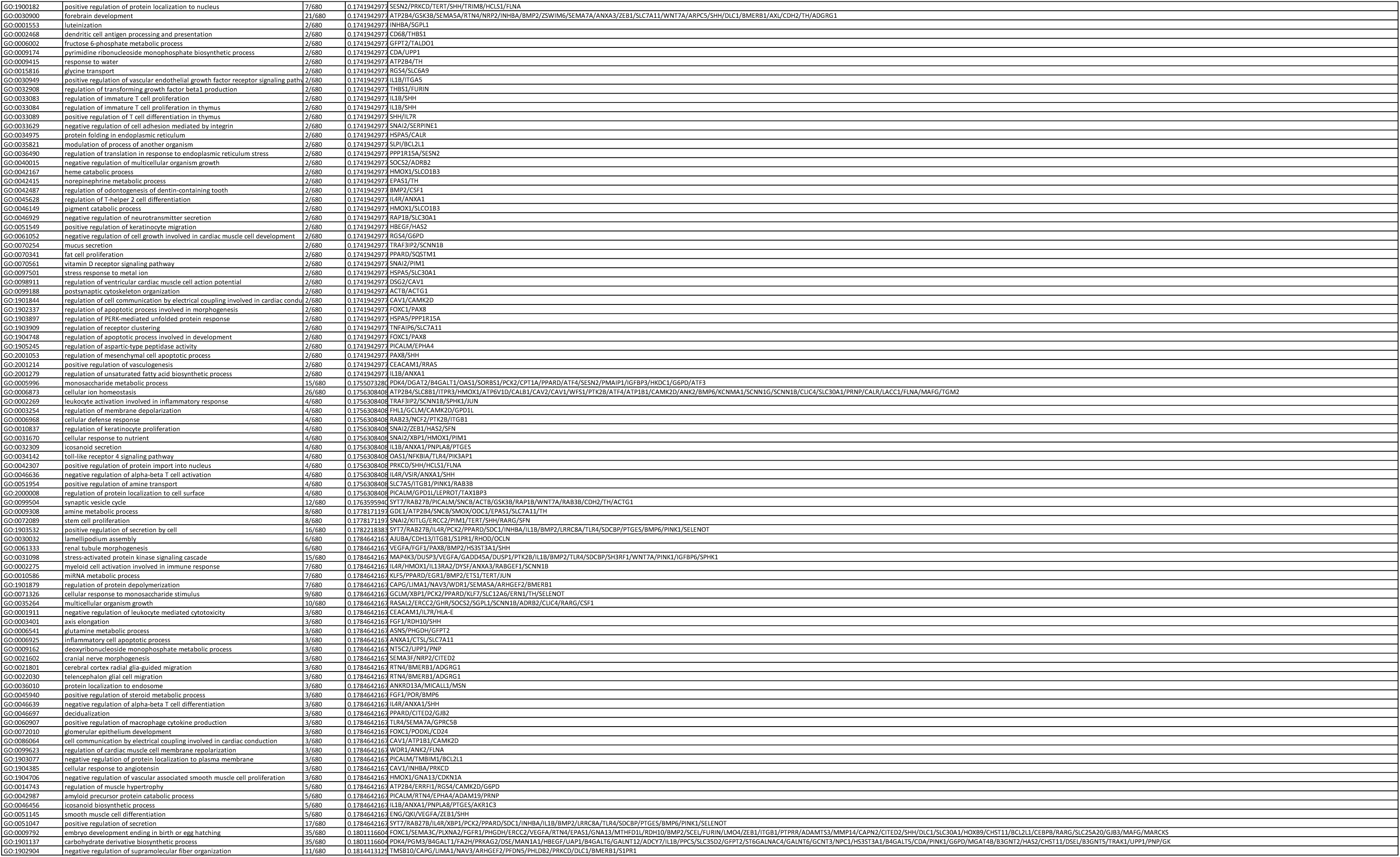
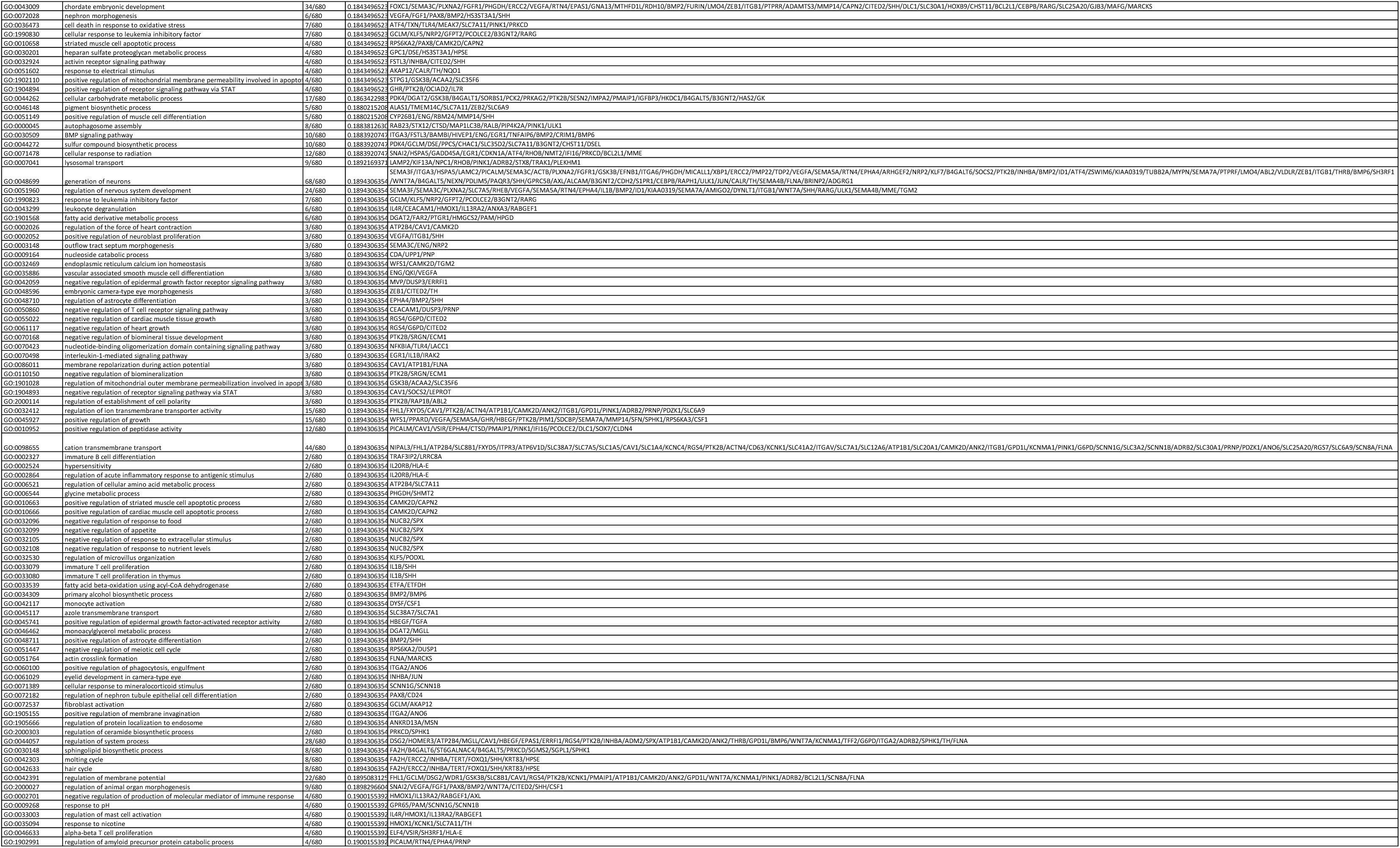
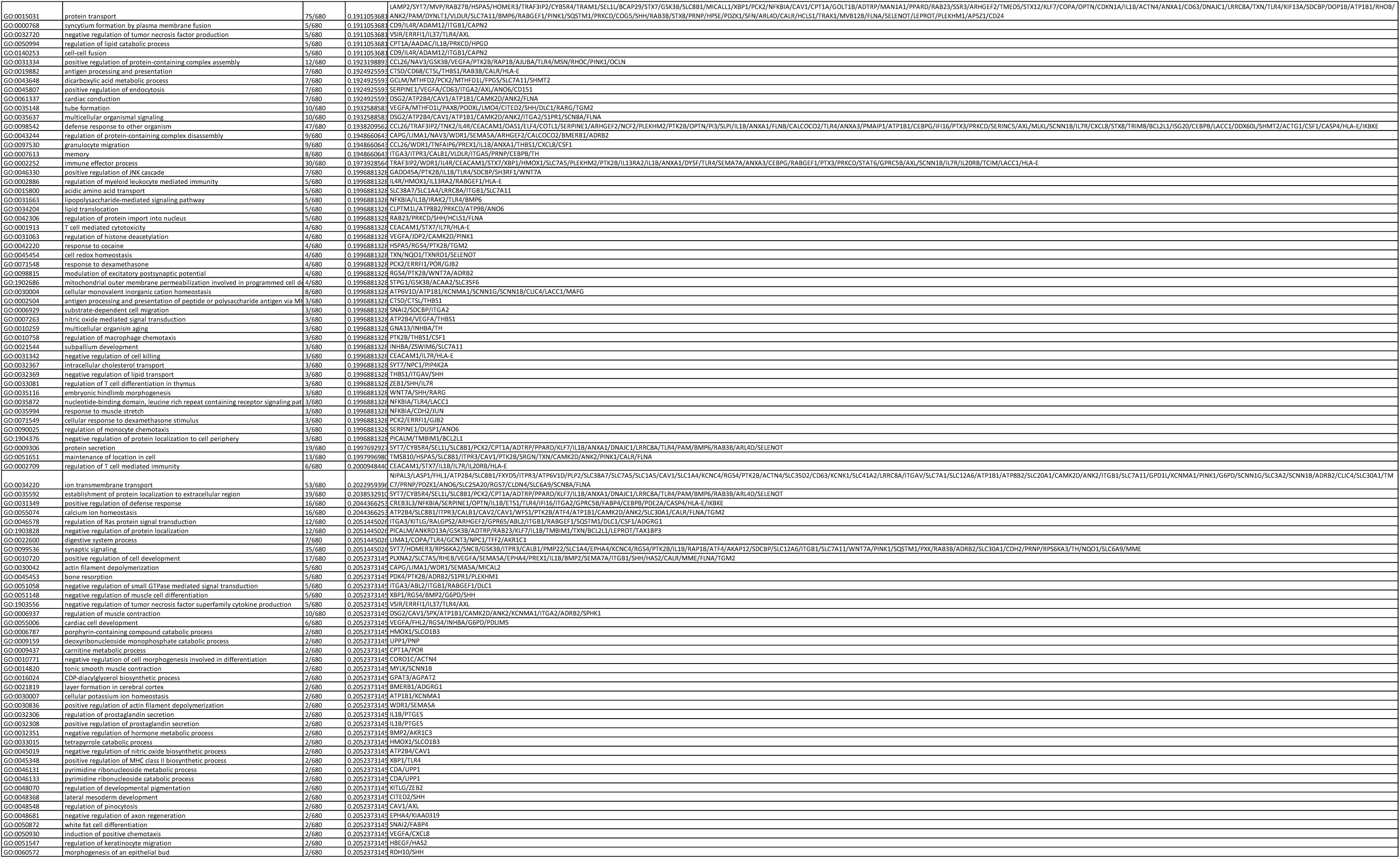
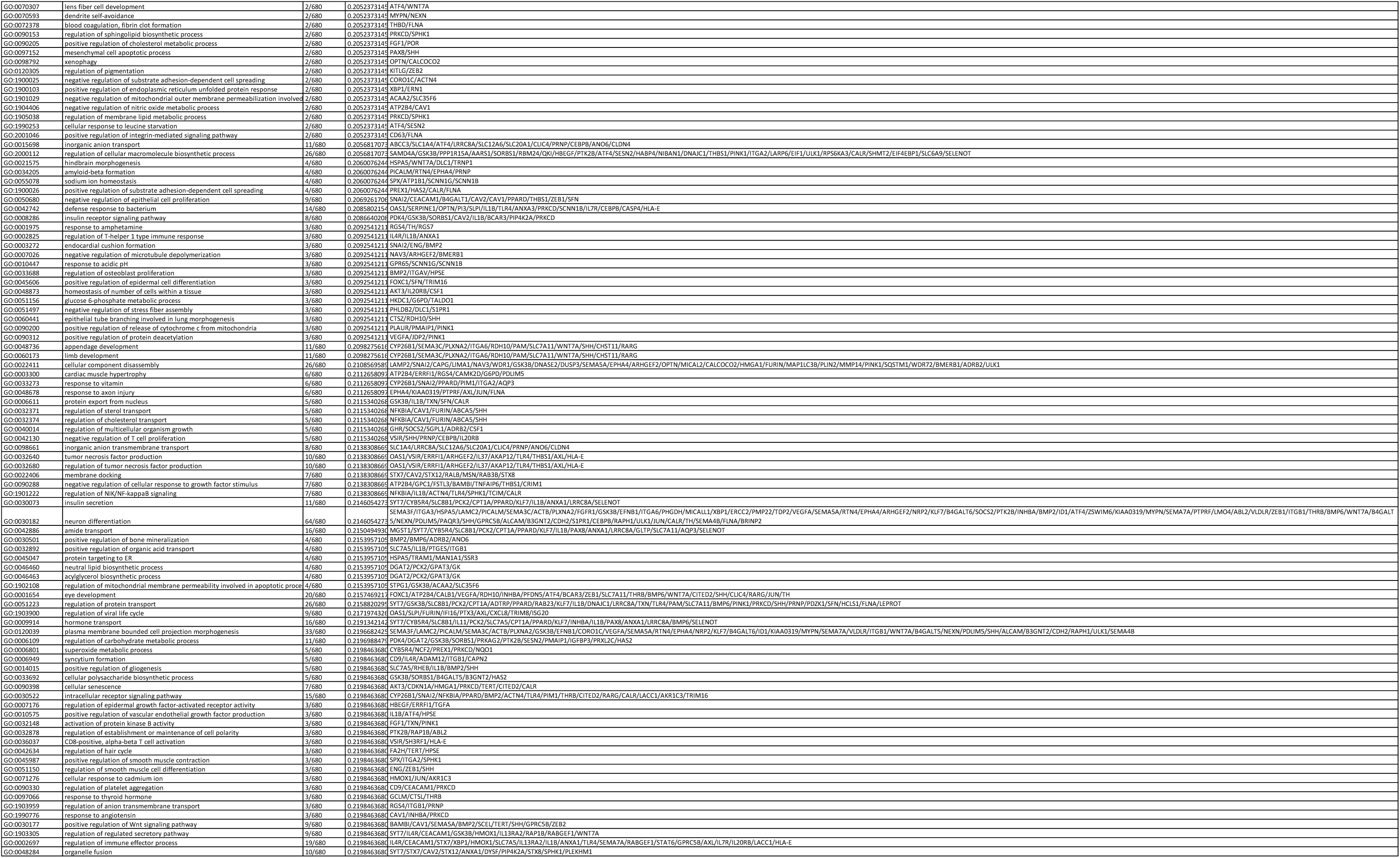
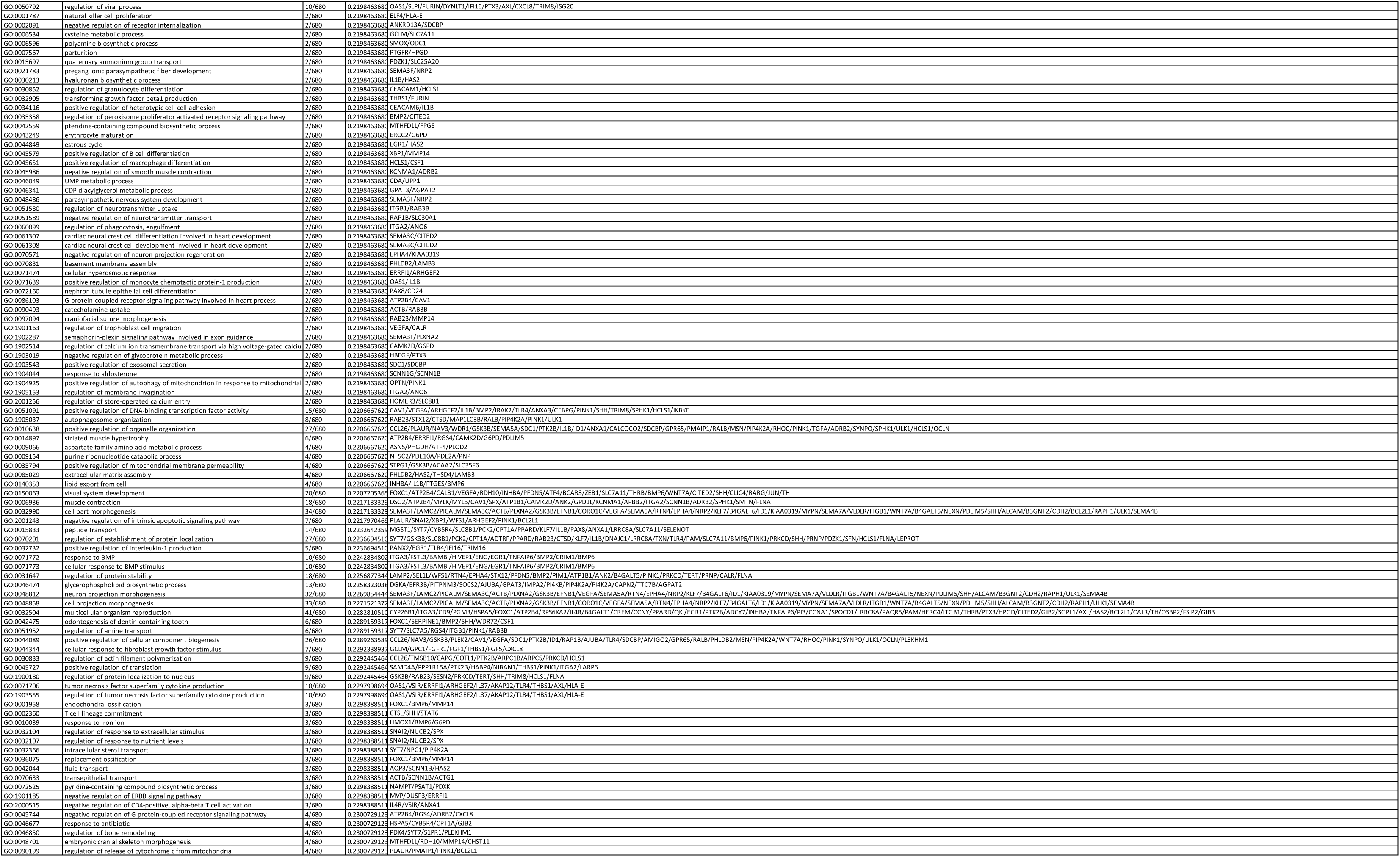
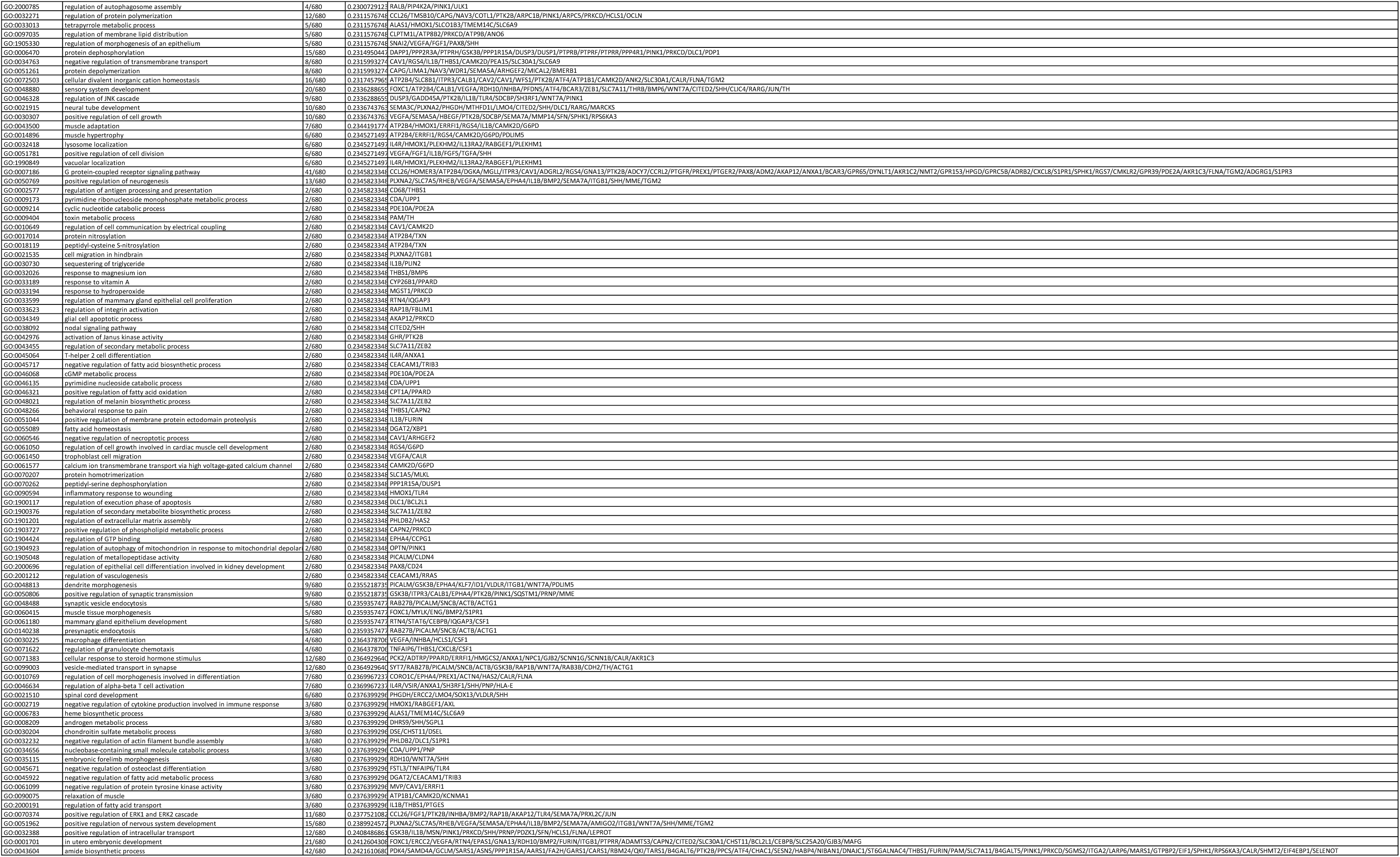
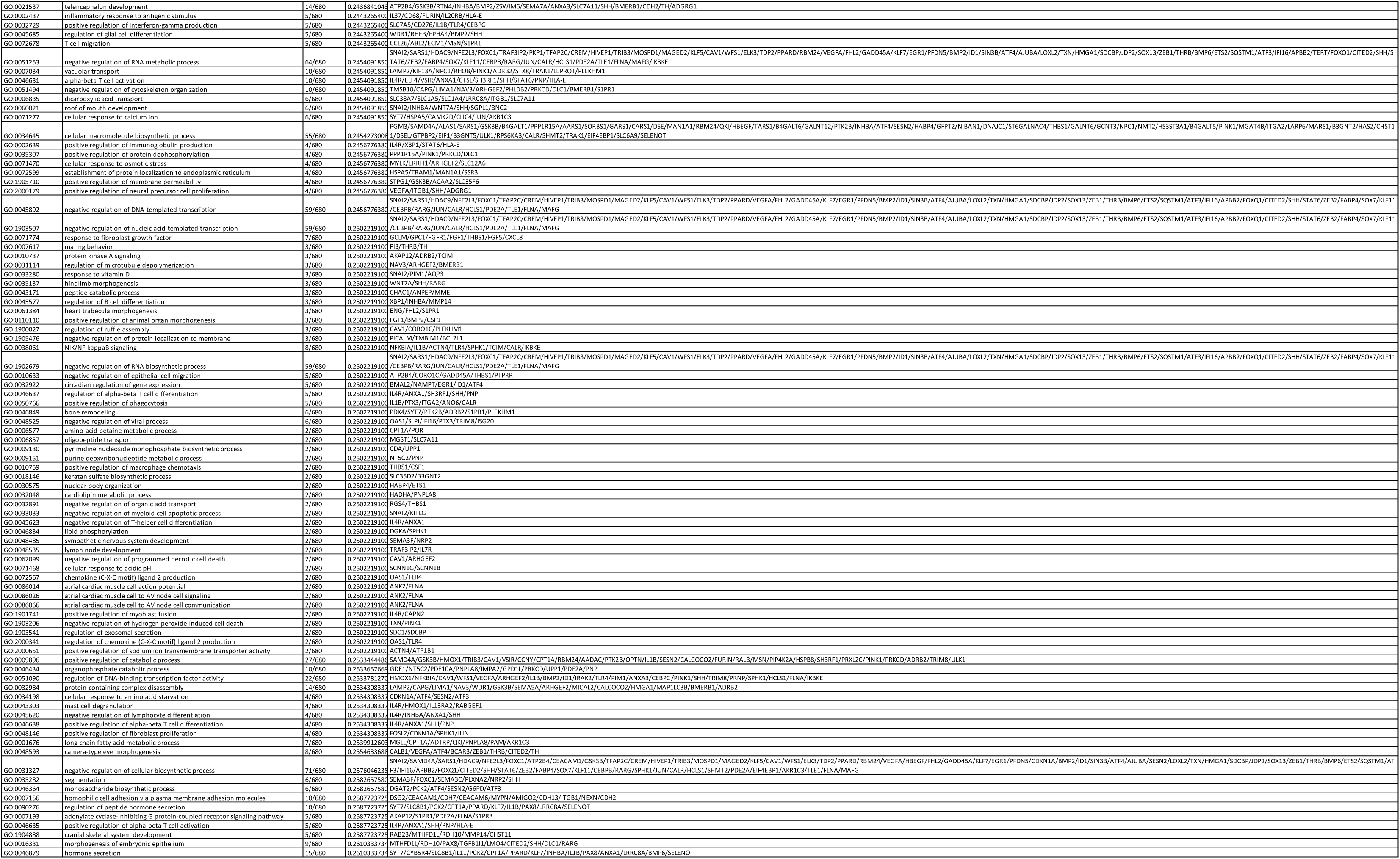
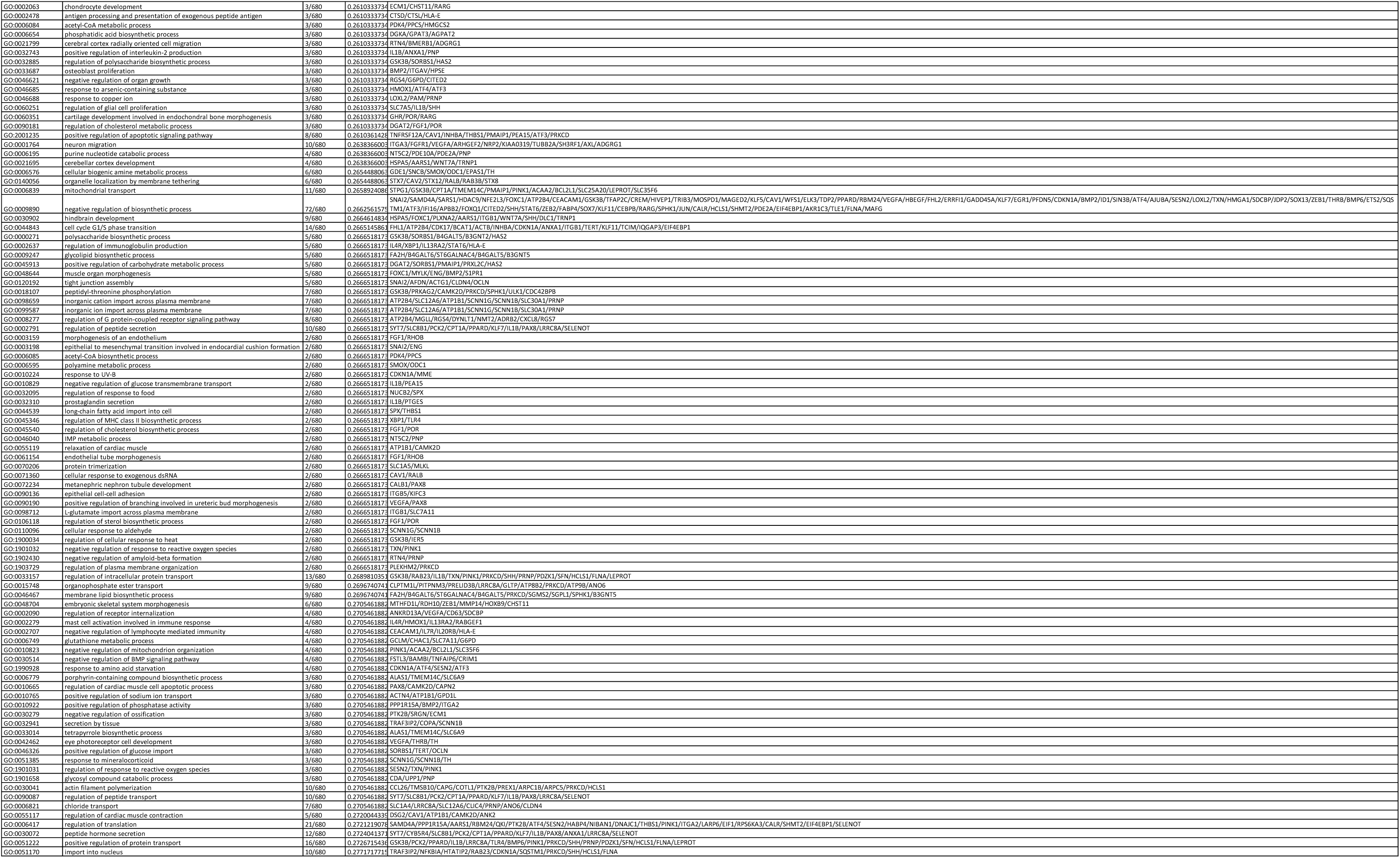
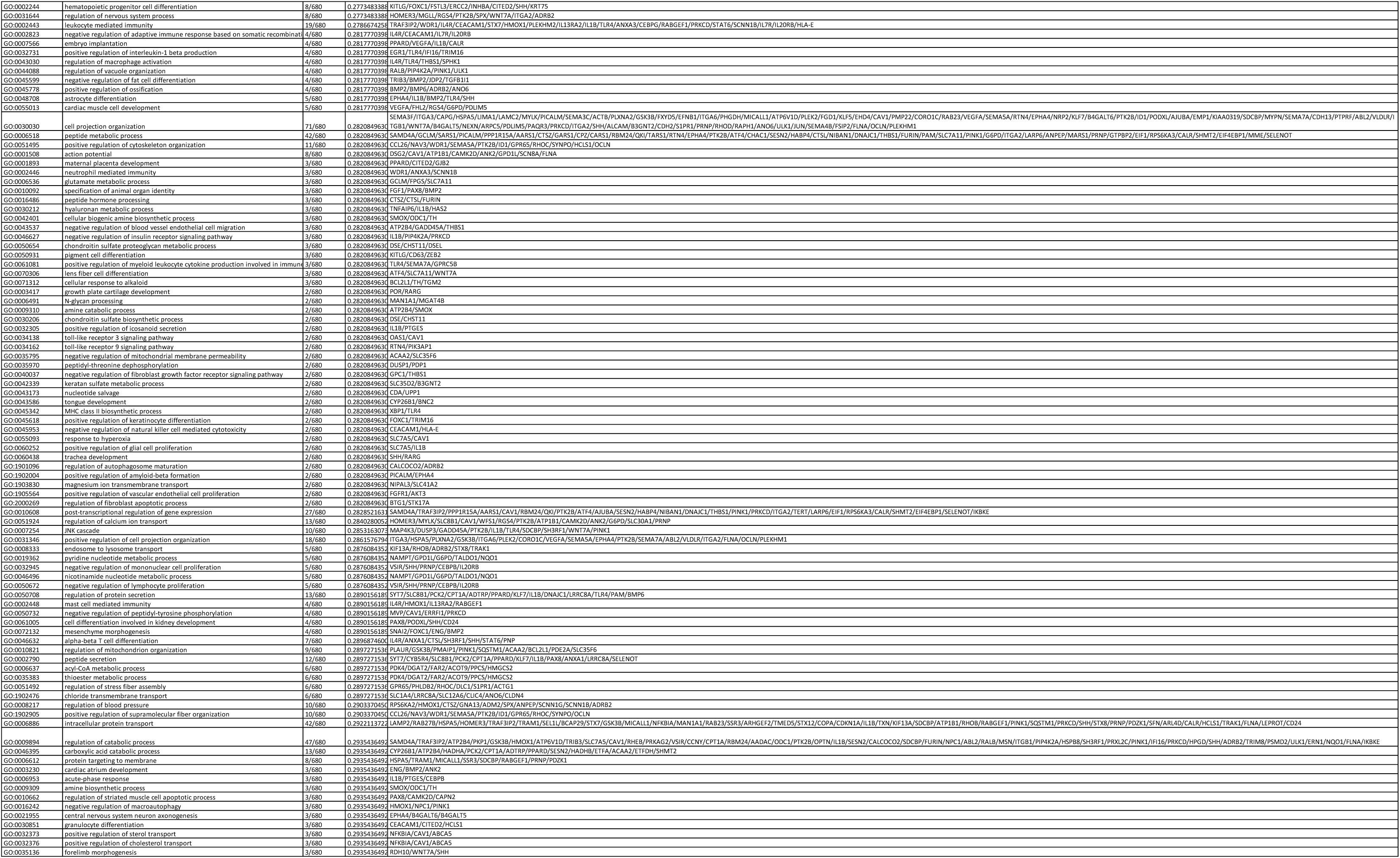
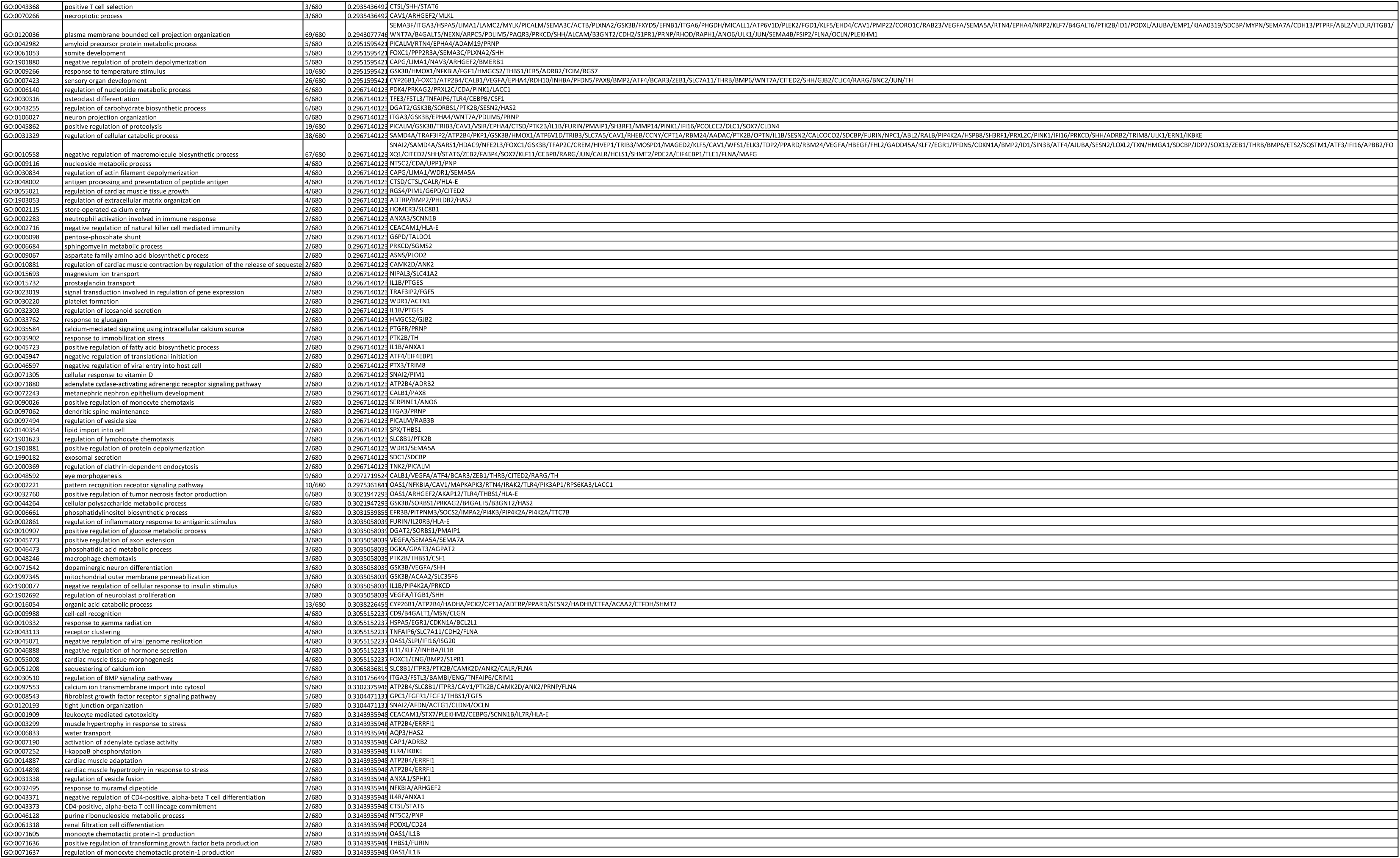
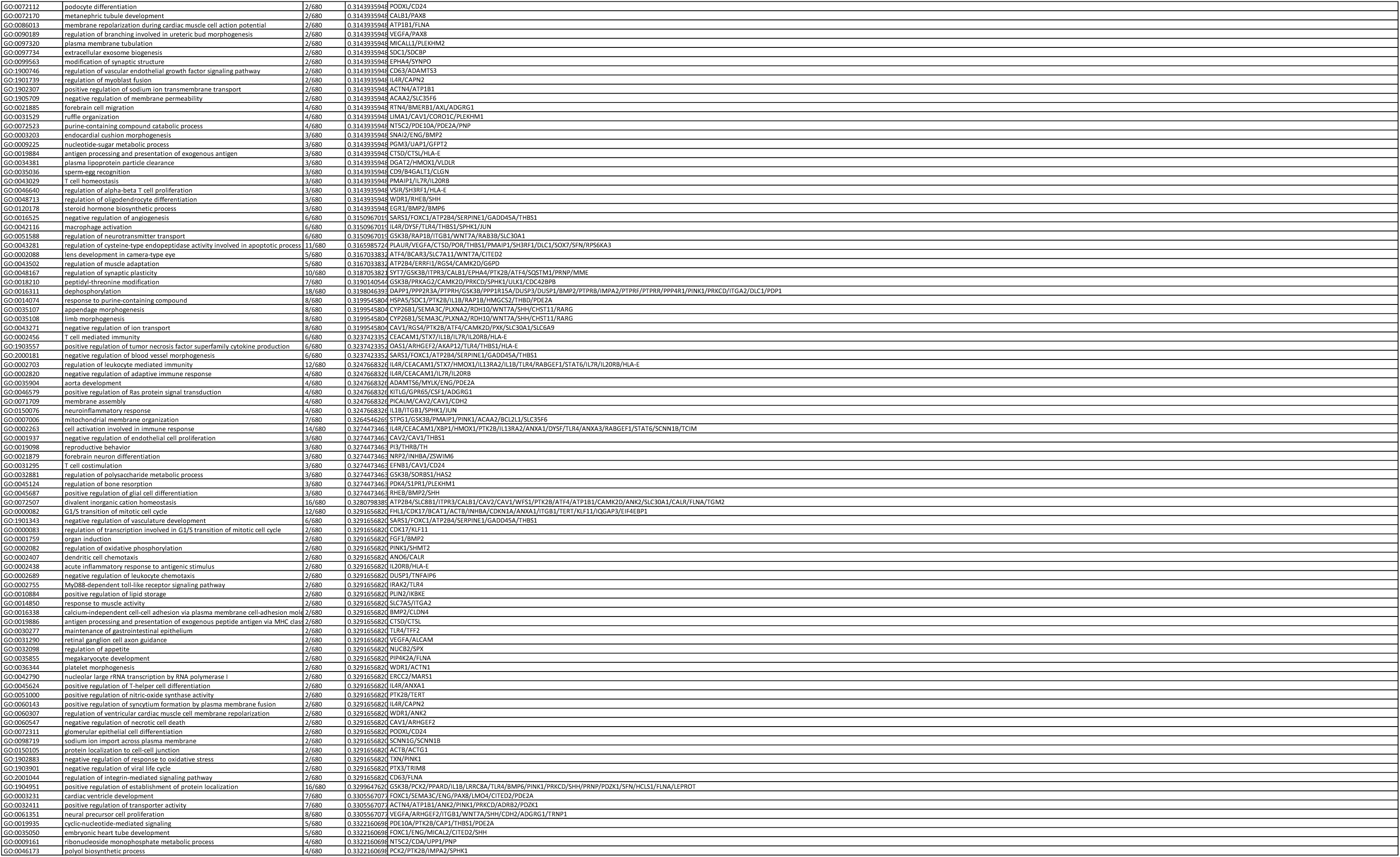
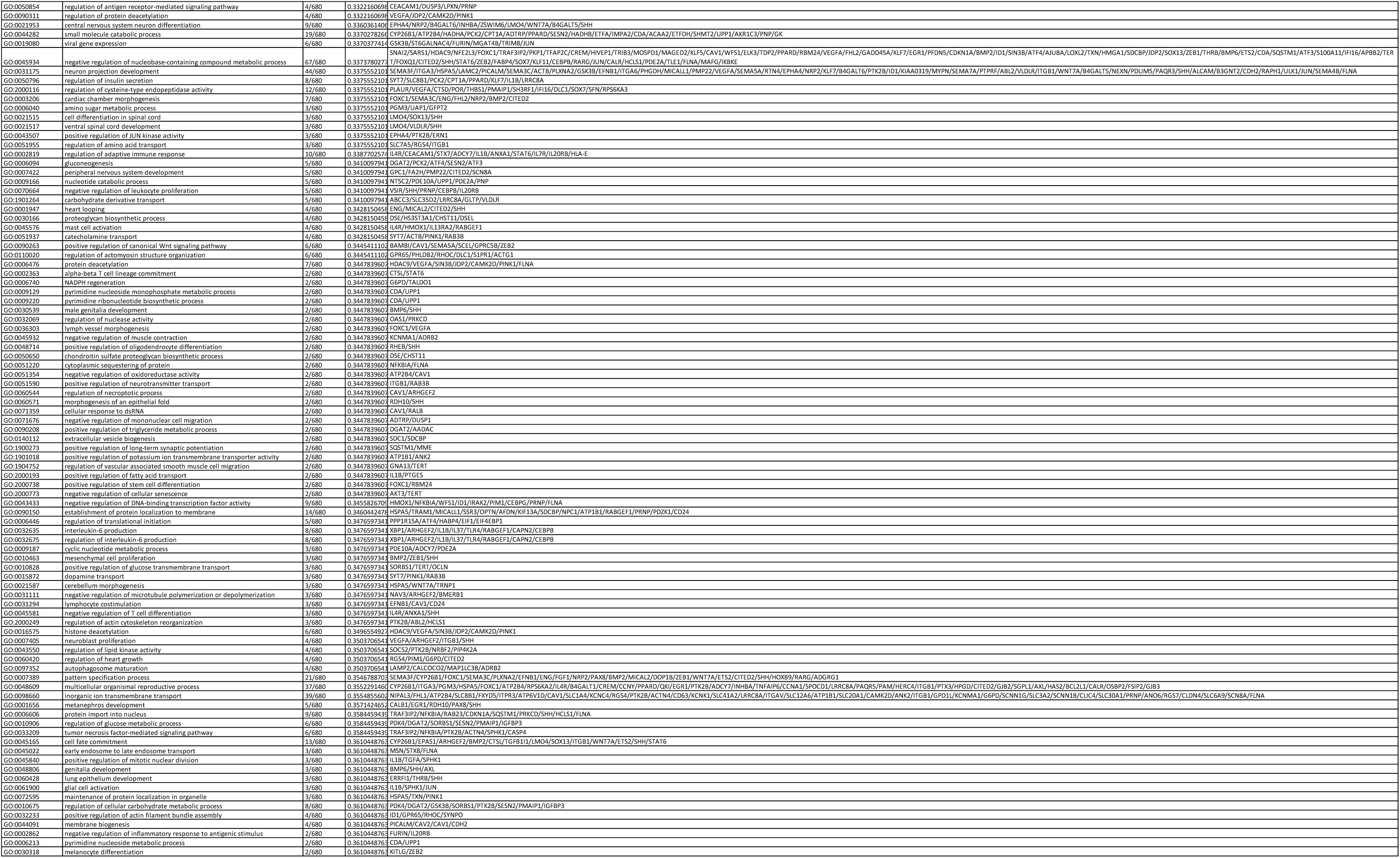
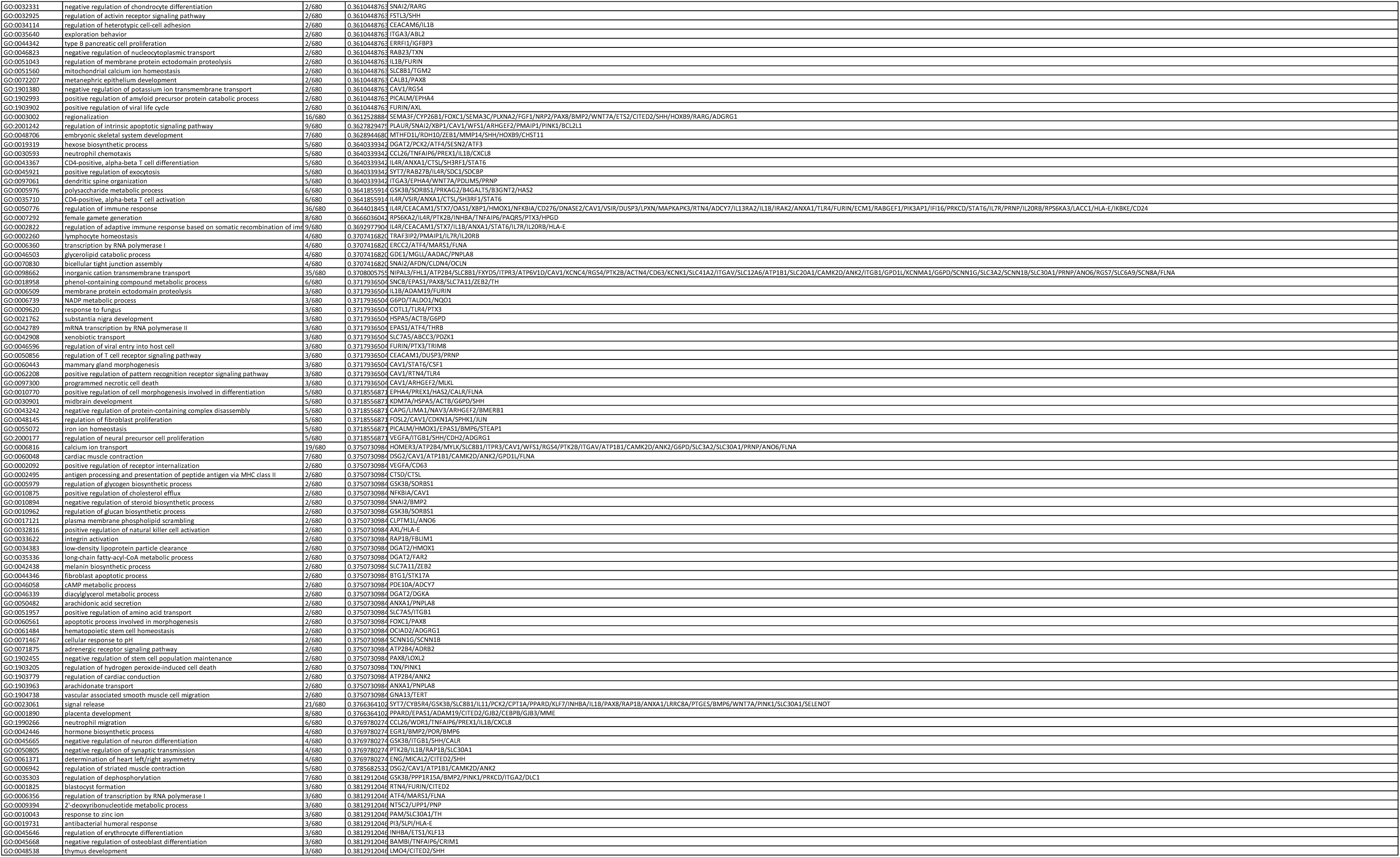
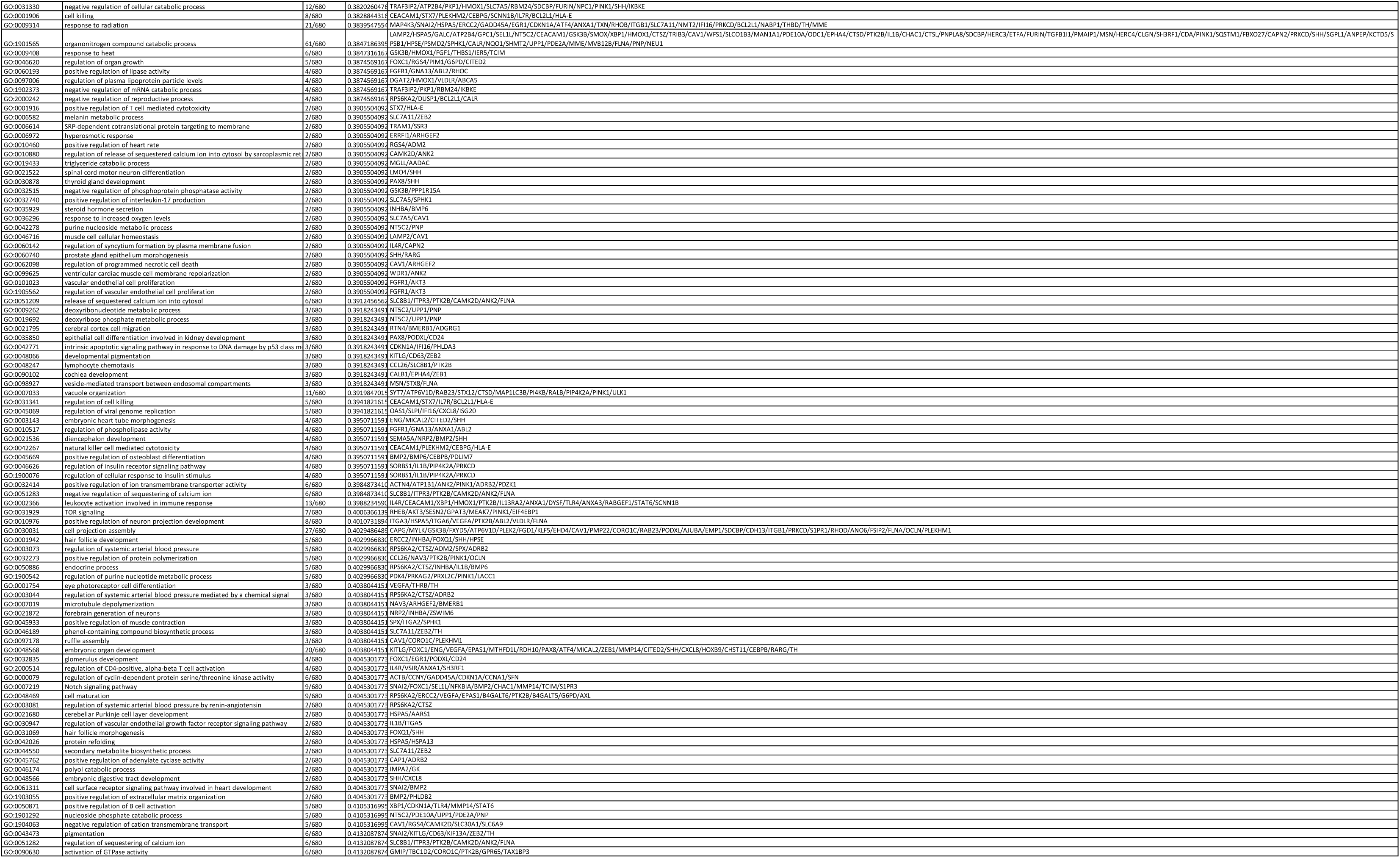
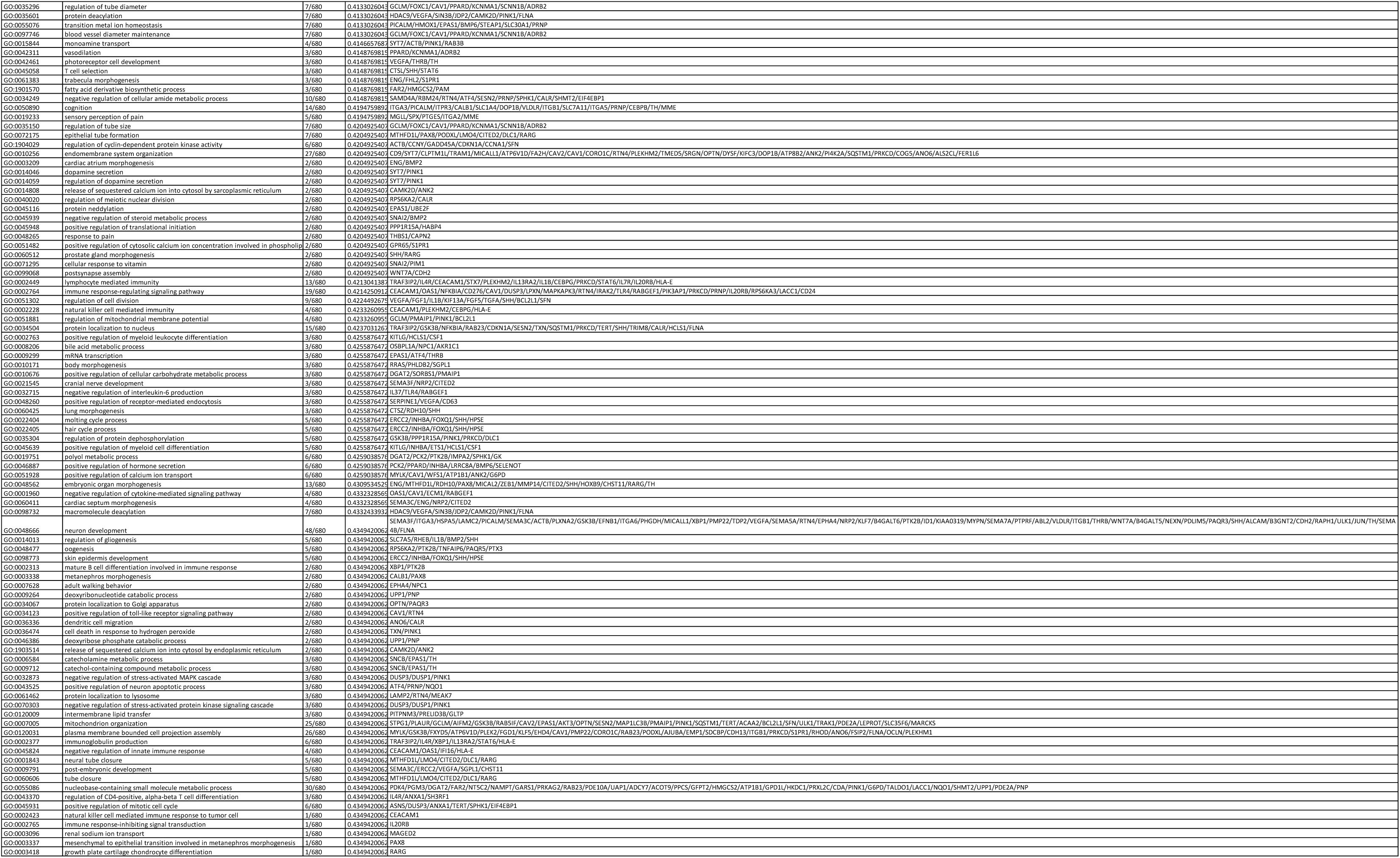
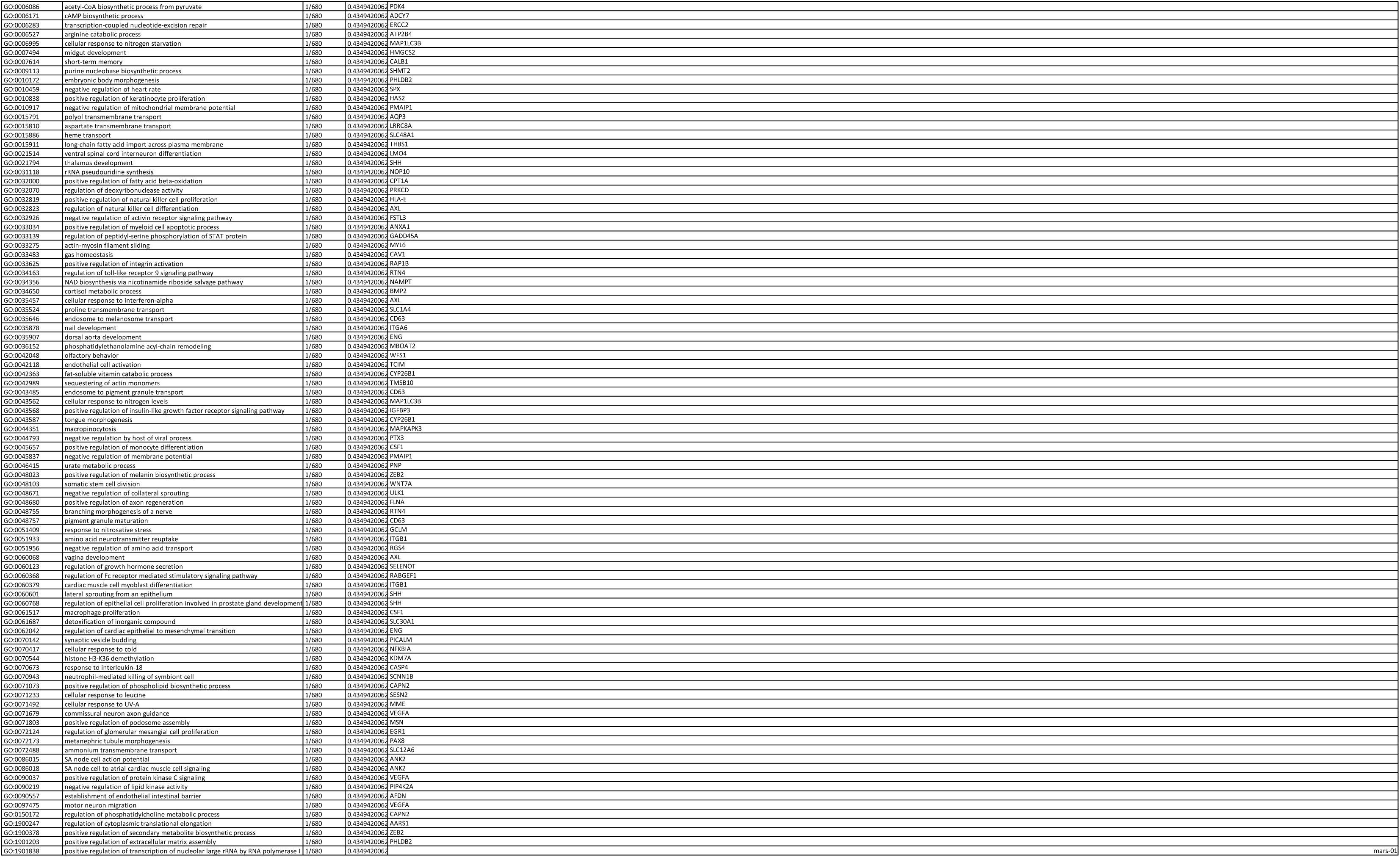
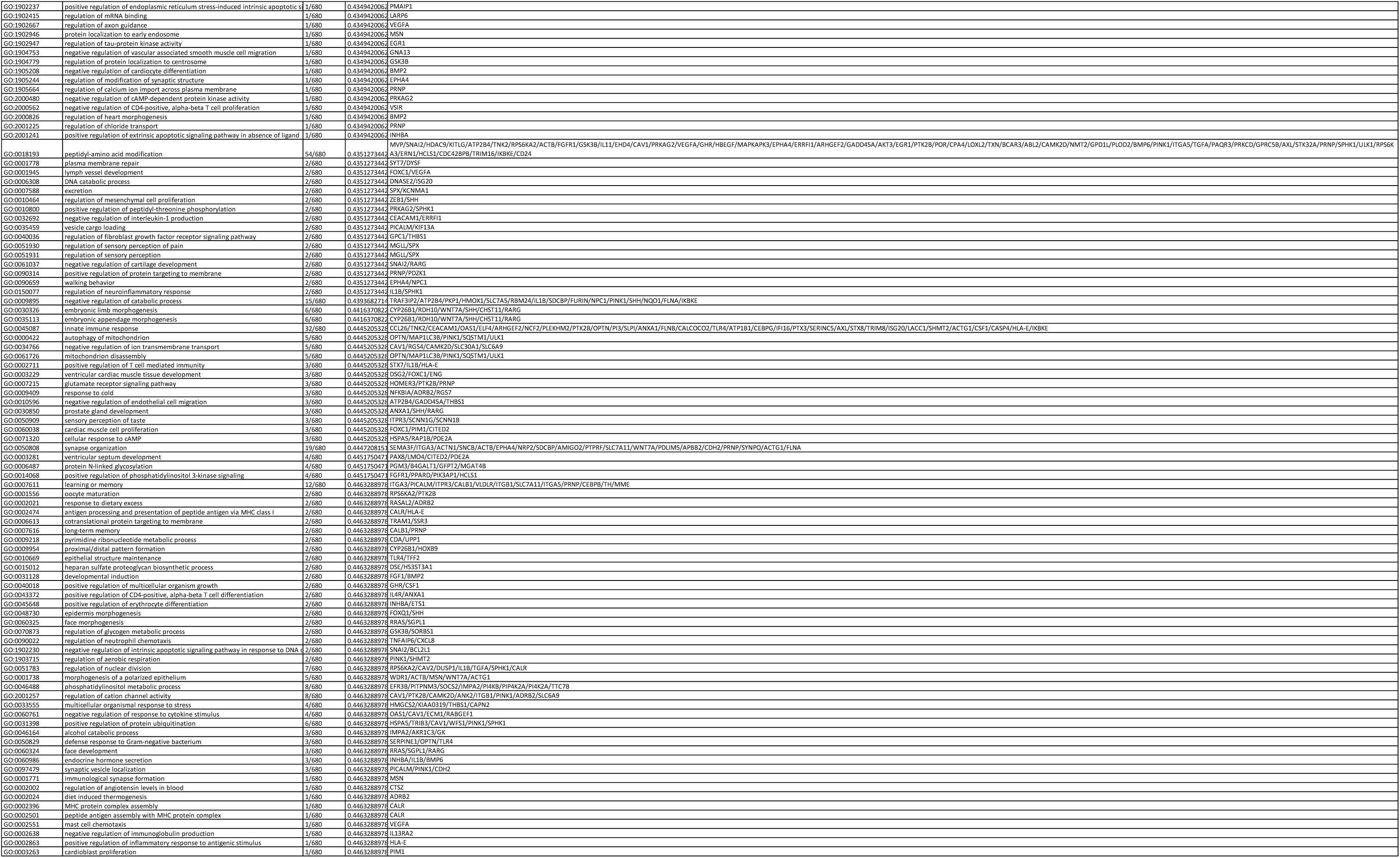
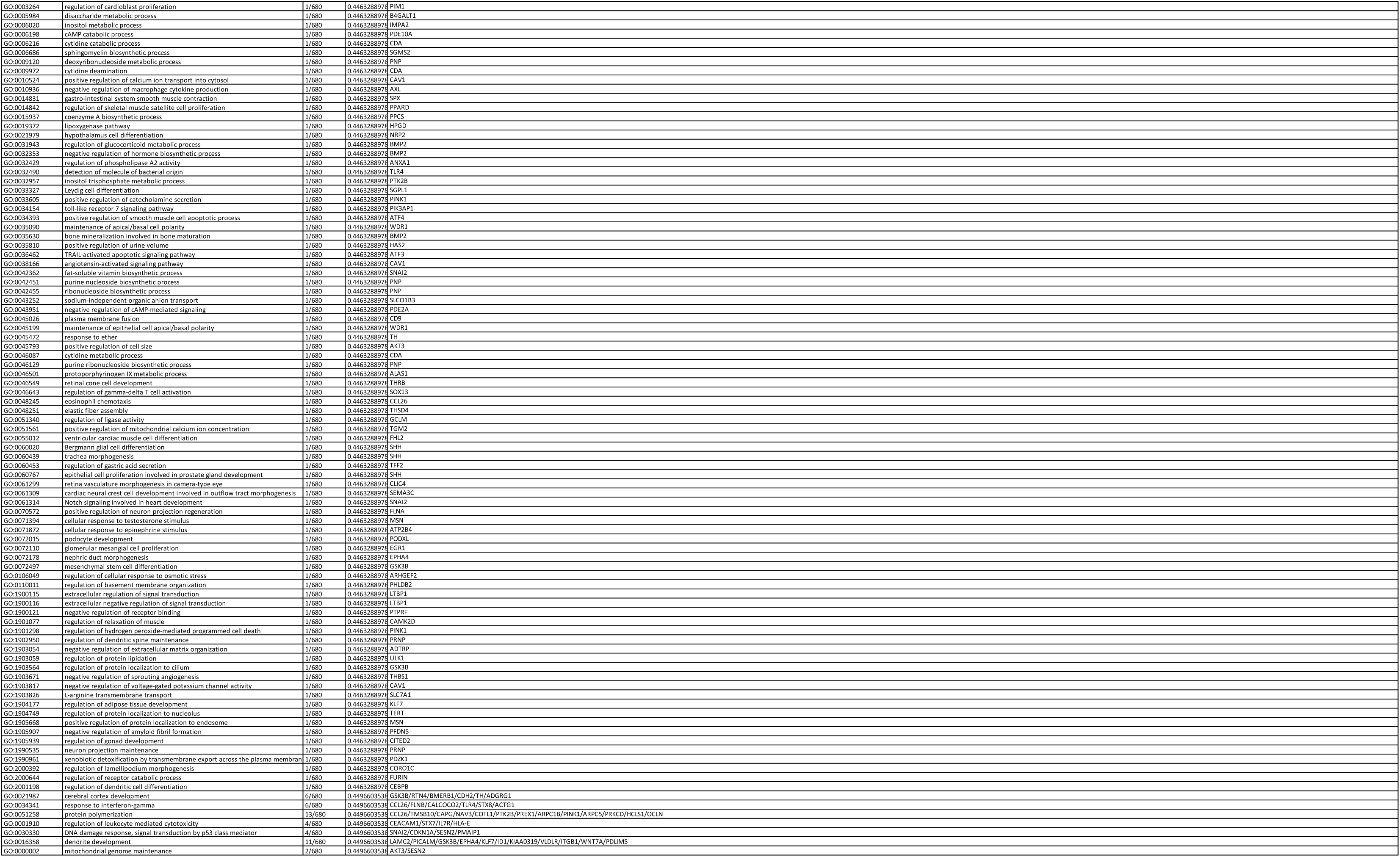
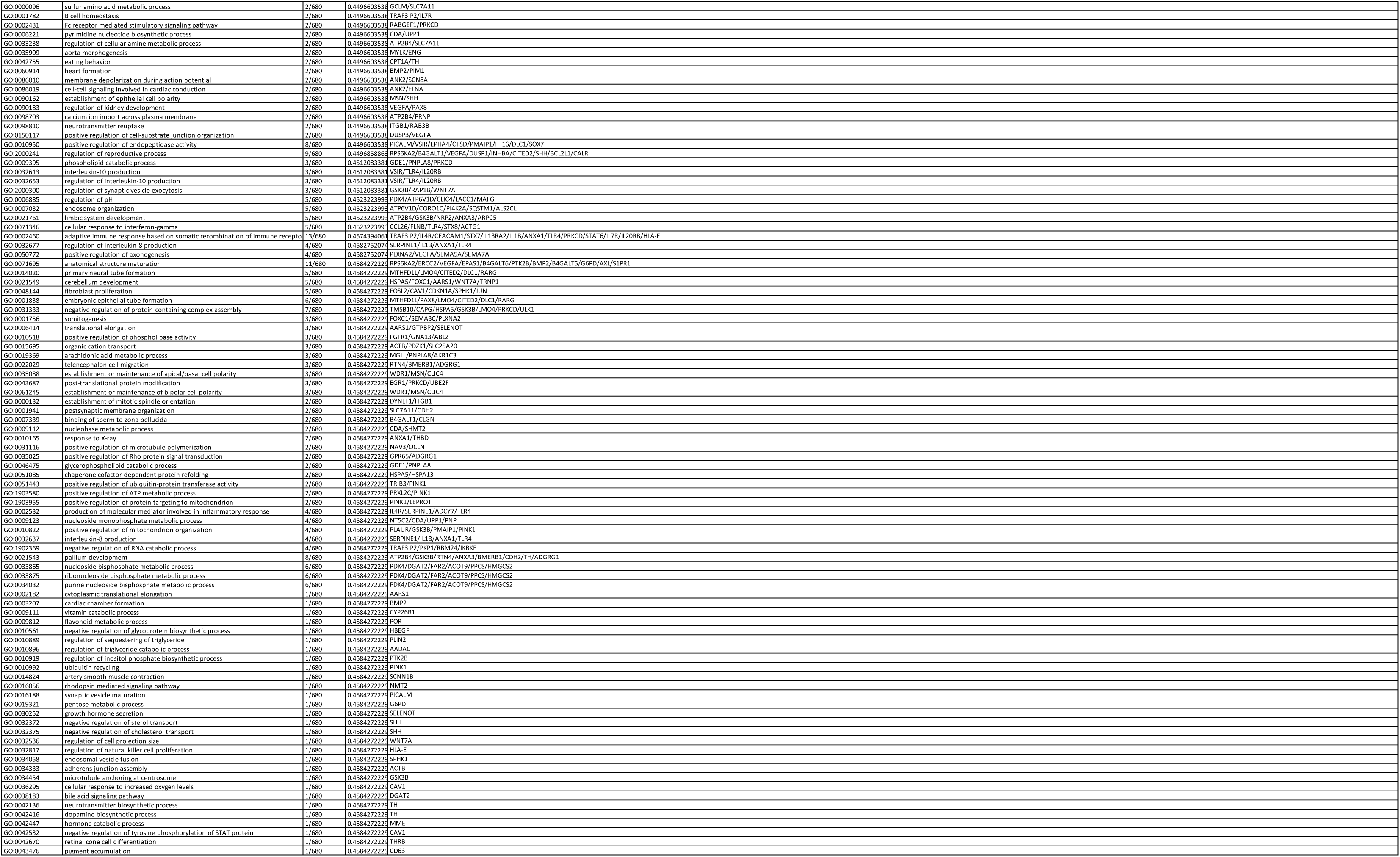
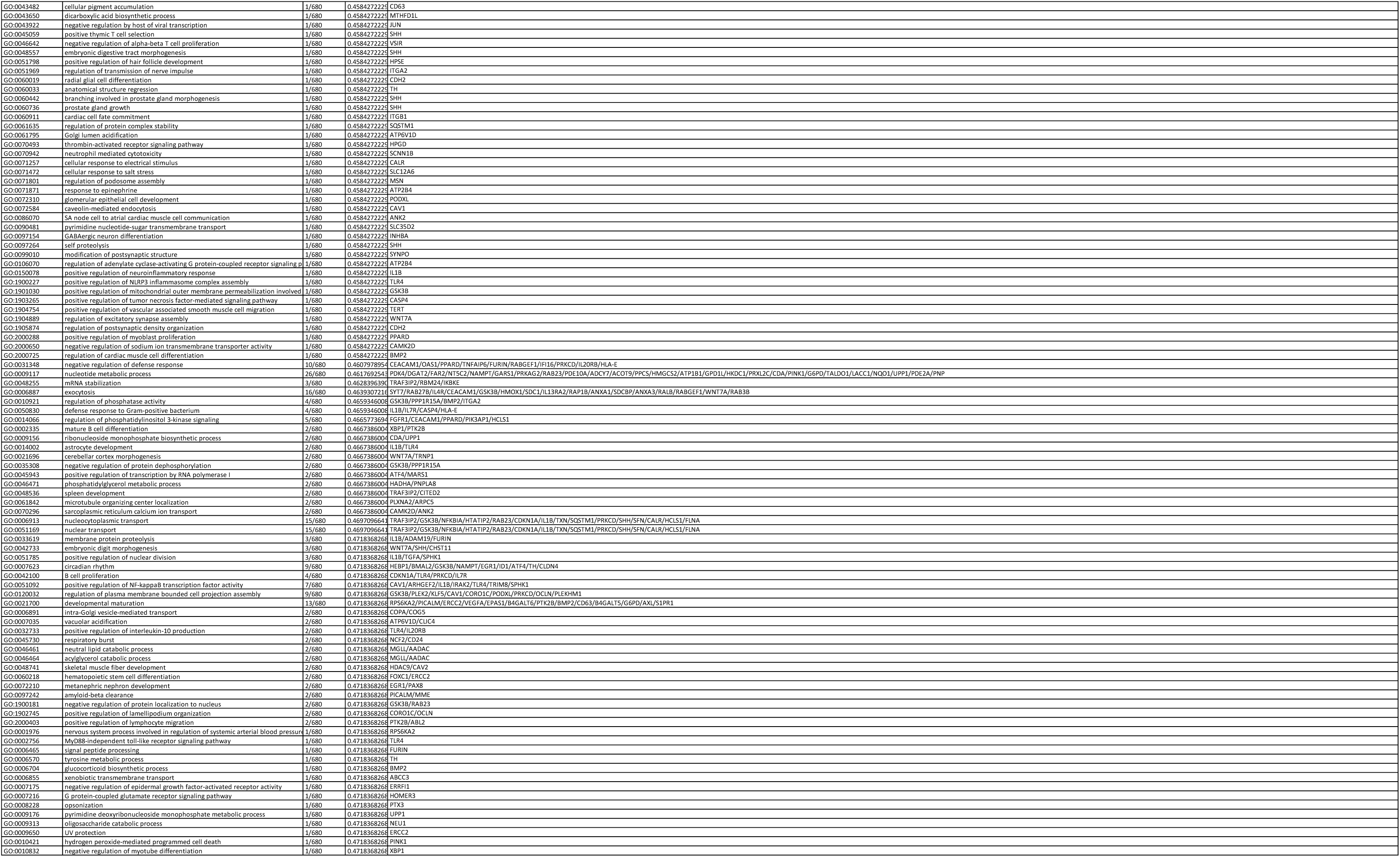
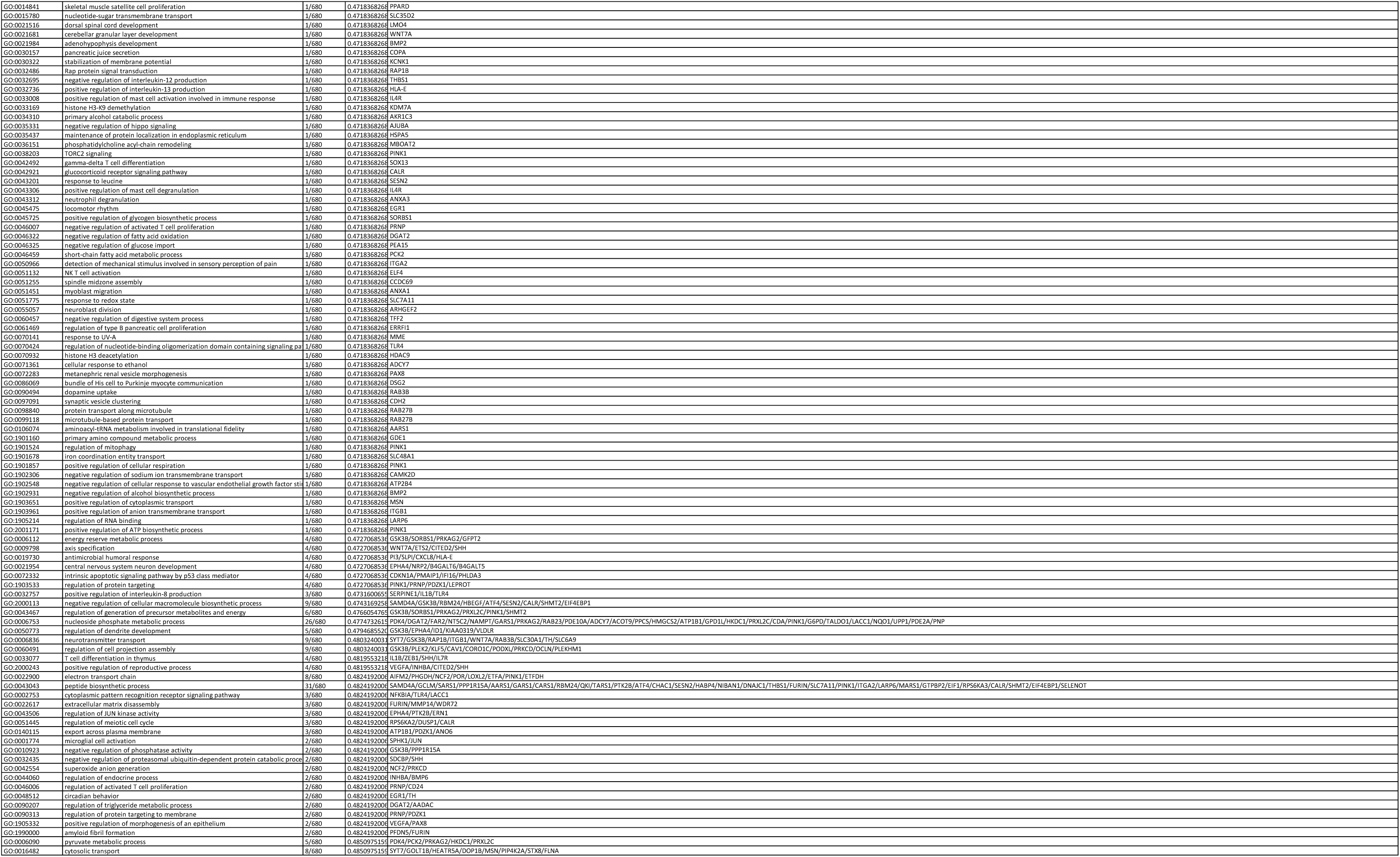
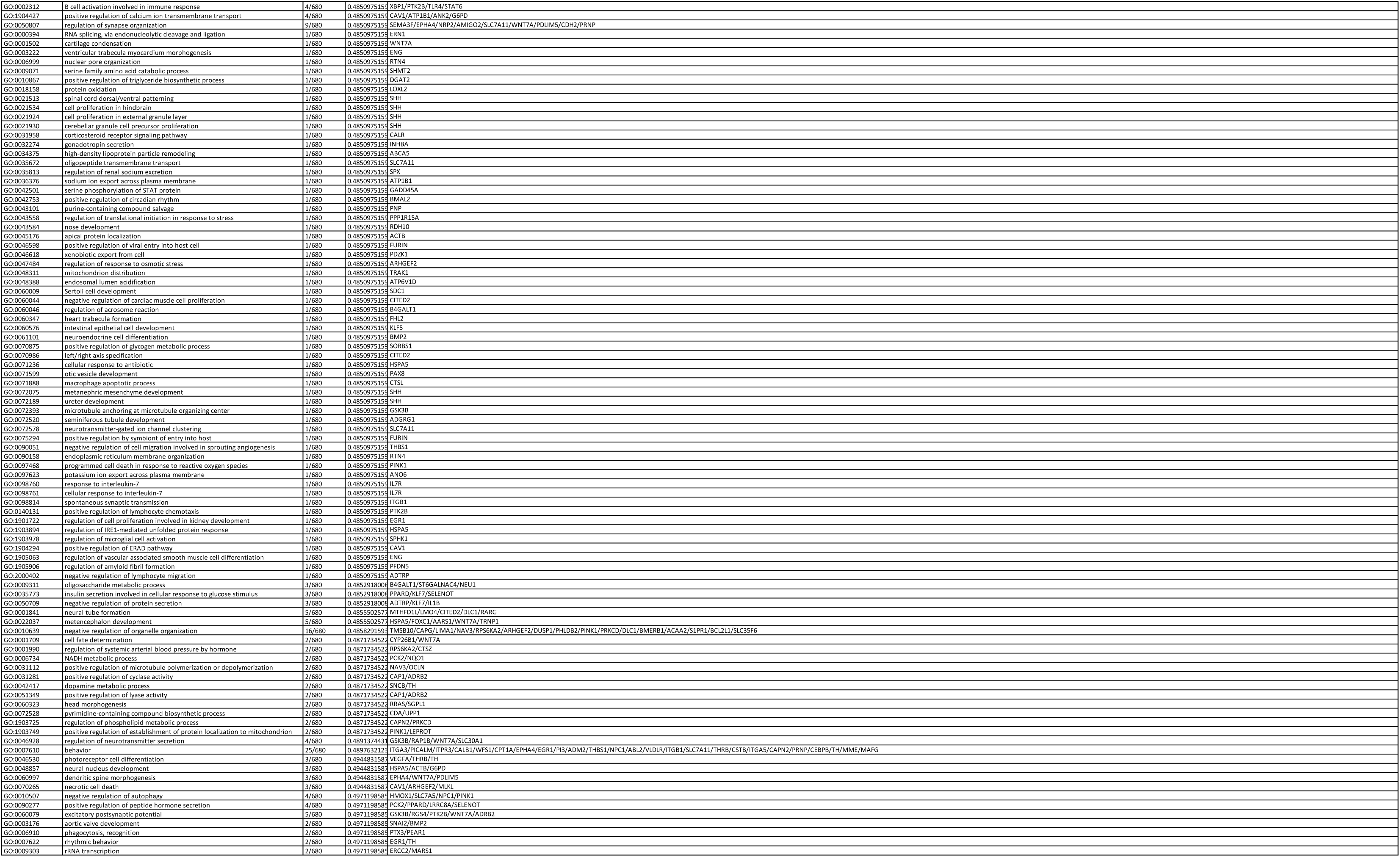
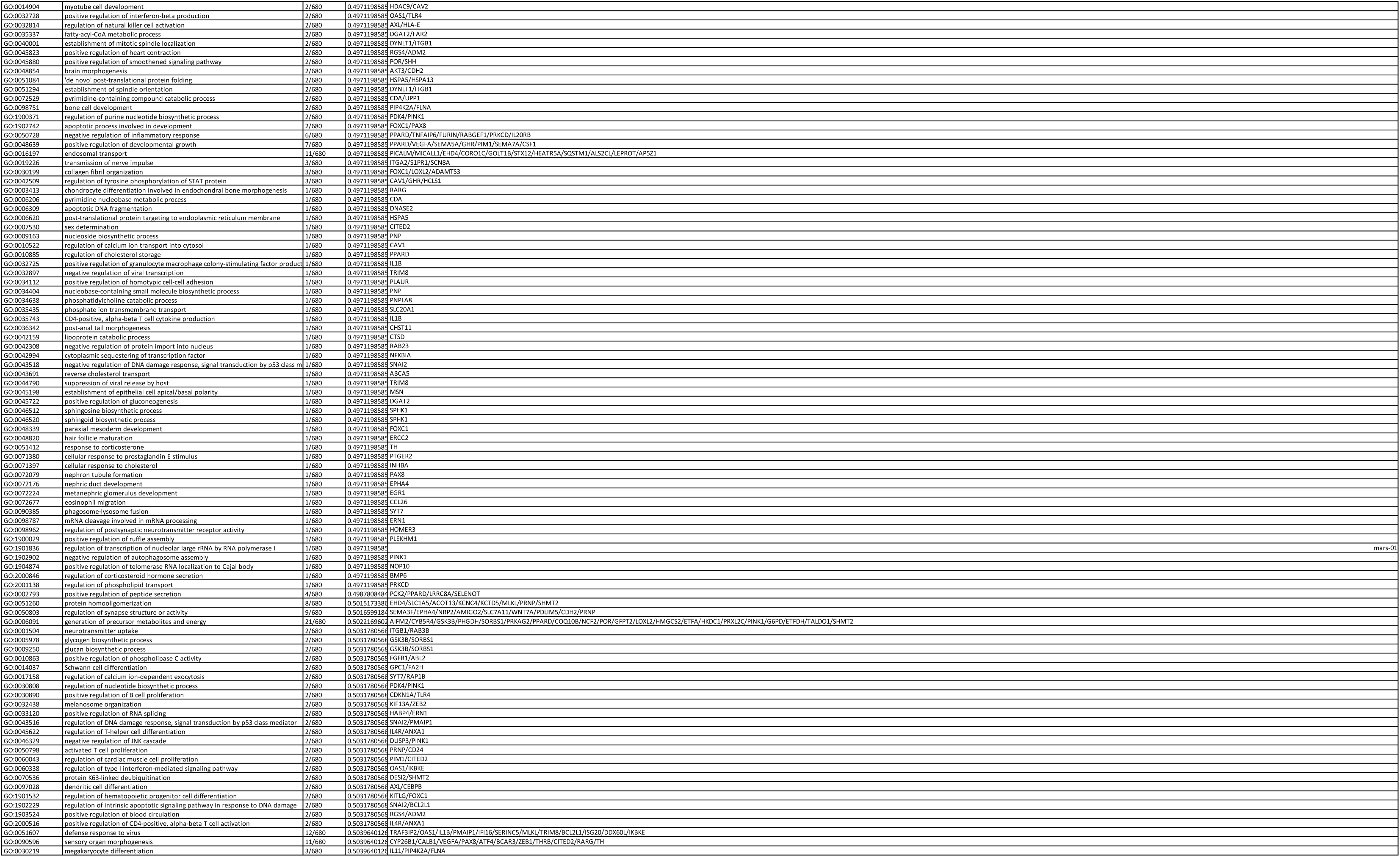
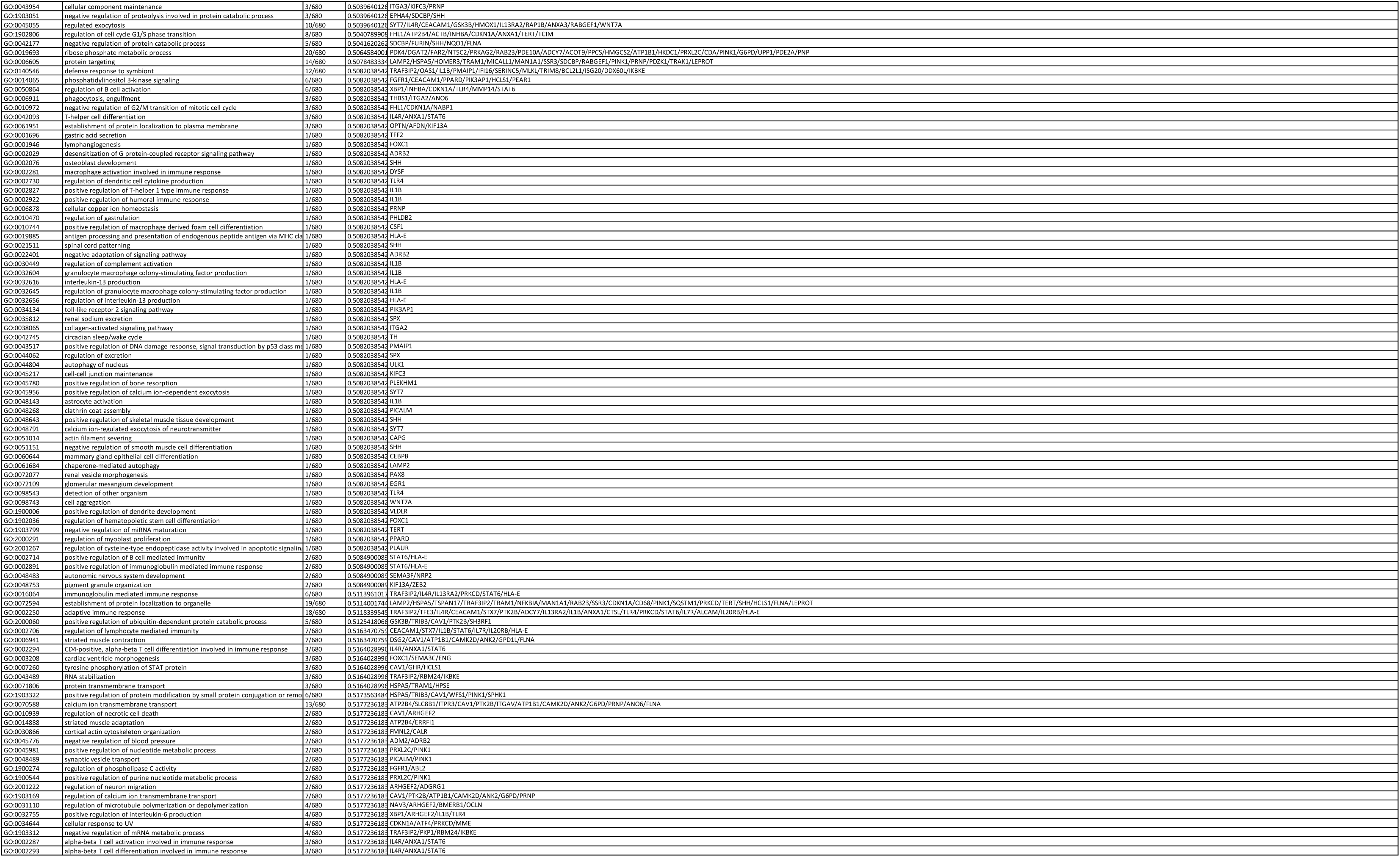
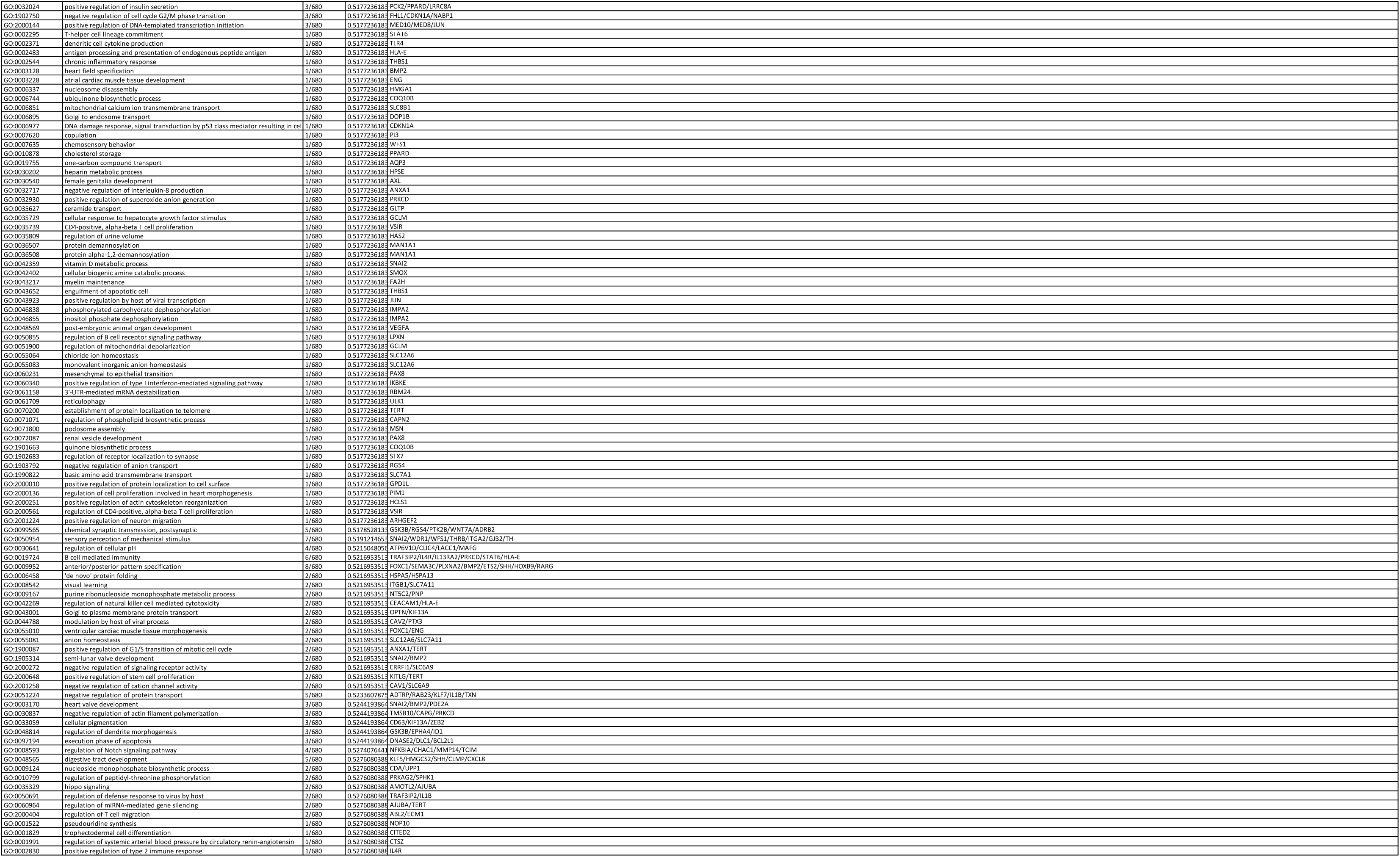
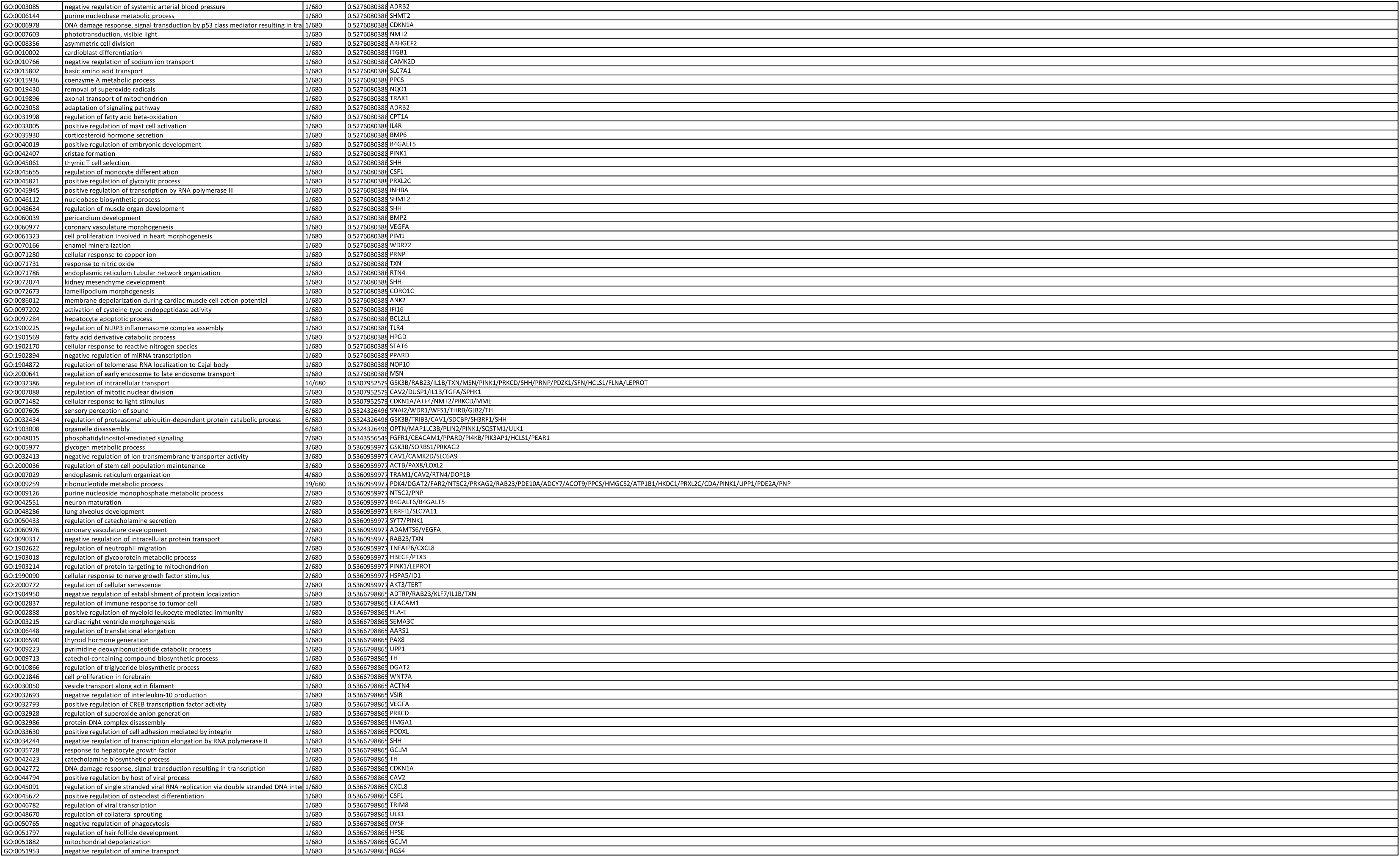
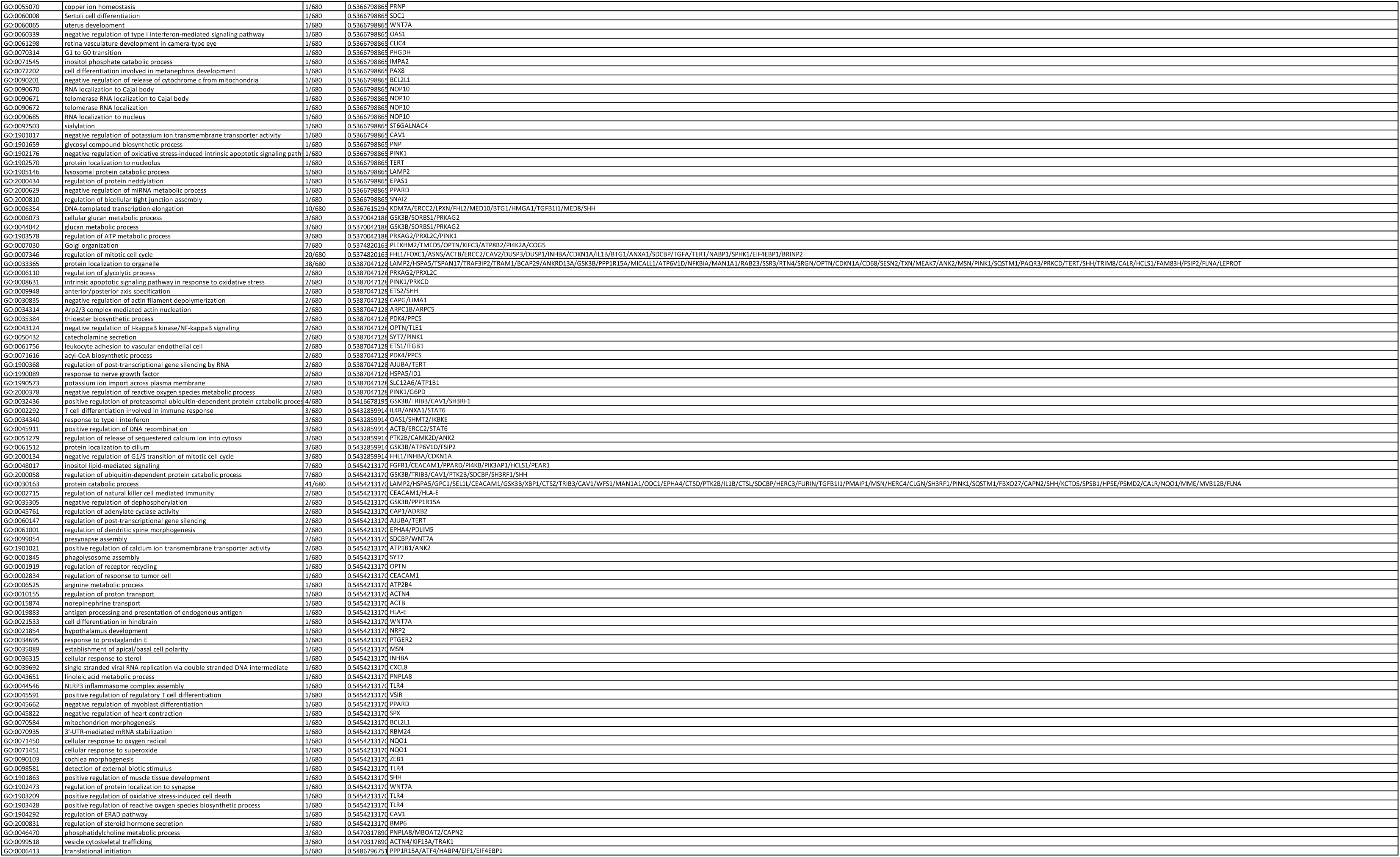
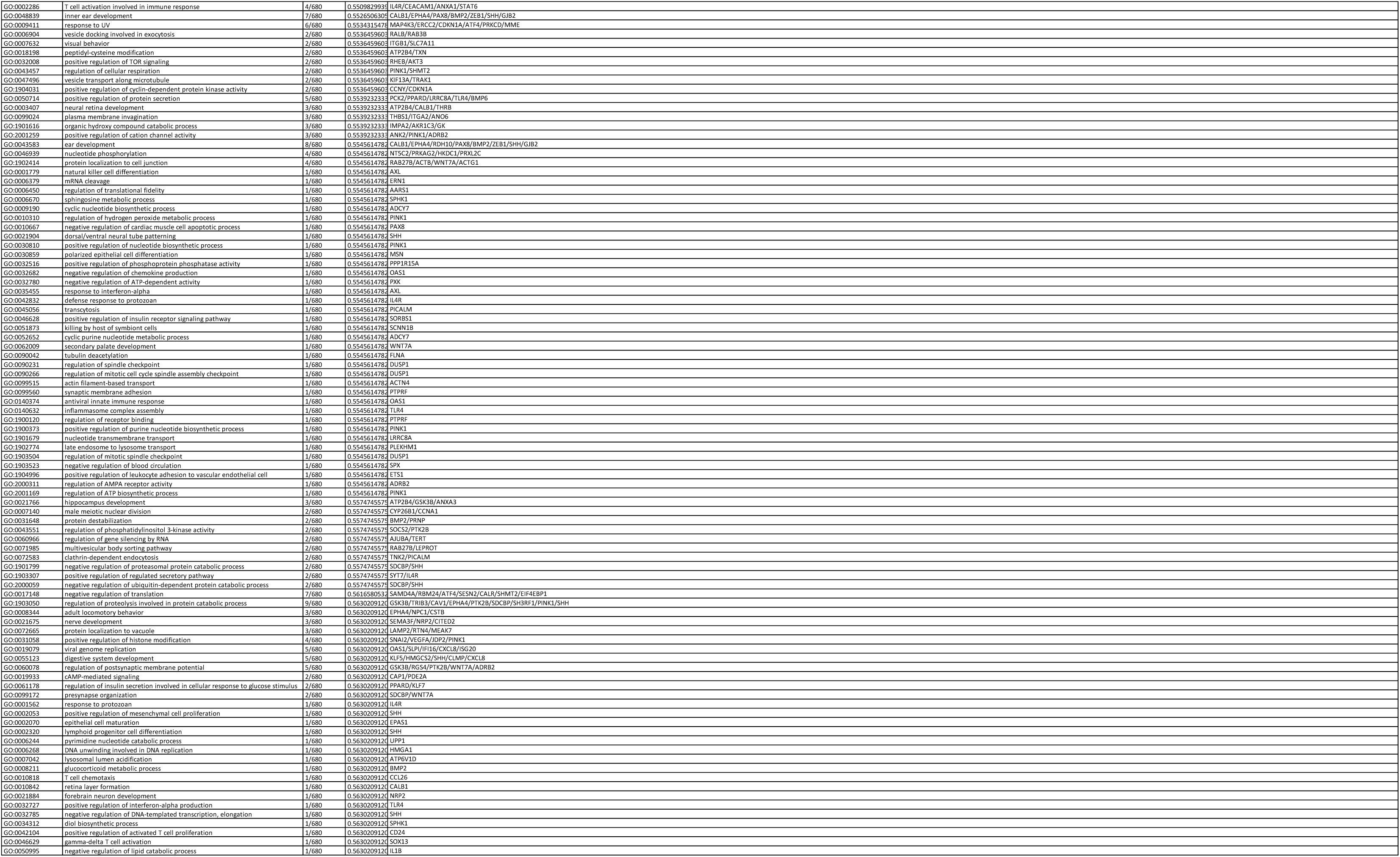
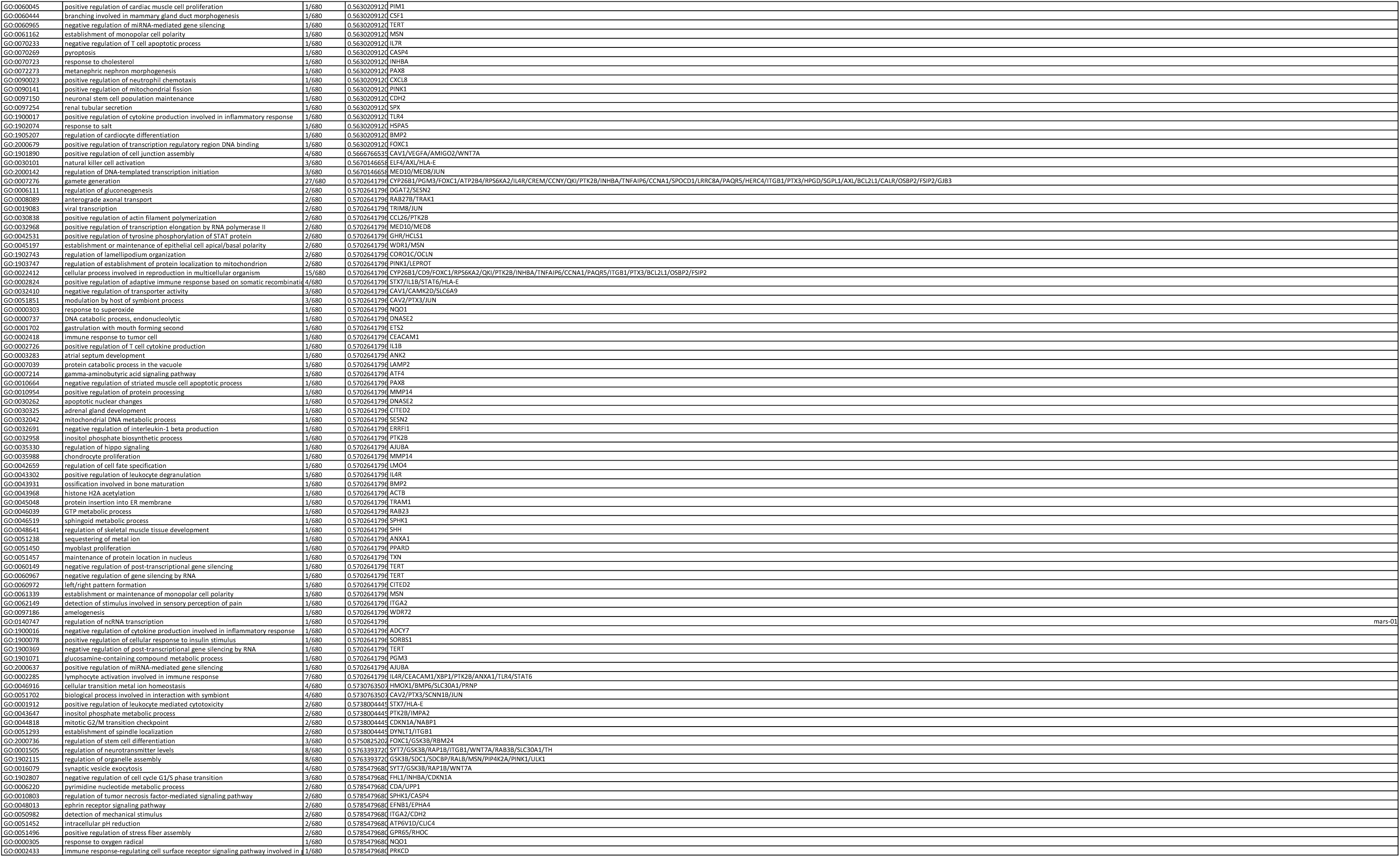
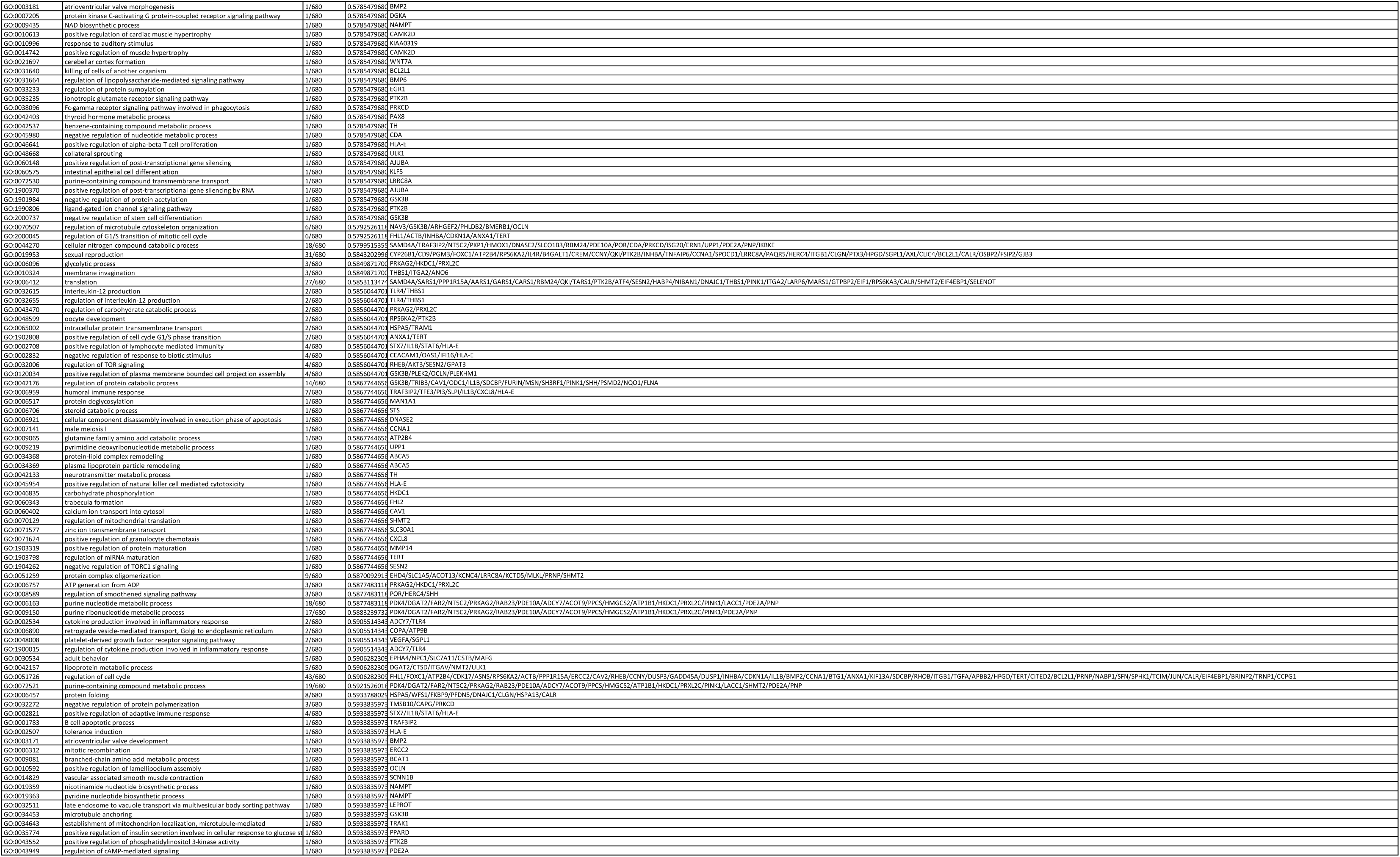
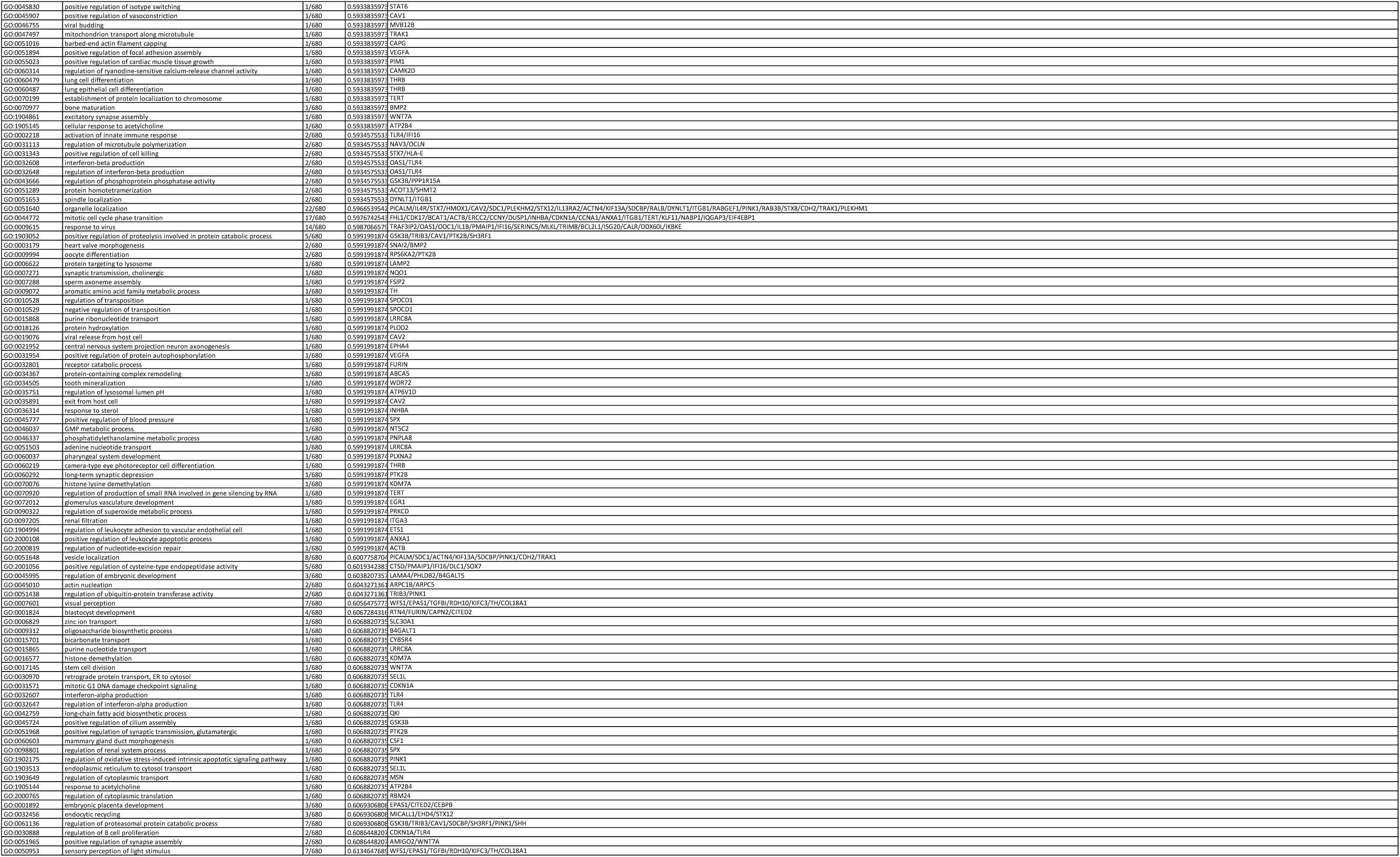
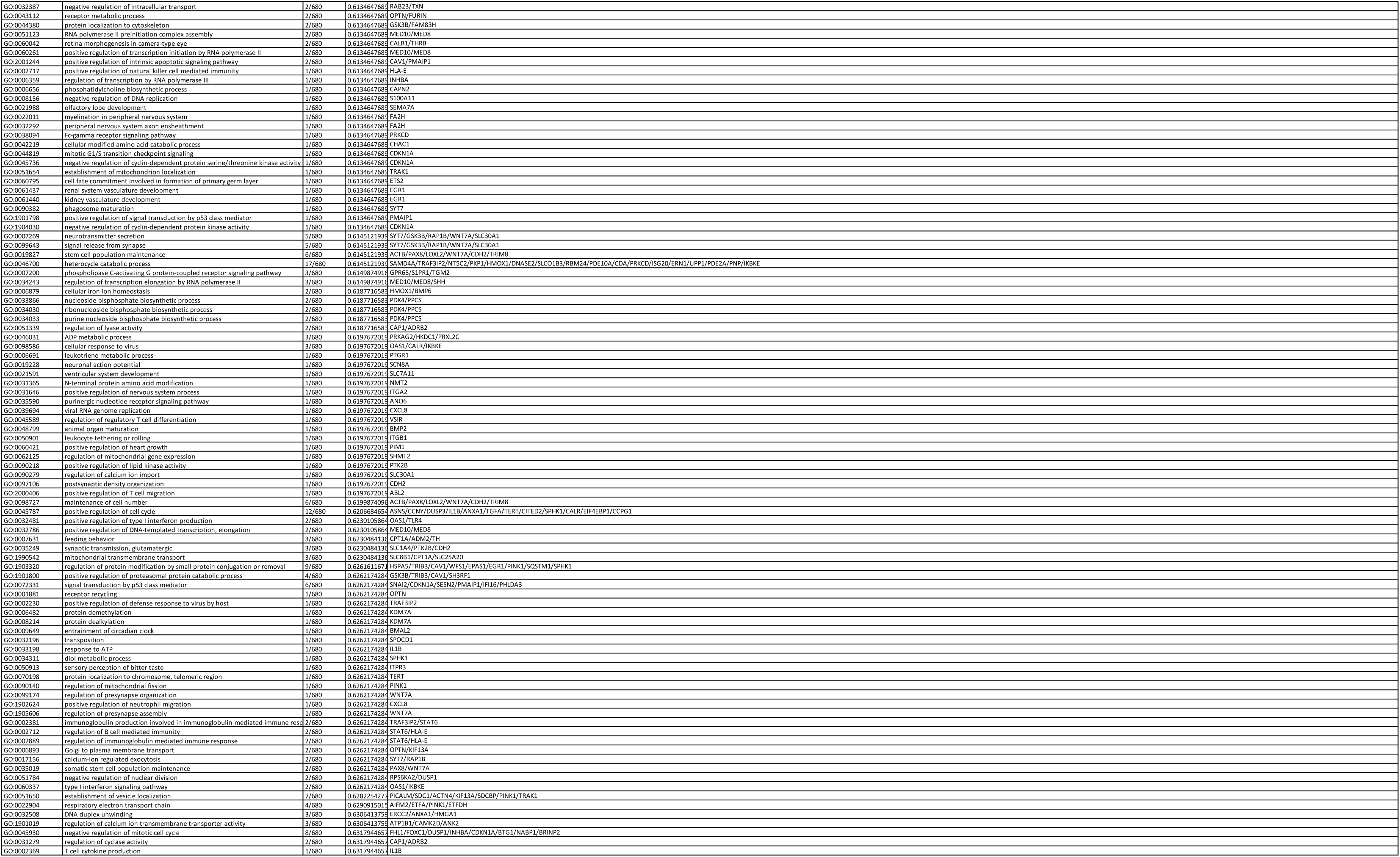
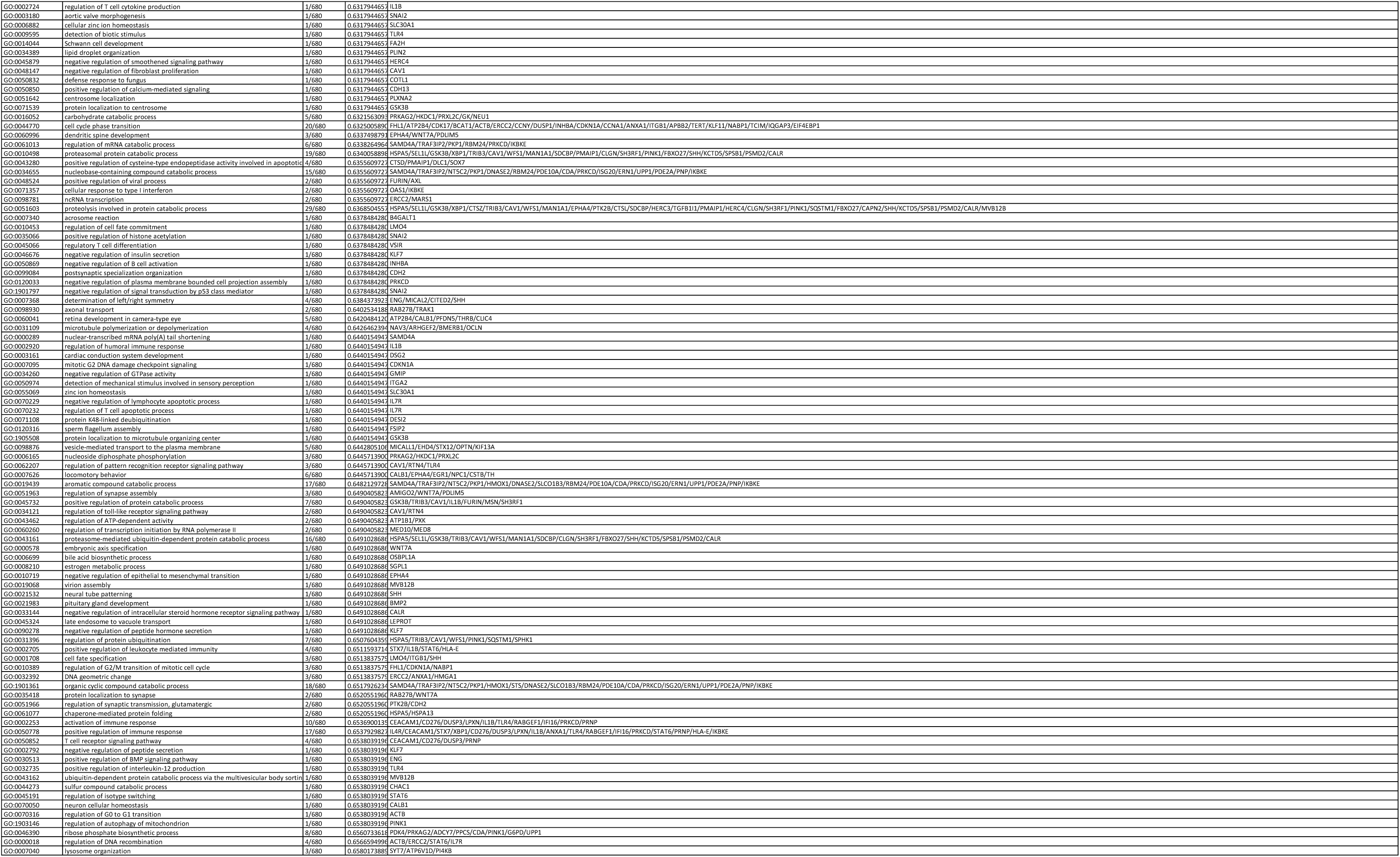
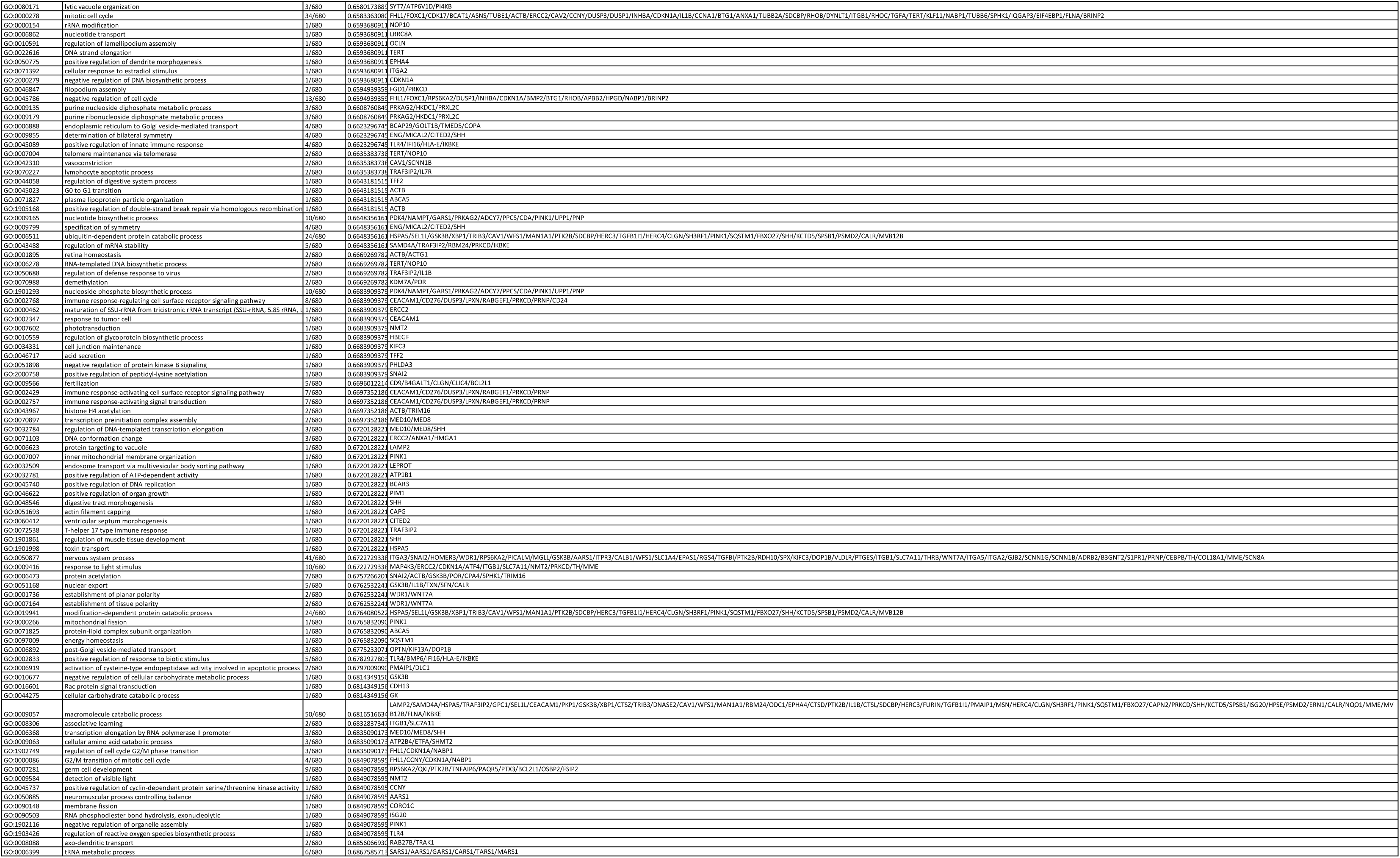
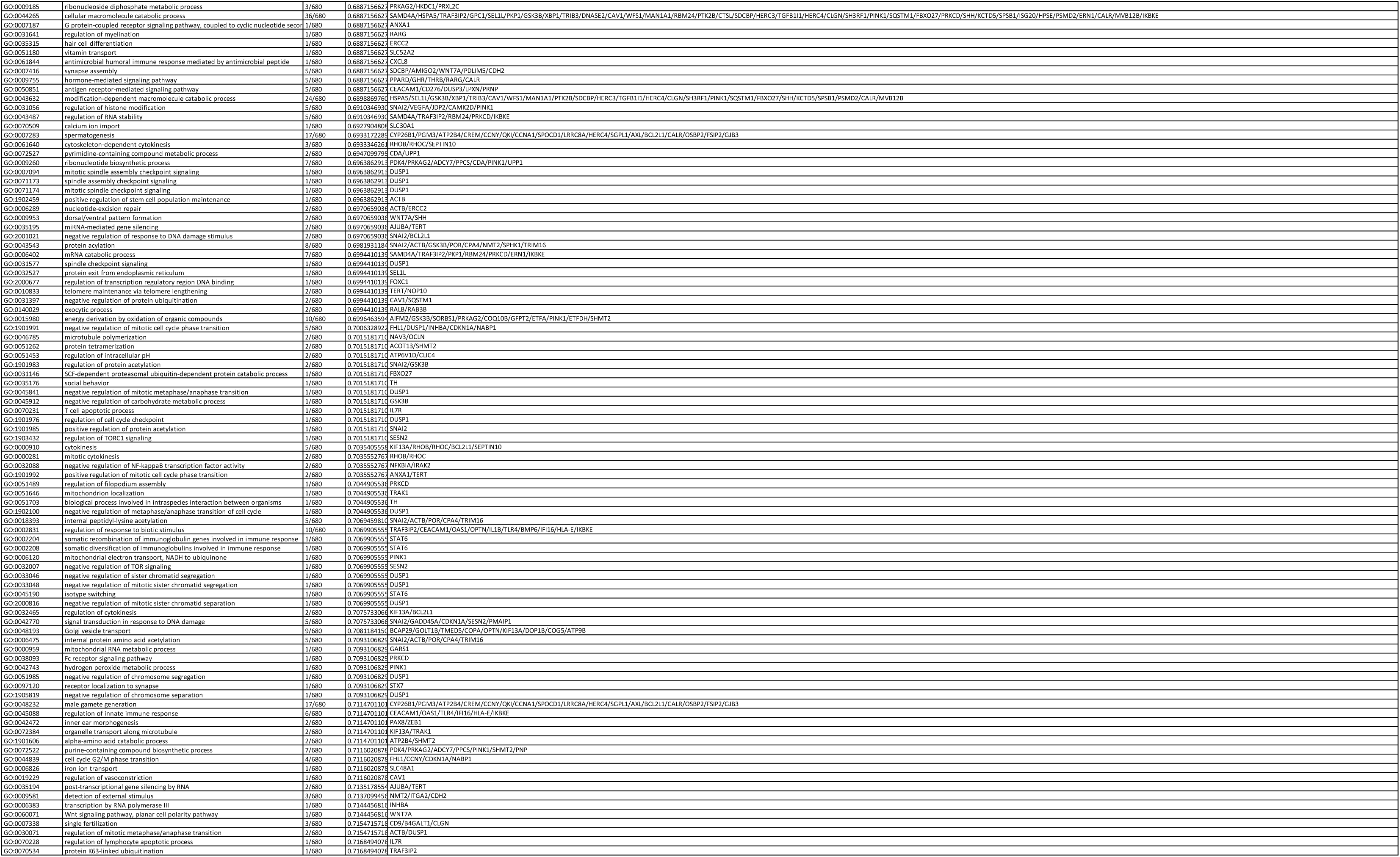
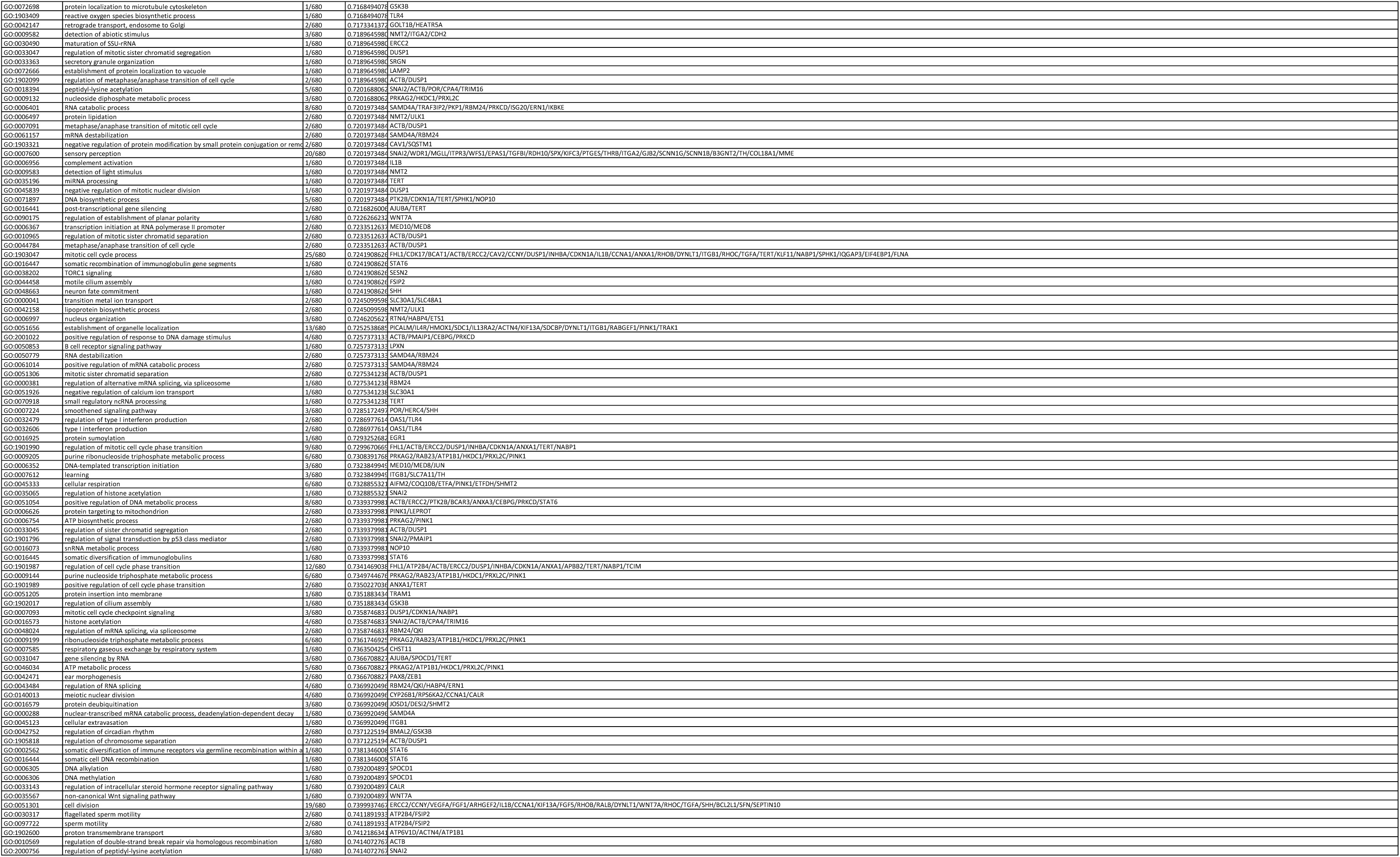
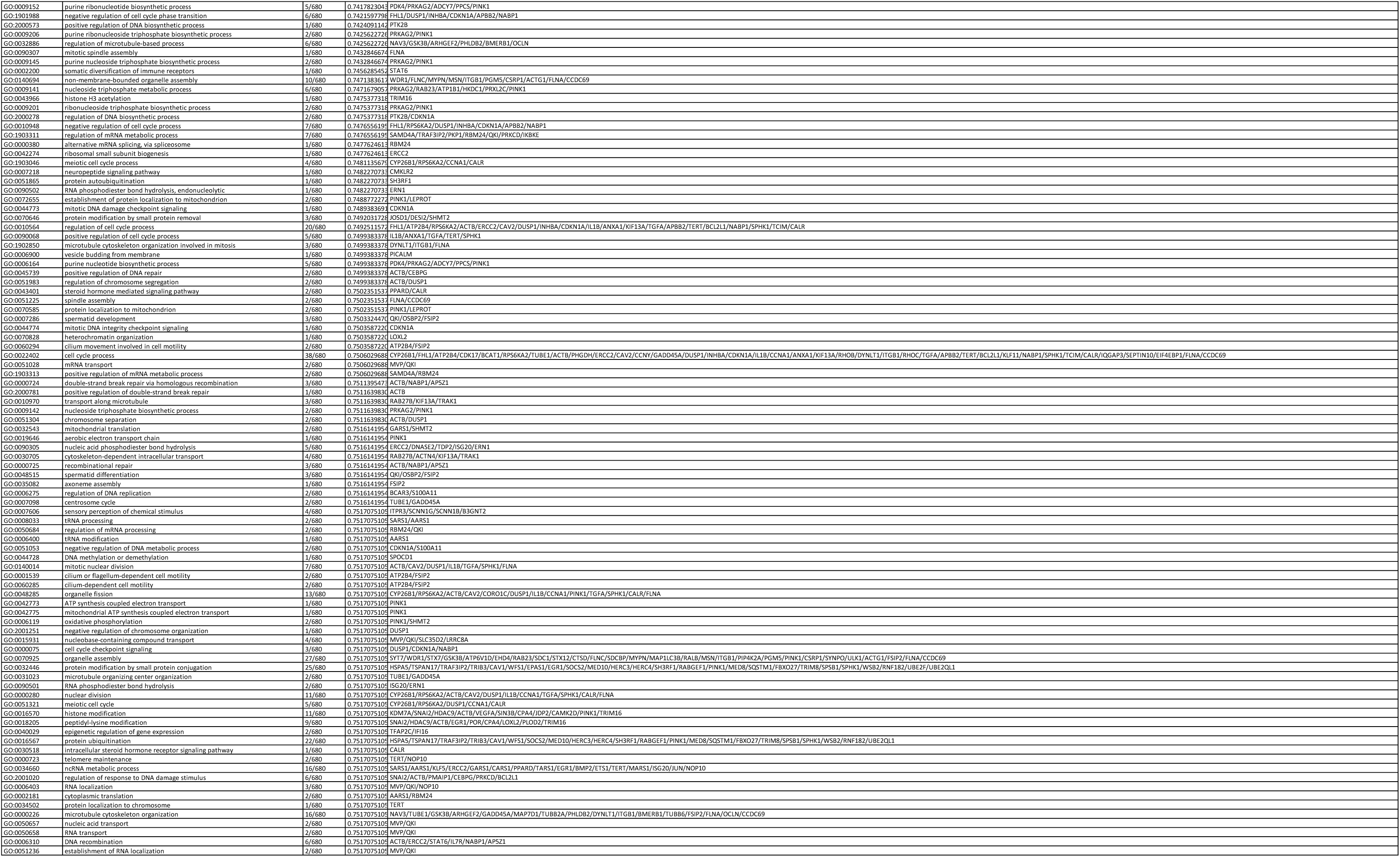
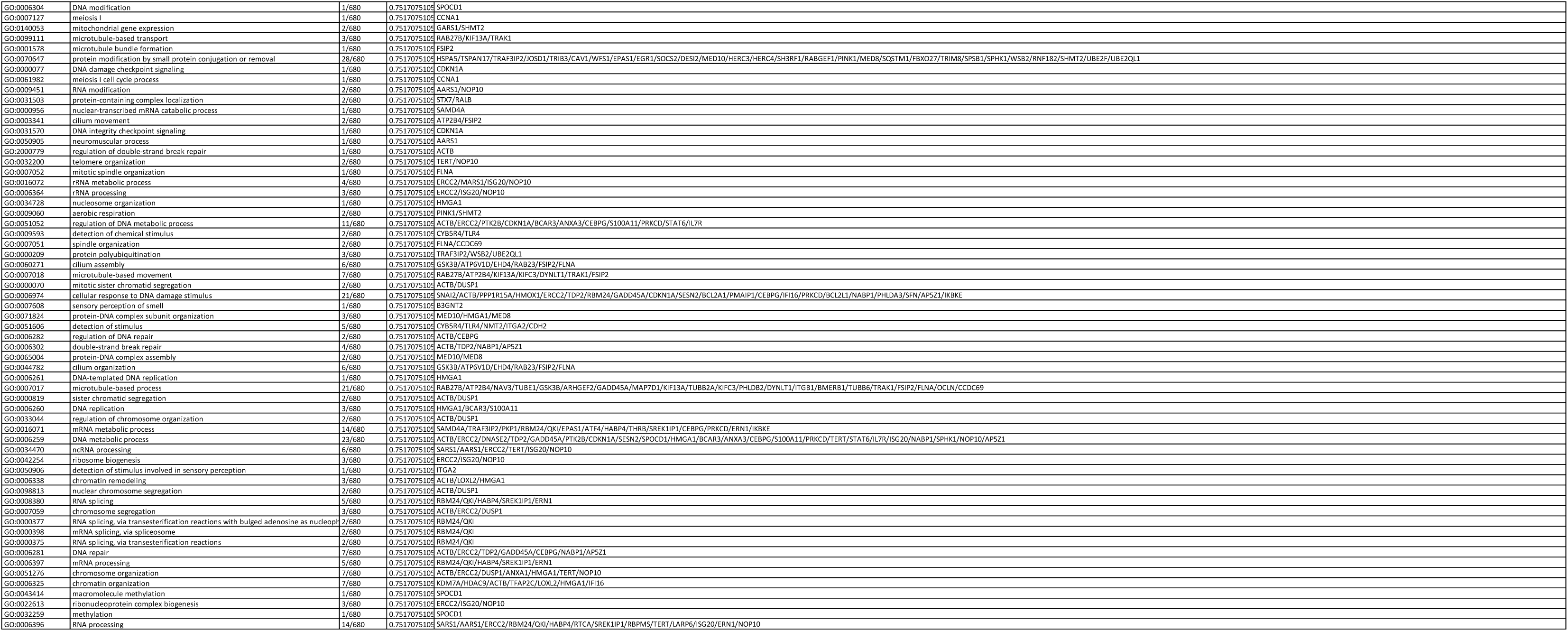
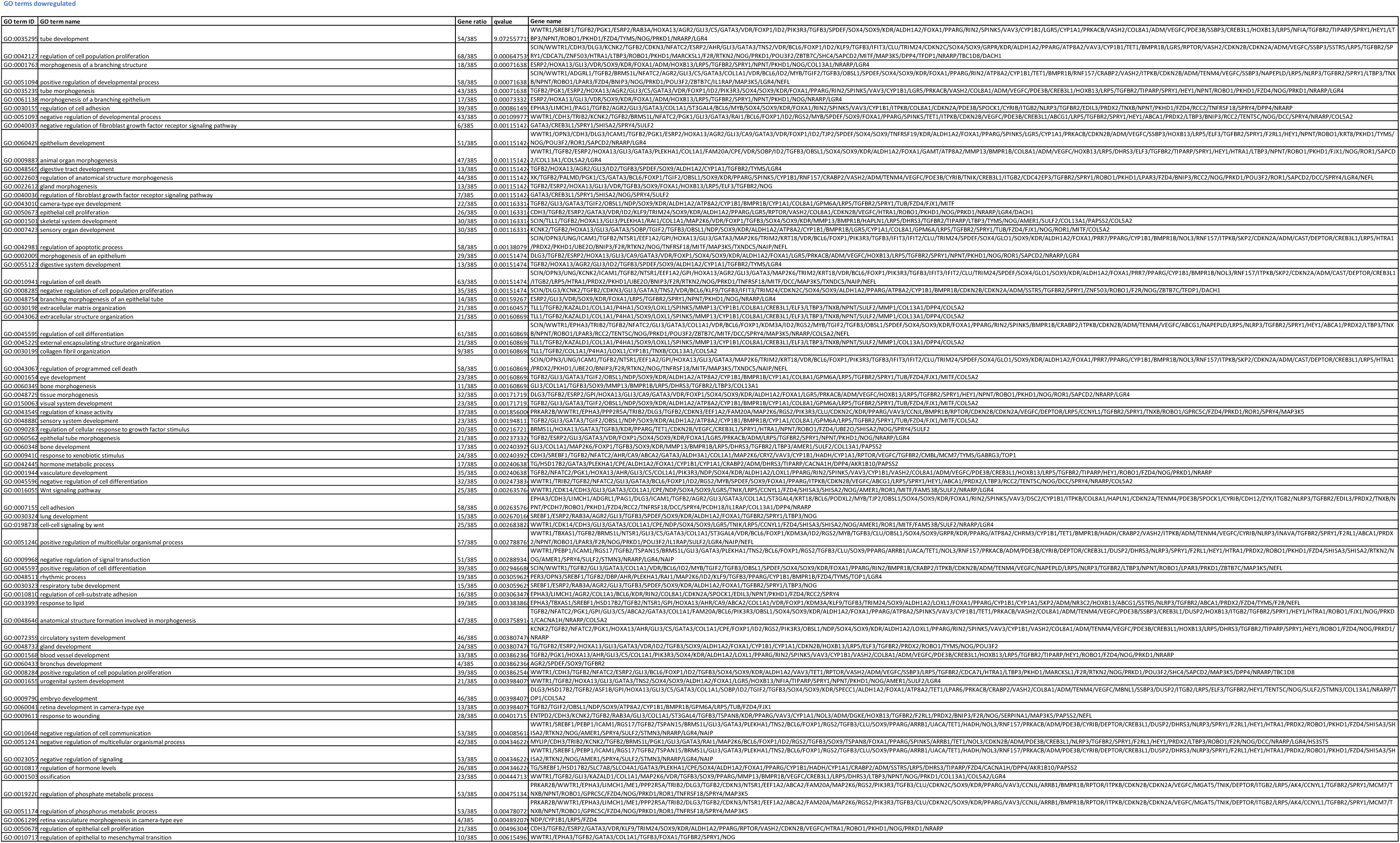
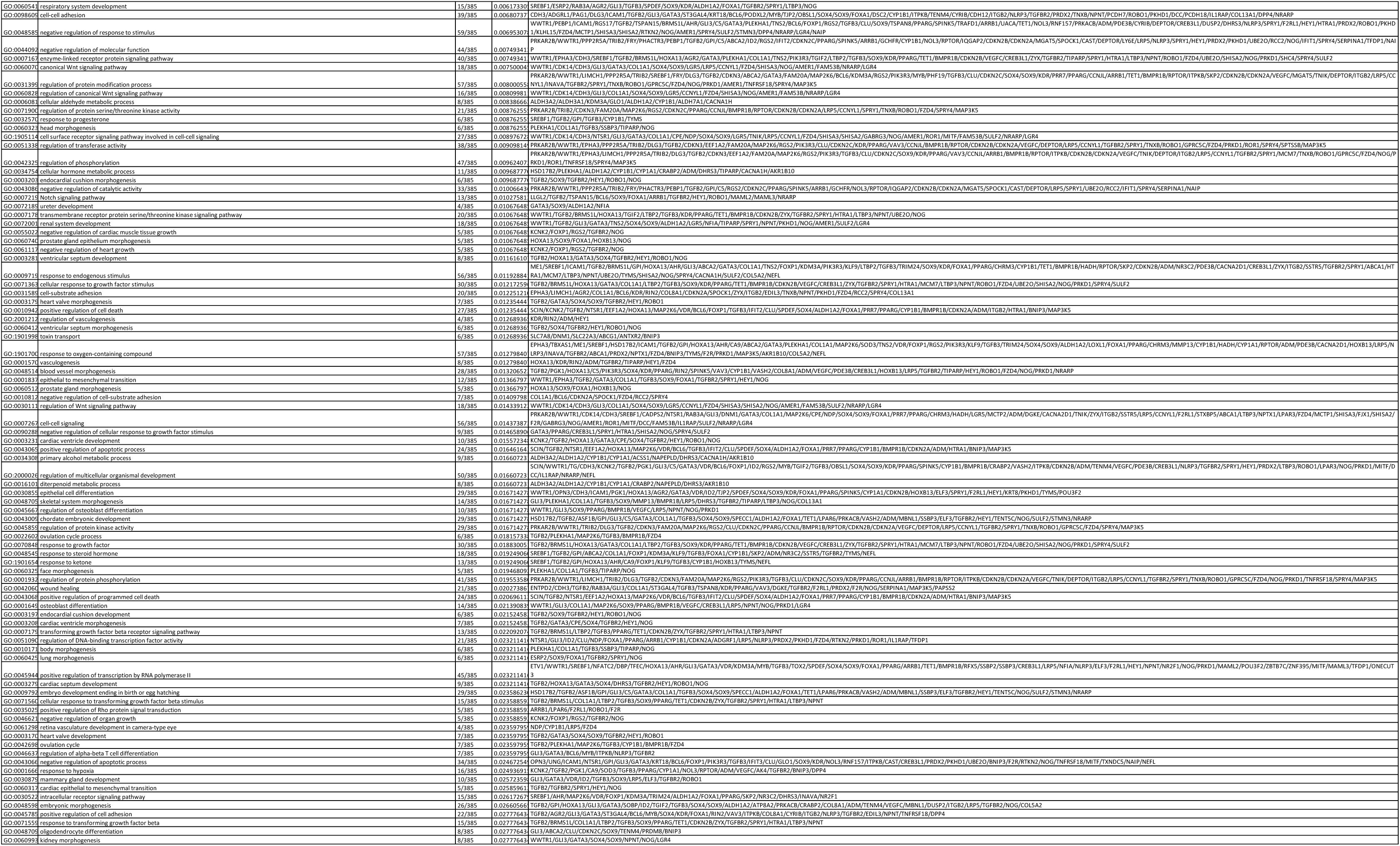
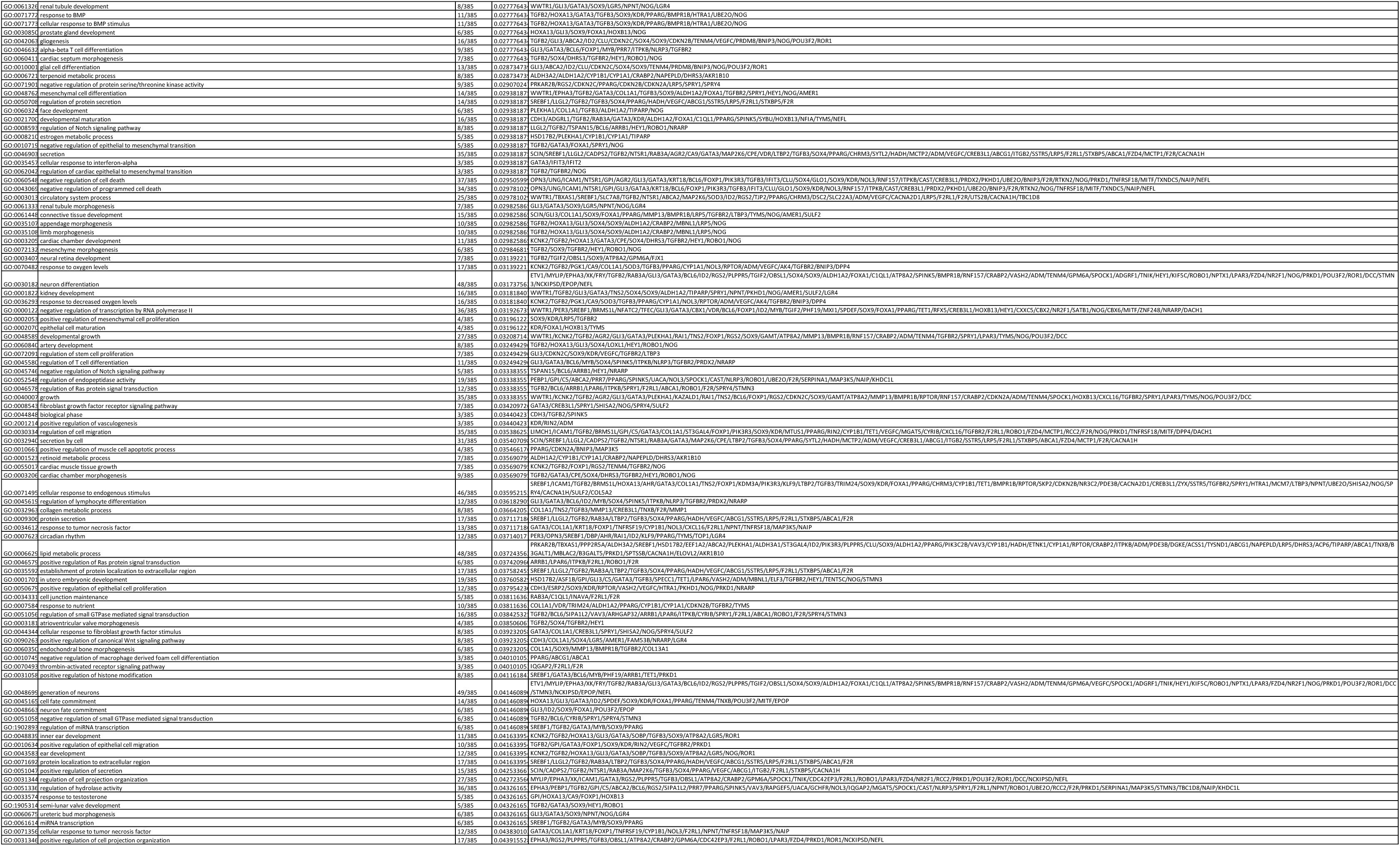
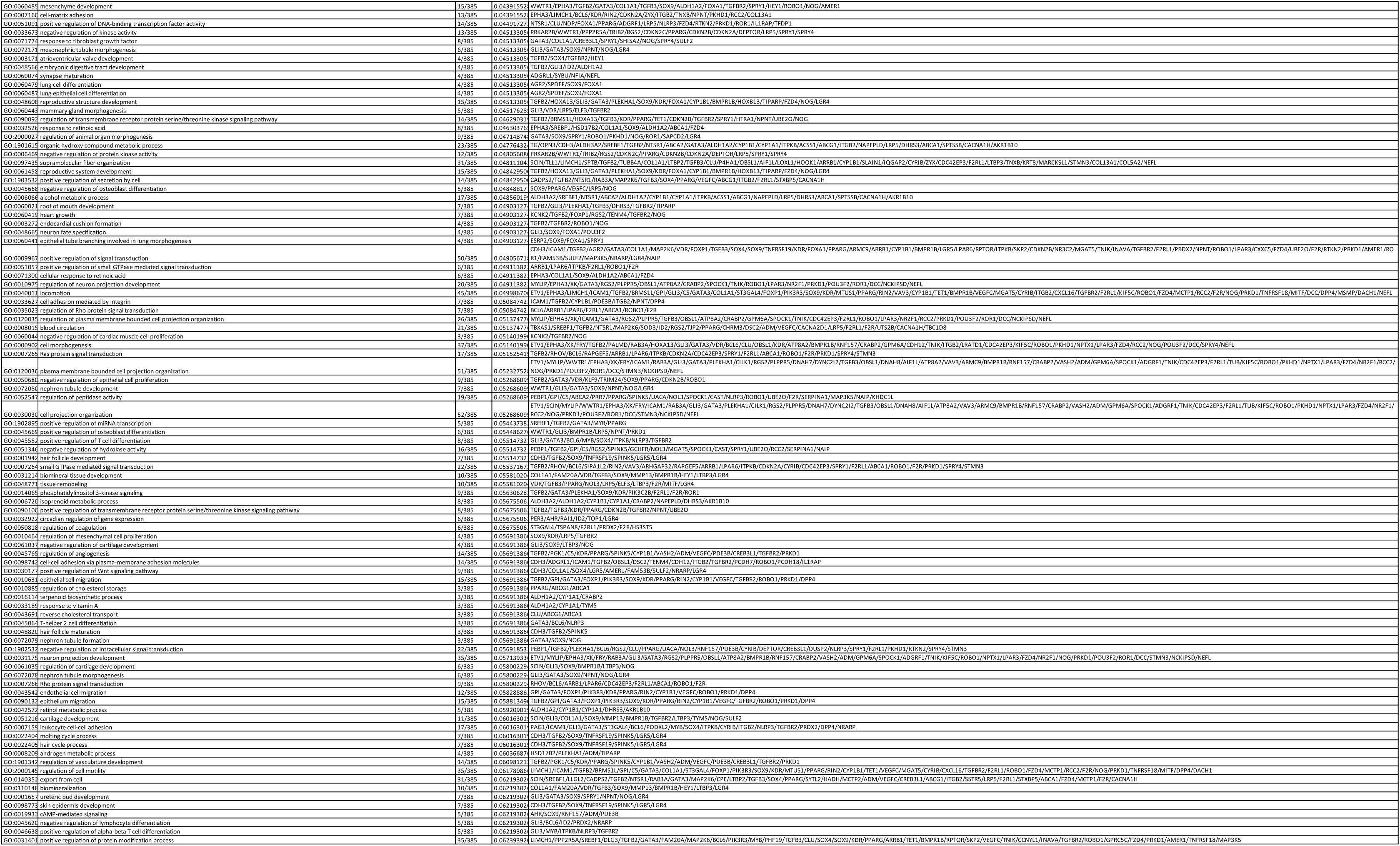
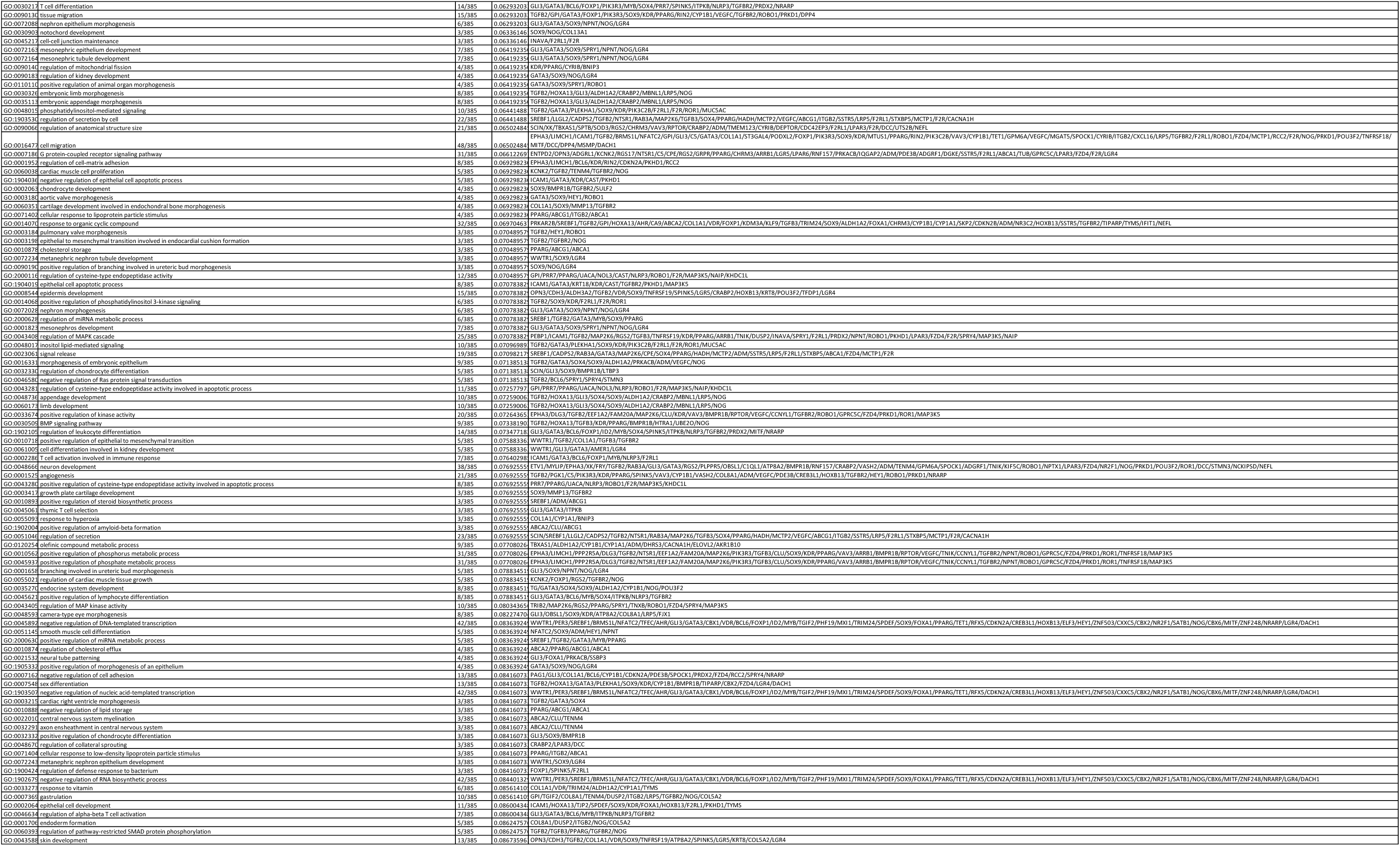
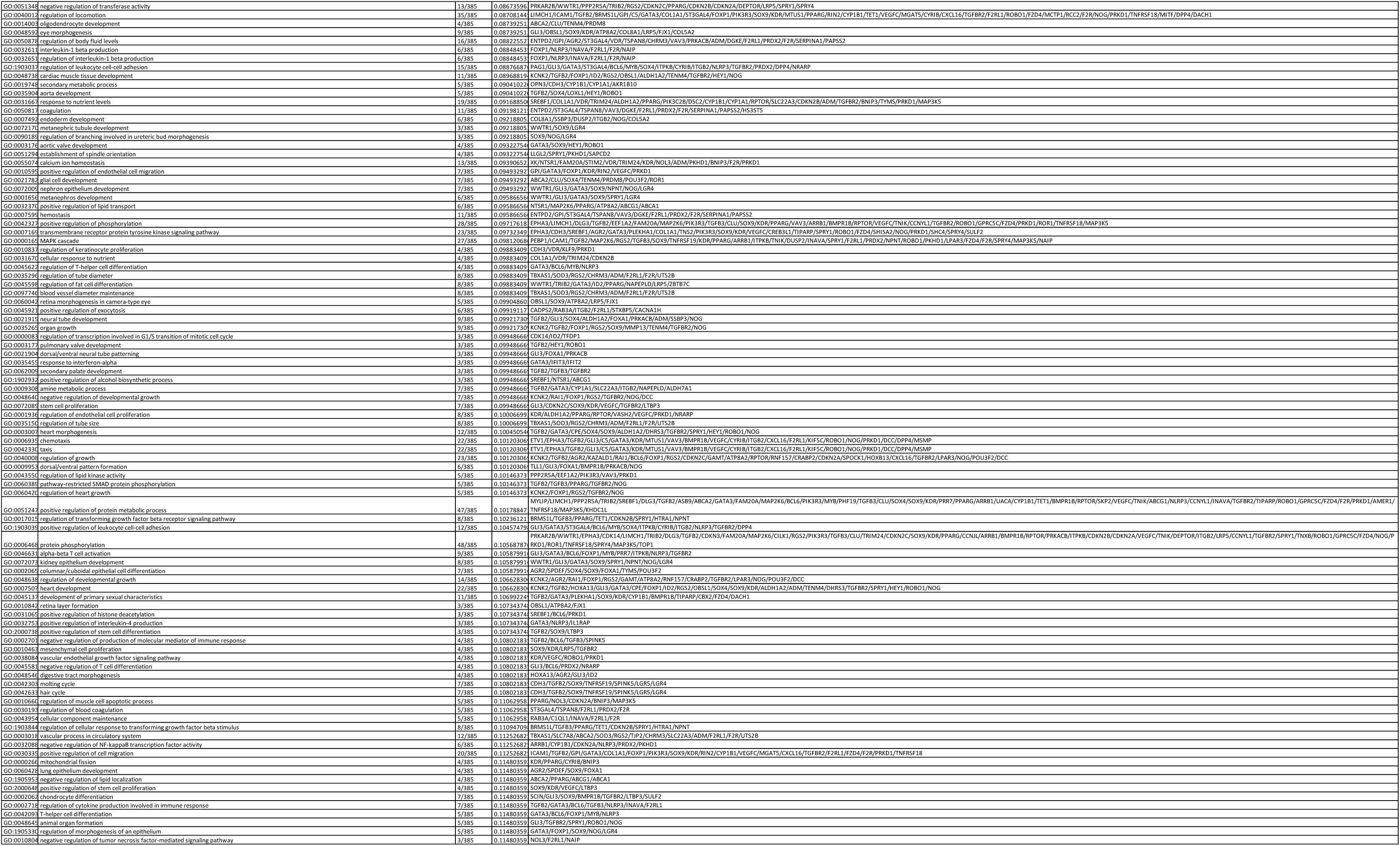
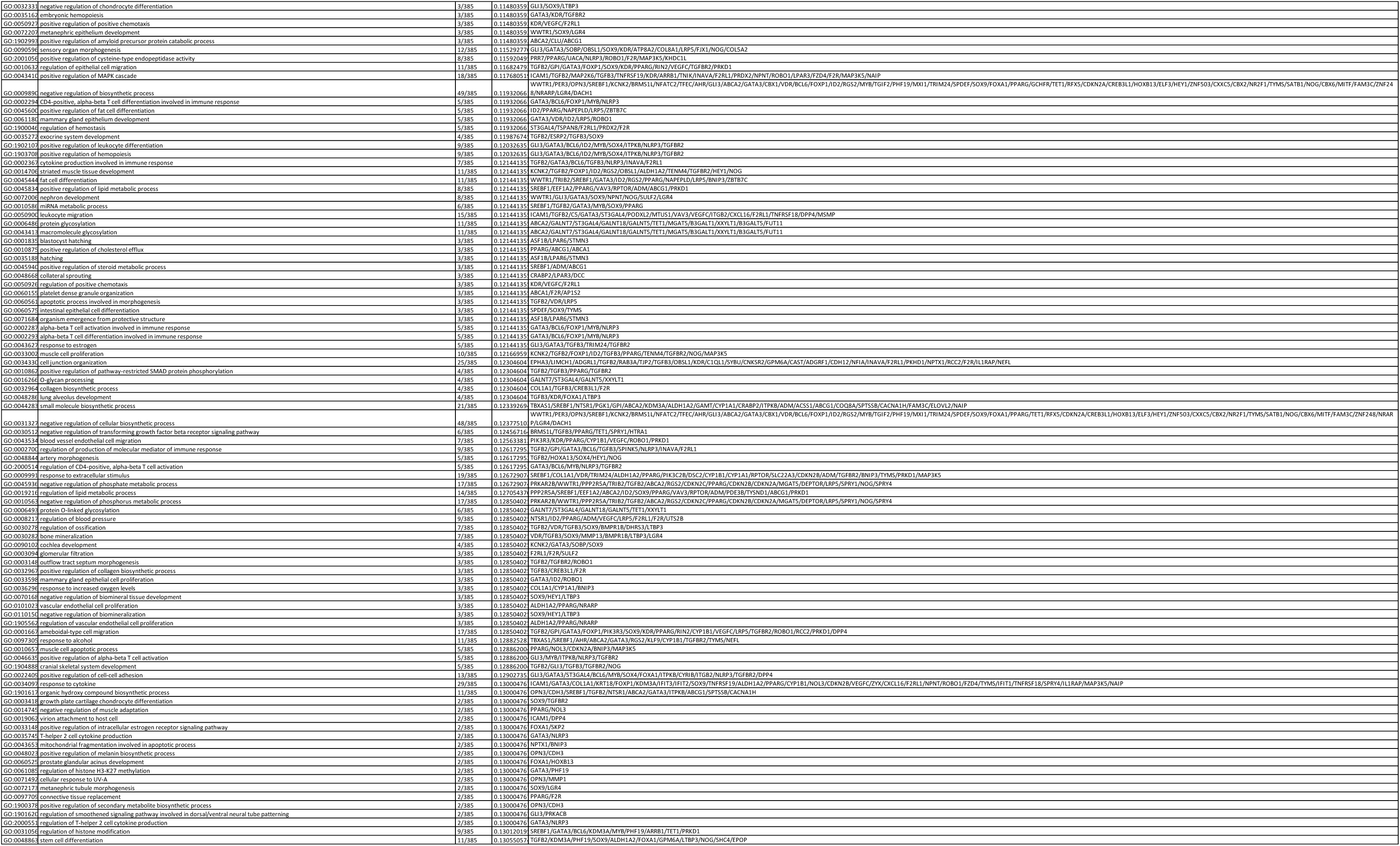
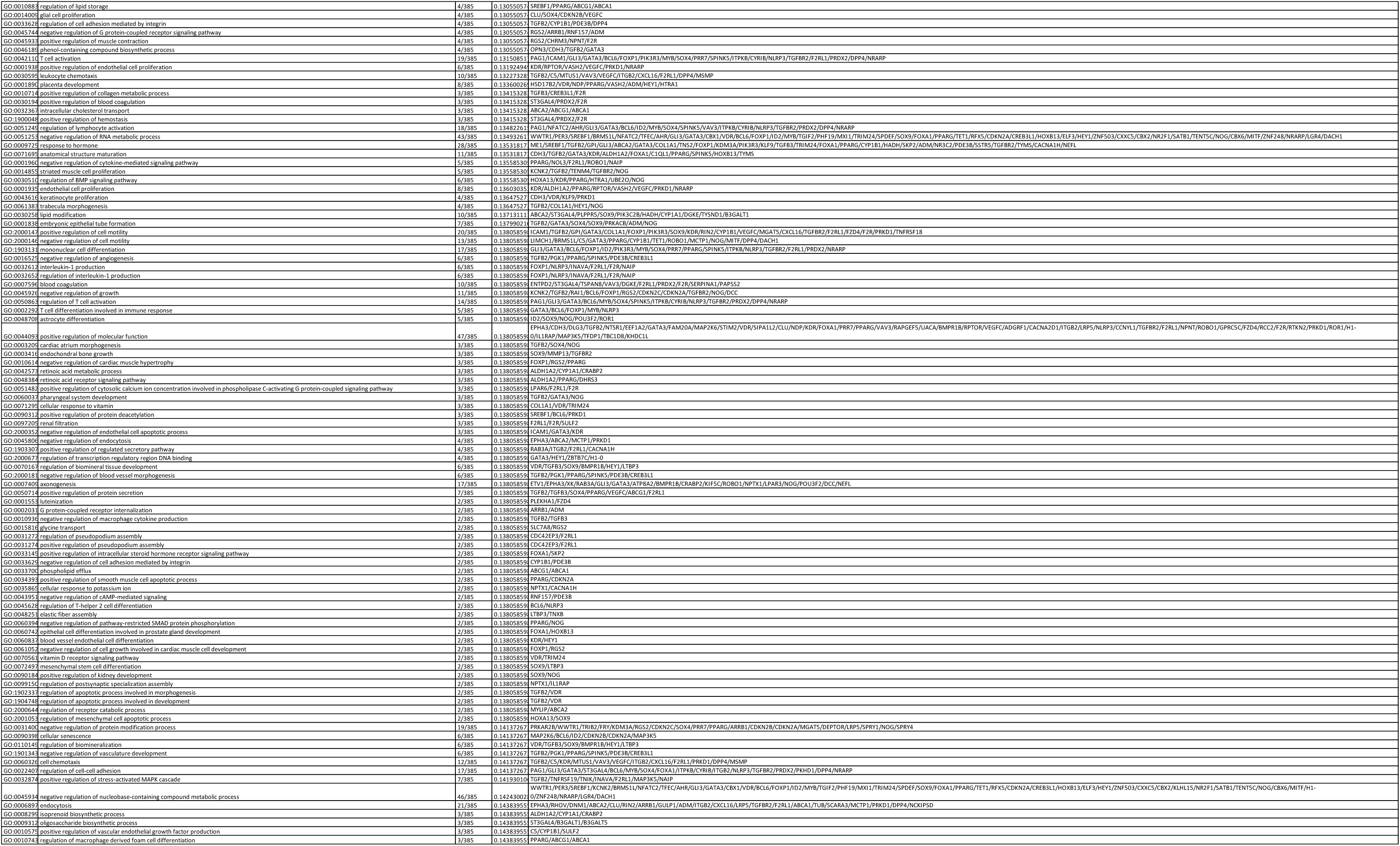
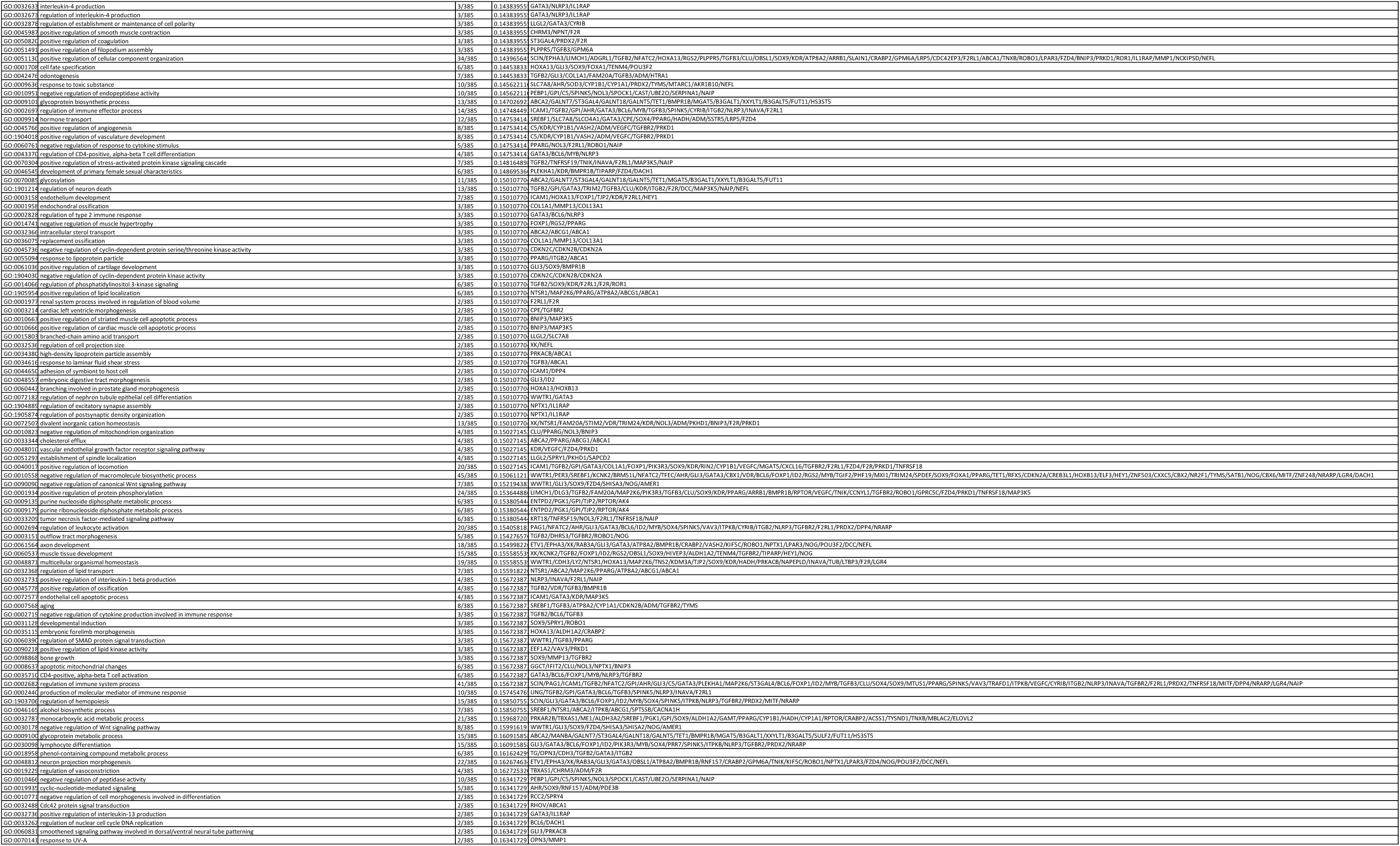
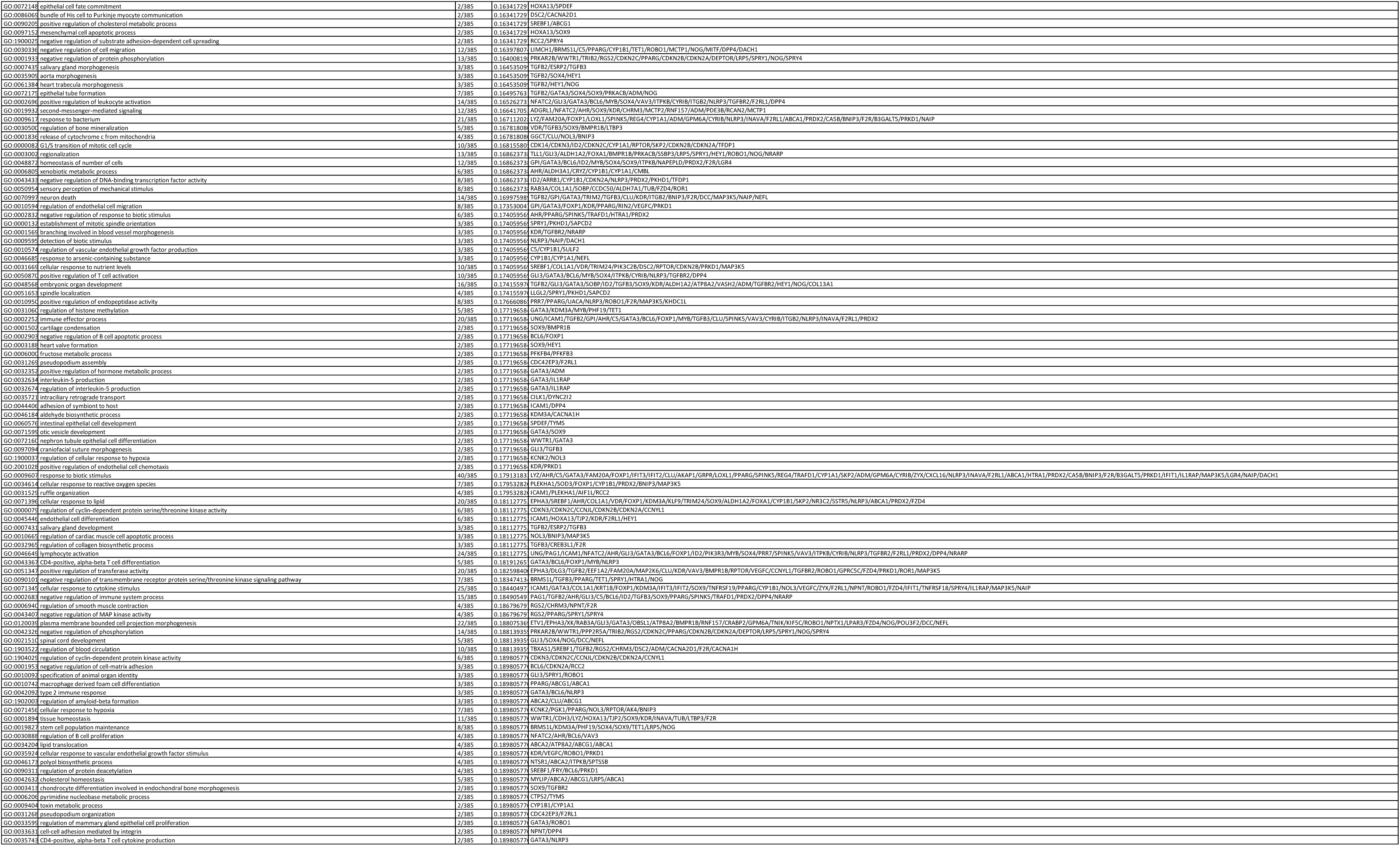
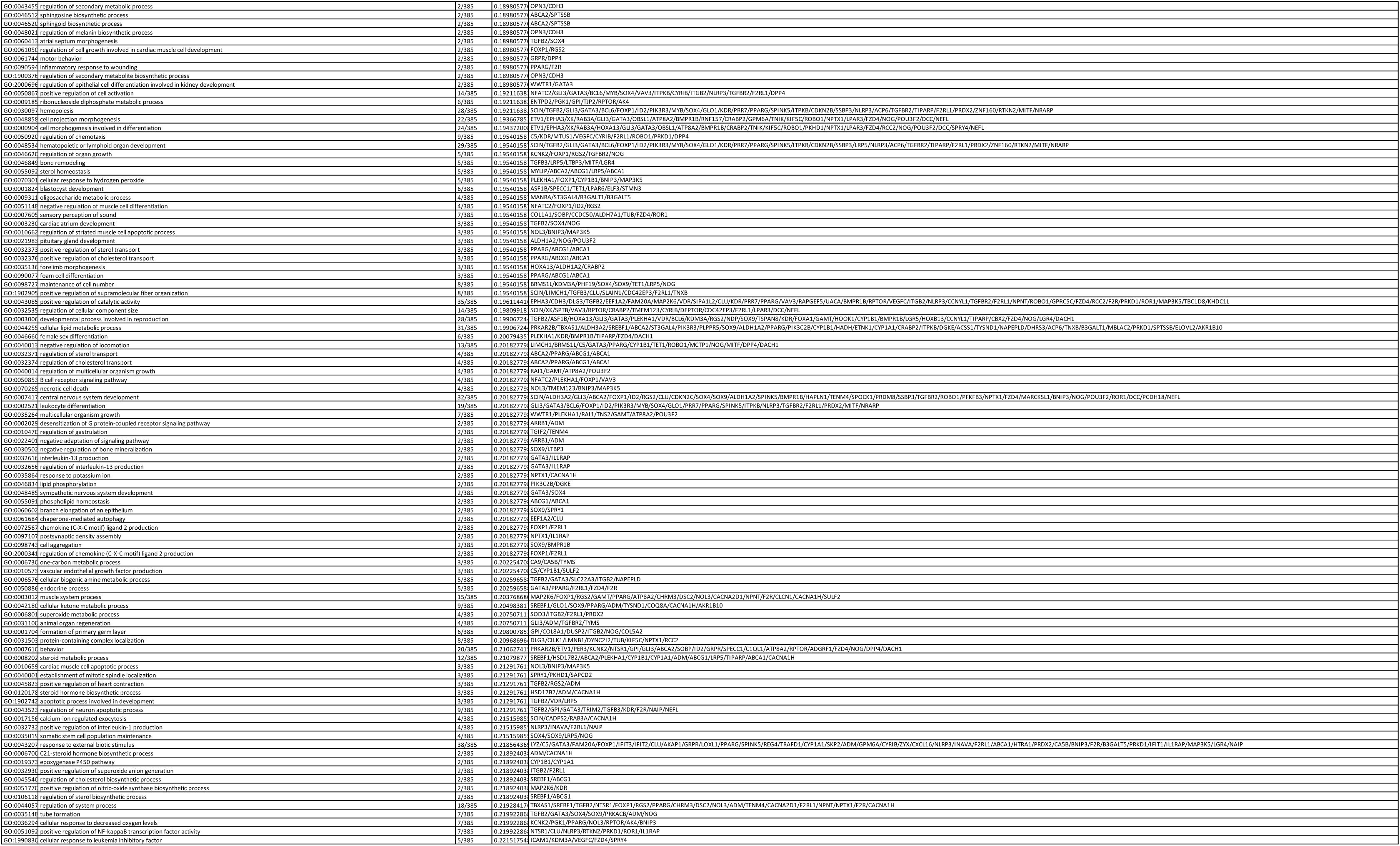
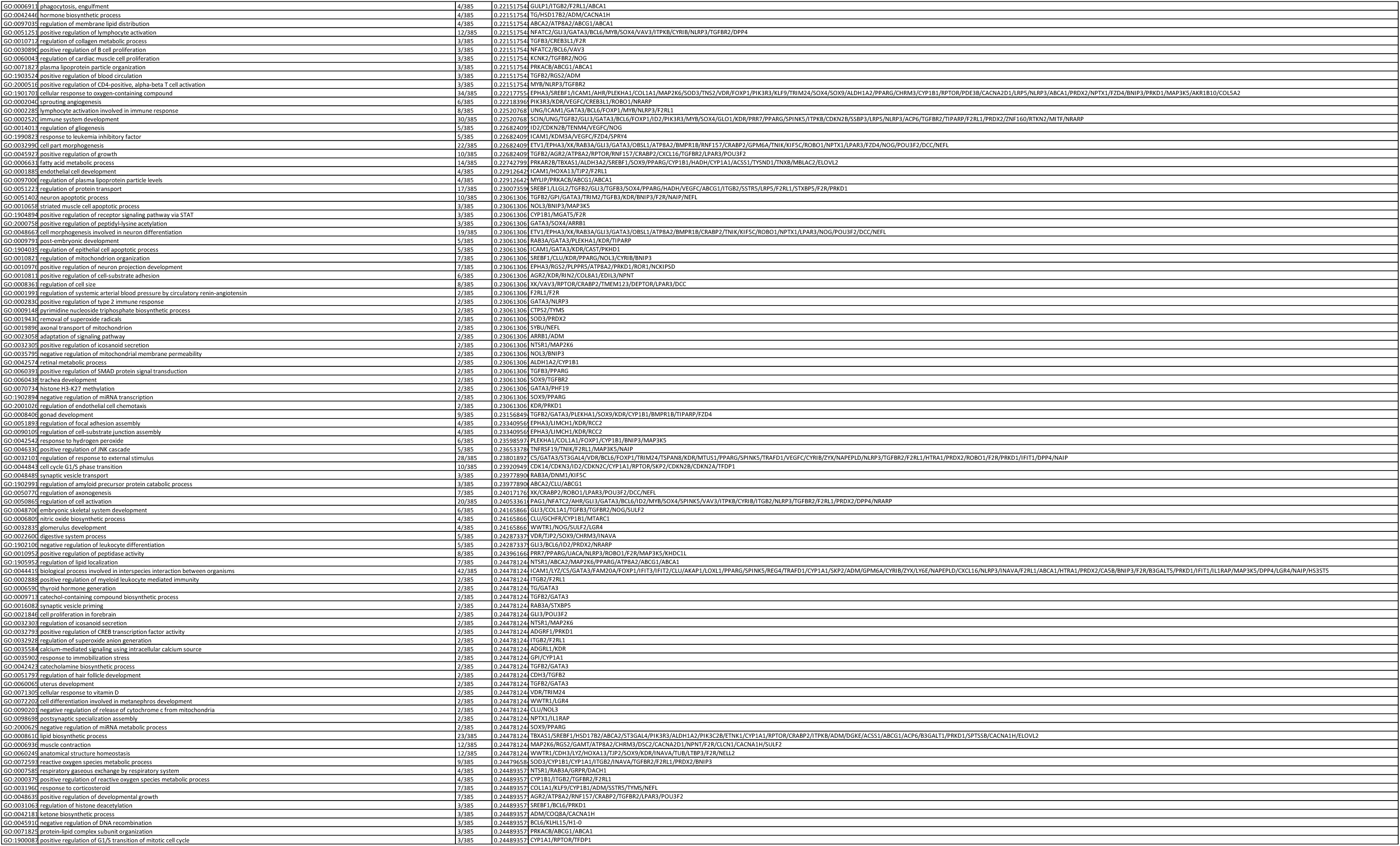
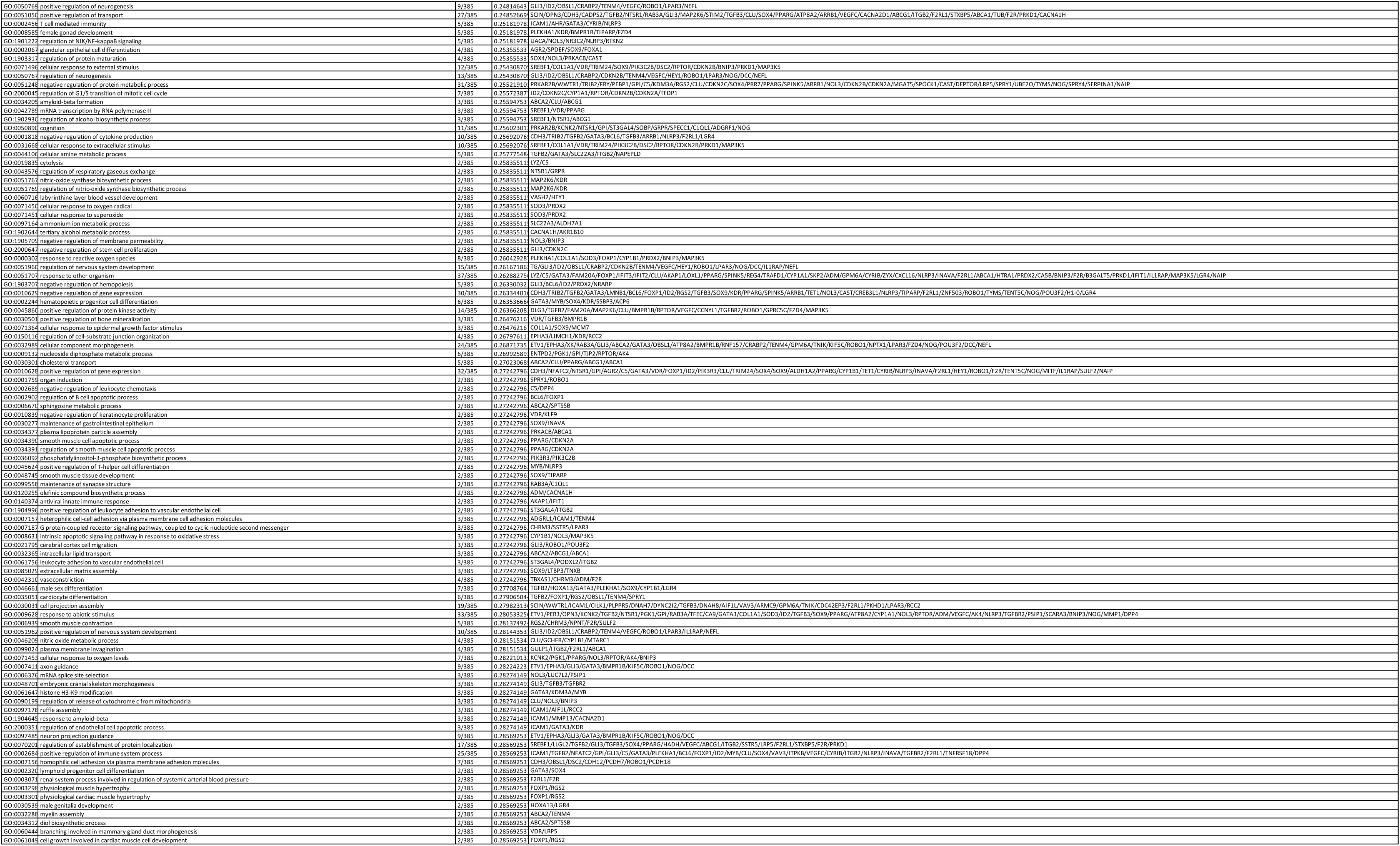
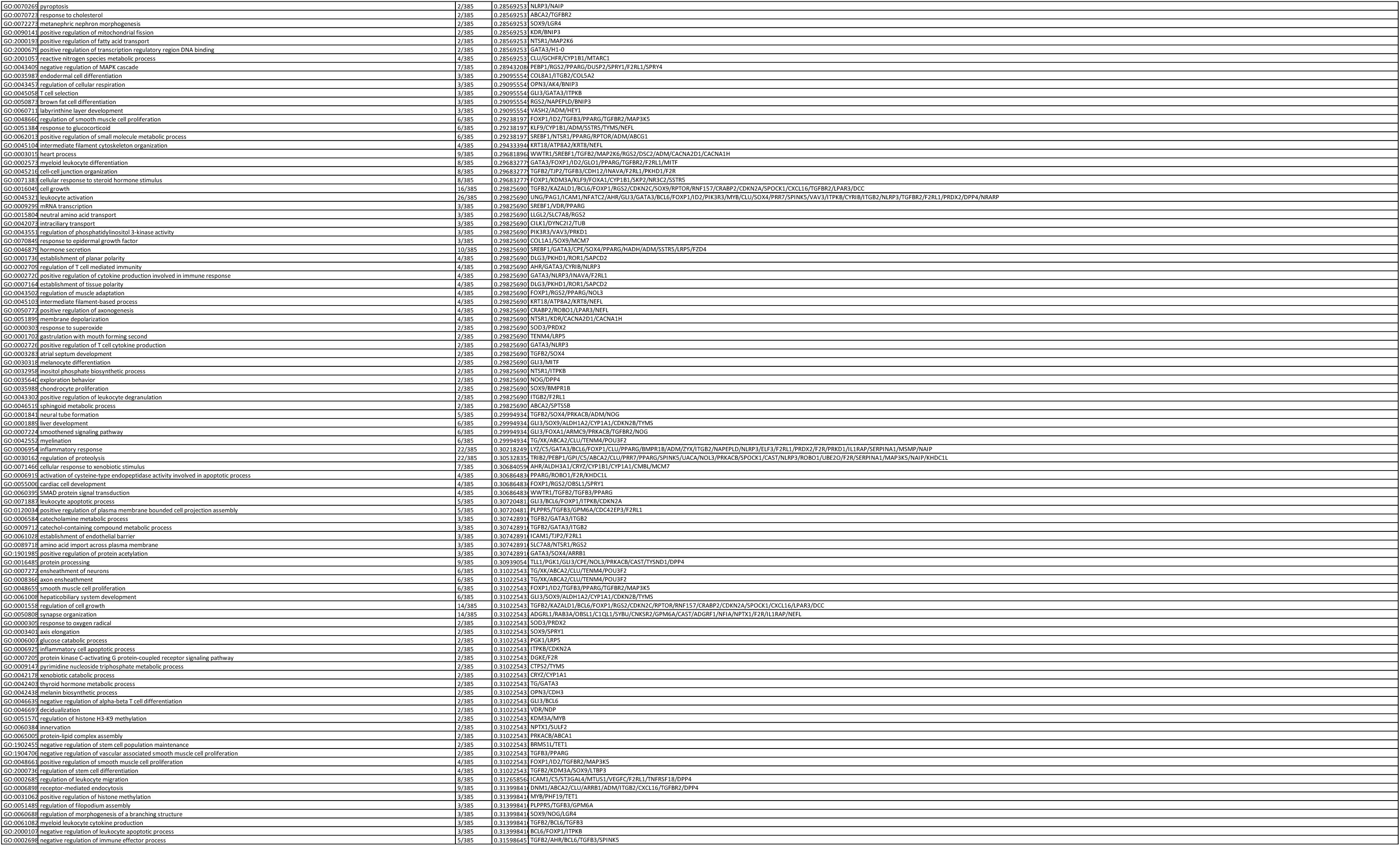
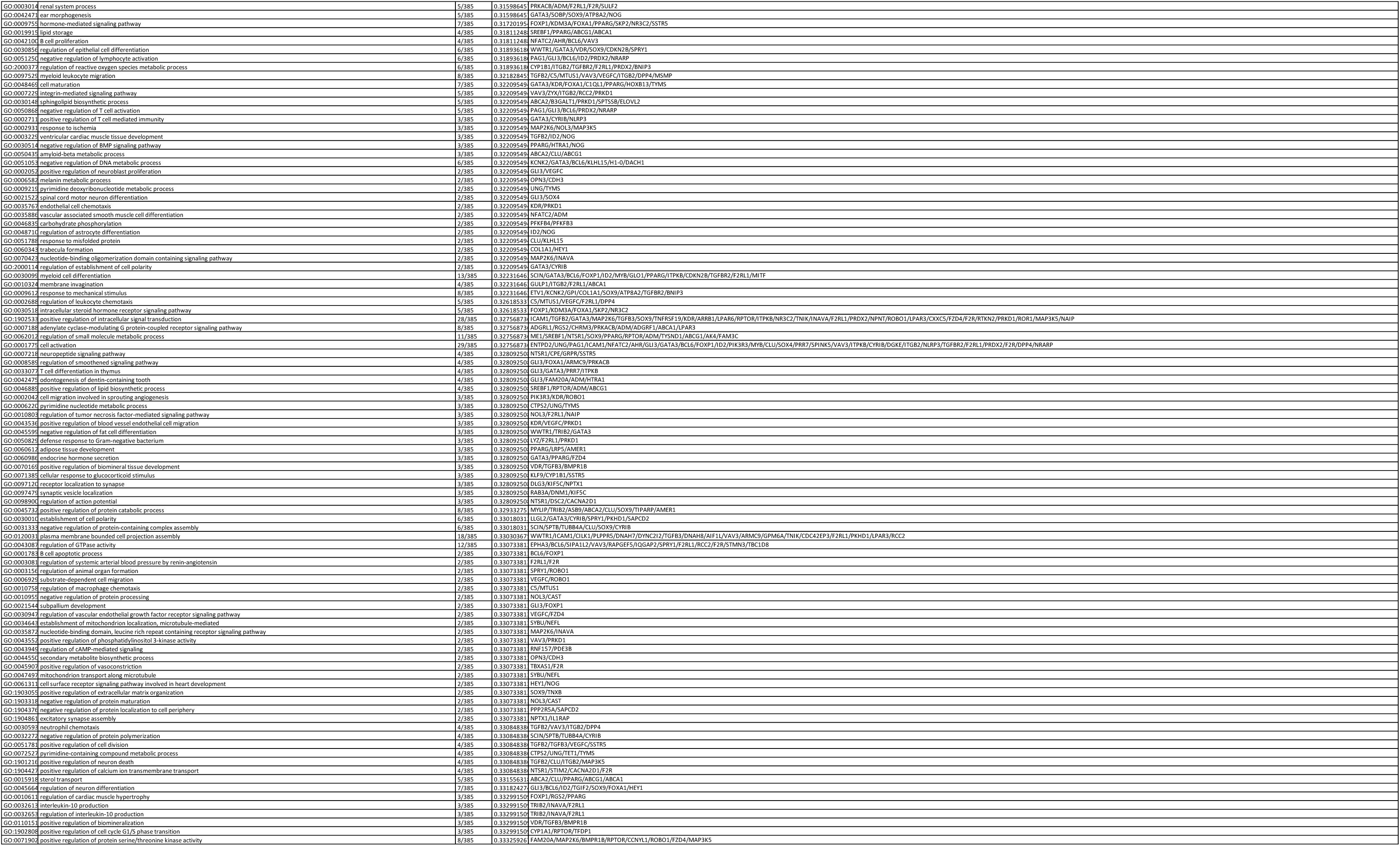
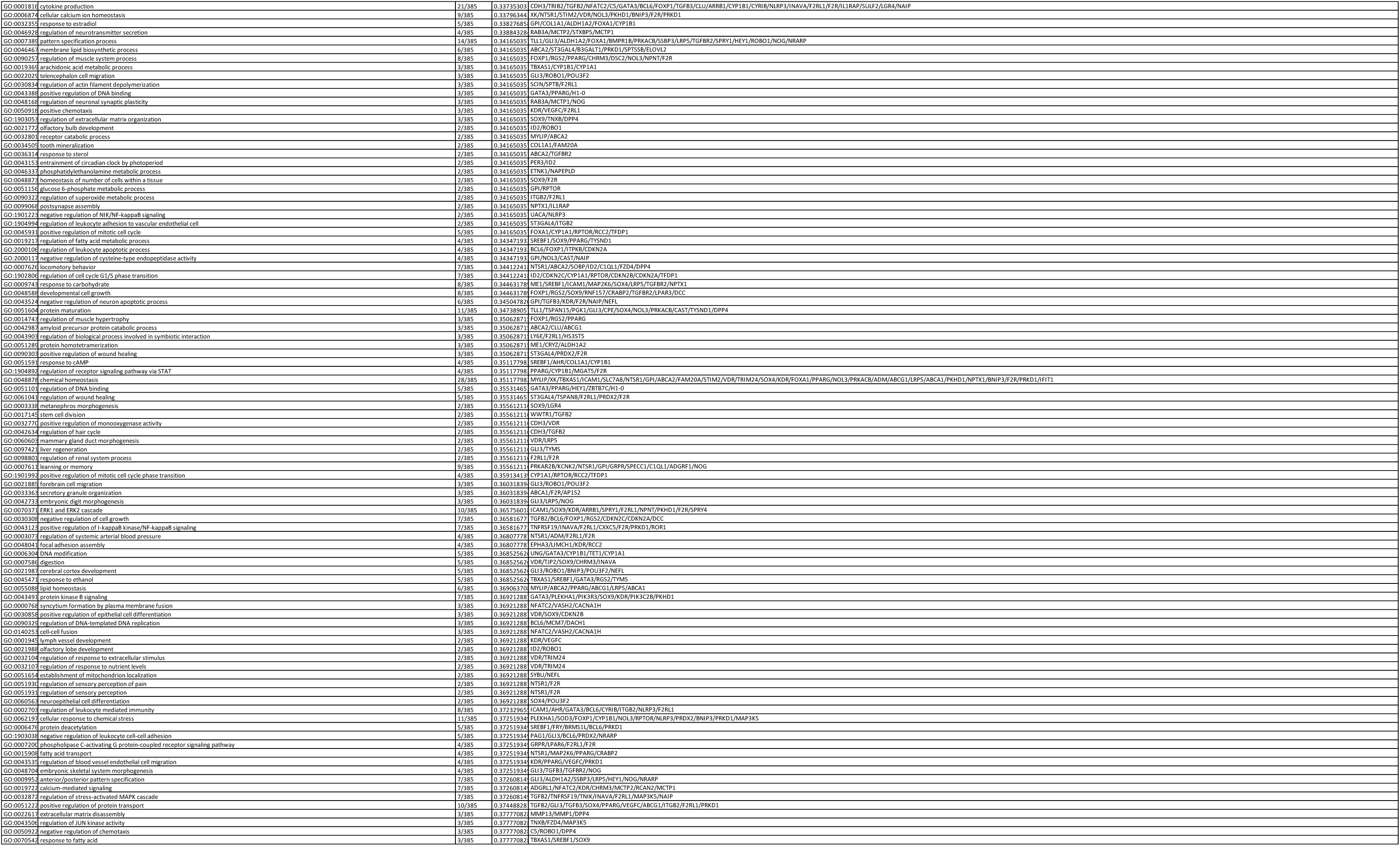
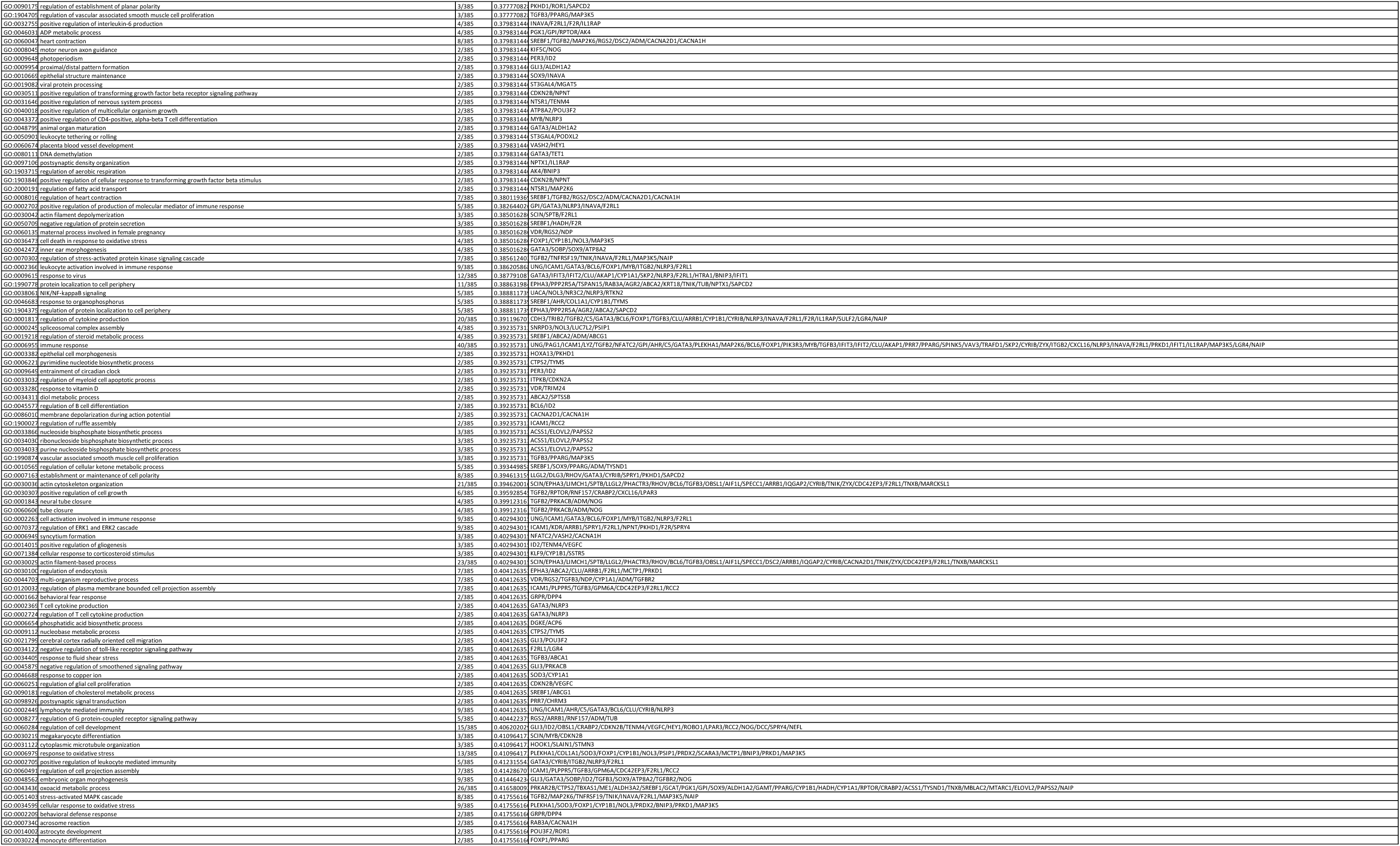
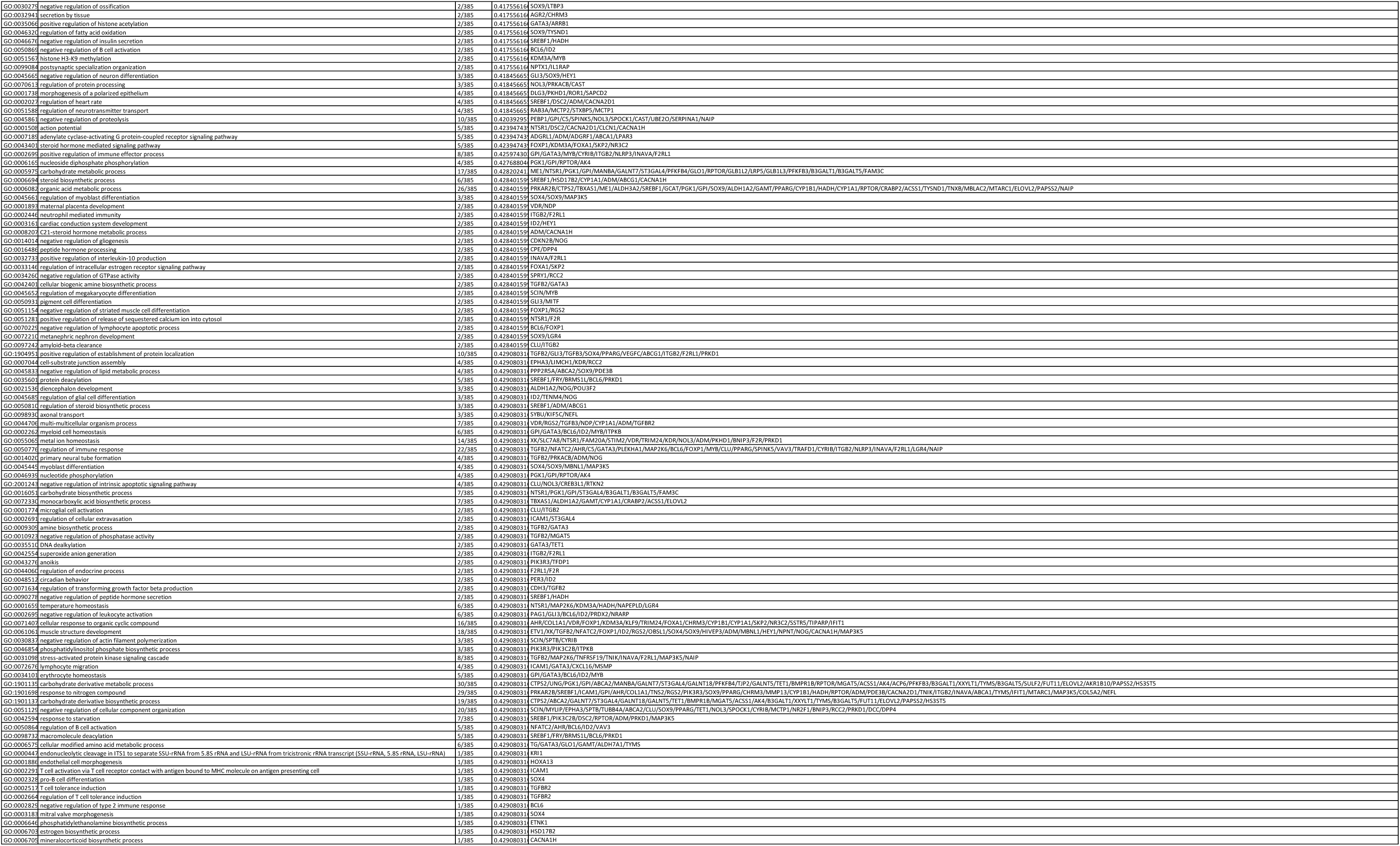
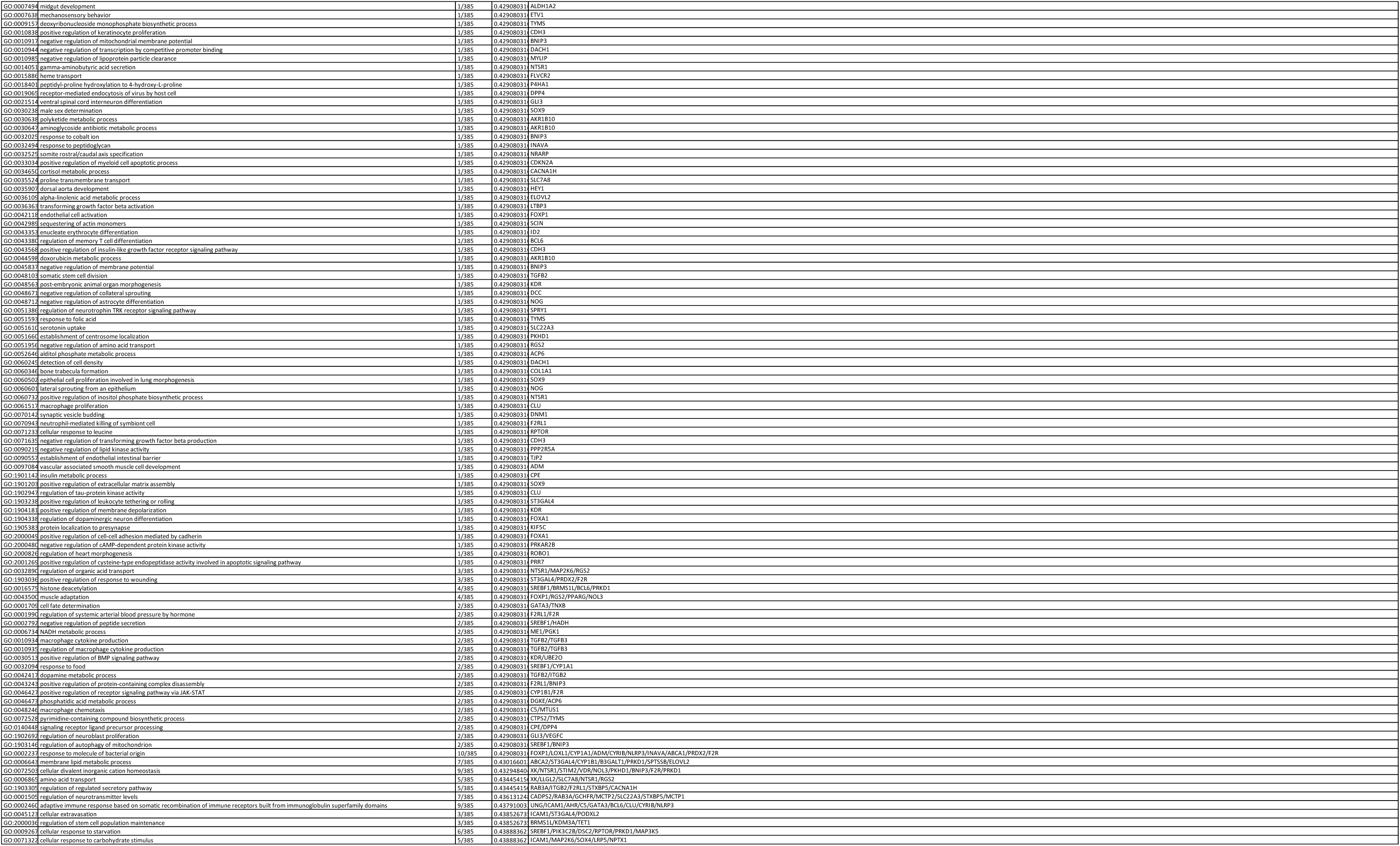
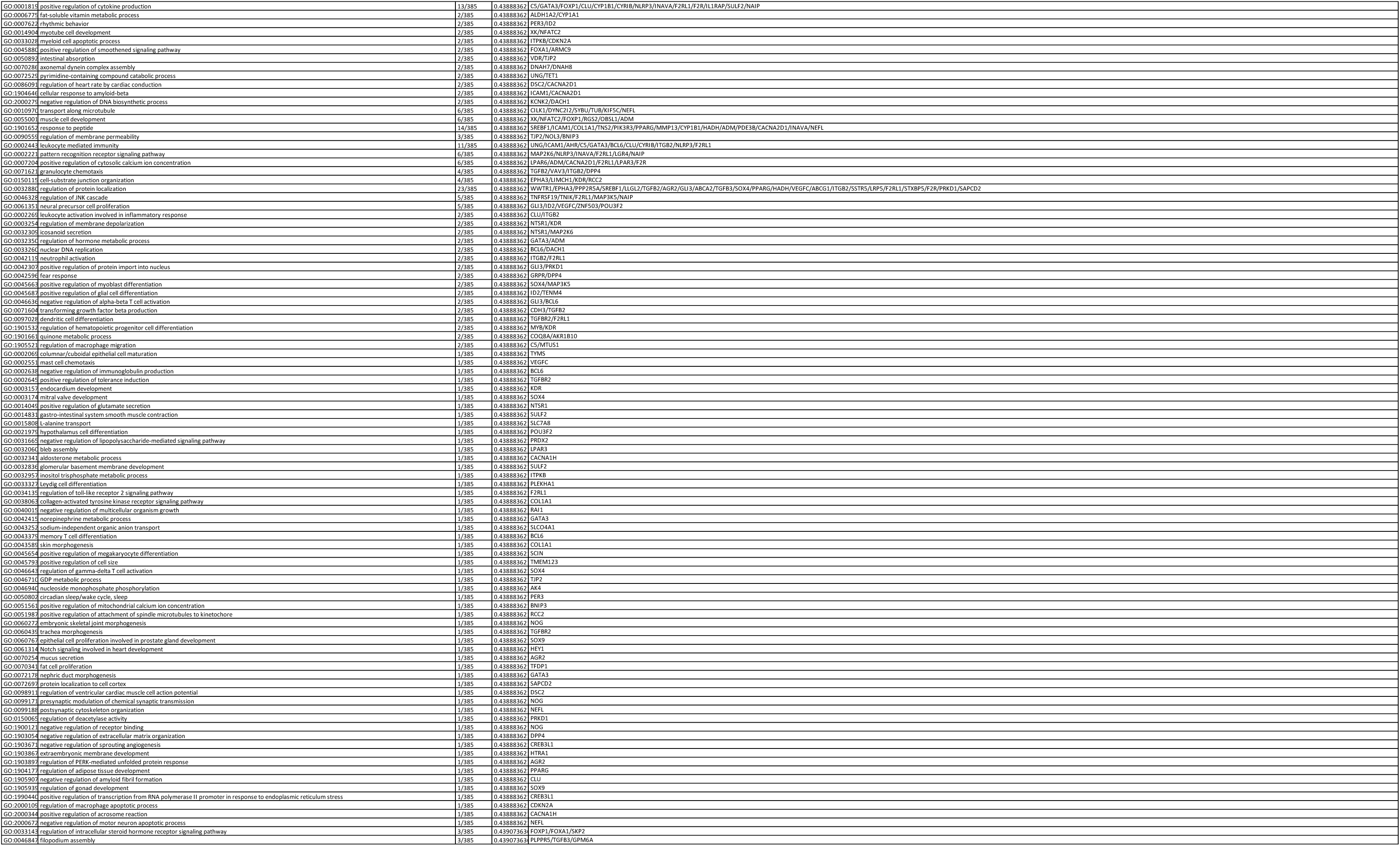
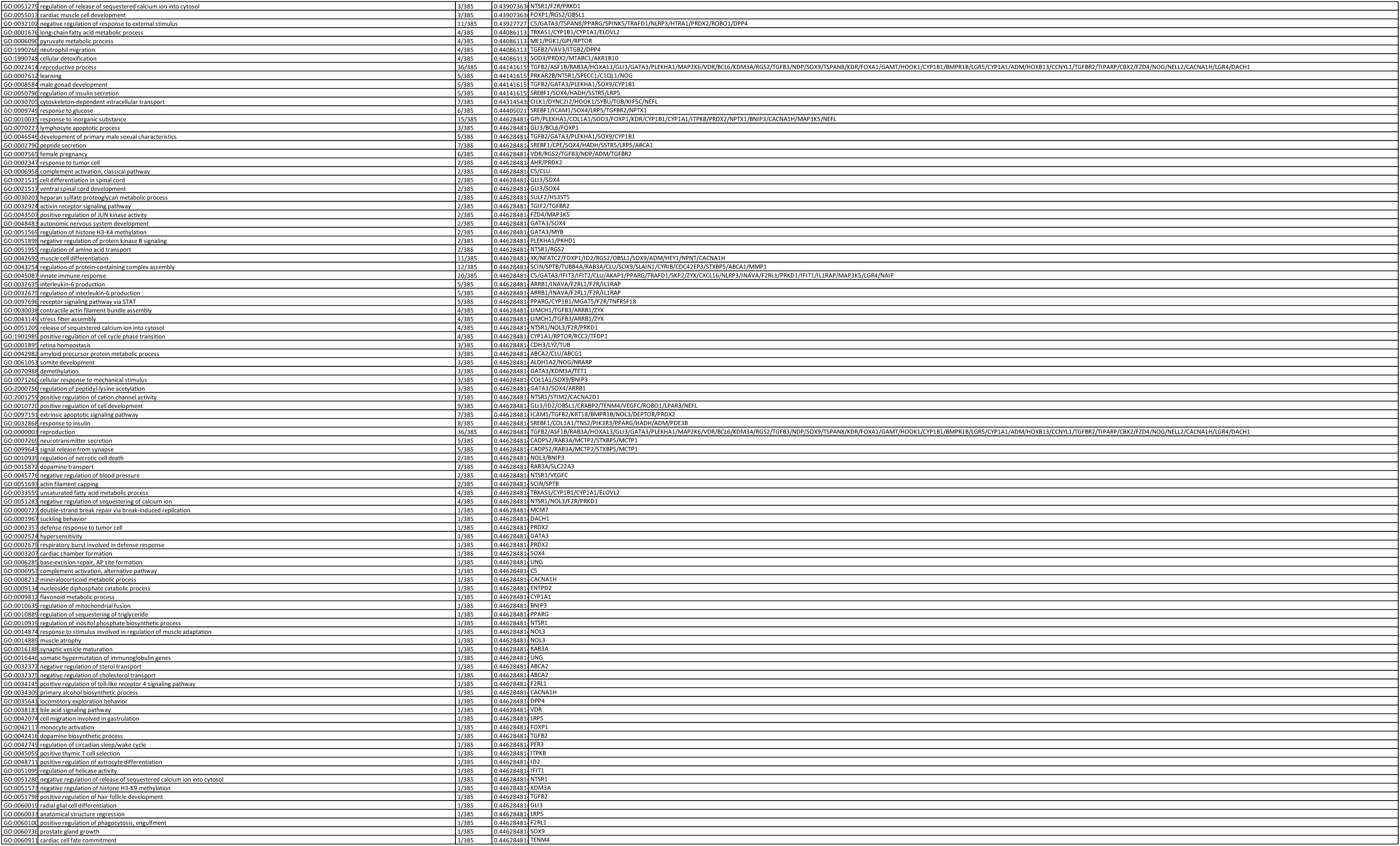
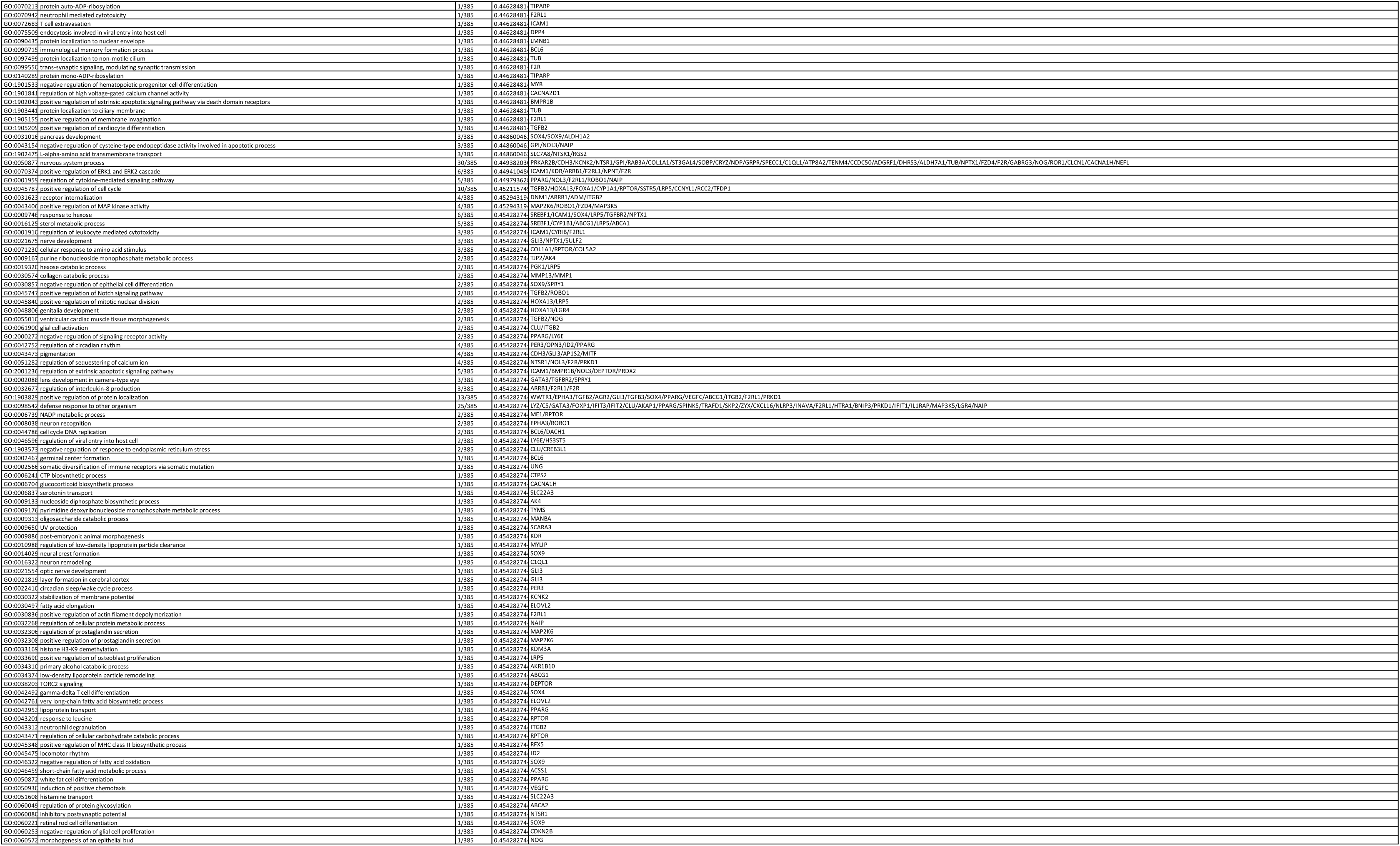
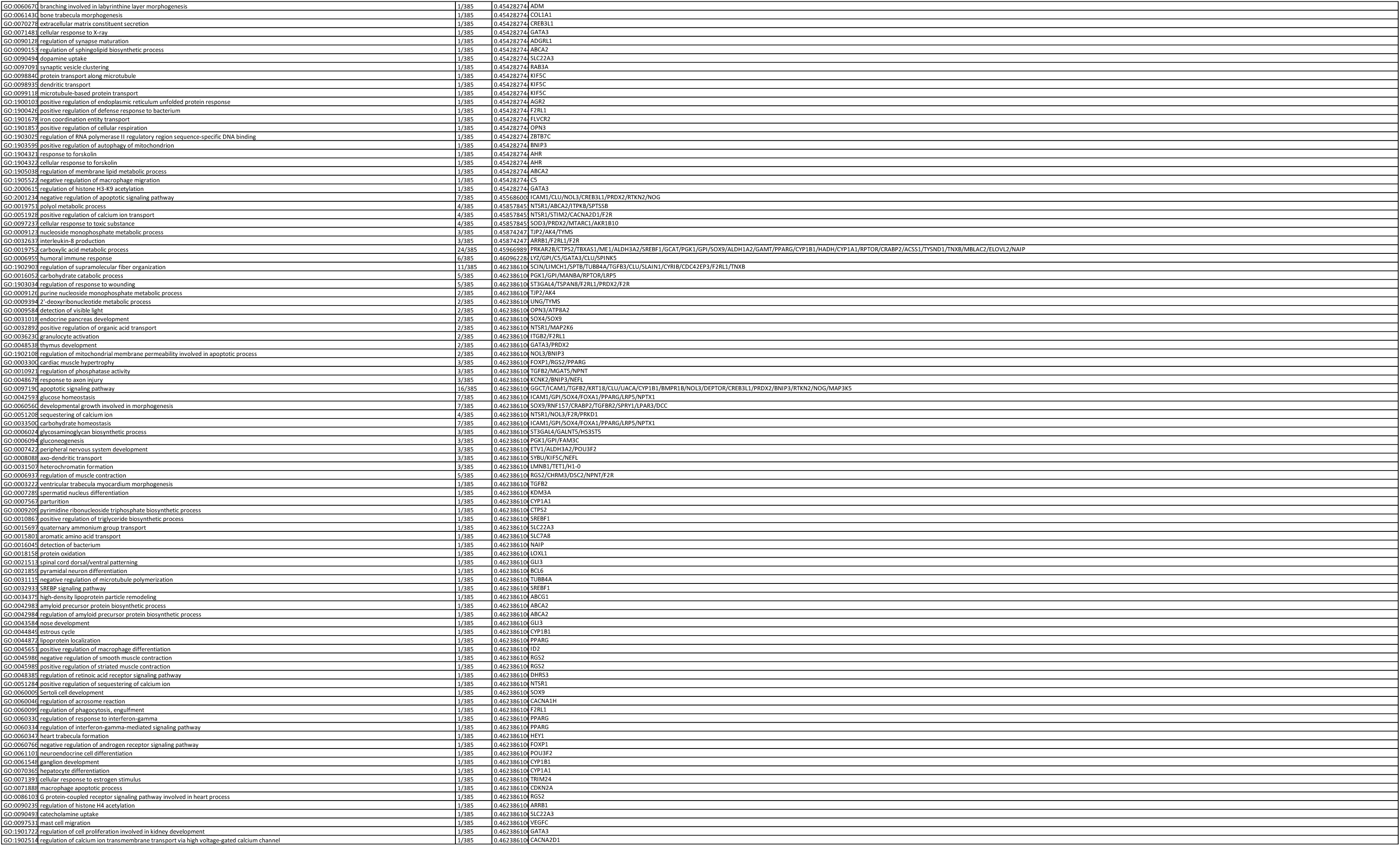
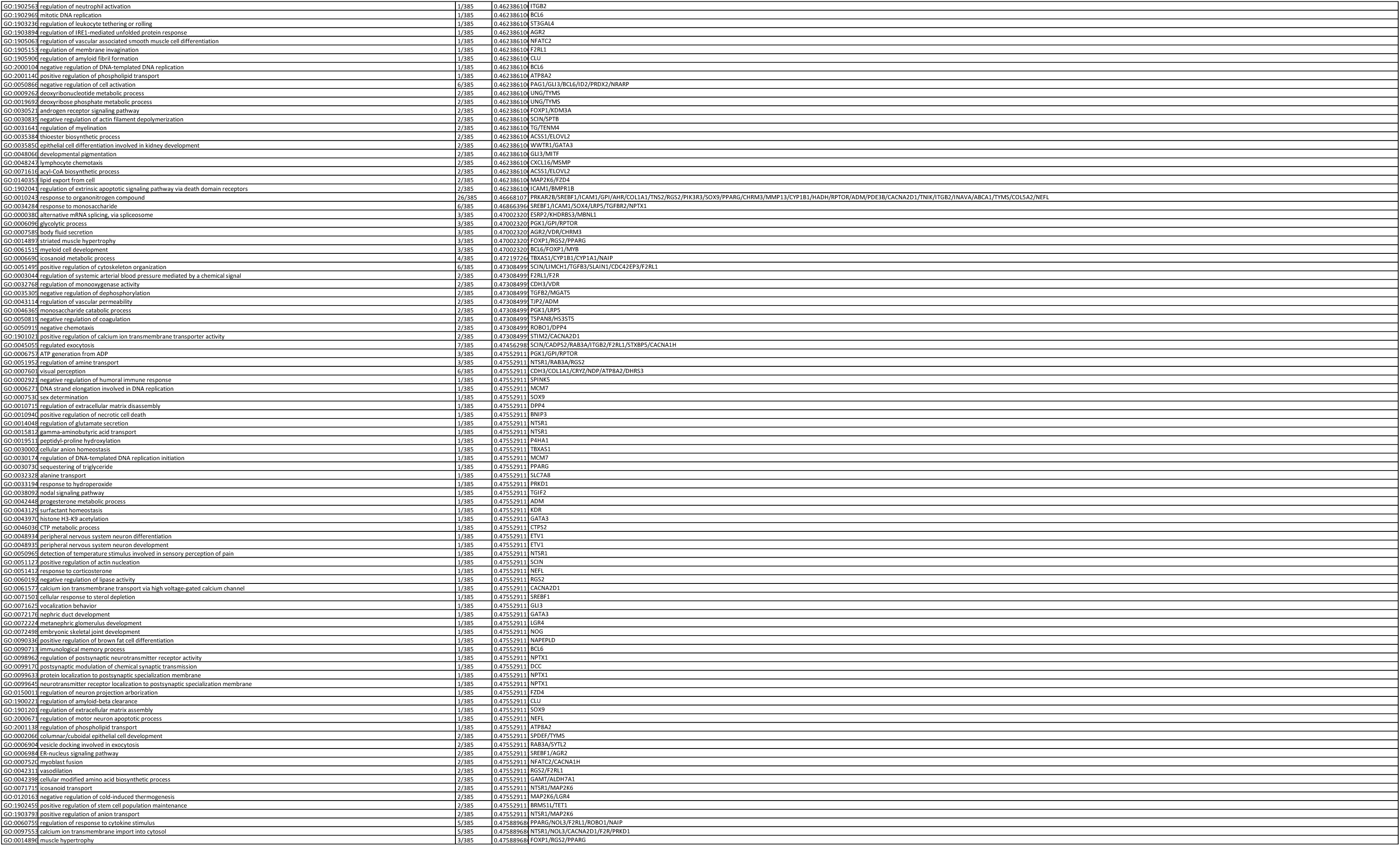
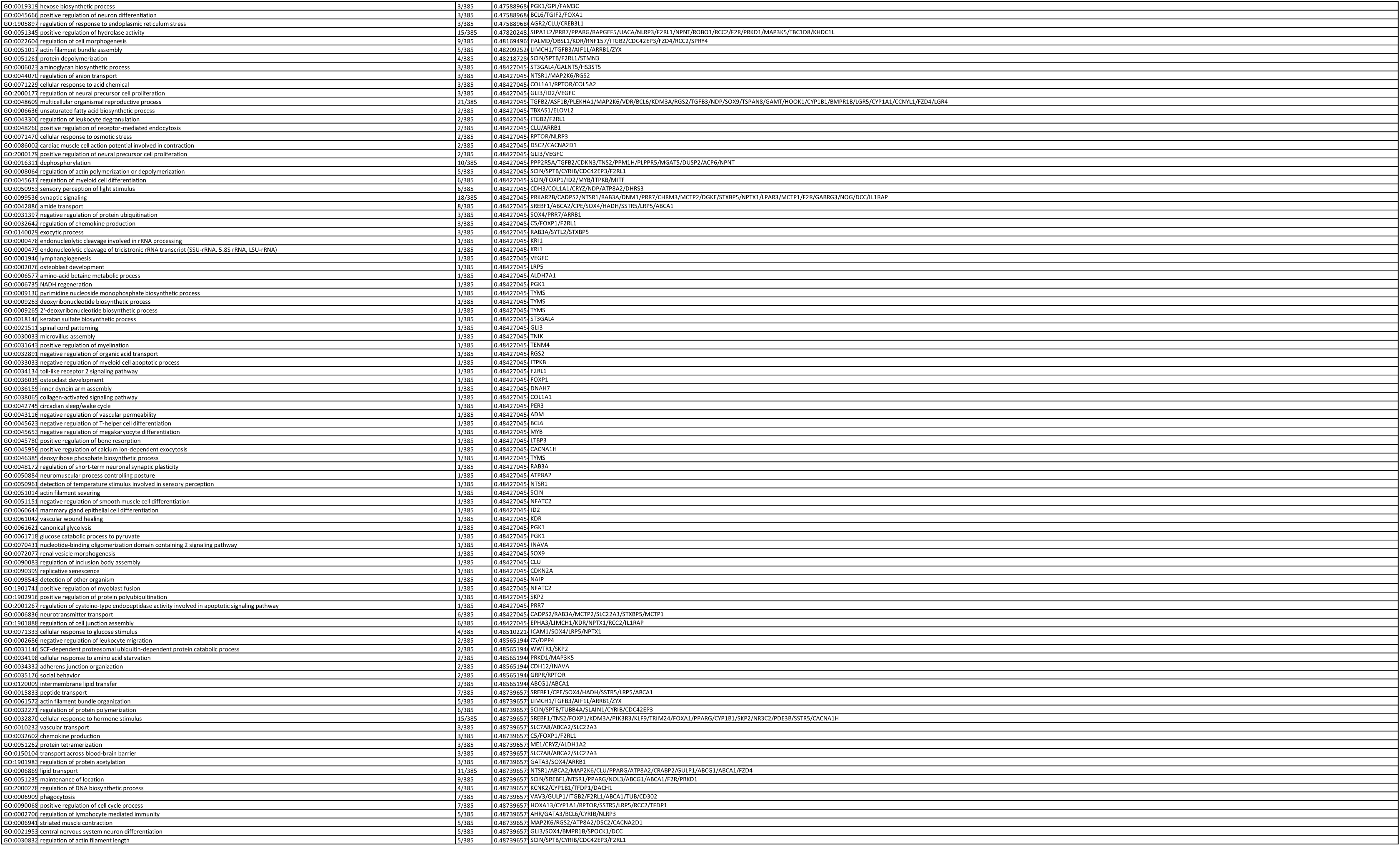
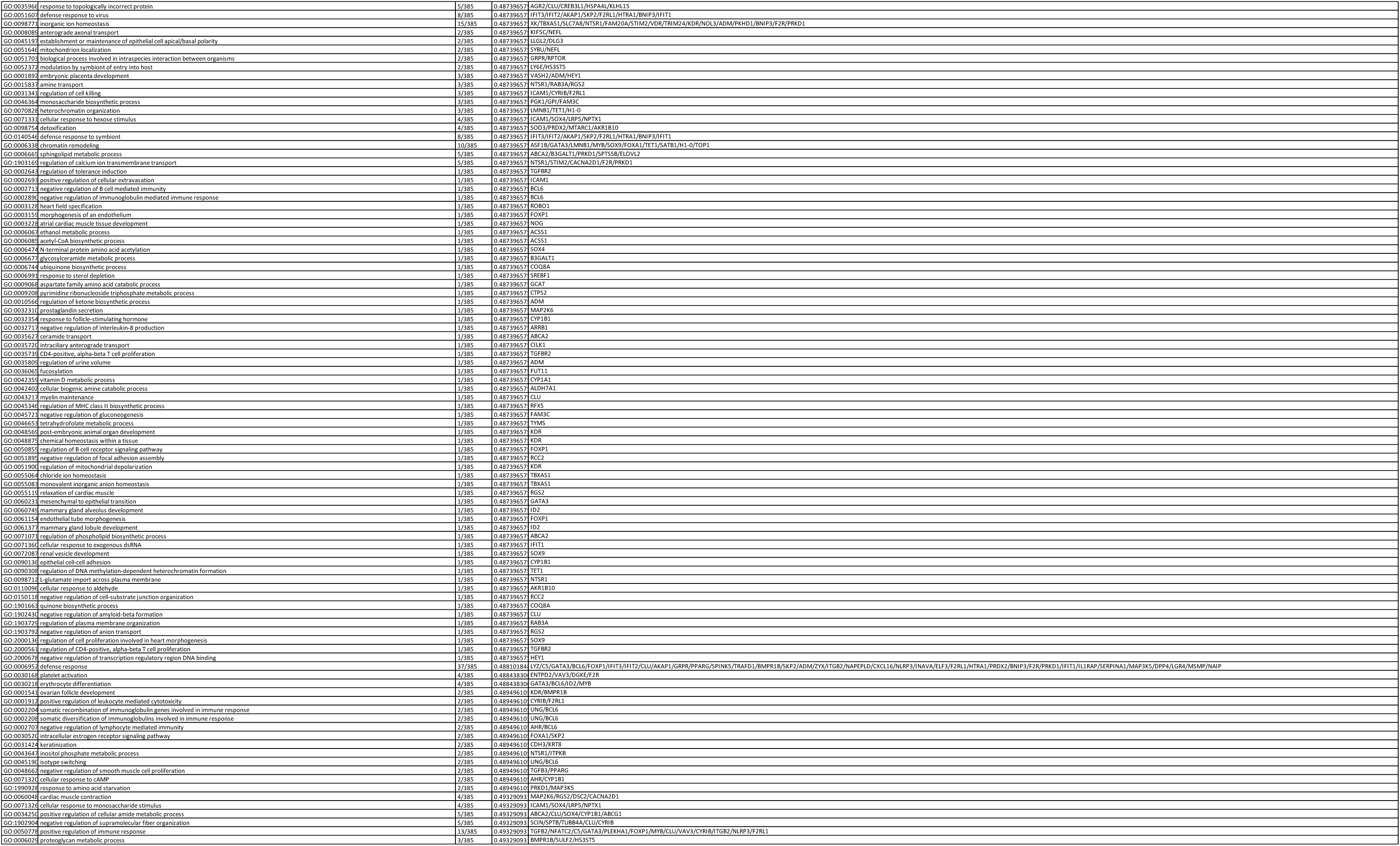
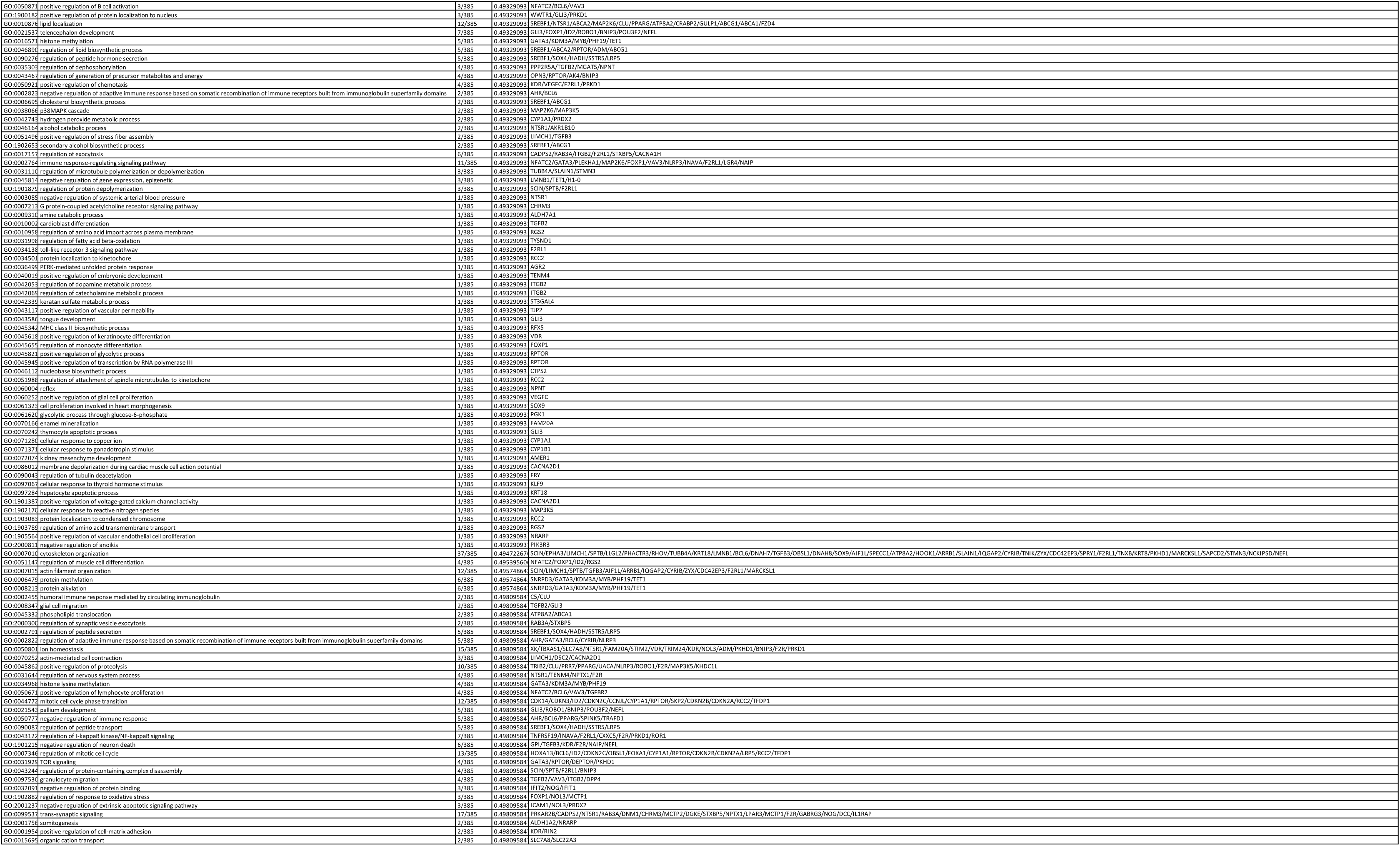
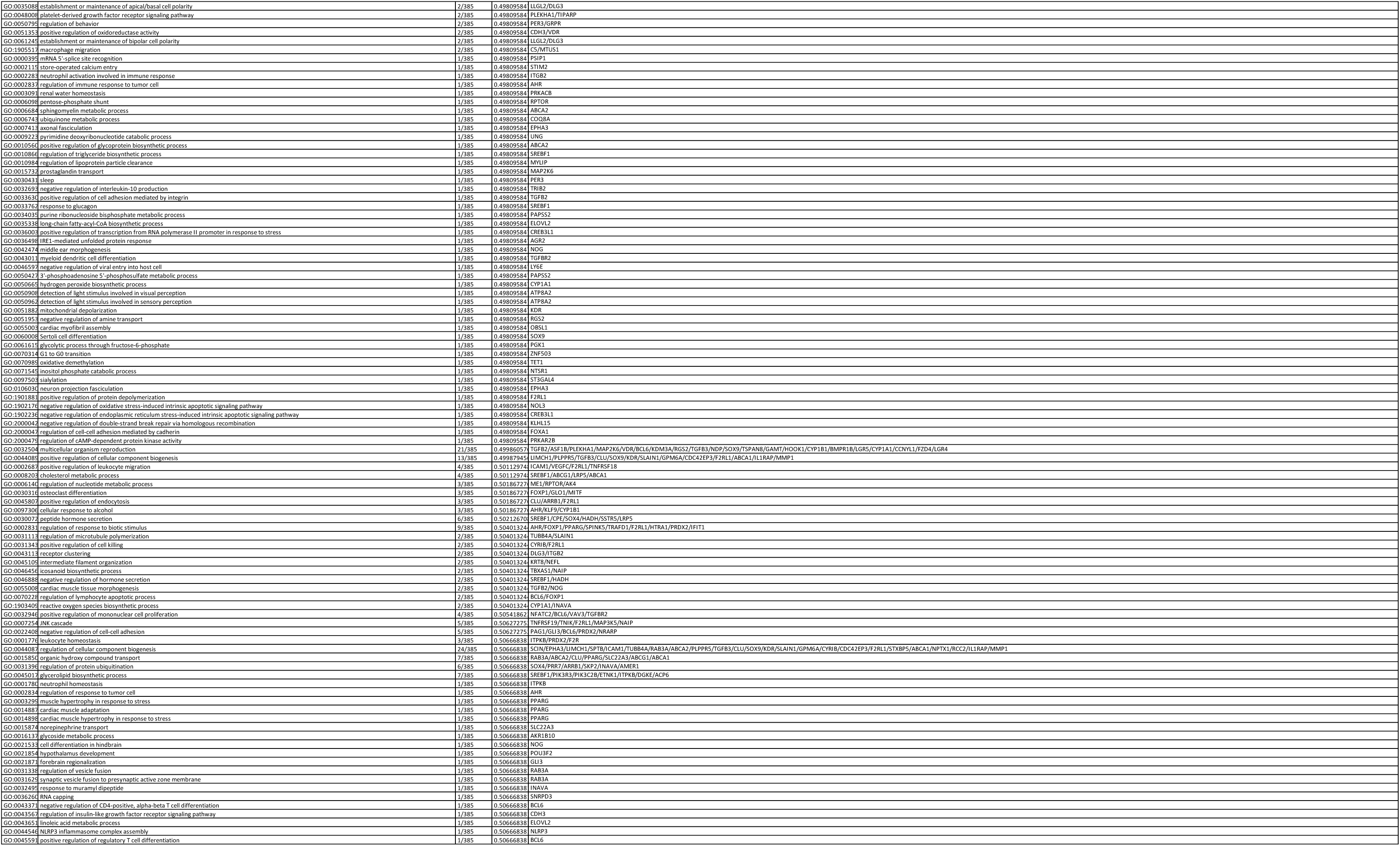
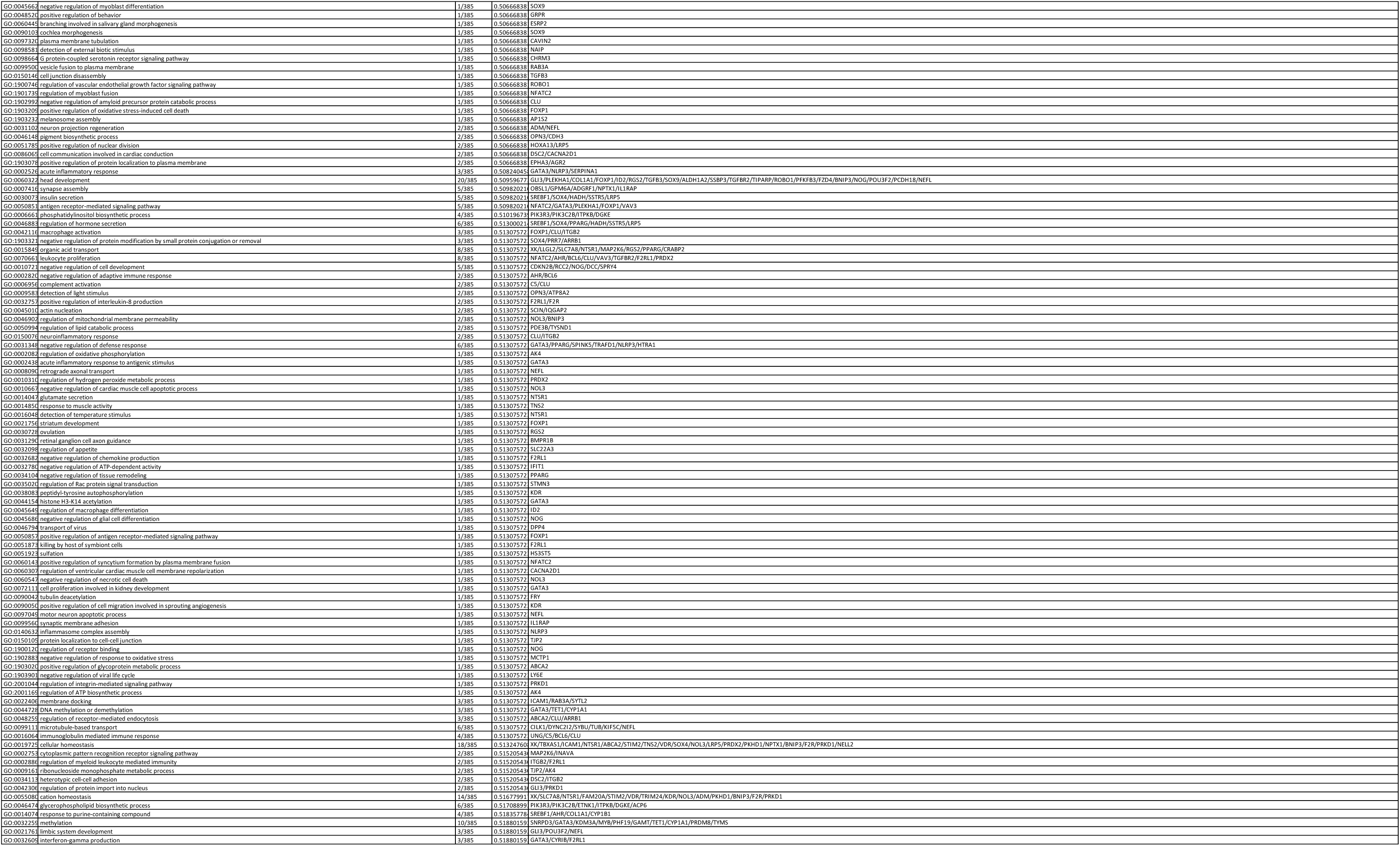
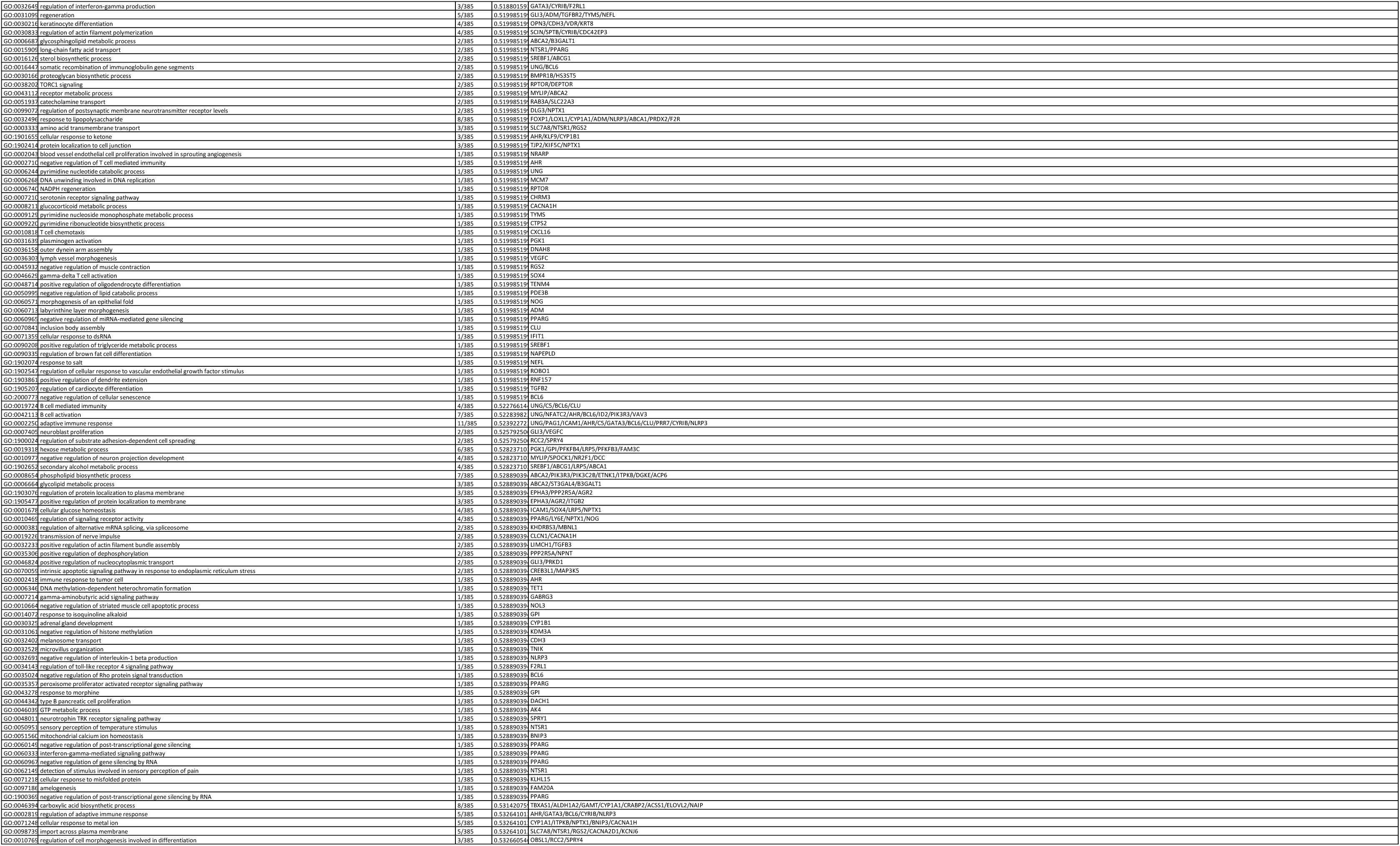
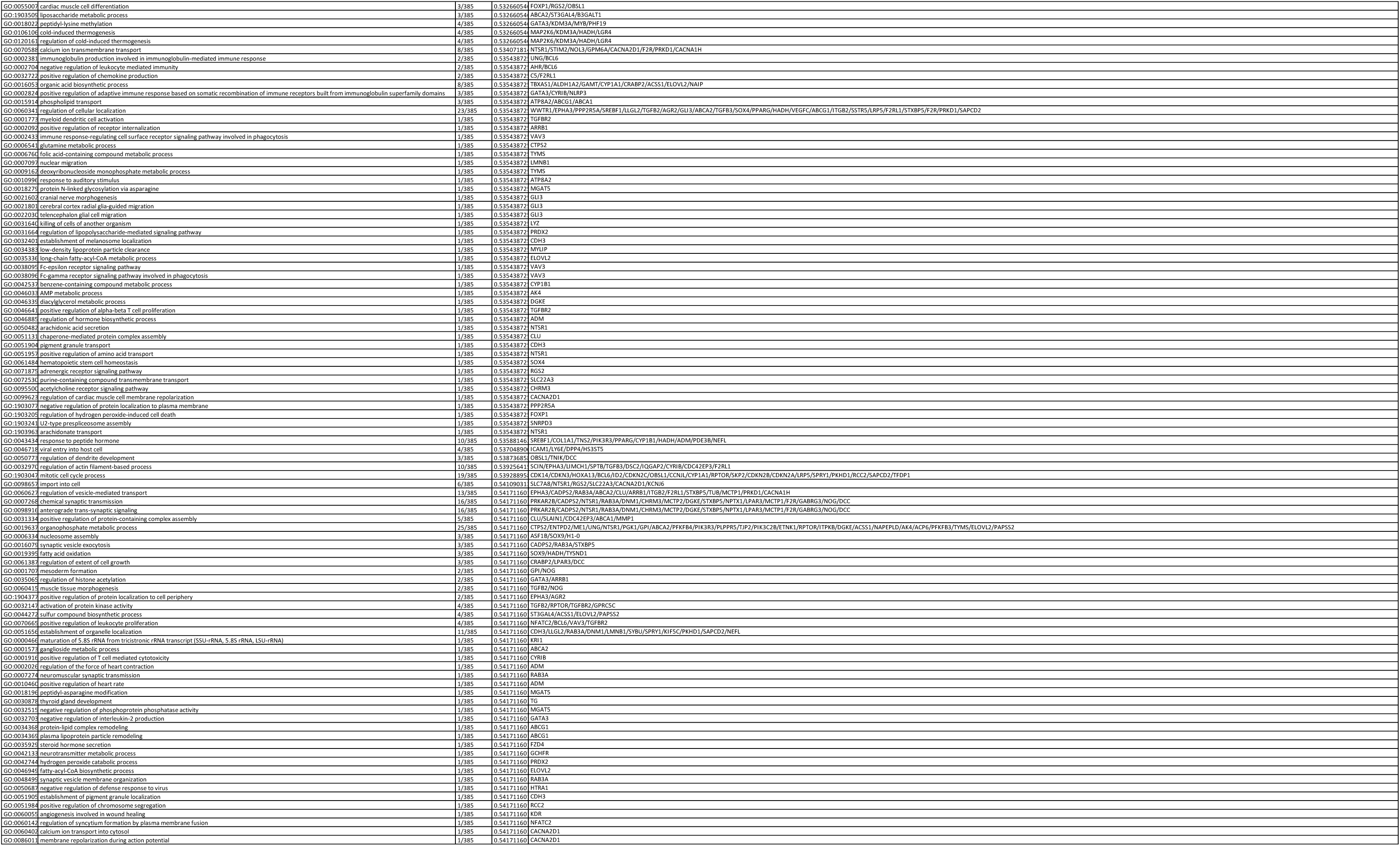
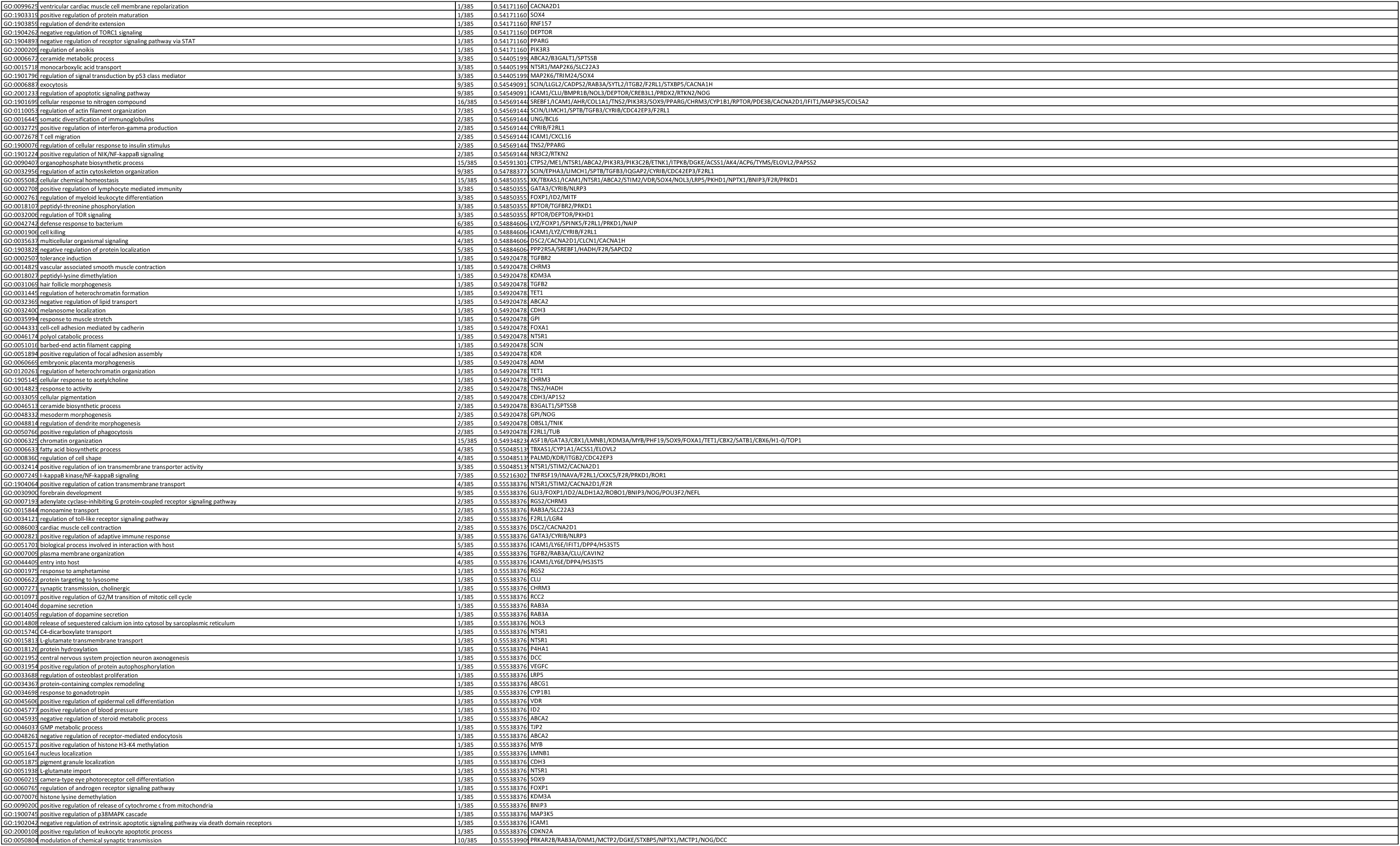
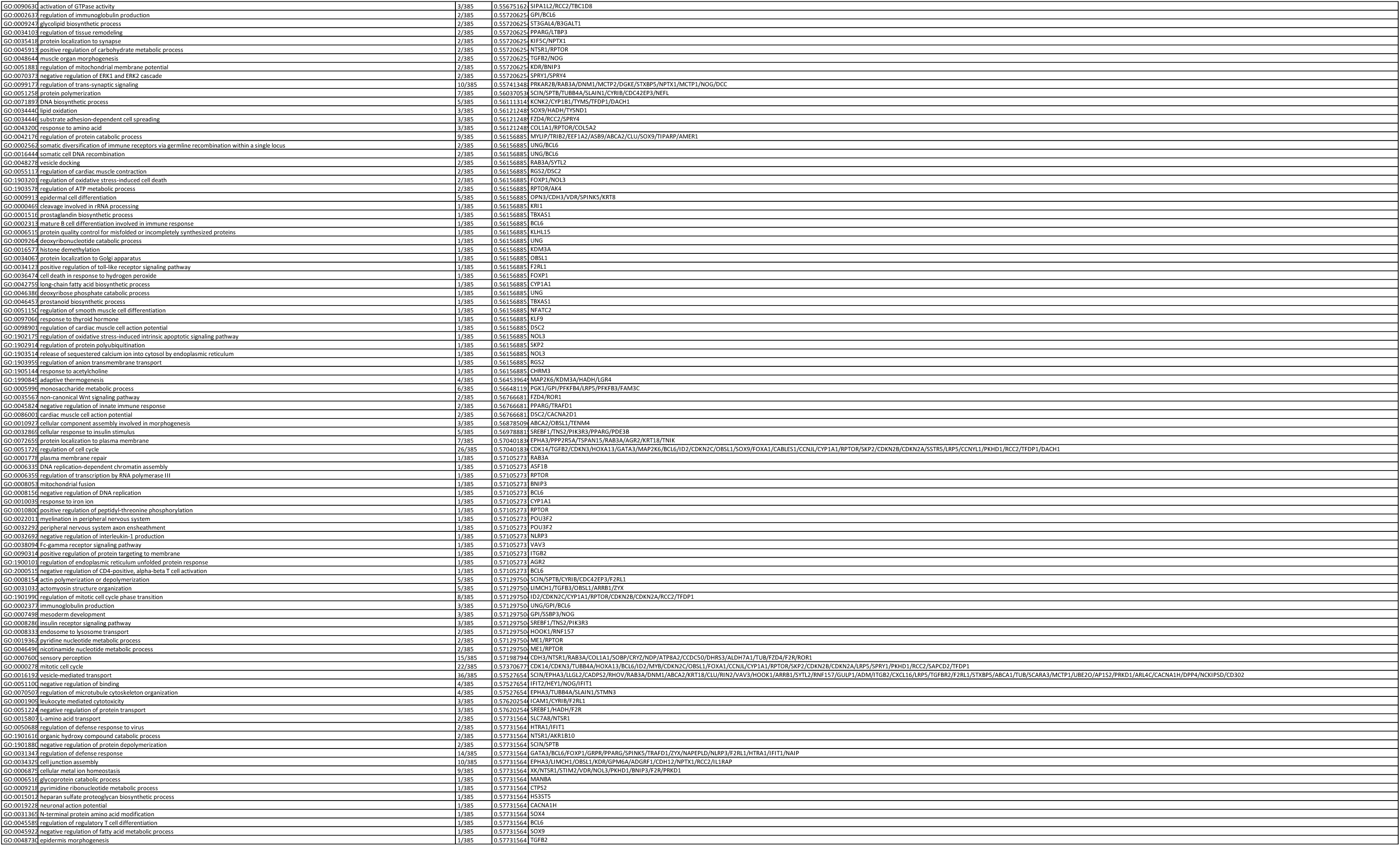
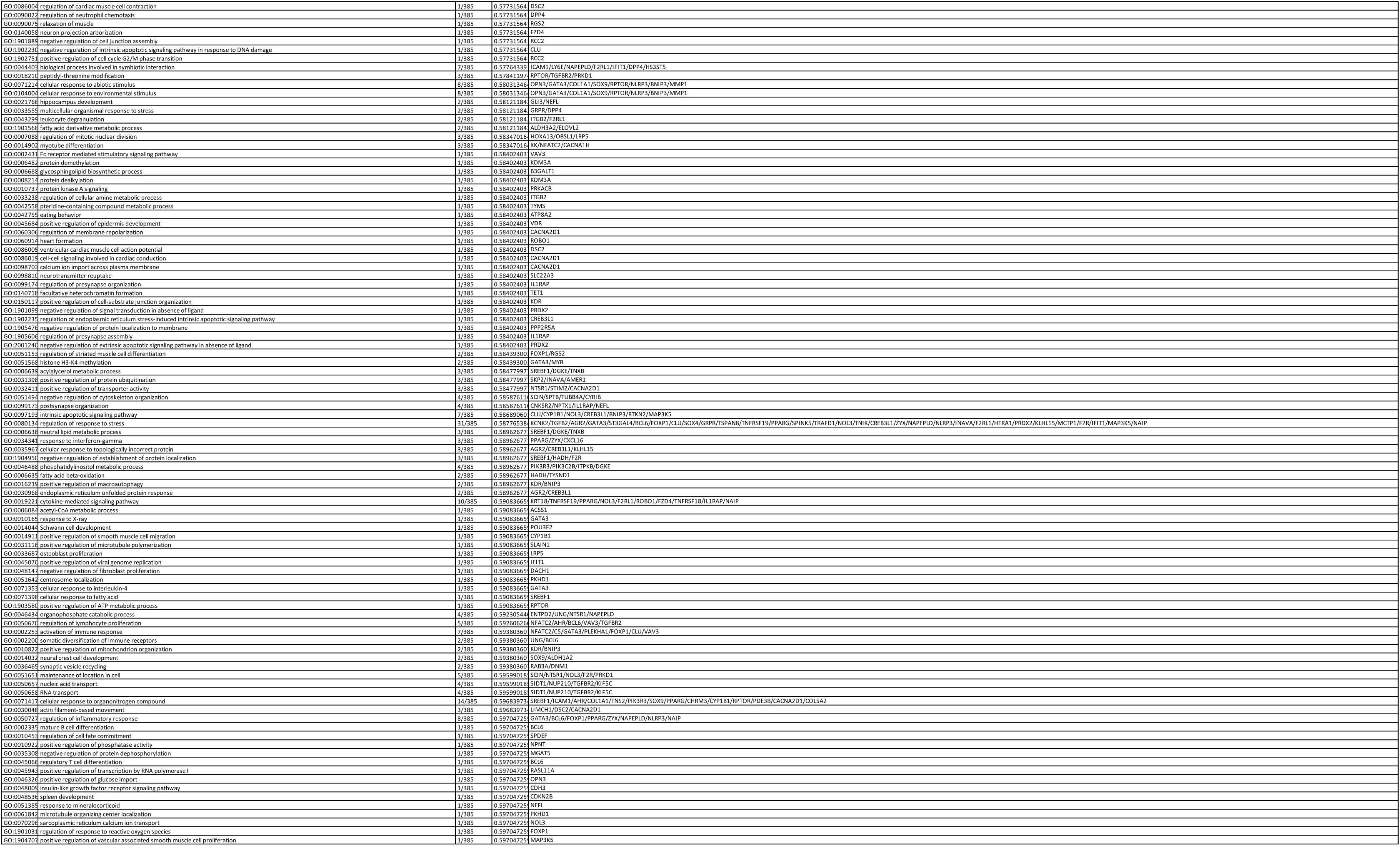
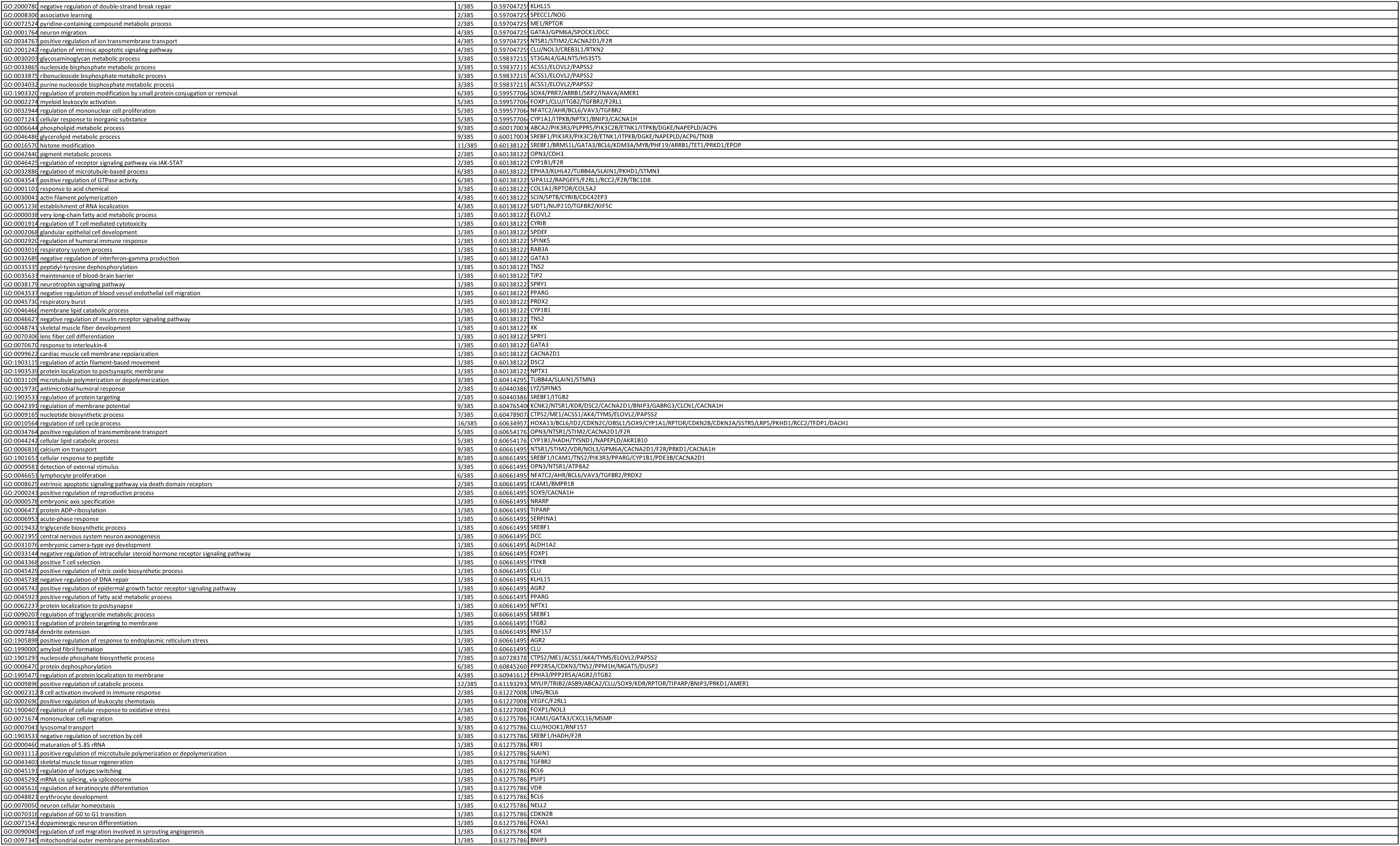
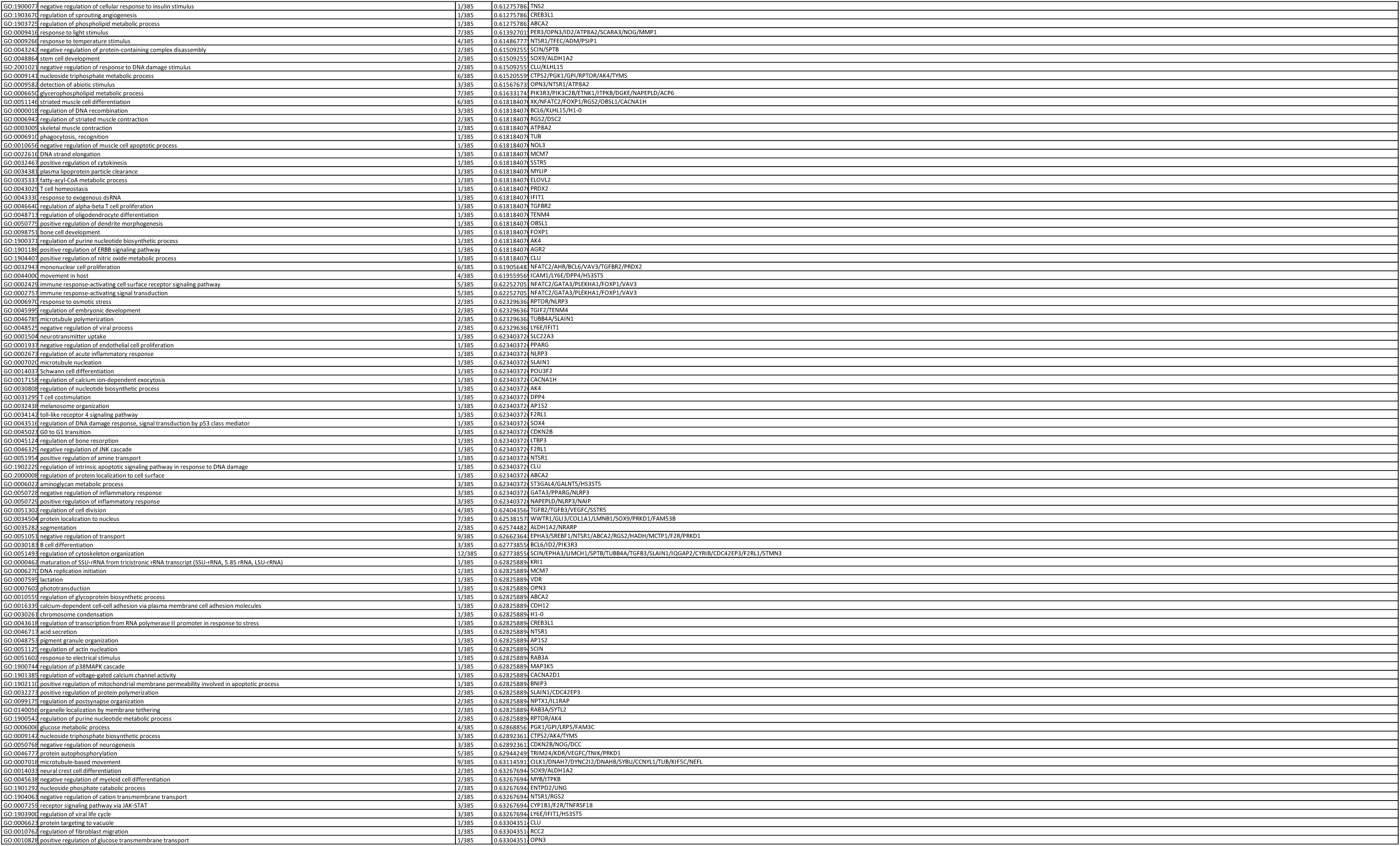
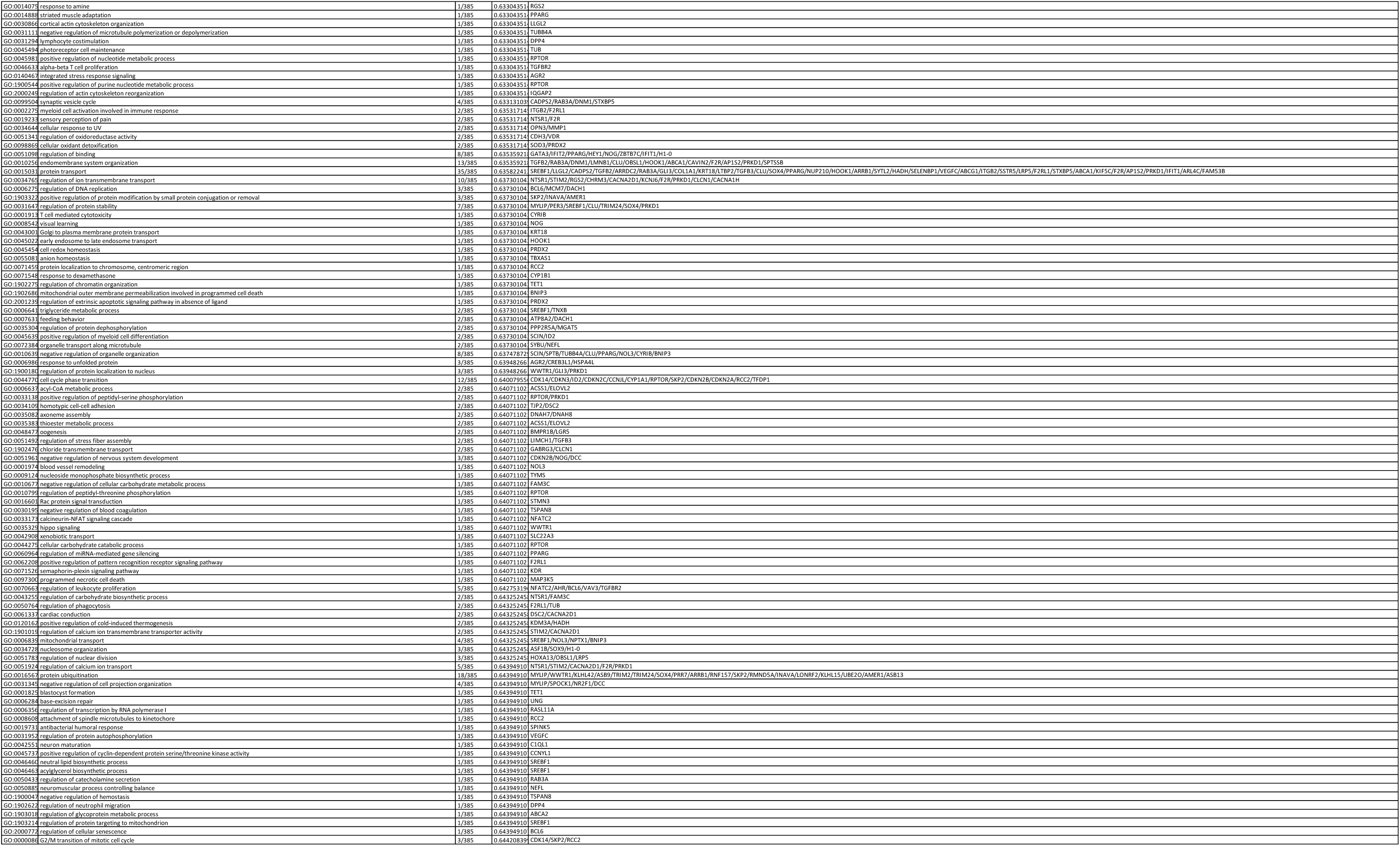
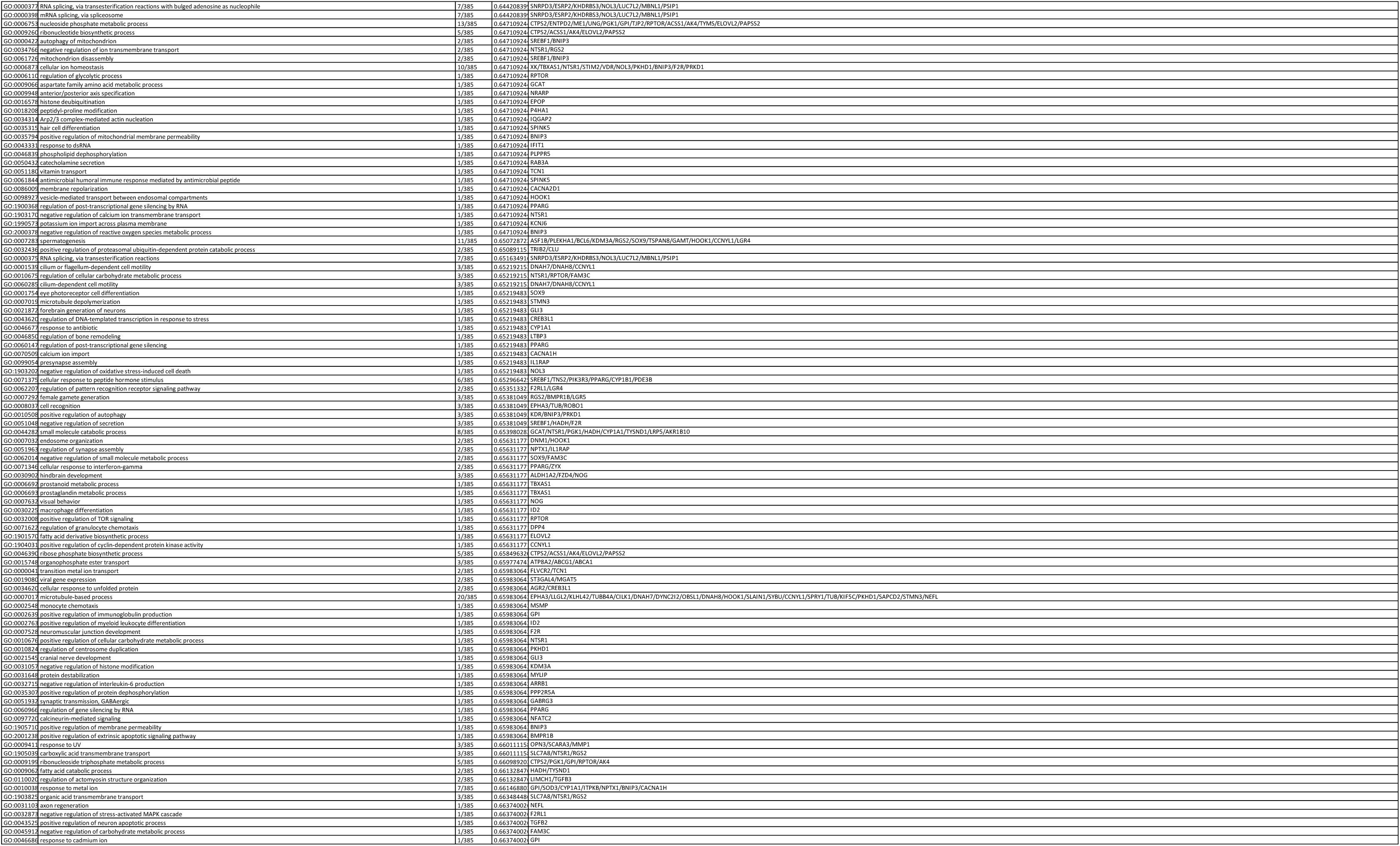
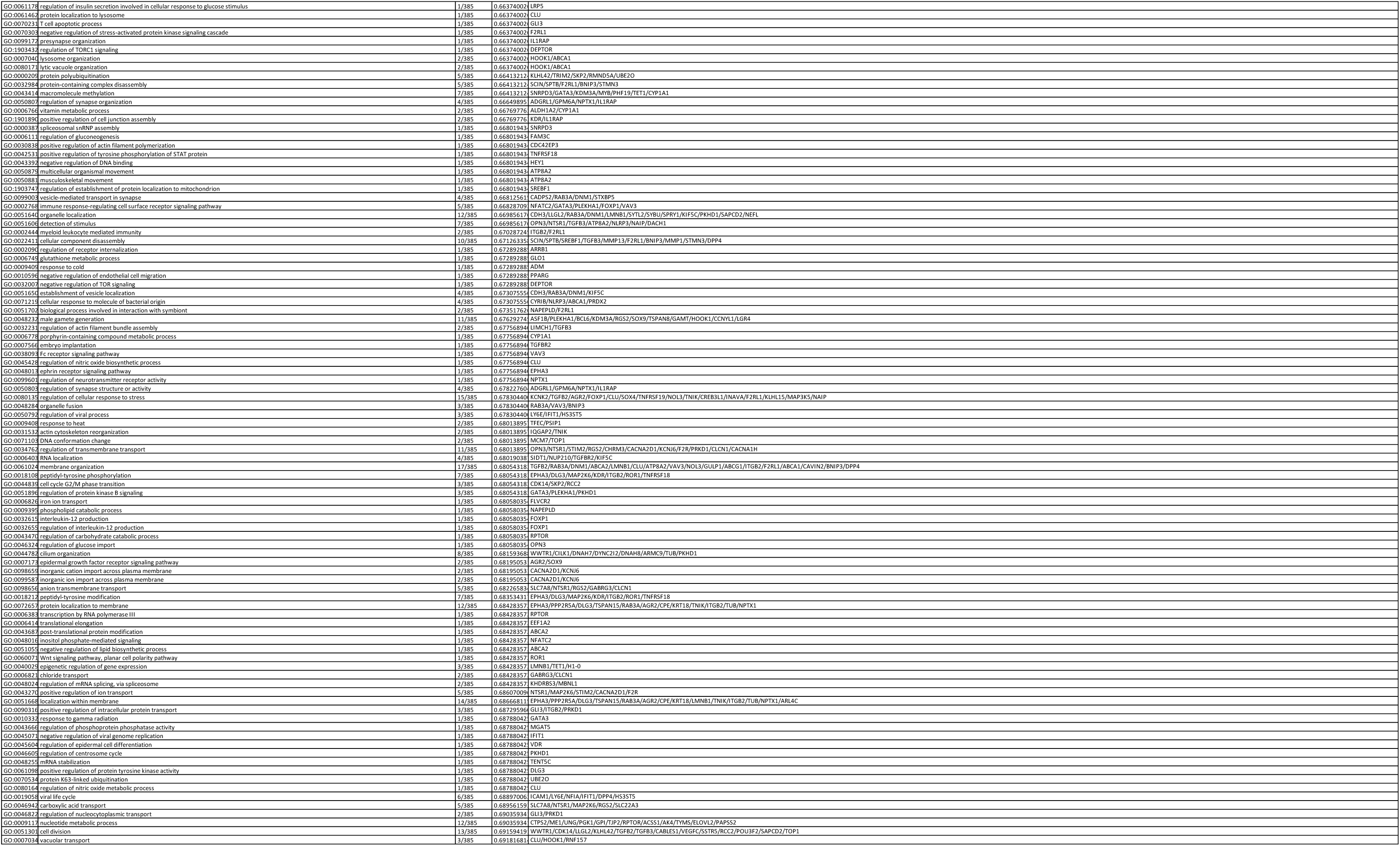
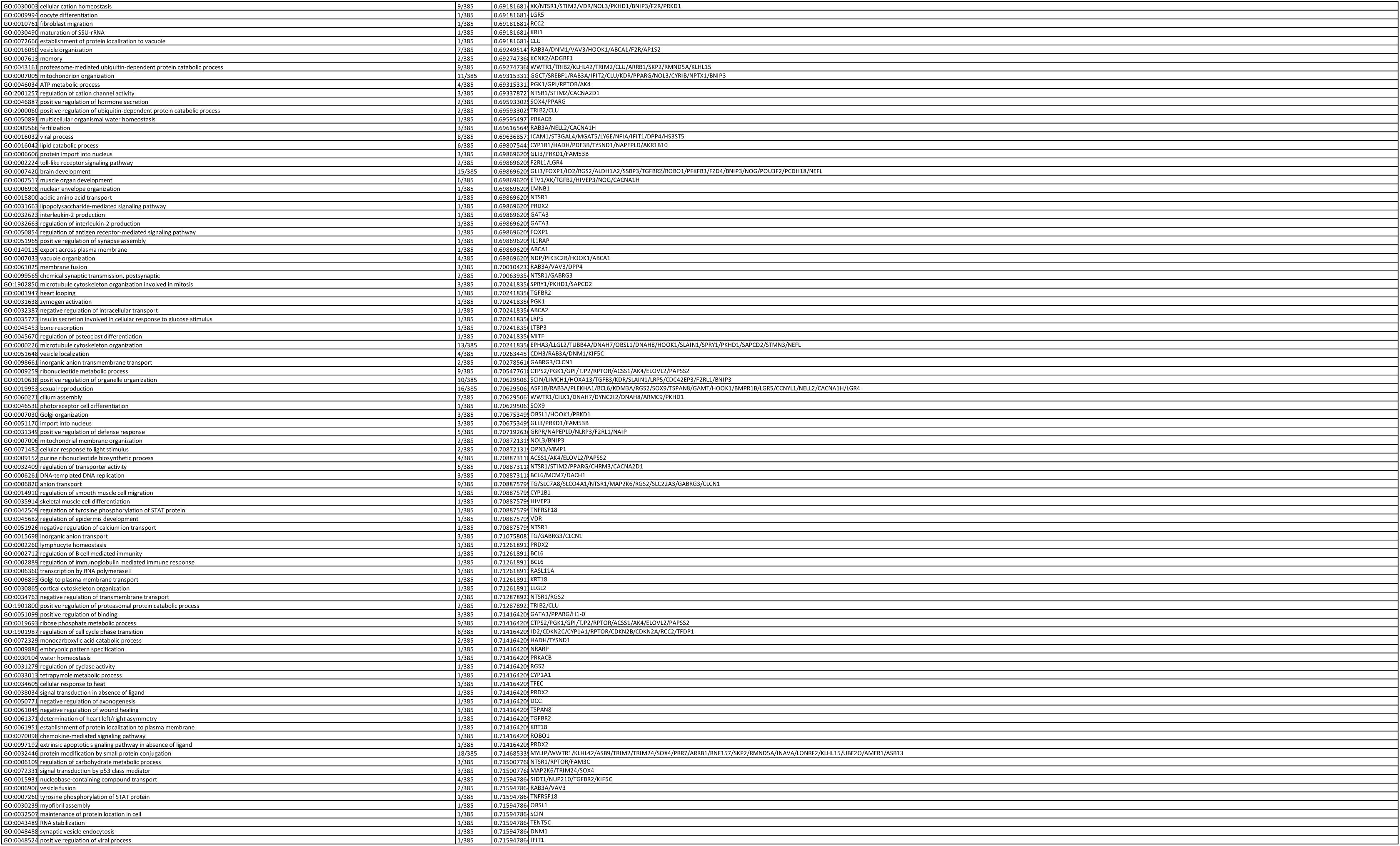
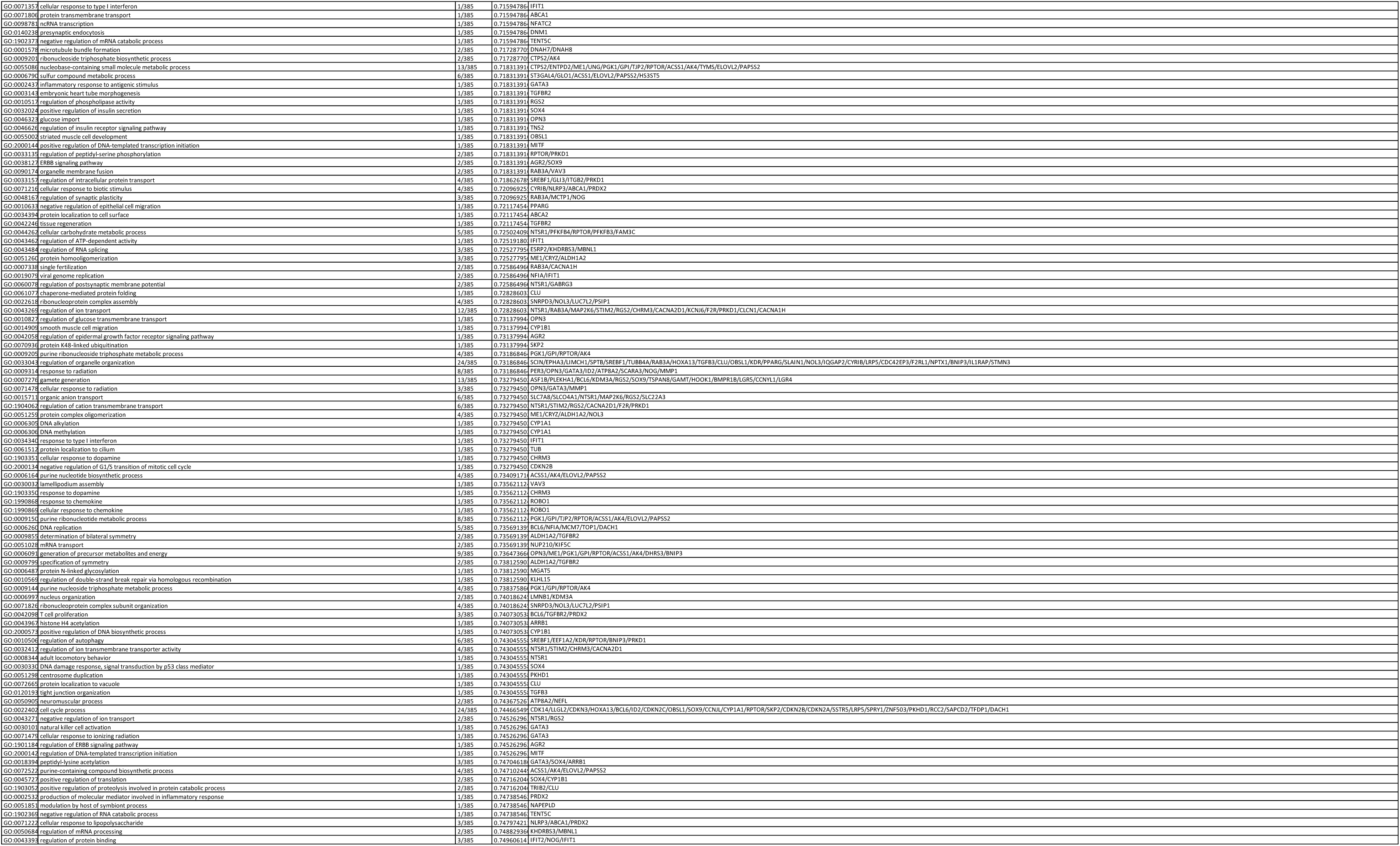
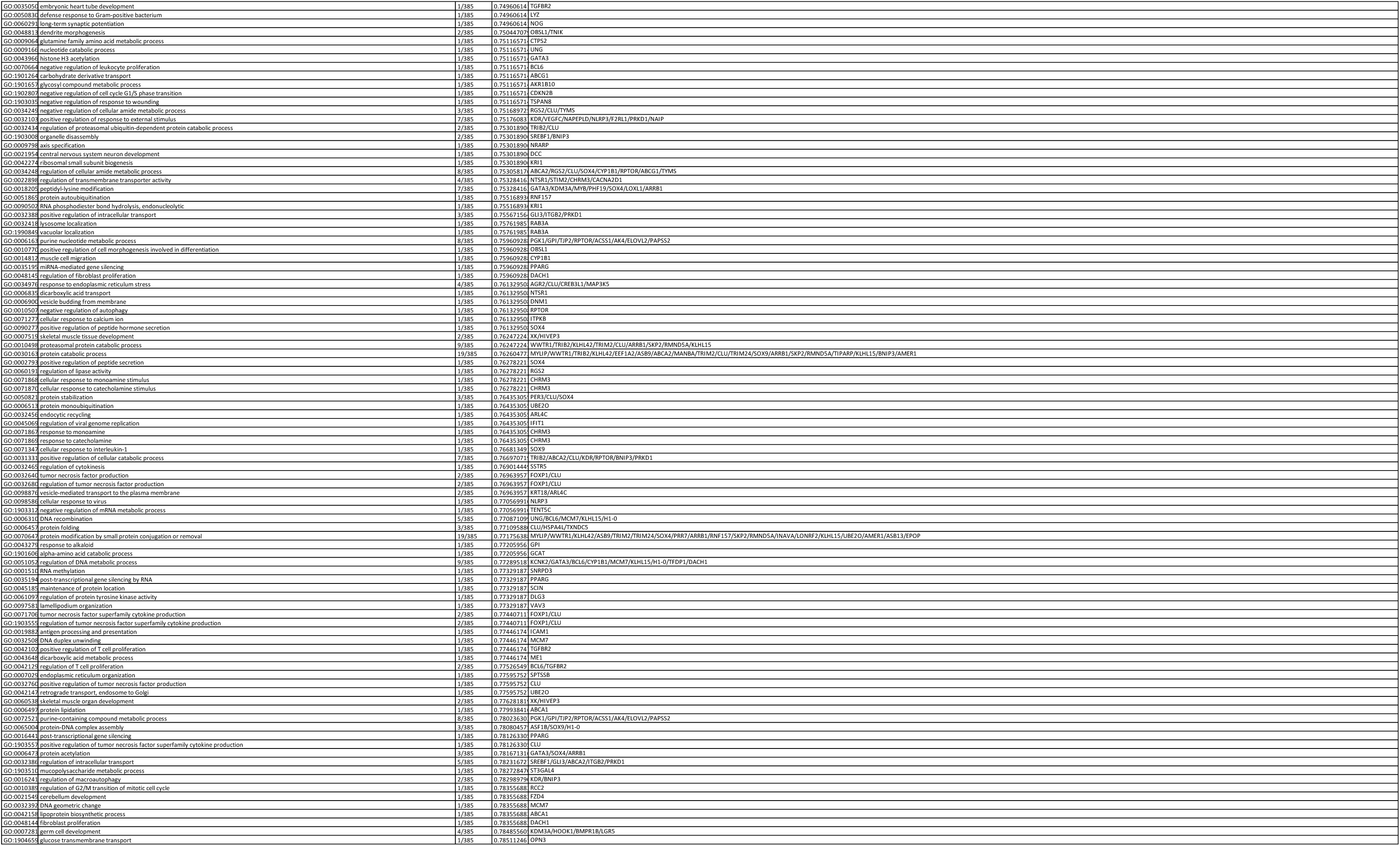
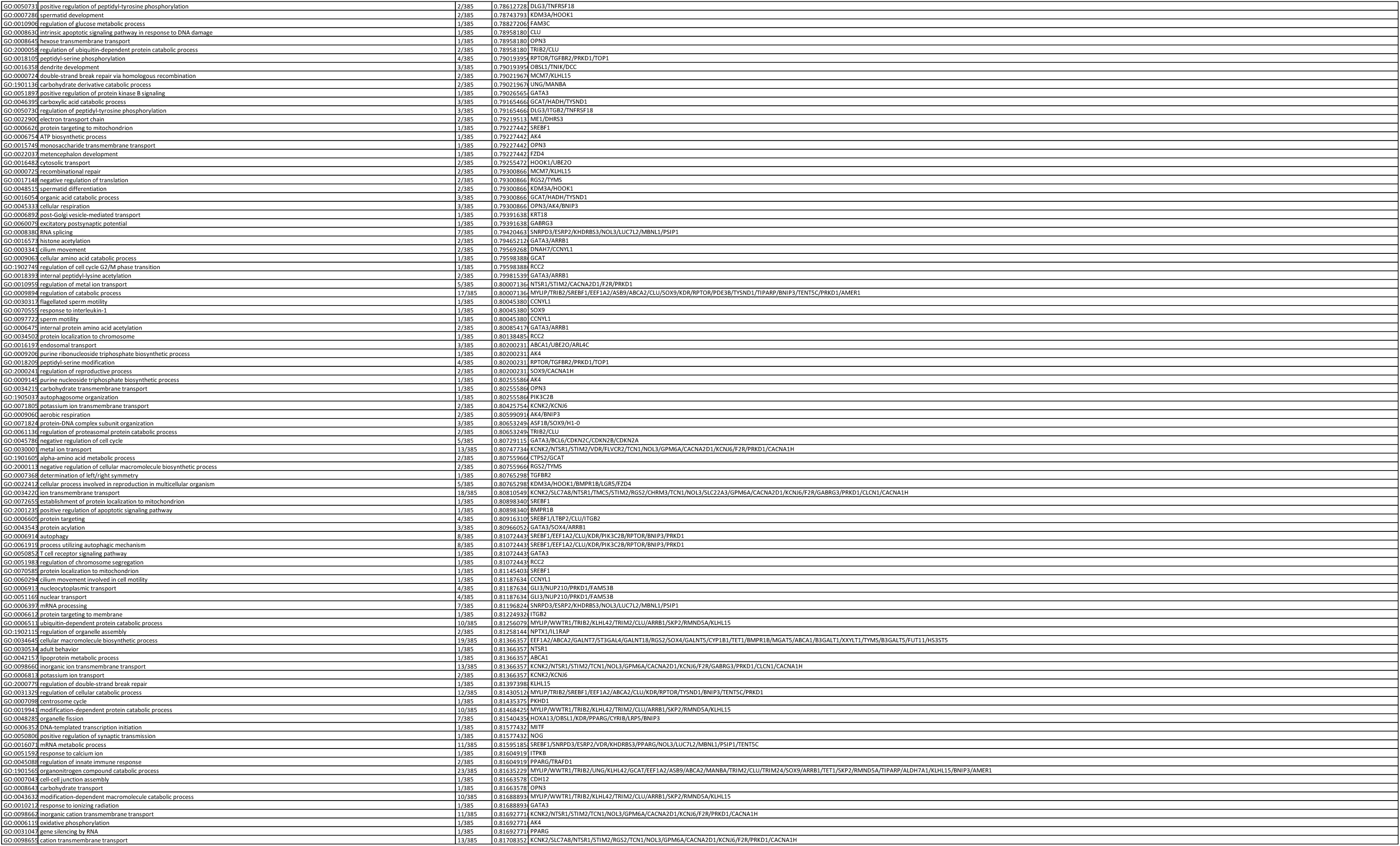
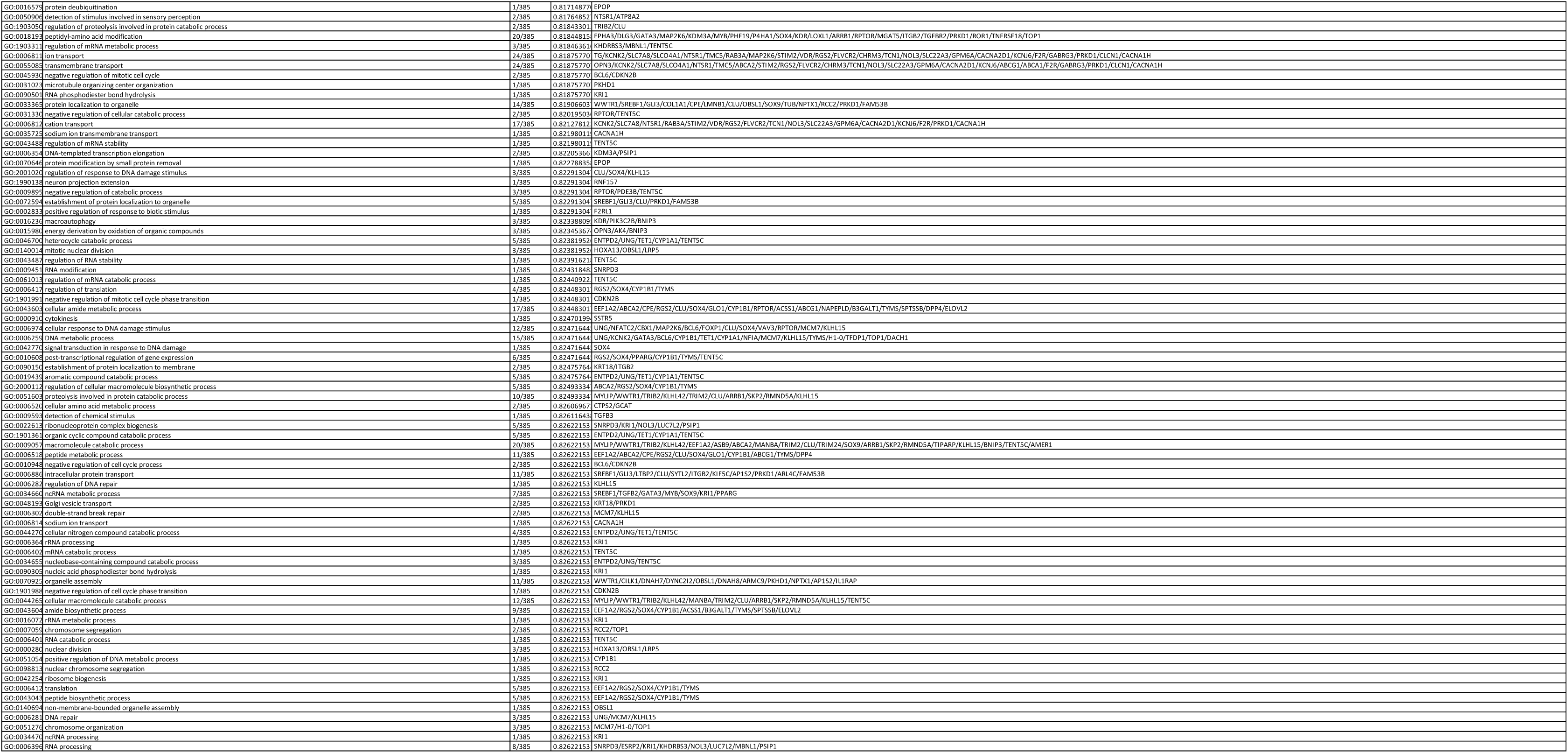
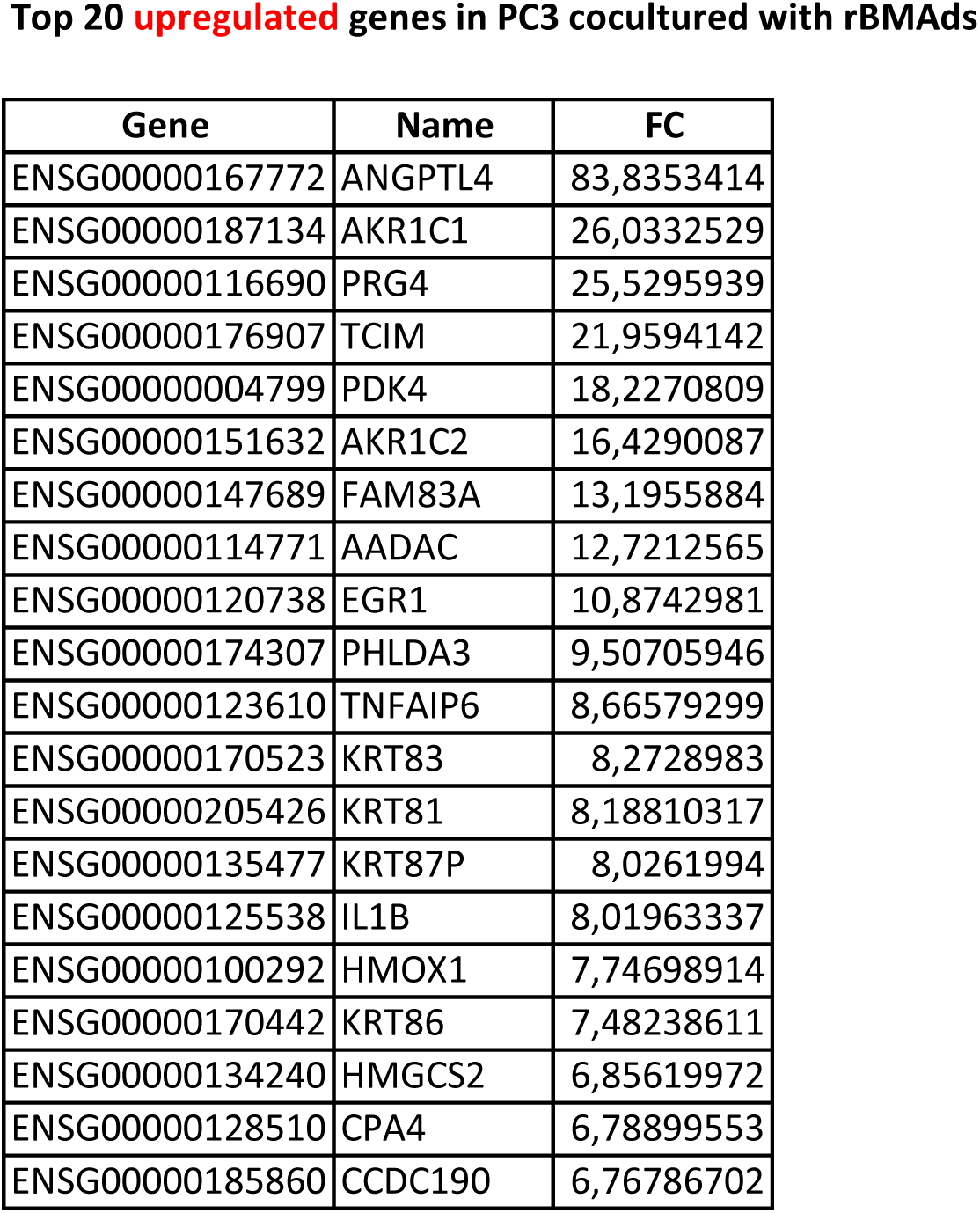
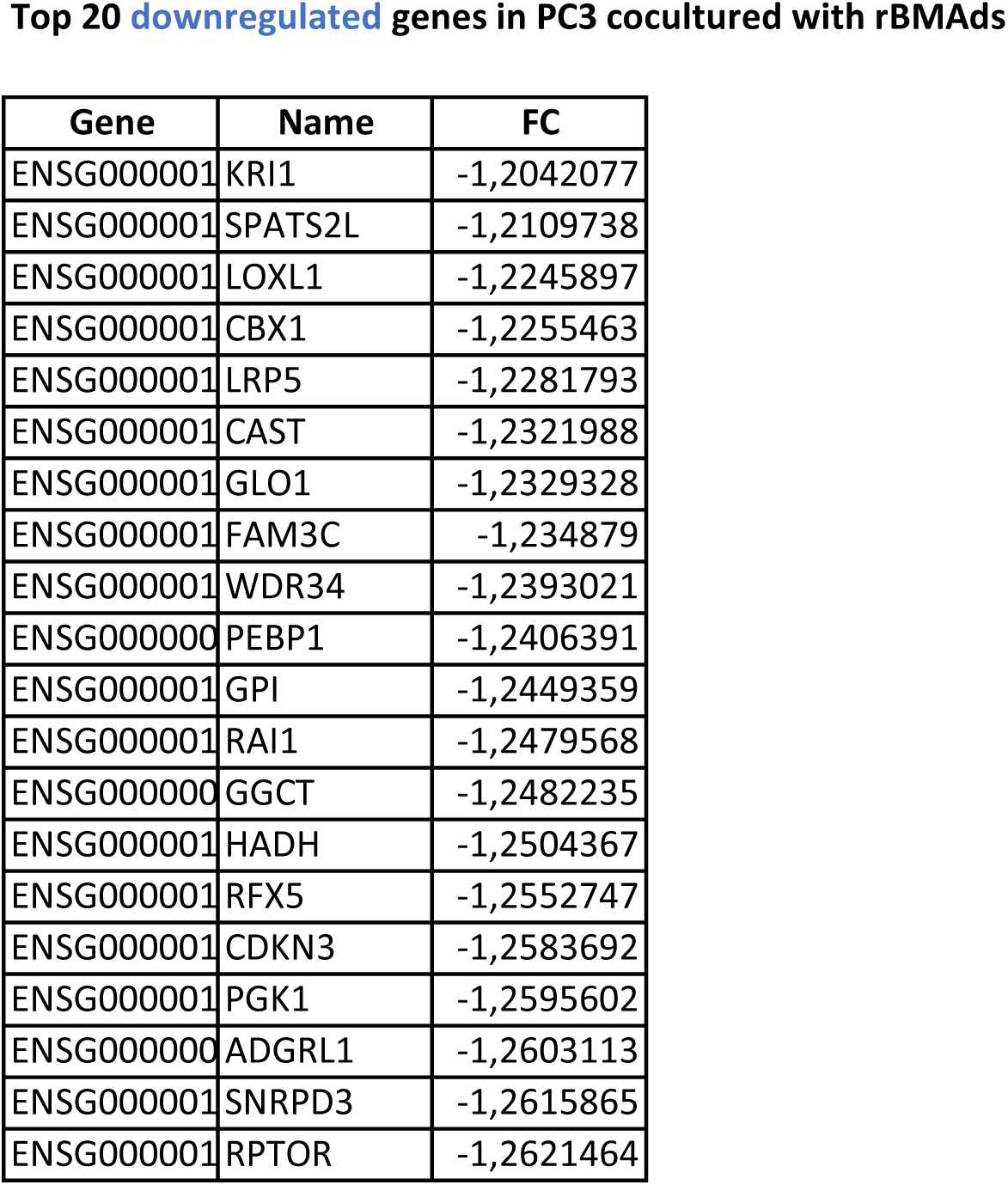
Gene Ontology (GO) analysis of differentially expressed genes in PC3 cells cocultured or not with rBMAds. List of upregulated (A) and downregulated (B) GO terms by the coculture with rBMAds including GO term ID, GO term name, Gene Ratio (number of up- and downregulated genes relative to the total number of genes within each GO term) and q-value of each GO term and gene name present in each GO term. (C) Top 20 of the most upregulated and downregulated genes in PC3 COC. Fold change (FC) values are represented. Data obtained from RNA-seq analysis.

Taken together, these results showed that rBMAds promote migration and invasion of PCa cells suggesting that these adipocytes actively contribute to metastatic dissemination within the bone microenvironment leading to the polymetastatic nature of late-stage disease.

### Adipocyte-derived FFAs drive PCa cell motility

Our results highlight that changes in the transcriptional landscape are key to the observed increase in motility induced by coculture, as the most upregulated pathways were related to cell migration. Since FFAs act as regulators of gene expression that impact tumor progression^47^, we first investigated whether these lipids were involved in the increased migration observed. Conditioned medium (CM) was prepared from rBMAds, and lipids were removed using lipid removal reagent. FFA quantification confirmed a significant reduction in lipid content in the delipidated CM (dCM) compared to the untreated CM (Fig. 4A). To assess the functional impact of lipid depletion, PC3 cells were treated with either CM or dCM, and intracellular lipid accumulation was evaluated. Bodipy staining showed a clear reduction in LDs in PC3 cells exposed to dCM (Fig. 4B), which was further supported by a decrease in TG content (Fig. 4C). Boyden chamber assays demonstrated that CM increased PC3 cell migration (consistent with coculture experiments), whereas this effect was lost when cells were exposed to dCM (Fig. 4D).

**Figure 4:**
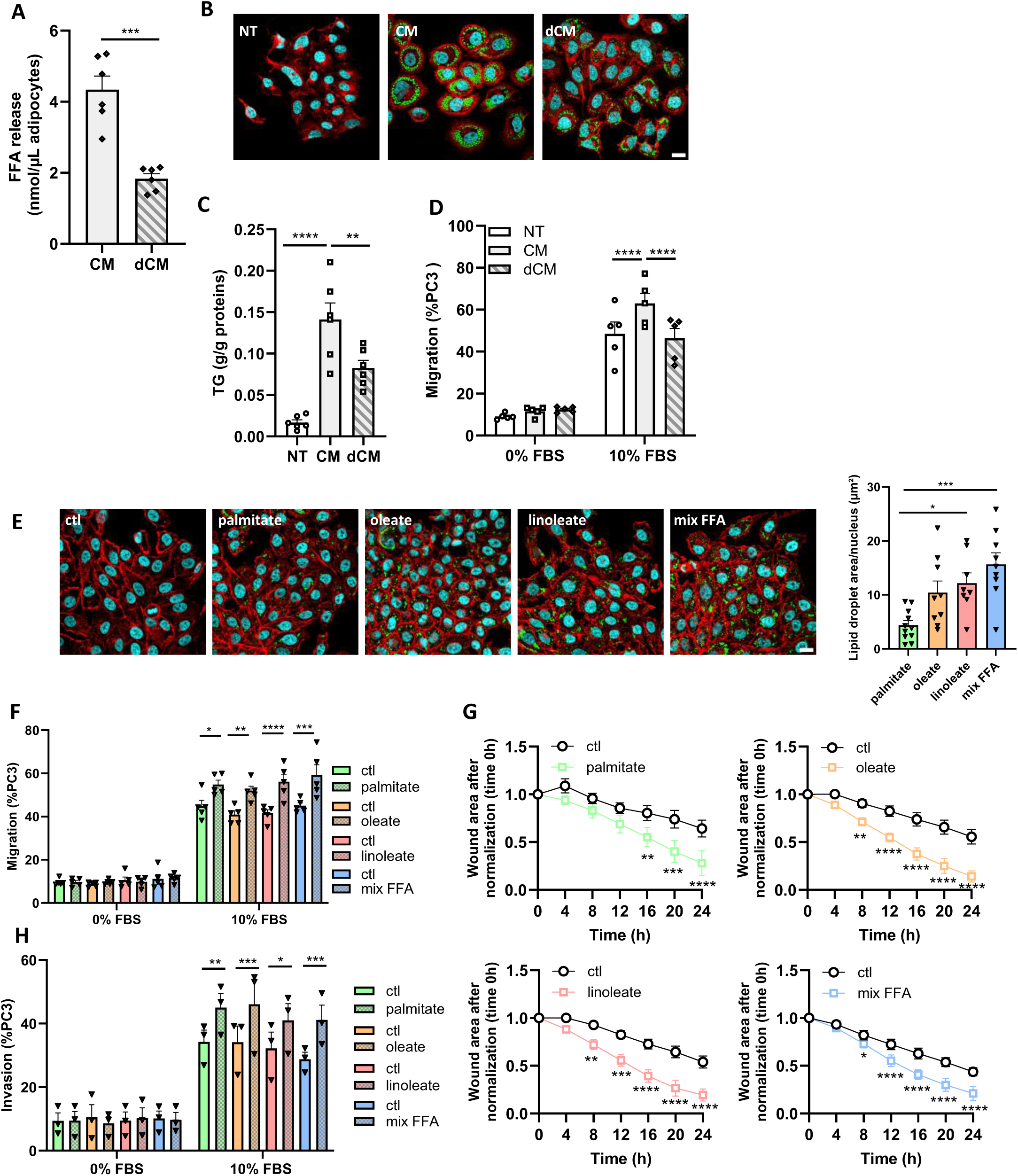
Adipocyte-derived FFAs drive PCa cell motility. **A.** FFA contained in delipidated (dCM) or control conditioned medium (CM) (n=6). **B.** Representative images of PC3 non-treated (NT) or treated with CM or dCM during 72h. Cells were stained with rhodamine-phalloidin (red) for F-Actin, DAPI (cyan) for nuclei and Bodipy 493/503 (green) for neutral lipids. Scale bar, 20µm. **C.** Quantification of TG contained in PC3 NT or treated with CM or dCM during 72h (n=6). **D.** Boyden chamber assay of PC3 previously treated during 72h without (NT) or with CM or dCM (n=5). **E.** Representative images of PC3 cultivated in presence or not (ctl) of exogenous free fatty acid (FFA): 100µM palmitate, 100µM oleate, 100µM linoleate or the mix of the three-FFA at 100µM each (mix FFA). Cells were stained with rhodamine-phalloidin (red) for F-Actin, DAPI (cyan) for nuclei and Bodipy 493/503 (green) for neutral lipids. Scale bar, 20µm. Quantification of lipid droplet area was performed with ImageJ/Fiji® software (Image J, Bethesda, MD, USA) and normalized by the number of nucleus (n=10 palmitate, n=9 oleate, n=7 linoleate, n=10 mix FFA). **F.** Boyden chamber assay of PC3 previously treated or not with palmitate, oleate, linoleate or the mix FFA for 72h (n=5). **G.** Scratch assay of PC3 previously treated or not with palmitate, oleate, linoleate or the mix FFA for 72h. The wound was normalized to time 0 (n=5). **H.** Invasion assay of PC3 previously treated or not with palmitate, oleate, linoleate or the mix FFA for 72h (n=3). Bar plot and curve represent mean ± SEM, ns non-significant, *P < 0.05, **P < 0.01, ***P < 0.001, ****P < 0.0001; (A) was analyzed by paired Student’s t-test, (E) was analyzed by one-way ANOVA with Tukey’s multiple comparison test, (C, D, F, G, H) were analyzed by two-way ANOVA with post hoc Sidak’s multiple comparisons test.

To further evaluate the contribution of FFAs, PC3 cells were treated for 72 hours with palmitate, oleate, or linoleate, the most abundant FFAs released by rBMAds (as shown in Fig. 1E). Exogenous FFA treatment led to neutral lipid accumulation (Fig. 4E) and increased cell motility in both Boyden chamber assays (Fig. 4F) and scratch assays (Fig. 4G). Similarly, FFA treatment enhanced migration of Du145 and LNCaP cells in the Boyden chamber assay (Supplementary Fig. 4A-B). In addition, exogenous FFA treatment increase invasion of PC3 (Fig. 4H). These findings demonstrate that FFAs released by rBMAds contribute to the pro-migratory phenotype of PCa cells.

### ANGPTL4 as a FFA-responsive driver of PCa cell migration

Among the top 10 most upregulated genes in the coculture condition, ANGPTL4 exhibited the highest induction, with an 80-fold increase in expression in PC3 COC compared to PC3 NC (Fig. 5A). ANGPTL4 is a glycoprotein primarily known for its role in lipid metabolism, particularly as an inhibitor of lipoprotein lipase (LPL), thereby regulating plasma TG levels^48^. ANGPTL4 has also been implicated in various cancer-related processes, including cell survival, proliferation, angiogenesis, and metastasis, acting in both autocrine and paracrine manners^49^. In most models, ANGPTL4 promotes tumor cell motility via cytoskeletal remodeling, integrin activation, and extracellular matrix (ECM) degradation^50–54^. In PCa, immunohistochemical analysis of ANGPTL4 expression in PCa tissues from radical prostatectomy showed that its expression was significantly correlated with PSA (Prostate Specific Antigen) recurrence after treatment^55^. Interestingly, ANGPTL4 is a known target of the PPARγ transcription factor, which binds and responds to diverse lipid metabolites^48,56–58^.

**Figure 5:**
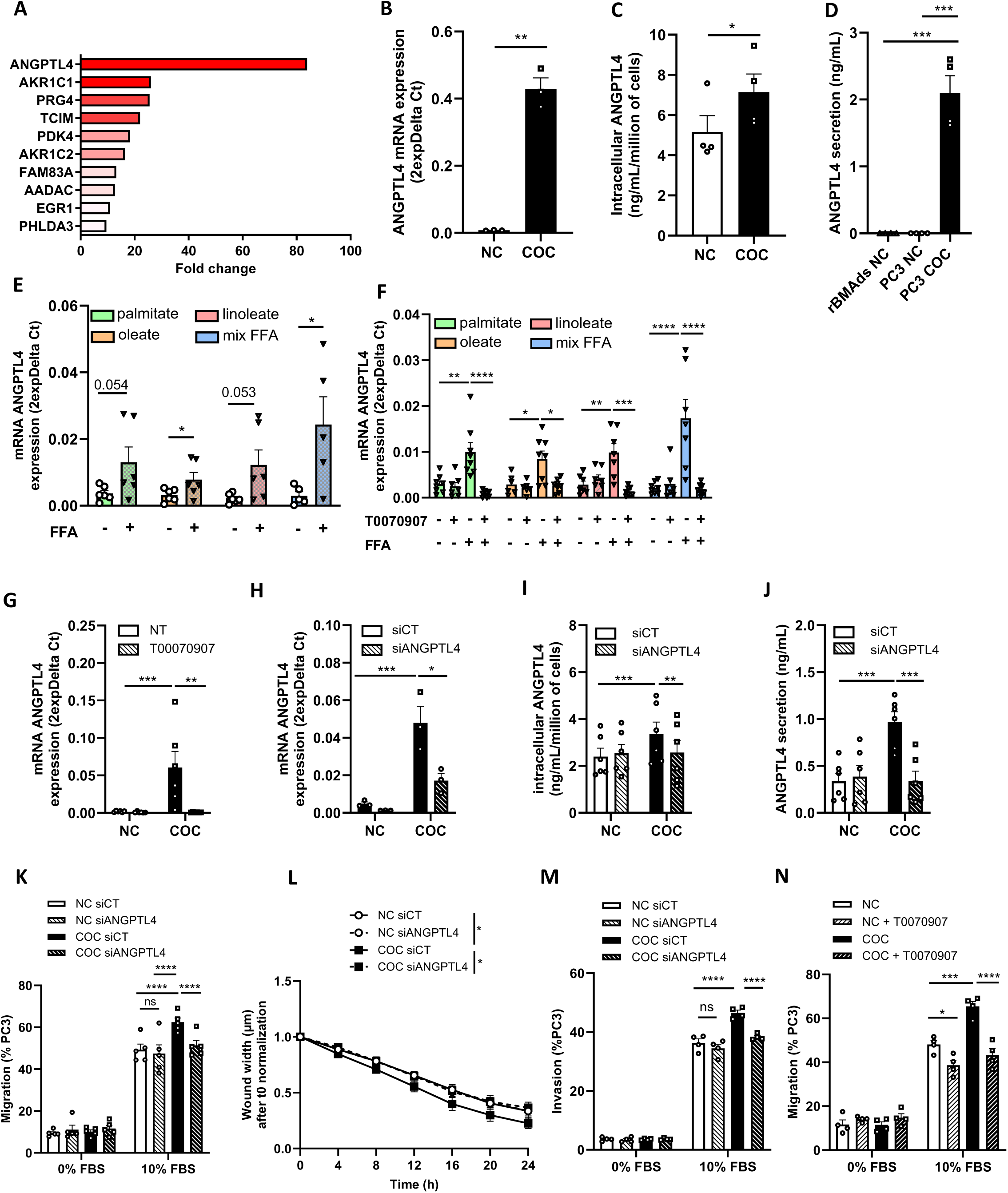
ANGPTL4 as a FFA-responsive driver of PCa cell migration. **A.** Fold change of the top 10 most upregulated genes induced by coculture in PC3 and identified by RNAseq. **B.** mRNA expression of ANGPTL4 in PC3 NC and COC after 72h of culture (n=3). **C.** Intracellular level of ANGPTL4 in PC3 NC and COC after 72h of culture (n=4). **D.** Quantification of ANGPTL4 contained in the medium of rBMAds NC, PC3 NC and PC3 COC after 72 hours of culture (n=4). **E.** mRNA expression of ANGPTL4 in PC3 after 72h of treatment with exogenous FFA: palmitate, oleate, linoleate or mix of FFA or without treatment (ctl) (n=6). **F.** mRNA expression of ANGPTL4 in PC3 treated 24h with exogenous FFA in presence or in absence of PPARγ inverse agonist (10µM T0070907) (n=8). **G.** mRNA expression of ANGPTL4 in PC3 NC and COC during 24h in presence or in absence (non-treated, NT) of PPARγ inverse agonist (10µM T0070907) (n=6). **H.** mRNA expression of ANGPTL4 in PC3 NC and COC transfected with siRNA control (siCT) or with siRNA against ANGPTL4 (siANGPTL4) and after 24h of coculture (n=3). **I.** Intracellular level of ANGPTL4 in PC3 NC and COC transfected with siCT or siANGPTL4 and after 24h of culture (n=6). **J.** Quantification of ANGPTL4 contained in the medium of PC3 NC and COC transfected with siCT or siANGPTL4 and after 24h of culture (n=6). **K.** Boyden chamber assay of PC3 NC and COC transfected with siCT or siANGPTL4 (n=5) after 24h of culture. **L.** Scratch assay on PC3 NC and PC3 COC transfected with siCT or siANGPTL4 was performed after 24h of culture. The wound was normalized to time 0 (n=5). **M.** Invasion assay of PC3 NC and COC transfected with siCT or siANGPTL4 after 24h of culture (n=4). **N.** Boyden chamber assay of PC3 NC and COC treated or not during 24h with PPARγ inverse agonist (10µM T0070907) (n=4). Bar pot and curve represent mean ± SEM, ns non-significant, *P < 0.05, **P < 0.01, ***P < 0.001, ****P < 0.0001; (B, C, E) were analyzed by paired Student’s t-test except for (E) with unpaired Student’s t-test, (D) was analyzed by one-way ANOVA with Tukey’s multiple comparison test, (F-N) were analyzed by two-way ANOVA with Tukey’s multiple comparisons test.

Based on RNA-seq results, we confirmed that PC3 COC exhibit a significant increase in ANGPTL4 mRNA levels using independent RNA extracts (Fig. 5B) as well as increased protein expression (Fig. 5C), compared to PC3 NC. Similar results were obtained in DU145 cells, which also exhibited increased ANGPTL4 expression following coculture (Supplementary Fig. 5A–B). In contrast, ANGPTL4 expression was undetectable in LNCaP cells under both basal conditions and after coculture (data not shown), despite the increased migration observed in these cells, highlighting the biological heterogeneity of PCa and indicating that not all cell lines recapitulate the same molecular response to adipocyte-derived FA. As previously shown^56–59^, ANGPTL4 may be secreted, and we confirmed its secretion in both PC3 (Fig. 5D) and Du145 cells after coculture (Supplementary Fig. 5C), whereas it was undetectable in rBMAds or cancer cells cultivated alone (Fig. 5D; Supplementary Fig. 5C).

To assess whether ANGPTL4 overexpression could be driven by exogenous lipids, PC3 cells were treated during 72 h with palmitate, oleate or linoleate alone or in combination (mix FFA). This led to a significant increase in ANGPTL4 gene expression in PC3 (Fig. 5E) and Du145 cells (Supplementary Fig. 5D). Given that ANGPTL4 is regulated by PPARγ, whose activity is modulated by FFAs^56,60,61^, we tested this regulatory axis. Treatment of PCa cells for 24 hours with FFAs in the presence or absence of T0070907, a PPARγ inverse agonist, showed that FFA-induced ANGPTL4 expression was completely abolished by the antagonist (Fig. 5F; Supplementary Fig. 5E). Similarly, 24h of coculture-induced ANGPTL4 expression was also blocked by PPARγ inverse agonist (Fig. 5G; Supplementary Fig. 5F). To test whether ANGPTL4 contributes functionally to PCa migration, we silenced ANGPTL4 using siRNA prior to 24h of coculture. Silencing was confirmed at both mRNA (Fig. 5H) and protein levels (Fig. 5I-J) in PC3, as well as in Du145 cells (Supplementary Fig. 5G-I). Knockdown of ANGPTL4 in PCa cells significantly reduced migration in Boyden chamber assay (Fig. 5K, Supplementary Fig. 5J) and wound closure in scratch assay (Fig. 5L) which has been enhanced in co-culture. ANGPTL4 silencing also decreased invasion capacities of PC3 cells, reinforcing its role in motility (Fig. 5M). Finally, to confirm the role of PPARγ activity in this process, PCa cells were treated with a PPARγ inverse agonist during 24 h coculture with rBMAds. This treatment effectively abolished the migration-promoting effect of coculture, confirming that PPARγ activity is required for the observed pro-migratory phenotype (Fig. 5N; Supplementary Fig. 5K). Mechanistically, ANGPTL4 has been reported to promote migration through actin cytoskeleton remodeling and pseudopodia formation^53^ and to induce an EMT-like transcriptional program^62–64^. To further investigate the mechanisms underlying ANGPTL4-dependent cell migration, we reconstructed an ANGPTL4-associated gene network in Cytoscape using RNA-seq-derived differentially expressed genes enriched in cell migration GO terms and their interactions with ANGPTL4. This analysis revealed a highly interconnected network rather than a linear signaling cascade (Supplementary Fig. 5L), suggesting that ANGPTL4 promotes cell migration by orchestrating a coordinated transcriptional program involving multiple complementary signaling pathways.

Taken together, our findings identify ANGPTL4 as a critical FFA-responsive effector that mediates PCa cell migration in response to adipocyte-derived lipid signals. Herein, we show that ANGPTL4 expression is robustly induced in cocultured PCa cells and this effect is driven by FFAs and depends on PPARγ activity. Most importantly, functional experiments using siRNA-mediated knockdown clearly demonstrate that downregulation of ANGPTL4 significantly impairs the adipocyte-driven migration and invasion of PCa cells. Taken together, these results establish a direct functional link between ANGPTL4 expression and tumor cell motility in aggressive cell lines. Thus, ANGPTL4 is not only a marker of lipid-induced transcriptional reprogramming but also a key driver of the migratory phenotype in PCa.

### High expression of ANGPTL4 contribute to poor overall survival in PCa patients with bone metastases

To determine whether ANGPTL4 expression is specifically induced by rBMAds, PC3 cells were cocultured with either human rBMAds or adipocytes isolated from human periprostatic adipose tissue (PPAT) found at the primary tumor site in PCa. Although PPAT adipocytes also induced ANGPTL4 expression in PCa cells compared with NC cells, this induction was 2.5-fold higher in cells cocultured with rBMAds (Fig. 6A). Consistent with these findings, analysis of a public datasets of human PCa samples revealed increased ANGPTL4 expression in bone lesions compared with primary tumors (Fig. 6B). Thus, although ANGPTL4 expression may be initiated within the primary tumor microenvironment, it is markedly amplified following colonization of the bone marrow niche where it promotes PCa cell migration and metastatic outgrowth.

**Figure 6:**
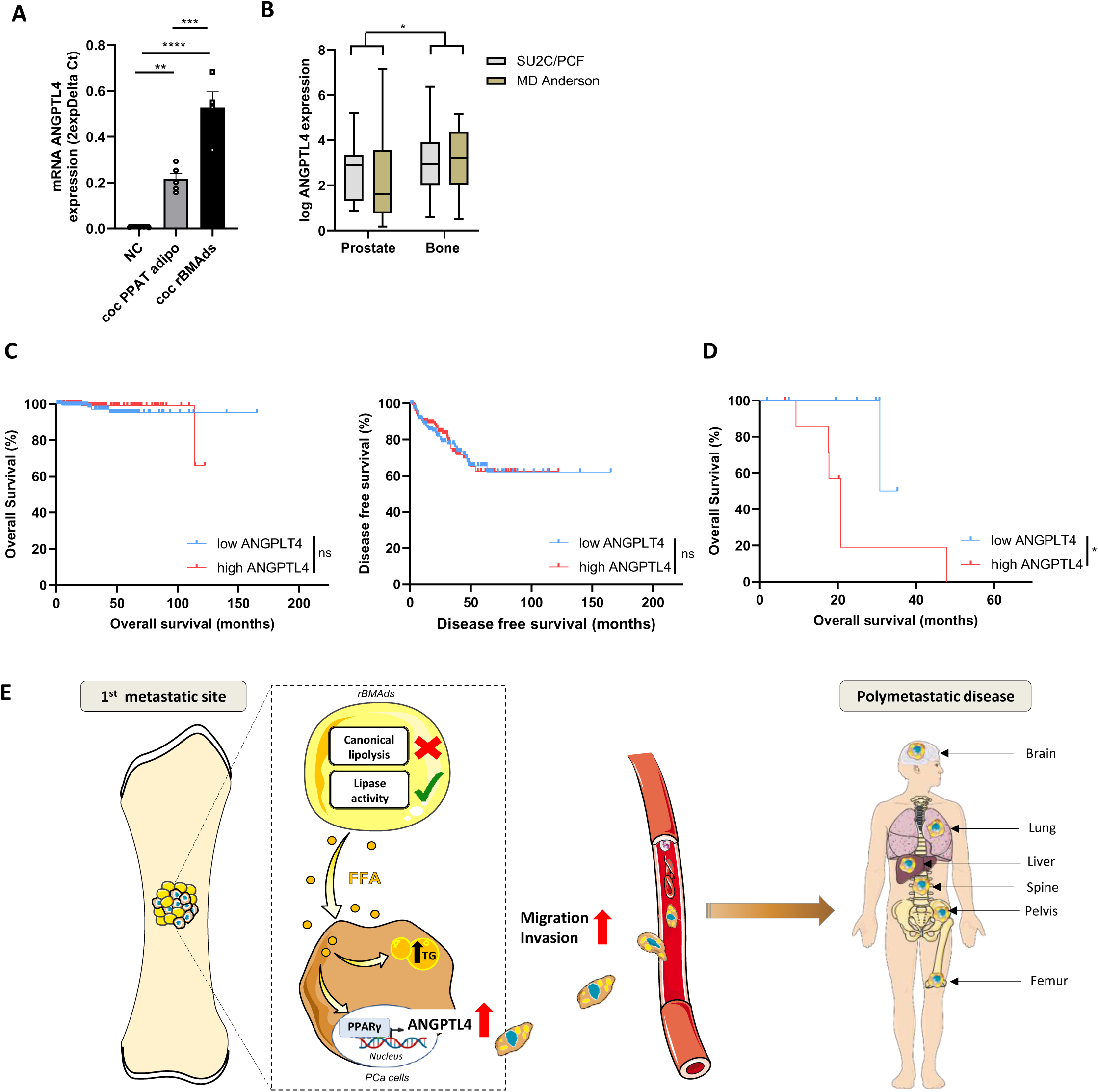
High expression of ANGPTL4 contribute to poor overall survival in PCa patients with bone metastases. **A.** mRNA expression of ANGPTL4 in PC3 NC and COC after 72h of coculture with periprostatic adipocytes or rBMAds (n=5 periprostatic adipocytes, n=4 rBMAds). **B**. ANGPTL4 expression in primary prostate tumors and bone metastases in the *Metastatic Prostate Adenocarcinoma* (*SU*2*C*/*PCF Dream Team*^38^, prostate n=7 and bone n=73) and the *Prostate Cancer MDA PCa PDX* (*MD Anderson*^39^ ; prostate n=21 and bone n=9). **C.** Overall survival (left) and disease-free survival (right) of primary PCa patients with low and high ANGPTL4 expression in the TGA dataset (overall survival: low, n=151; high, n=151; disease-free survival: low, n=153; high, n=153). ns non-significant. **D.** Overall survival of bone metastasis PCa patients with low (n=8) and high (n=8) level of ANGPTL4 expression. *P < 0.05. **E.** Proposed model of how rBMAds participate to tumor progression. At the bone metastatic site, a metabolic crosstalk occurs between rBMAds and PCa cells. rBMAds release FFA via a non-canonical lipolytic pathway. These FFAs are taken up by PCa cells inducing a transcriptional reprogramming, leading to the induction of ANGPTL4 on a PPARγ dependent mechanism that promotes motility. Through this signaling pathway, rBMAds may promote the development of polymetastatic disease by facilitating the establishment of multiple metastatic sites in the same organ or in different organs (brain, lung, liver, spine, pelvis).

We next investigated the clinical relevance of ANGPTL4 expression across different stages of PCa progression. Analysis of The Cancer Genome Atlas (TCGA) cohort comprising 459 primary PCa samples revealed that high ANGPTL4 expression in primary tumors was not associated with either overall or disease-free survival (Fig. 6C), indicating that the clinical relevance of ANGPTL4 is not linked to the primary tumor. In contrast, analysis of a clinically annotated dataset^38^ of 26 bone metastasis cases, revealed that high ANGPTL4 expression was associated with reduced overall survival (Fig. 6D), identifying ANGPTL4 as a marker of poor clinical outcome at the terminal stage of the disease. Interestingly, ANGPTL4 expression was maintained across metastases at other anatomical sites, including the liver, lung and lymph nodes, and was further increased in liver metastases (Supplementary Fig. 6A). Given that bone represents the predominant metastatic site in PCa, accounting for approximately 80% of metastatic lesions^5^, and that secondary metastases arise from tumor cells initially established in the bone microenvironment^65,66^, these observations suggest that sustained ANGPTL4 expression is maintained during late-stage metastatic dissemination.

Collectively, our findings support a model in which ANGPTL4 is selectively induced by rBMAds within the bone marrow niche, thereby promoting aggressive disease and contributing to poor clinical outcome in patients with PCa bone metastases.

### Concluding remarks

Using a 3D model of primary human rBMAds, we identify a previously unrecognized metabolic crosstalk between rBMAds and PCa cells, in which FFAs are released via an unconventional lipase-dependent mechanism. These FFAs are taken up by PCa cells, triggering gene remodeling through PPARγ-dependent induction of ANGPTL4 and enhancing migration and invasion. While ANGPTL4 expression may be initiated at the primary tumor site, our findings suggest that it is strongly amplified within the bone marrow adipocyte-rich metastatic niche, where it promotes tumor cell motility and metastatic outgrowth. As summarized in Fig. 6E, rBMAd-driven invasion may not only facilitate bone colonization and local tumor progression but also promote the subsequent dissemination of PCa cells to secondary osseous and extra-osseous sites, thereby contributing to the systemic, polymetastatic progression of advanced disease.

Supporting the translational potential of these observations, preclinical studies in other cancer types show that blocking antibodies against ANGPTL4 can inhibit tumor cell colonization *in vivo*^67^ and reduce EMT in 3D tumor models^68^. In addition to rBMAds, cBMAds are also present within the bone marrow. Although they are not typically found in close proximity to PCa metastatic lesions, cBMAds may contribute to tumor progression through similar or distinct mechanisms, an open question that warrants further investigation. Collectively, these results identify ANGPTL4 as a promising target to limit the migratory and invasive properties of PCa cells, warranting validation in *in vivo* models that faithfully recapitulate successive metastatic colonization, which are currently lacking for PCa.

## Supplementary Figure legends

**<u>Supplementary figure 1:</u> Development of 3D culture model of rBMAds and FFA transfer in PCa cells in coculture.**

**A.** Maximum intensity projection of Z-stack acquired through confocal microscopy. Representative images of rBMAds embedded in fibrin gels and stained with Bodipy 493/503 (green), after their isolation (Day 0) or after 3 days (Day 3) or 5 days (Day 5) of culture. Scale bar, 100µm. **B.** Distribution of rBMAds diameter (µm) at Day 0, Day 3 and Day 5 of culture (n=9 paired samples) and the mean diameter (µm) of rBMAds. **C.** Quantification of adiponectin released by rBMAds at Day 1, Day 2 and Day 3 of culture (n=4). **D.** Representative images of PC3, Du145 and LNCaP cultivated alone (NC) or with rBMAds (COC) at 72h of culture. Cells were stained with rhodamine-phalloidin (red) for F-Actin, DAPI (blue) for nuclei and Bodipy 493/503 (green) for neutral lipids. Scale bar, 20µm. **E.** Quantification of triglycerides (TGs) contained in PC3, Du145 and LNCaP NC and COC (n=4 PC3, n=5 Du145, LNCaP). **F.** mRNA expression of CD36 in U937 ctl and KO for CD36 (U937 KO CD36) used as controls and in PCa cell lines (n=3). **G**. CD36 expression by flow cytometry in U937 ctl and KO CD36 and PCa cell lines (n=3). **H.** mRNA expression of FATP genes family, SLC27A1-5 in PC3, Du145 and LNCaP (n=5 PC3; n=3 Du145 and LNCaP**). I.** mRNA expression of SLC27A4 coding for FATP4 protein in PC3 transfected with siRNA control (siCT) or siSLC27A4 (n=4**). J.** Western Blot of FATP4 protein in PC3 transfected with siCT or siSLC27A4. **K.** Quantification of Bodipy FLC16 uptake in PC3 transfected with siCT or siSLC27A4 after 30min, 1h, 2h, 3h and 4h of incubation with Bodipy FLC16 (n=5). **L.** Quantification of fluorescent oleate uptake in PC3 transfected with siCT or siSLC27A4 after 30min, 1h, 2h, 3h and 4h of incubation with fluorescent oleate (n=5). **M.** Quantification by flow cytometry of fluorescence intensity of neutral lipids stained by Bodipy in PC3 NC and COC transfected by siCT or siSLC27A4 (n=6). Bars represent mean ± SEM, ns non-significant; *P < 0.05, **P < 0.01, ***P < 0.001; ****P < 0.0001; N.D.: Non Detected, (B, C, F, H) were analyzed by one-way ANOVA with Tukey’s multiple comparisons test. (E, K, L, N) were analyzed by two-way ANOVA with Sidak’s multiple comparisons test, (I) was analyzed by Student’s t-test.

**<u>Supplementary figure 2:</u> rBMAds exhibit non canonical lipolysis with uncoupled FFA and glycerol release. A.** Glycerol and FFA release by SCAds and rBMAds after 3h, 18h and 24h of stimulation or not with isoproterenol 1µM (n=4). **B.** FFA and glycerol released in culture medium of PC3 NC, rBMAds NC and PC3 COC with or without orlistat (100µM) or paraoxon-ethyl (100µM) treatment during 24h (n=10). Bars represent mean ± SEM, ns non-significant; *P < 0.05, **P < 0.0 ****P < 0.0001. (A) was analyzed by two-way ANOVA with Sidak’s multiple comparisons test, (B) was analyzed by one-way ANOVA with Tukey’s multiple comparisons test.

**<u>Supplementary figure 3:</u> rBMAds reprogram PCa cells toward a pro-migratory and invasive phenotype.**

**A.** Boyden chamber assay of Du145 NC and COC (n=5). **B.** Boyden chamber assay of LNCaP NC and COC (n=6). **C.** Invasion assay for 24h of Du145 NC and COC (n=5). **D.** Invasion assay for 24h of LNCaP NC and COC (n=4). **E.** Representative images of Du145 NC and COC during 72h and stained with rhodamine-phalloidin (red) for F-Actin, and DAPI (cyan) for nuclei. Scale bar, 20µm. **F**. Number of Du145 NC and COC after 24h, 48h or 72h of culture (n=4) **G.** Number of LNCaP NC and LNCaP COC after 24h, 48h, or 72h of culture (n=5). Bar plot represent mean ± SEM, ns non- significant, *P < 0.05, **P < 0.01; (A-D, G) were analyzed by two-way ANOVA with Sidak’s multiple comparisons test for (A, B) and (F) by Wilcoxon test.

**<u>Supplementary figure 4:</u> Adipocyte-derived FFAs drive PCa cell motility.**

**A.** Boyden chamber assay of Du145 previously treated or not (ctl) with palmitate, oleate, linoleate or the mix of FFA for 72h (n=6). **B**. Boyden chamber assay of LNCaP previously treated or not (ctl) with palmitate, oleate, linoleate or the mix of FFA for 72h (n=7). Bar plot represent mean ± SEM, **P < 0.01, ***P < 0.001, ****P < 0.0001; (A-B) were analyzed by two-way ANOVA with Tukey’s multiple comparisons test.

**<u>Supplementary figure 5:</u> ANGPTL4 as a FFA-responsive driver of PCa cell migration.**

**A.** mRNA expression of ANGPTL4 in Du145 NC and COC, after 72h of culture (n=3). **B.** Intracellular level of ANGPTL4 in Du145 NC and COC, after 72h of culture (n=4). **C.** Quantification of ANGPTL4 contained in the medium of rBMAds cultivated alone (rBMAds NC), Du145 NC and Du145 COC (n=3). **D.** mRNA expression of ANGPTL4 in Du145 after 72h of treatment of exogenous FFA: palmitate, oleate, linoleate or mix of FFA or without treatment (n=5). **E.** mRNA expression of ANGPTL4 in Du145 treated 24h with exogenous FFA in presence or in absence of PPARγ inverse agonist (10µM T0070907) (n=7). **F.** mRNA expression of ANGPTL4 in Du145 NC and COC during 24h in presence or in absence (non-treated, NT) of PPARγ inverse agonist (10µM T0070907) (n=3). **G.** mRNA expression of ANGPTL4 Du145 NC and COC transfected with siRNA control (siCT) or with siRNA against ANGPTL4 (siANGPTL4) (n=4). **H.** Intracellular level of ANGPTL4 in Du145 NC and COC, transfected with siCT or siANGPLT4 and after 24h of culture (n=5). **I.** Quantification of ANGPTL4 contained in the medium of Du145 NC and COC transfected with siCT or siANGPTL4 and after 24h of culture (n=5). **J.** Boyden chamber assay of Du145 NC and COC and transfected with siCT or siANGPTL4 (n=5). **K.** Boyden chamber assay of Du145 NC and COC treated or not during 24h with PPARγ inverse agonist (10µM T0070907). (n=5). **L.** Cytoscape network of ANGPTL4 interactors and interactors associated with the GO term « cell migration ». ANGPTL4 is filled in red, and its neighbours/interactors are framed in red. Cell migration-related major hubs, namely ITGB1 and IL1B, are filled in blue and yellow, respectively. Neighbours/interactors of those two genes are framed accordingly. The two genes that are neighbours of both ITGB1 and IL1B, i.e. JUN and CAV1, are filled in green. Bar plot represent mean ± SEM, ns non-significant, *P < 0.05, **P < 0.01, ***P < 0.001, ****P < 0.0001; (A, B) were analyzed by Student’s t-test, (C) was analyzed by one-way ANOVA with Tukey’s multiple comparisons test, (D-K) were analyzed by two-way ANOVA with Sidak’s multiple comparisons test for (D) and Tukey’s multiple comparisons test for (E-K).

**<u>Supplementary figure 6:</u> ANGPTL4 expression is maintained in other metastatic sites. A.** ANGPTL4 expression in bone, liver, lung and lymph node from the *Metastatic Prostate Adenocarcinoma* (*SU*2*C*/*PCF Dream Team*^38^, bone n=73, liver n=39, lung n=6, lymph node n=115), GSE118435 (bone n=6, liver n=14, lung n=6, lymph node n=10) and GSE126078 (bone n=9, liver n=60, lung n=10, lymph node n=28) datasets.

## Acknowledgements

This study was supported by the French National Cancer Institute (INCA PLBio 2020-2024) and the *Ligue Nationale contre le Cancer* (équipe labélisée). Marine Hernandez received a PhD fellowship from French ministry of higher education and research and from the *Ligue Nationale contre le Cancer*. Sauyeun Shin received a post-doctoral fellowship from INCa PLBio (2020-28). Alessandro Taccini received a European PhD fellowship (MSCA-2022-DN-01; PROSTAMET # 101120283). This work benefited from the TRI (*Toulouse Réseau Imagerie*) imaging facility, member of the national infrastructure France-BioImaging supported by the French National Research Agency (ANR-10-INBS-04). Lipidomic analysis were performed at MetaToul (Toulouse metabolomics & fluxomics facilities, www.mth-metatoul.com on the site dedicated to Lipids Analysis Inserm I2MC) and was supported by the MetaboHUB infrastructure funded by the Agence Nationale de la Recherche under the France 2030 program (MetaboHUB ANR-11-INBS-0010; MetEx+ ANR-21-ESRE-0035; MetaboHUB (JVCE) ANR-24-INBS-0012). We thank Carine Valle from Genomics and Transcriptomics platform and Clementine Decamps from Bioinformatics platform, Technological cluster of the Cancer Research Center of Toulouse (INSERM-UMR1037) for the analysis of the transcriptional study. We are grateful to the genotoul bioinformatics platform Toulouse Occitanie (<u>Bioinfo Genotoul,</u> https://doi.org/10.15454/1.5572369328961167E12) for providing computing resources used for alignment. We thank Dr Guillaume Bossis IGMM, Montpellier, France for providing leukemia cell lines U937 control and knock-out (KO) for CD36.

## Author contributions

N.R set up the conditions for harvesting BMAT and SCAT in close collaboration with CA and CM and supervised the samples collection. MM handled the BMAT and SCAT. MH performed most of the experiments with the help of SS and AT. CV performed RNA-seq experiment. DR analyzed RNAseq data and prepared the related figures. SD performed and prepared the figures of the imaging experiments. JBM and NG conducted lipidomic analysis. MH, CM and CA wrote the manuscript. CA and CM conceived the idea for this project and CA supervised the study.

## Data availability

RNA sequencing data generated in this publication (PC3 cocultivated or not with rBMAds) have been deposited in NCBI’s Gene Expression Omnibus and are accessible through GEO Series accession number GSE312982 (https://www.ncbi.nlm.nih.gov/geo/query/acc.cgi?acc=GSE312982).

