## Supplementary figures and images for "Human bone marrow adipocytes drive prostate cancer bone metastasis progression via lipid-mediated induction of Angiopoietin-like 4"

### Figure S1

**Supplementary figure 1:**

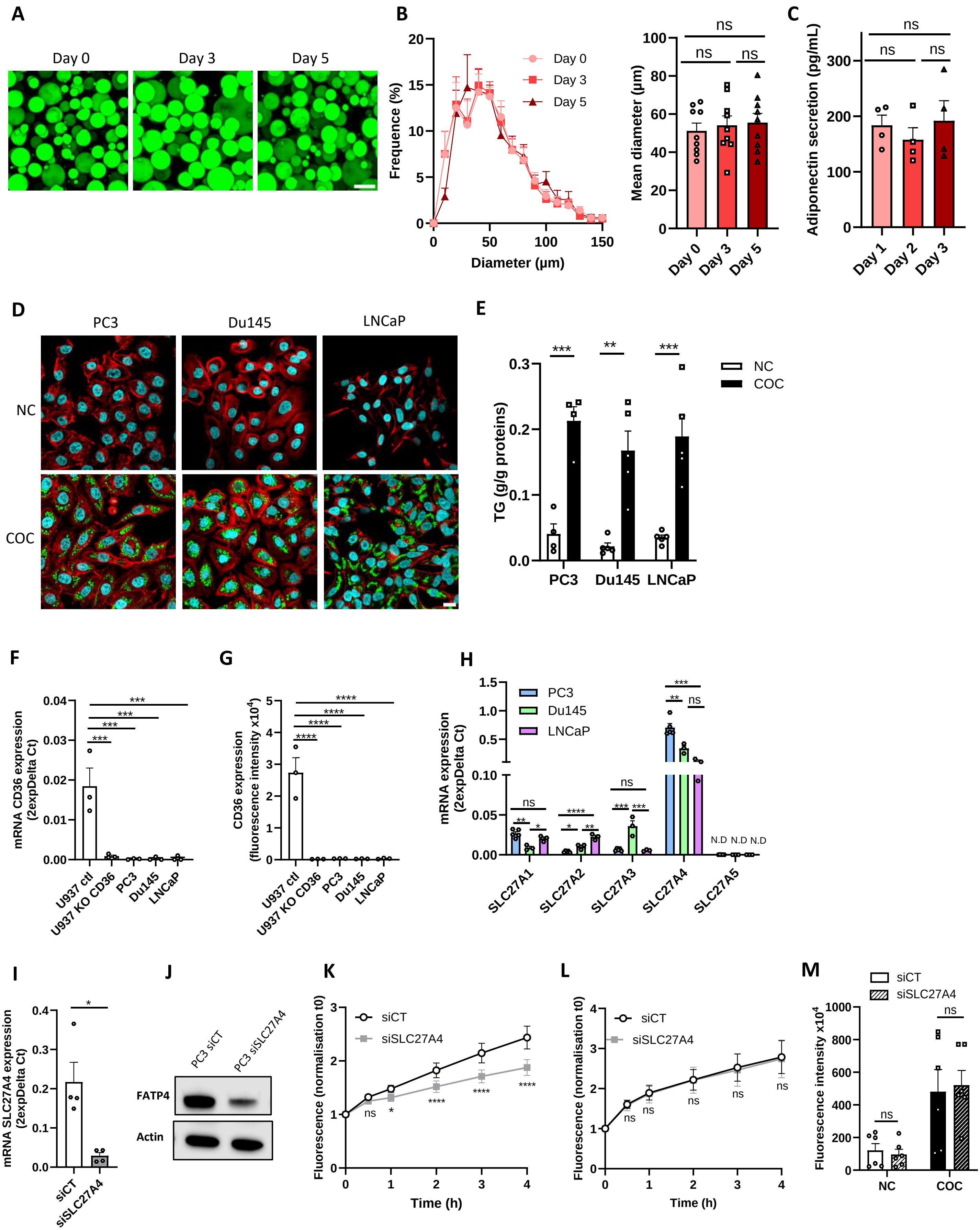

### Figure S2

#### Supplementary figure 2:

**A**

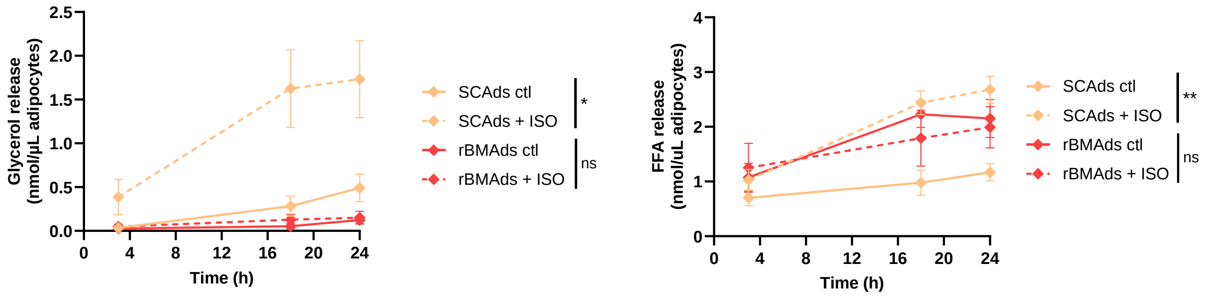


**B**

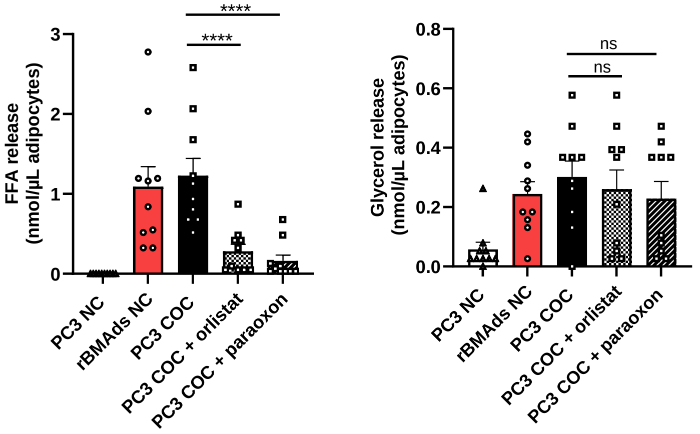

### Figure S3

#### Supplementary figure 3:

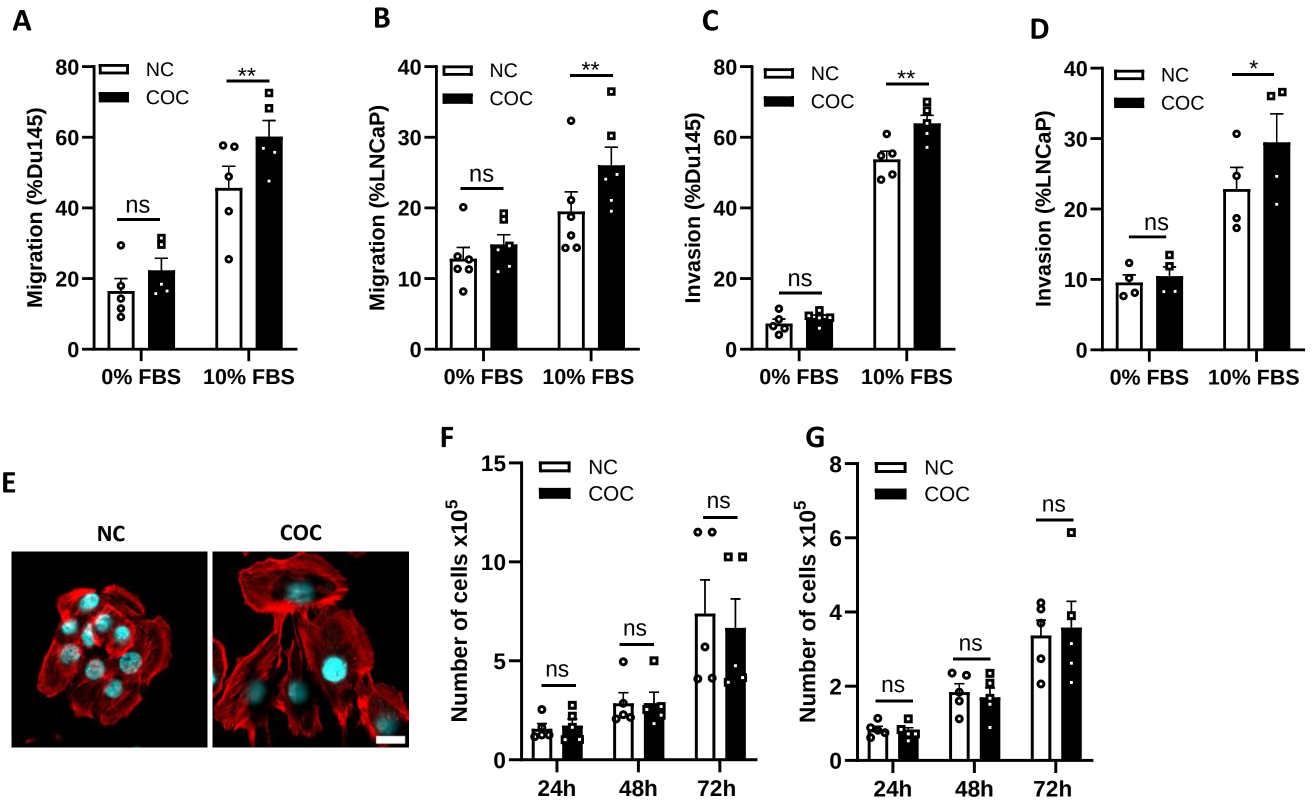

### Figure S4

**Supplementary figure 4 :**

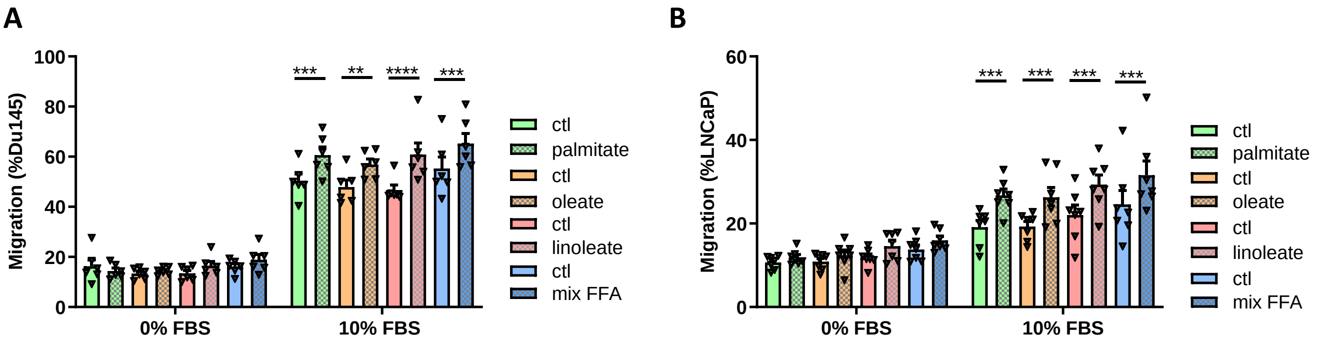

### Figure S5

#### Supplementary figure 5

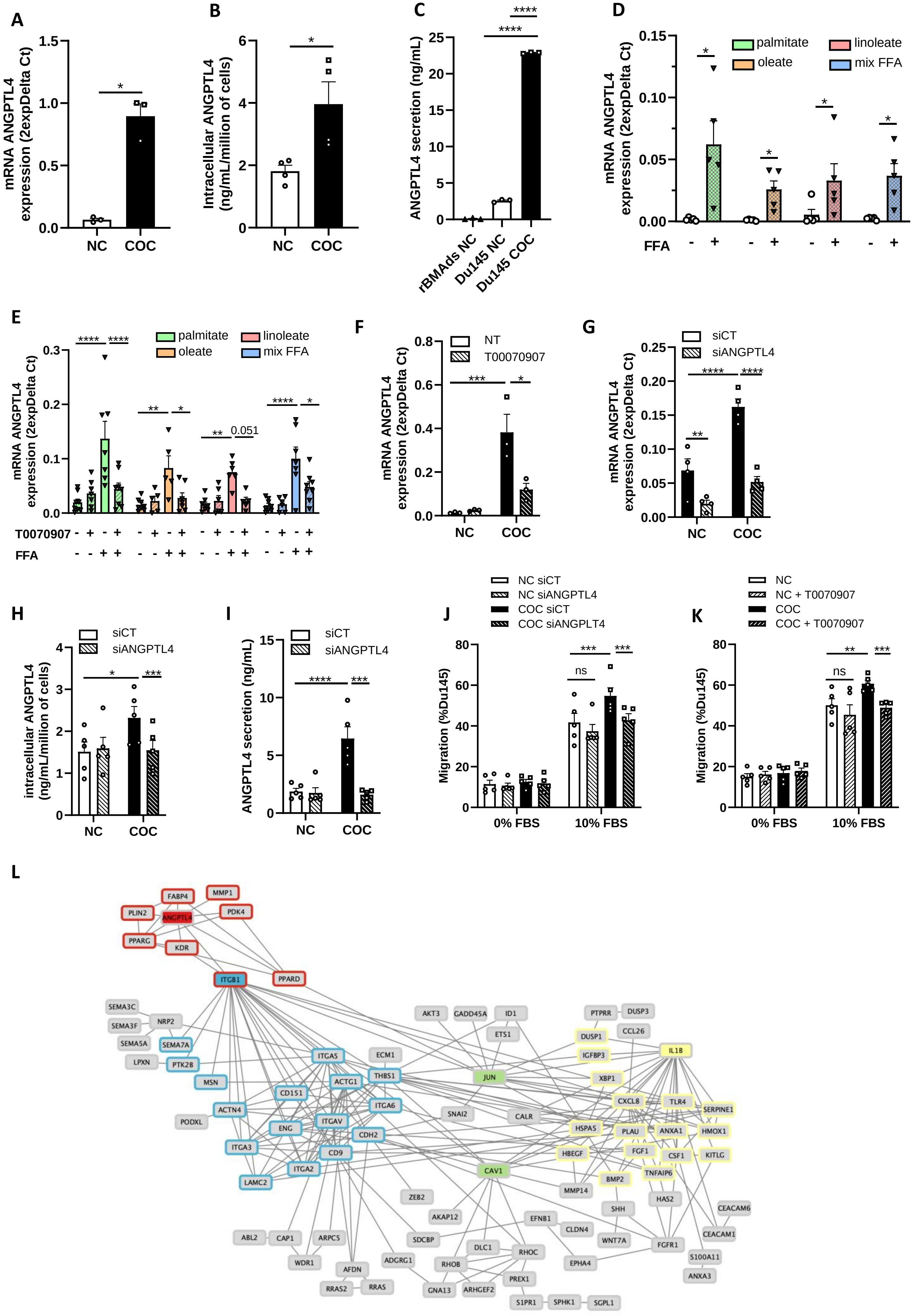

### Figure S6

#### Supplementary figure 6 :

**A**

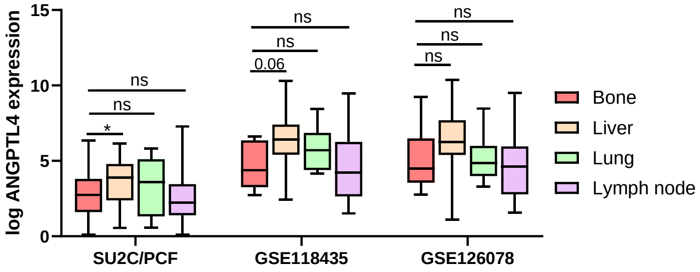
