## Supplementary material for "Human bone marrow adipocytes drive prostate cancer bone metastasis progression via lipid-mediated induction of Angiopoietin-like 4": Table S1

| Figure | #patient | Gender | Age | BMI | experiments |
|---|---|---|---|---|---|
| **Figure 1A** | 596 | M | 67 | 25.64 | microscopy PC3 |
| **Figure 1B** | 450 | M | 71 | 29.39 | neutral lipid quantification (lipidomic) |
| | 452 | M | 68 | 25.25 | neutral lipid quantification (lipidomic) |
| | 455 | M | 83 | 24.21 | neutral lipid quantification (lipidomic) |
| | 623 | M | 48 | 22.49 | neutral lipid quantification (lipidomic) |
| | 641 | M | 56 | 23.67 | neutral lipid quantification (lipidomic) |
| | 655 | M | 68 | 21.88 | neutral lipid quantification (lipidomic) |
| | 681 | M | 37 | 24.84 | neutral lipid quantification (lipidomic) |
| | 686 | M | 69 | 26.73 | neutral lipid quantification (lipidomic) |
| | 718 | M | 88 | 27.18 | neutral lipid quantification (lipidomic) |
| **Figure 1C** | 544 | W | 79 | 24.97 | microscopy transfert Bodipy FLC16 |
| **Figure 1D** | 570 | M | 52 | 28.37 | quantification of Bodipy FLC16 uptake (cytometry) |
| | 571 | M | 88 | 23.51 | quantification of Bodipy FLC16 uptake (cytometry) |
| | 585 | M | 80 | 26.61 | quantification of Bodipy FLC16 uptake (cytometry) |
| **Figure 1E + 1F** | 450 | M | 71 | 29.39 | FFA species (lipidomic) |
| | 452 | M | 68 | 25.25 | FFA species (lipidomic) |
| | 455 | M | 83 | 24.21 | FFA species (lipidomic) |
| | 463 | M | 73 | 26.30 | FFA species (lipidomic) |
| **Figure 2B** | 555 | W | 68 | 21.64 | lipolysis |
| | 573 | M | 63 | 22.72 | lipolysis |
| | 578 | W | 70 | 24.57 | lipolysis |
| | 619 | W | 65 | 24.17 | lipolysis |
| | 625 | W | 63 | 30.47 | lipolysis |
| | 629 | W | 67 | 19.95 | lipolysis |
| | 637 | M | 77 | 24.69 | lipolysis |
| | 638 | W | 76 | 30.30 | lipolysis |
| | 639 | W | 68 | 26.77 | lipolysis |
| | 814 | W | 59 | 25.47 | lipolysis |
| **Figure 2C + 2D** | 635 | W | 38 | 30.80 | FFA and glycerol quantification |
| | 636 | W | 70 | 28.89 | FFA and glycerol quantification |
| | 639 | W | 68 | 26.77 | FFA and glycerol quantification |
| | 642 | M | 58 | 29.41 | FFA and glycerol quantification |
| | 968 | W | 80 | 23.74 | FFA and glycerol quantification |
| | 999 | M | 58 | 26.30 | FFA and glycerol quantification |
| | 1015 | M | 44 | 22.84 | FFA and glycerol quantification |
| **Figure 2E** | 962 | W | 90 | 22.83 | FFA and glycerol quantification (paraoxon ethyl. orlistat) |
| | 968 | W | 80 | 23.74 | FFA and glycerol quantification (paraoxon ethyl. orlistat) |
| | 986 | W | 55 | 27.69 | FFA and glycerol quantification (paraoxon ethyl. orlistat) |
| | 1000 | W | 80 | 21.48 | FFA and glycerol quantification (paraoxon ethyl. orlistat) |
| | 1005 | M | 72 | 27.77 | FFA and glycerol quantification (paraoxon ethyl. orlistat) |
| | 1010 | W | 76 | 21.67 | FFA and glycerol quantification (paraoxon ethyl. orlistat) |
| | 1011 | M | 84 | 26.30 | FFA and glycerol quantification (paraoxon ethyl. orlistat) |
| **Figure 3A-B-C** | 593 | M | 68 | 21.78 | RNAseq |
| | 596 | M | 67 | 25.64 | RNAseq |
| | 598 | M | 60 | 24.73 | RNAseq |
| | 599 | M | 75 | 25.91 | RNAseq |
| | 604 | M | 72 | 23.31 | RNAseq |
| **Figure 3D** | 897 | M | 70 | 24.16 | Migration Boyden chamber PC3 |
| | 898 | M | 70 | 24.09 | Migration Boyden chamber PC3 |
| | 900 | M | 80 | 26.37 | Migration Boyden chamber PC3 |
| | 908 | M | 64 | 22.99 | Migration Boyden chamber PC3 |
| | 909 | M | 70 | 25.82 | Migration Boyden chamber PC3 |
| **Figure 3E** | 533 | M | 76 | 23.88 | Scratch test PC3 |
| | 547 | M | 69 | 23.74 | Scratch test PC3 |
| | 550 | M | 85 | 27.18 | Scratch test PC3 |
| | 551 | M | 67 | 26.88 | Scratch test PC3 |
| **Figure 3F** | 1149 | M | 86 | 27.40 | Invasion assay PC3 |
| | 1155 | M | 71 | 23.99 | Invasion assay PC3 |
| | 1160 | M | 68 | 22.96 | Invasion assay PC3 |
| | 1161 | M | 62 | 26.51 | Invasion assay PC3 |
| | 1165 | M | 67 | 22.15 | Invasion assay PC3 |
| **Figure 3G** | 596 | M | 67 | 25.64 | microscopy PC3 |
| **Figure 3H** | 581 | M | 79 | 24.77 | PC3 numbers |
| | 582 | M | 64 | 28.41 | PC3 numbers |
| | 583 | M | 92 | 27.1 | PC3 numbers |
| | 584 | M | 36 | 22.89 | PC3 numbers |
| **Figure 3I** | 1203 | M | 52 | 24.62 | BrdU PC3 |
| | 1208 | M | 71 | 27.25 | BrdU PC3 |
| | 1215 | M | 77 | 29.4 | BrdU PC3 |

| | | | | | |
|---|---|---|---|---|---|
| | 956 | M | 71 | 20.76 | FFA quantification delipidated CM |
| | 957 | W | 61 | 34.15 | FFA quantification delipidated CM |
| | 958 | M | 65 | 23.16 | FFA quantification delipidated CM |
| | 992 | M | 54 | 24.69 | FFA quantification delipidated CM |
| | 993 | M | 67 | 25.59 | FFA quantification delipidated CM |
| **Figure 4A** | 999 | M | 58 | 26.30 | FFA quantification delipidated CM |
| | 956 | M | 71 | 20.76 | TG quantification delpidated CM |
| | 958 | M | 65 | 23.16 | TG quantification delpidated CM |
| | 999 | M | 58 | 26.30 | TG quantification delpidated CM |
| | 1005 | M | 72 | 27.77 | TG quantification delpidated CM |
| | 1006 | M | 83 | 25.38 | TG quantification delpidated CM |
| **Figure 4B-C** | 1008 | M | 75 | 25.65 | TG quantification delpidated CM |
| | 956 | M | 71 | 20.76 | Migration Boyden chamber treatment delipidated CM |
| | 957 | W | 61 | 34.15 | Migration Boyden chamber treatment delipidated CM |
| | 958 | M | 65 | 23.16 | Migration Boyden chamber treatment delipidated CM |
| | 992 | M | 54 | 24.69 | Migration Boyden chamber treatment delipidated CM |
| | 993 | M | 67 | 25.59 | Migration Boyden chamber treatment delipidated CM |
| **Figure 4D** | 999 | M | 58 | 26.30 | Migration Boyden chamber treatment delipidated CM |
| | 593 | M | 68 | 21.78 | RNAseq |
| | 596 | M | 67 | 25.64 | RNAseq |
| | 598 | M | 60 | 24.73 | RNAseq |
| | 599 | M | 75 | 25.91 | RNAseq |
| **Figure 5A** | 604 | M | 72 | 23.31 | RNAseq |
| | 958 | M | 65 | 23.16 | mRNA quantification ANGPTL4 in PC3 |
| | 960 | M | 68 | 24.82 | mRNA quantification ANGPTL4 in PC3 |
| **Figure 5B** | 961 | M | 80 | 23.62 | mRNA quantification ANGPTL4 in PC3 |
| | 1067 | M | 46 | 24.69 | intracellular ANGPTL4 in PC3 |
| | 1077 | M | 64 | 24.76 | intracellular ANGPTL4 in PC3 |
| | 1079 | M | 68 | 28.23 | intracellular ANGPTL4 in PC3 |
| **Figure 5C** | 1082 | M | 54 | 18.08 | intracellular ANGPTL4 in PC3 |
| | 1052 | M | 43 | 27.77 | ANGPTL4 secretion PC3 |
| | 1053 | M | 54 | 24.69 | ANGPTL4 secretion PC3 |
| | 1055 | M | 71 | 25.35 | ANGPTL4 secretion PC3 |
| **Figure 5D** | 1056 | M | 86 | 27.28 | ANGPTL4 secretion PC3 |
| | 1086 | M | 38 | 22.84 | mRNA ANGPTL4 expression with inverse agonist PPARg treatment PC3 |
| | 1090 | M | 78 | 26.83 | mRNA ANGPTL4 expression with inverse agonist PPARg treatment PC3 |
| | 1095 | M | 68 | 23.36 | mRNA ANGPTL4 expression with inverse agonist PPARg treatment PC3 |
| | 1102 | M | 74 | 27.73 | mRNA ANGPTL4 expression with inverse agonist PPARg treatment PC3 |
| | 1105 | M | 59 | 25.73 | mRNA ANGPTL4 expression with inverse agonist PPARg treatment PC3 |
| **Figure 5G** | 1107 | M | 74 | 27.31 | mRNA ANGPTL4 expression with inverse agonist PPARg treatment PC3 |
| | 955 | M | 59 | 24.31 | mRNA quantification ANGPTL4 siANGPTL4 treatment PC3 |
| | 967 | M | 44 | 21.55 | mRNA quantification ANGPTL4 siANGPTL4 treatment PC3 |
| **Figure 5H** | 970 | M | 75 | 26.22 | mRNA quantification ANGPTL4 siANGPTL4 treatment PC3 |
| | 1127 | M | 68 | 17.86 | intracellular and secretion of ANGPTL4 after siANGPTL4 PC3 |
| | 1133 | M | 67 | 26.49 | intracellular and secretion of ANGPTL4 after siANGPTL4 PC3 |
| | 1136 | M | 63 | 25.31 | intracellular and secretion of ANGPTL4 after siANGPTL4 PC3 |
| | 1139 | M | 69 | 28.70 | intracellular and secretion of ANGPTL4 after siANGPTL4 PC3 |
| | 1143 | M | 55 | 26.83 | intracellular and secretion of ANGPTL4 after siANGPTL4 PC3 |
| **Figure 5I-J** | 1145 | M | 79 | 26.78 | intracellular and secretion of ANGPTL4 after siANGPTL4 PC3 |
| | 955 | M | 59 | 24.31 | Migration Boyden chamber after siANGPTL4 PC3 |
| | 967 | M | 44 | 21.55 | Migration Boyden chamber after siANGPTL4 PC3 |
| | 969 | M | 70 | 25.03 | Migration Boyden chamber after siANGPTL4 PC3 |
| | 970 | M | 75 | 26.22 | Migration Boyden chamber after siANGPTL4 PC3 |
| **Figure 5K** | 971 | M | 31 | 26.67 | Migration Boyden chamber after siANGPTL4 PC3 |
| | 1024 | M | 71 | 25.88 | Scratch test PC3 after siANGPTL4 |
| | 1025 | M | 86 | 19.81 | Scratch test PC3 after siANGPTL4 |
| | 1030 | M | 57 | 27.47 | Scratch test PC3 after siANGPTL4 |
| | 1041 | M | 78 | 26.73 | Scratch test PC3 after siANGPTL4 |
| **Figure 5L** | 1043 | M | 52 | 27.17 | Scratch test PC3 after siANGPTL4 |
| | 1186 | M | 58 | 24.42 | Invasion assay PC3 after siANGPTL4 |
| | 1185 | M | 64 | 22.76 | Invasion assay PC3 after siANGPTL4 |
| | 1189 | M | 91 | 20.34 | Invasion assay PC3 after siANGPTL4 |
| **Figure 5M** | 1190 | M | 61 | 26.31 | Invasion assay PC3 after siANGPTL4 |
| | 1133 | M | 67 | 26.49 | Migration Boyden chamber after inverse agonsit PPARg treatment |
| | 1135 | M | 72 | 25.46 | Migration Boyden chamber after inverse agonsit PPARg treatment |
| | 1136 | M | 63 | 25.31 | Migration Boyden chamber after inverse agonsit PPARg treatment |
| **Figure 5N** | 1139 | M | 69 | 28.70 | Migration Boyden chamber after inverse agonsit PPARg treatment |
| | AB809 (periprostatic adipocytes) | M | 66 | 24.27 | mRNA quantification ANGPTL4 PC3 |
| | VJ810 (periprostatic adipocytes) | M | 53 | 25.06 | mRNA quantification ANGPTL4 PC3 |
| | CJ811 (periprostatic adipocytes) | M | 61 | 24.44 | mRNA quantification ANGPTL4 PC3 |

| | | | | | |
|---|---|---|---|---|---|
| | MB912 (periprostatic adipocytes) | M | 68 | 27.17 | mRNA quantification ANGPTL4 PC3 |
| | HG 913 (periprostatic adipocytes) | M | 74 | 26.17 | mRNA quantification ANGPTL4 PC3 |
| | 1304 | M | 56 | 22.55 | mRNA quantification ANGPTL4 PC3 |
| | 1328 | M | 42 | 27.1 | mRNA quantification ANGPTL4 PC3 |
| | 958 | M | 65 | 23.16 | mRNA quantification ANGPTL4 PC3 |
| | 960 | M | 68 | 24.82 | mRNA quantification ANGPTL4 PC3 |
| **Figure 6A** | 961 | M | 80 | 23.62 | mRNA quantification ANGPTL4 PC3 |
| | 759 | W | 66 | 27.10 | microscopy rBMAds |
| | 811 | M | 57 | 24.72 | microscopy rBMAds |
| **Supplementary Fig.1A** | 808 | M | 69 | 25.10 | microscopy rBMAds |
| | 807 | M | 70 | 24.15 | size rBMAds |
| | 808 | M | 69 | 25.10 | size rBMAds |
| | 809 | W | 80 | 22.31 | size rBMAds |
| | 822 | W | 62 | 18.73 | size rBMAds |
| | 827 | W | 58 | 28.71 | size rBMAds |
| | 830 | M | 71 | 24.86 | size rBMAds |
| | 881 | W | 72 | 19.83 | size rBMAds |
| | 882 | M | 72 | 21.48 | size rBMAds |
| **Supplementary Fig.1B** | 883 | W | 80 | 24.01 | size rBMAds |
| | 1128 | W | 80 | 25.71 | adiponectin secretion |
| | 1129 | W | 75 | 28.70 | adiponectin secretion |
| | 1131 | W | 56 | 26.35 | adiponectin secretion |
| **Supplementary Fig.1C** | 1137 | W | 73 | 21.48 | adiponectin secretion |
| | 655 | M | 68 | 21.88 | microscopy Du145 |
| | 700 | M | 44 | 27.17 | microscopy LNCaP |
| **Supplementary Fig. 1D** | 547 | M | 69 | 23.74 | microscopy PC3 |
| | 790 | M | 72 | 27.45 | TG quantification PC3 |
| | 839 | M | 46 | 28.39 | TG quantification PC3 |
| | 923 | M | 71 | 26.53 | TG quantification Du145 + LNCaP |
| | 958 | M | 65 | 23.16 | TG quantification PC3 |
| | 960 | M | 68 | 24.82 | TG quantification PC3 + Du145 |
| | 961 | M | 80 | 23.62 | TG quantification Du145 |
| | 950 | M | 77 | 27.17 | TG quantification Du145 + LNCaP |
| | 1005 | M | 72 | 27.77 | TG quantification LNCaP |
| | 1018 | M | 66 | 24.34 | TG quantification LNCaP |
| **Supplementary Fig. 1E** | 1019 | M | 54 | 22.71 | TG quantification LNCaP |
| | 1349 | M | 69 | 27.68 | lipid accumulation in PC3 siSLC27A4 |
| | 1366 | M | 43 | 19.87 | lipid accumulation in PC3 siSLC27A4 |
| | 1367 | M | 66 | 24.44 | lipid accumulation in PC3 siSLC27A4 |
| | 1370 | M | 25 | 21.26 | lipid accumulation in PC3 siSLC27A4 |
| | 1371 | M | 57 | 25.14 | lipid accumulation in PC3 siSLC27A4 |
| **Supplementary Fig. 1M** | 1372 | M | 56 | 30.86 | lipid accumulation in PC3 siSLC27A4 |
| | 1360 | W | 81 | 25.71 | long lipolysis |
| | 1363 | M | 64 | 20.76 | long lipolysis |
| | 1364 | W | 63 | 23.43 | long lipolysis |
| **Supplementary Fig.2A** | 1365 | W | 63 | 21.97 | long lipolysis |
| | 1347 | W | 98 | 27.34 | FFA and glycerol quantification in coculture (paraoxon ethyl. orlistat) |
| | 1353 | M | 47 | 24.96 | FFA and glycerol quantification in coculture (paraoxon ethyl. orlistat) |
| | 1358 | M | 62 | 24.66 | FFA and glycerol quantification in coculture (paraoxon ethyl. orlistat) |
| | 1213 | M | 78 | 24.93 | FFA and glycerol quantification in coculture (paraoxon ethyl. orlistat) |
| | 1214 | M | 55 | 24.69 | FFA and glycerol quantification in coculture (paraoxon ethyl. orlistat) |
| | 1220 | M | 61 | 25.62 | FFA and glycerol quantification in coculture (paraoxon ethyl. orlistat) |
| | 1221 | M | 51 | 24.77 | FFA and glycerol quantification in coculture (paraoxon ethyl. orlistat) |
| | 1222 | M | 58 | 27.73 | FFA and glycerol quantification in coculture (paraoxon ethyl. orlistat) |
| | 1223 | M | 62 | 22.46 | FFA and glycerol quantification in coculture (paraoxon ethyl. orlistat) |
| **Supplementary Fig.2B** | 1363 | M | 64 | 20.76 | FFA and glycerol quantification in coculture (paraoxon ethyl. orlistat) |
| | 653 | M | 79 | 26.22 | Migration Boyden chamber Du145 |
| | 655 | M | 68 | 21.88 | Migration Boyden chamber Du145 |
| | 681 | M | 37 | 24.84 | Migration Boyden chamber Du145 |
| | 900 | M | 80 | 26.37 | Migration Boyden chamber Du145 |
| **Supplementary Fig. 3A** | 909 | M | 70 | 25.82 | Migration Boyden chamber Du145 |
| | 671 | M | 88 | 25.51 | Migration Boyden chamber LNCaP |
| | 677 | M | 65 | 24 | Migration Boyden chamber LNCaP |
| | 681 | M | 37 | 24.84 | Migration Boyden chamber LNCaP |
| | 683 | M | 83 | 23.38 | Migration Boyden chamber LNCaP |
| | 900 | M | 80 | 26.37 | Migration Boyden chamber LNCaP |
| **Supplementary Fig. 3B** | 909 | M | 70 | 25.82 | Migration Boyden chamber LNCaP |
| | 1161 | M | 62 | 26.51 | Invasion assay Du145 |
| | 1173 | M | 96 | 27.68 | Invasion assay Du145 |
| | 1174 | M | 73 | 26.30 | Invasion assay Du145 |

| | | | | | |
|---|---|---|---|---|---|
| | 1175 | M | 65 | 22.95 | Invasion assay Du145 |
| **Supplementary Fig. 3C** | 1176 | M | 63 | 27.64 | Invasion assay Du145 |
| | 1290 | M | 37 | 25.11 | Invasion assay LNCaP |
| | 1298 | M | 77 | 26.22 | Invasion assay LNCaP |
| | 1299 | M | 64 | 26.74 | Invasion assay LNCaP |
| **Supplementary Fig. 3D** | 1301 | M | 40 | 28.08 | Invasion assay LNCaP |
| **Supplementary Fig. 3E** | 655 | M | 68 | 21.88 | microscopy Du145 |
| | 662 | M | 57 | 24.84 | Du145 numbers |
| | 670 | M | 70 | 26.42 | Du145 numbers |
| | 672 | M | 49 | 29.83 | Du145 numbers |
| | 937 | M | 52 | 23.37 | Du145 numbers |
| **Supplementary Fig.3F** | 1153 | M | 74 | 22.76 | Du145 numbers |
| | 1153 | M | 74 | 22.76 | LNCaP numbers |
| | 1167 | M | 39 | 21.15 | LNCaP numbers |
| | 1215 | M | 77 | 29. 4 | LNCaP numbers |
| | 1217 | M | 86 | 23. 94 | LNCaP numbers |
| **Supplementary Fig.3G** | 1224 | M | 48 | 24. 62 | LNCaP numbers |
| | 958 | M | 65 | 23.16 | mRNA quantification ANGPTL4 Du145 |
| | 960 | M | 68 | 24.82 | mRNA quantification ANGPTL4 Du145 |
| **Supplementary Fig. 5A** | 961 | M | 80 | 23.62 | mRNA quantification ANGPTL4 Du145 |
| | 1082 | M | 54 | 18.08 | intracellular ANGPTL4 in Du145 |
| | 1084 | M | 75 | 21.50 | intracellular ANGPTL4 in Du145 |
| | 1086 | M | 38 | 22.84 | intracellular ANGPTL4 in Du145 |
| **Supplementary Fig. 5B** | 1088 | M | 65 | 21.91 | intracellular ANGPTL4 in Du145 |
| | 1059 | M | 70 | 27.77 | ANGPTL4 secretion PC3 and Du145 |
| | 1061 | M | 76 | 24.91 | ANGPTL4 secretion PC3 and Du145 |
| **Supplementary Fig. 5C** | 1063 | M | 60 | 23.12 | ANGPTL4 secretion PC3 and Du145 |
| | 1086 | M | 38 | 22.84 | mRNA ANGPTL4 expression with inverse agonist PPARg treatment Du145 |
| | 1090 | M | 78 | 26.83 | mRNA ANGPTL4 expression with inverse agonist PPARg treatment Du145 |
| **Supplementary Fig. 5F** | 1105 | M | 59 | 25.73 | mRNA ANGPTL4 expression with inverse agonist PPARg treatment Du145 |
| | 969 | M | 70 | 25.03 | mRNA quantification ANGPTL4 siANGPTL4 treatment Du145 |
| | 971 | M | 31 | 26.67 | mRNA quantification ANGPTL4 siANGPTL4 treatment Du145 |
| | 1215 | M | 88 | 29. 4 | mRNA quantification ANGPTL4 siANGPTL4 treatment Du145 |
| | 1217 | M | 86 | 23. 94 | mRNA quantification ANGPTL4 siANGPTL4 treatment Du145 |
| **Supplementary Fig. 5G** | 1219 | M | 54 | 20. 78 | mRNA quantification ANGPTL4 siANGPTL4 treatment Du145 |
| | 1127 | M | 68 | 17.86 | intracellular and secretion of ANGPTL4 after siANGPTL4 Du145 |
| | 1133 | M | 67 | 26.49 | intracellular and secretion of ANGPTL4 after siANGPTL4 Du145 |
| | 1136 | M | 63 | 25.31 | intracellular and secretion of ANGPTL4 after siANGPTL4 Du145 |
| | 1139 | M | 69 | 28.70 | intracellular and secretion of ANGPTL4 after siANGPTL4 Du145 |
| **Supplementary Fig. 5H-I** | 1143 | M | 55 | 26.83 | intracellular and secretion of ANGPTL4 after siANGPTL4 Du145 |
| | 967 | M | 44 | 21.55 | Migration Boyden chamber after siANGPTL4 Du145 |
| | 969 | M | 70 | 25.03 | Migration Boyden chamber after siANGPTL4 Du145 |
| | 970 | M | 75 | 26.22 | Migration Boyden chamber after siANGPTL4 Du145 |
| | 971 | M | 31 | 26.67 | Migration Boyden chamber after siANGPTL4 Du145 |
| **Supplementary Fig. 5J** | 983 | M | 50 | 25.77 | Migration Boyden chamber after siANGPTL4 Du145 |
| | 1143 | M | 55 | 26.83 | Migration Boyden chamber after inverse agonsit PPARg treatment |
| | 1145 | M | 79 | 26.78 | Migration Boyden chamber after inverse agonsit PPARg treatment |
| | 1147 | M | 72 | 27.15 | Migration Boyden chamber after inverse agonsit PPARg treatment |
| | 1148 | M | 72 | 19.57 | Migration Boyden chamber after inverse agonsit PPARg treatment |
| **Supplementary Fig. 5K** | 1149 | M | 86 | 27.40 | Migration Boyden chamber after inverse agonsit PPARg treatment |
